# RIGATonI: An R software for Rapid Identification of Genomic Alterations in Tumors affecting lymphocyte Infiltration

**DOI:** 10.1101/2024.03.02.583103

**Authors:** Raven Vella, Emily L. Hoskins, Lianbo Yu, Julie W. Reeser, Michele R. Wing, Eric Samorodnitsky, Leah Stein, Elizabeth G. Bruening, Anoosha Paruchuri, Michelle Churchman, Nancy Single, Wei Chen, Aharon G. Freud, Sameek Roychowdhury

## Abstract

Tumor genomic alterations have been associated with altered tumor immune microenvironments and therapeutic outcomes. These studies raise a critical question: are there additional genomic variations altering the immune microenvironment in tumors that can provide insight into mechanisms of immune evasion? This question is the backbone of precision immuno-oncology. Current computational approaches to estimate immunity in bulk RNA sequencing (RNAseq) from tumors include gene set enrichment analysis and cellular deconvolution, but these techniques do not consider the spatial organization of lymphocytes or connect immune phenotypes with gene activity. Our new software package, Rapid Identification of Genomic Alterations in Tumors affecting lymphocyte Infiltration (RIGATonI), addresses these two gaps in separate modules: the Immunity Module and the Function Module. Using pathologist-reviewed histology slides and paired bulk RNAseq expression data, we trained a machine learning algorithm to detect high, medium, and low levels of immune infiltration (Immunity Module). We validated this technique using a subset of pathologist-reviewed slides not included in the training data, multiplex immunohistochemistry, flow cytometry, and digital staining of The Cancer Genome Atlas (TCGA). In addition to immune infiltrate classification, RIGATonI leverages another novel machine learning algorithm for the prediction of gain- and loss-of-function genomic alterations (Function Module). We validated this approach using clinically relevant and function-impacting genomic alterations from the OncoKB database. Combining these two modules, we analyzed all genomic alterations present in solid tumors in TCGA for their resulting protein function and immune phenotype. We visualized these results on a publicly available website. To illustrate RIGATonI’s potential to identify novel genomic variants with associated altered immune phenotypes, we describe increased anti-tumor immunity in renal cell carcinoma tumors harboring 14q deletions and confirmed these results with previously published single-cell RNA sequencing. Thus, we present our R package and online database, RIGATonI: an innovative software for precision immuno-oncology research.

---

Immune surveillance is crucial for the eradication of cancer cells^1^. Investigating factors impacting the tumor immune microenvironment can aid in understanding this process^1^. The tumor microenvironment is composed of tumor cells, along with surrounding immune cells, stroma cells and tissue matrix, and microbiota^1^. Each of these elements vary across patients and tumor types, producing a spectrum of immune phenotypes and, consequently, clinical outcomes to immunotherapy^1^. In particular, the field of precision immuno-oncology requires approaches designed to discover relationships between tumor genomic alterations and their associated microenvironments^2, 3^. There are several techniques to assess the tumor immune microenvironment using bulk tumor RNAseq data; however, there are no approaches designed to rapidly detect associations between tumor genomic alterations and the quality of tumor inflammation in big data repositories. Thus, there is a need to develop computational tools specifically designed to assess immunity in large databases of tumor transcriptomes.

Current methods for detecting altered immunity in tumors (**Figure 1A**) include gene set enrichment analyses (e.g., ImSig^4^, Thorsson *et. al.*^5^), cellular deconvolution techniques (e.g., MCP-counter^6^, quantiseqR^7^, CIBERSORT^8^), and ensemble results provided in databases (e.g., TCIA^9^, TIMER2.0^10^, TIMEDB^11^). Gene set enrichment analysis involves assessing the expression of immune-related gene sets and then clustering cancers based on the results. This is routinely employed in large studies of immunotherapy outcomes to reveal the tumor features that underscore interpatient differences^12, 13^. These approaches are often not optimized for application to data outside the study within which they are built because they are not intended for robust analysis across databases. Alternatively, cellular deconvolution tools attempt to emulate flow cytometry or immunohistochemistry by estimating the number of specific immune cells in a sample^6–8^. Importantly, cellular deconvolution does not make clear distinctions between immune phenotypes broadly and rather leaves interpretation up to the user. Databases of cellular deconvolution results have emerged, which include ensemble analyses across The Cancer Genome Atlas (TCGA) and other sources^10, 11^. These databases provide information about associations between genomic alterations and immune phenotypes; however, they do not separate analyses based on the functional status of the genomic alteration instead combining all alterations in a gene of interest regardless of their molecular impact^9–11^. None of these techniques were specifically developed to classify immune phenotypes and discern associations with functionally relevant tumor genomic alterations.

**Figure 1.**
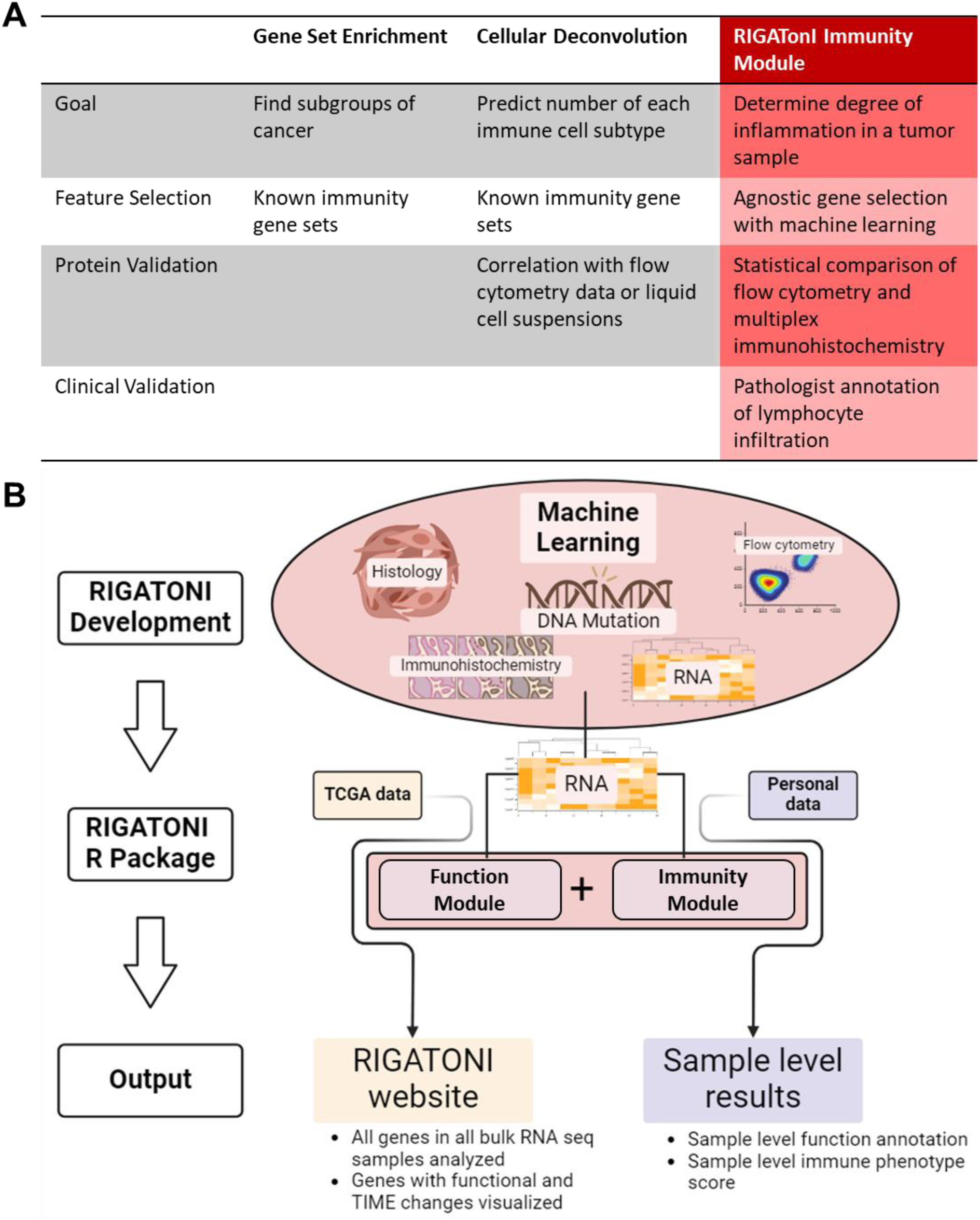
Overview of RIGATonI as a novel agnostic approach to measure immune infiltration from bulk tumor RNASeq. **A.** Current approaches to estimate immunity from bulk tumor RNAseq include gene set enrichment^4, 5^ and cellular deconvolution^6–8^ based methods. In contrast, RIGATonI utilizes a gene agnostic approach to classify immune cell infiltration in tumors by training on pathologist-annotated digital slides. **B.** RIGATonI enables identification of candidate genomic alterations associated with altered immune infilration in subsets of cancer through evaluation of gene expression, genomic alterations, pathologist-classified tumors, and protein validation. The Immunity Module assesses immune infiltration from bulk RNAseq expression for individual samples. The Function Module predicts the function (loss or gain) of individual samples for a given gene of interest using bulk RNAseq expression.

To address these gaps, we developed <u>R</u>apid Identification of <u>G</u>enomic <u>A</u>lterations in <u>T</u>umors affecting lymphocyte Infiltration (RIGATonI). RIGATonI is composed of two machine learning modules to connect functional genomic alterations (Function Module) and altered immune phenotypes (Immunity Module) in an unbiased manner. Our approach is distinct from currently available bulk RNAseq methods in four ways: 1) model training with pathologist-defined immune infiltration as gold standard, 2) consideration for immune cell spatial characteristics (e.g., tertiary lymphoid structures, dispersion characteristics, and degree of infiltration), 3) rapid, robust, and precise immune phenotype classification across big data resources, and 4) gene candidate filtering that evaluates protein function prediction rather than gene mutation status alone (**Figure 1A**). To build and validate RIGATonI, we combined histologic features identified by computational staining^14^, pathologist-defined immune infiltration, protein-protein interaction networks^15^, genomic data, proteomic data, and transcriptomic data (**Figure 1B**).

First, we built the Immunity Module to predict immune phenotypes using bulk RNAseq expression. Contrasting other approaches, this module was not built using exclusively immune-related genes; instead, we performed unbiased feature selection to determine the best predictors (n=114) of tumor immunity (**Figure 1A**). We validated and fine-tuned our approach using manually reviewed tumor histology by pathologists, computational staining^14^, immunohistochemistry^16^, and flow cytometry^16^ (**Figure 1A**). Next, we developed the Function Module which can accurately predict the function of genomic alterations (copy number alterations, single nucleotide variations, and structural variations) from bulk RNAseq expression. We validated this module using data from the largest collection of functionally impactful, clinically relevant genomic alterations in cancer: OncoKB^17^. These two modules were combined to uncover connections between all the genomic alterations in solid tumors in TCGA and the immune phenotypes of samples harboring these alterations (**Figure 1B**). We created an interactive visualization interface (https://rigatoni.osc.edu) to help researchers access our TCGA analysis results for individual genes (**Figure 1B**).

Additionally, we built an R package which can be used to perform the RIGATonI analyses on any samples of interest (**Figure 1B**).

To demonstrate the applications of the RIGATonI software, we explored a novel connection between 14q deletion in renal cell carcinoma and increased anti-cancer immunity discovered by RIGATonI’s analysis of TCGA. Using bulk RNA sequencing and single cell RNA sequencing from TCGA and Yu *et. al.*^18^ respectively, we show increased infiltration of CD8+ T cells, decreased pro-tumor immune checkpoint signatures, and increased CD8+ T cell proliferation, cytotoxicity, and inflammation.

Together, these results introduce RIGATonI: a unique and powerful tool designed with machine learning to identify immunologically impactful genomic alterations in cancer.

## RESULTS

### RIGATonI predicts immune phenotypes by utilizing histology and bulk RNAseq with high accuracy

RIGATonI’s Immunity Module was trained and validated using a comprehensive, pan-cancer dataset (OSU-ORIEN dataset) from The Ohio State University (OSU) including digital histology paired with bulk RNAseq (sequenced by Oncology Research Information Exchange Network, ORIEN). To ensure the OSU-ORIEN dataset included sufficient low, medium, and high immune phenotypes, we used a preliminary version of our machine learning algorithm developed using computational staining^14^ output from TCGA. We succeeded in doing so and produced a training data set of 403 tumors across 22 different cancer types (**Supplemental Figure 1**). Digital histology slides from these tumors were reviewed independently by two pathologists and were classified into low, medium, or high immune infiltration groups (**Figure 2A**). Pathologists used a semi-quantitative approach to estimate the percentage of tumor area occupied by lymphocytes. They also considered the distribution of these lymphocytes throughout the tumor area (e.g., deeply penetrating, semi-penetrating, or peripheral), and the overall quality of inflammation (e.g., presence or absence of tertiary lymphoid structures, signs of cytotoxic killing of tumor cells). To build the final model, we evaluated six different models using two different machine learning approaches and pathologist annotations both together and separately (**see Methods**). We selected genes for immune phenotype prediction within the Immunity Module using ElasticNet^19^, which yielded 114 transcriptomic features (**Supplemental Table 1**). All bulk RNAseq expression for these 114 genes were extracted and subsequent analyses utilized only this data. An XgBoost^20^ algorithm was trained using 334 of 403 tumors with Bayesian parameter optimization^21^ to classify tumors’ immune infiltration. We assessed the accuracy of the algorithm in a variety of ways.

**Figure 2.**
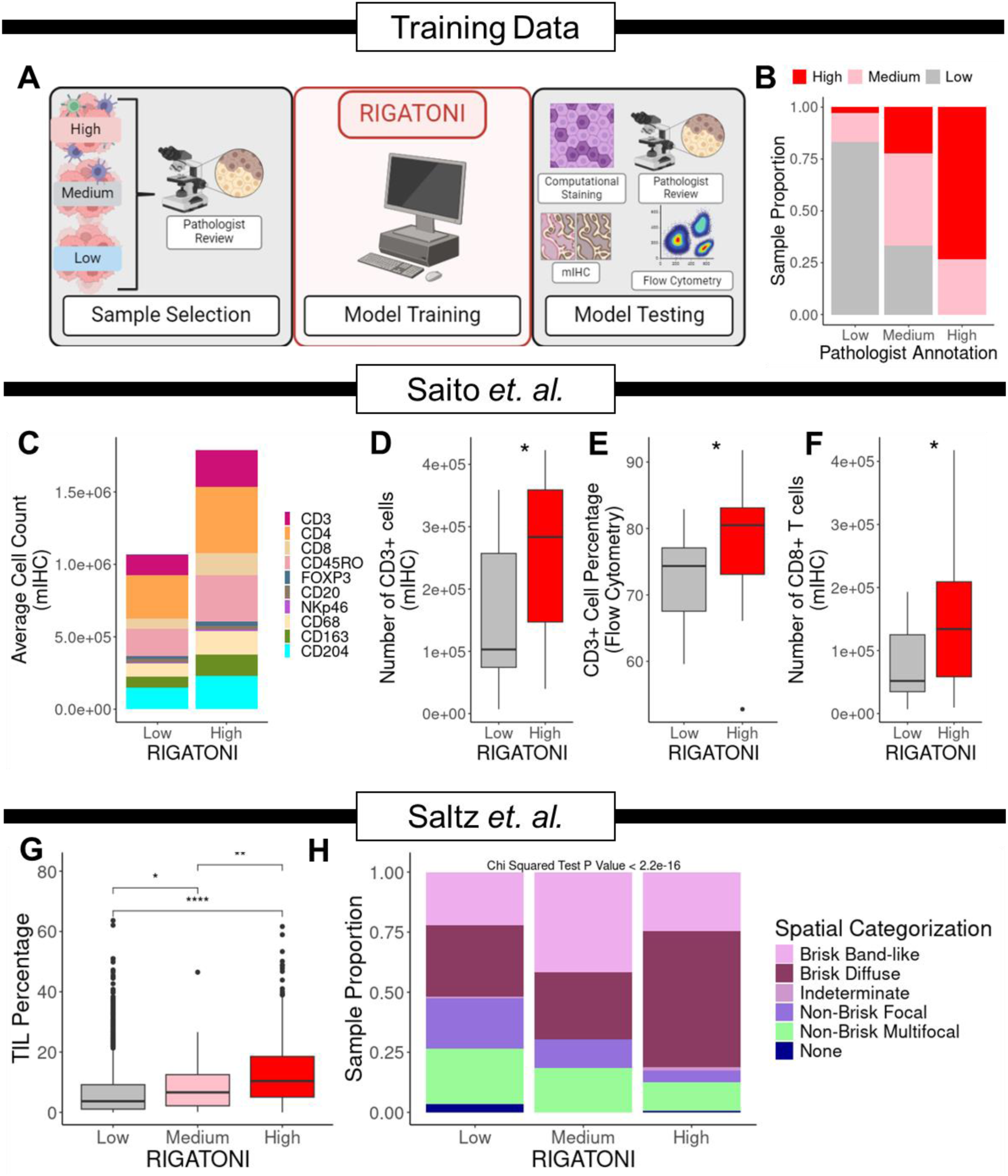
Development and validation of RIGATONI immune infiltrate classification. **A.** To train RIGATonI, we selected tumor samples based on a preliminary algorithm developed to identify a range of low to high lymphocyte infiltration phenotypes. Tumor samples were selected across diverse cancer types and two pathologists independently classified them as high, medium, or low degrees of infiltration. We built a machine learning model to predict these annotations and validated with a variety of data types. **B.** The accuracy of the model (Y-axis) was validated using a blind dataset (X-axis) not used for training data. RIGATonI-high annotations were 82% accurate for pathologist high and low annotations. RIGATonI-medium predictions were 63% accurate. **C.** Next, we investigated an independent dataset of gastric tumors with mIHC, flow cytometry, and tumor RNAseq data^16^. RIGATonI -high and -low samples corresponded to immune cell subsets as measured by mIHC. **D.** The counts of CD3+ T cells detected by mIHC were significantly increased in RIGATonI-high samples. **E**. Flow cytometry of these tumors demonstrated an increase in CD3-positive lymphocytes in RIGATonI -high vs. -low samples. **F.** The counts of CD8+ T cells detected by mIHC were significantly increased in RIGATonI-high samples. **G.** We also investigated our algorithm’s association with histologic features detected by convolutional neural networks in 5,202 tumors from TCGA^14^. The percentage of tumor infiltrating lymphocytes were measured by Saltz *et. al.*^14^ and corresponded with RIGATonI -low, -medium, and -high classifications. **H**. We evaluated spatial characteristics of lymphocyte infiltrates using Saltz *et. al.*^14^ approach. The overall distribution of spatial characteristics is significantly different across RIGATonI -low, -medium and -high subsets. RIGATonI-high samples often displayed brisk diffuse lymphocyte patterns shown in maroon. Significance values: p≤0.05: *, p≤0.01: **, p≤0.001: ***, p≤0.0001: ****.

First, we evaluated the accuracy of RIGATonI to classify 69 blind-set samples (**Figure 2B**). The overall accuracy of the model was 71.01% (95% confidence interval 58.84%-81.31%). The balanced accuracy was higher for tumors with high and low infiltration (82.04% and 82.58%, respectively) compared to those with medium infiltration (63.4%). A similar trend was observed for sensitivity and specificity (**Supplemental Table 2-4**). The accuracy was significantly different from the no information rate (p<0.01), indicating that the overall accuracy is significantly better than what could be achieved from random chance. We also failed to reject the null hypothesis of McNemar’s test, which indicates there is insufficient evidence that the predictions made by the algorithm are different from the true phenotypes (p>0.05). Detailed statistics and a confusion matrix of the results are available in **Supplemental Table 2-4**.

### RIGATonI’s Immunity Module corresponds with mIHC and flow cytometry features of increased lymphocyte infiltration

We validated the Immunity Module with a set of 32 gastric tumors from a recent study which provided matched multiplex immunohistochemistry (mIHC), flow cytometry, and bulk RNAseq^16^. Tumors classified “high” by RIGATonI (RIGATonI-high) had higher immune cell counts detected by mIHC (**Figure 2C**). Further analysis revealed that this increase was mainly due to a greater number of CD3+ cells in RIGATonI-high tumors compared to RIGATonI-low tumors (**Figure 2D**/**Supplemental Figure 2A)**. Further, flow cytometry data from the same study confirmed our findings. RIGATonI-high tumors showed a significantly higher percentage of lymphocytes (specifically CD3+ cells) compared to RIGATonI-low tumors (**Figure 2E/Supplemental Figure 2B)**. Additionally, we observed a significant increase in CD8+ T cells measured by mIHC in RIGATonI-high tumors compared to RIGATonI-low tumors (**Figure 2F**).

### RIGATonI phenotypes aligned with findings from computational staining of lymphocytes

We applied the RIGATonI Immunity Module to 5,202 tumors from TCGA and examined the output of an orthogonal approach for computational staining of digital pathology slides^14^. Saltz *et. al.*^14^ developed a convolutional neural network to detect the percentage and distribution of lymphocytes on digital histology images. First, we compared RIGATonI classifications to the predicted lymphocyte percentage on the slide (**Figure 2G**). We saw that RIGATonI-high tumors displayed significantly greater lymphocyte percentages compared to RIGATonI -medium or -low tumors (**Figure 2G**). Similarly, RIGATonI-medium tumors displayed significantly greater lymphocyte percentages compared to RIGATonI-low tumors (**Figure 2G**). Next, we compared RIGATonI classifications to five patterns of spatial attributes of tumors described by Saltz *et. al.*^14^: brisk band-like, brisk diffuse, indeterminant, non-brisk focal, non-brisk multifocal, or none (**Figure 2H**). We observed a significant difference in the spatial arrangements of lymphocytes between RIGATonI classifications (**Figure 2H**). Lymphocytes from RIGATonI-high tumors were more likely to have a brisk diffuse arrangement than the population (p<0.01), defined by a broad distribution of many lymphocytes throughout the histology slide^14^. RIGATonI-medium tumors display brisk band-like arrangements more often than the population (p<0.01). Brisk band-like infiltration patterns indicate the lymphocytes are clustered in a band across the slide, but that there are many lymphocytes^14^. RIGATonI-low tumors displayed a higher proportion of non-brisk focal lymphocyte arrangements (p<0.01). Non-brisk focal arrangements indicated negligible numbers of lymphocytes in a handful of locations across the image^14^. In summary, RIGATonI-high tumors exhibited extensive lymphocyte infiltration throughout the tissue slide, while RIGATonI-low tumors showed limited infiltration in isolated and/or scattered spots **(Figure 2H)**.

### RIGATonI includes an innovative Function Module which can accurately classify genomic alterations with molecular effects using protein-protein interaction networks

The Function Module first uses the STRING^15^ protein-protein interaction database to identify proteins which have direct and validated interactions with the protein of interest (**Figure 3A**). Both proteins which act on the protein of interest (upstream proteins) and proteins on which the protein of interest acts (downstream proteins) are considered. Next, we collect the gene names corresponding to the proteins into two lists. These two gene sets serve different purposes: upstream genes allow assessment of alterations which impact the expression level of the gene of interest; downstream genes allow us to assess the impact of alterations which may change the activity level of the gene of interest. We built a generalized linear model over a Poisson distribution using the extracted RNAseq counts of upstream and downstream genes using samples with no alteration in the gene of interest. These two models are then saved and used to predict the RNAseq counts of the gene of interest within the mutant samples. 95% prediction intervals are created for both the upstream and downstream models. Mutant samples with true counts below the 95% prediction interval of either model are classified as “LOF”. Conversely, samples with counts exceeding the 95% prediction interval are classified as “GOF”. Samples within the 95% prediction interval of both models are annotated as “unknown” (**Figure 3A**).

**Figure 3:**
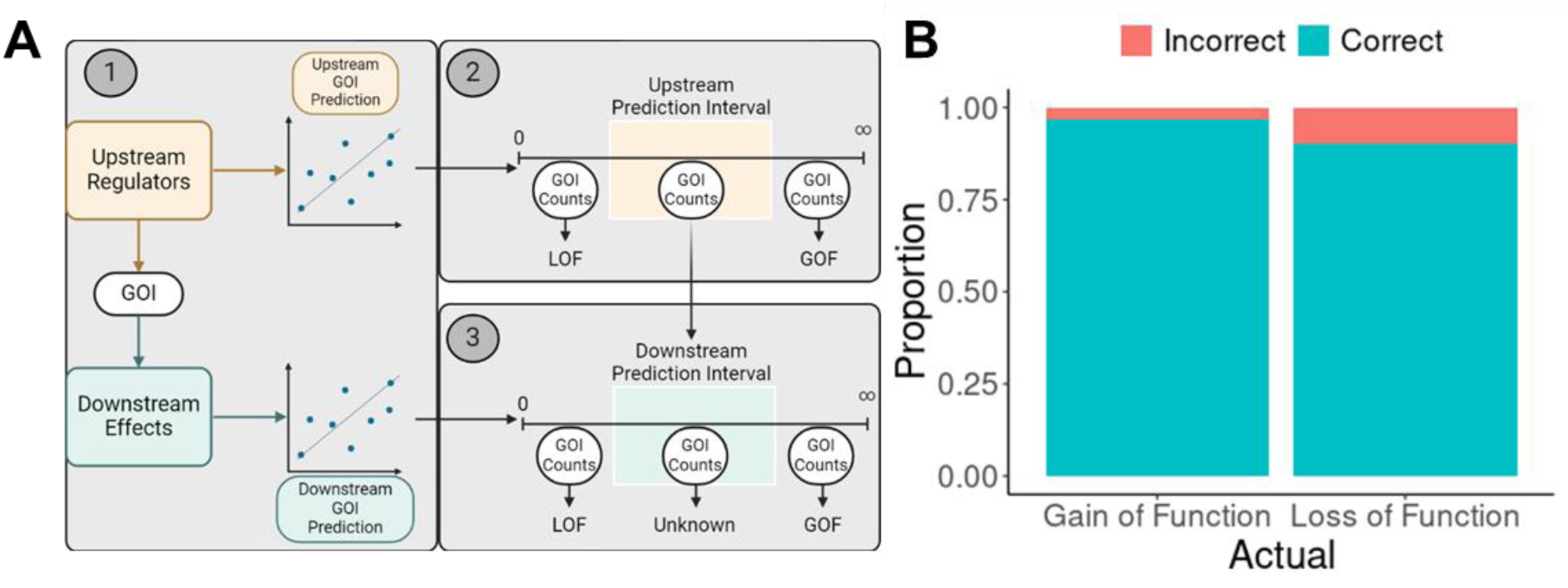
Function Module development and testing. **A.** The Function Module was developed using upstream regulators and downstream targets of a given gene of interest from the STRING^15^ database. Parallel linear models over a Poisson distribution were built. First, the model of the upstream regulators was assessed. Next, if the expression of the gene of interest falls within the prediction interval, gene expression predicted by the downstream targets was assessed. **B.** The Function Module was assessed using OncoKB^17^ as a ground truth. There were 291 genomic alterations with corresponding annotations in OncoKB^17^ and 96.8% of the gain-of-function calls (60/62) and 90.4% of the loss-of-function calls (207/229) were correctly classified.

To assess the Function Module’s accuracy, we analyzed genomic alterations from a total of 1008 oncogenes and tumor suppressor genes annotated by OncoKB^17^. When applied to 10,464 tumors from the TCGA, our algorithm successfully identified 400 GOF alterations and 966 LOF genomic alterations (SNVs, structural variations, and copy number variations) within these 1008 genes. Genomic alterations harbored by fewer than five tumors were not assessed. Fusions were excluded due to the high potential for false positives as many tumors harbored multiple fusions and imprecise breakpoints.

Of the 400 candidate GOF alterations uncovered, 62 were annotated in the OncoKB^17^ database. The algorithm correctly categorized 60 of 62 as GOF, and two were incorrectly categorized as LOF, yielding an accuracy of 96% for GOF (**Figure 3B**). Notably, 84.5% of the alterations annotated by the algorithm had no information available in OncoKB^17^; most of these variants were whole gene amplifications (**Supplemental Table 5).** Similarly, we validated the accuracy of this algorithm for LOF alterations. Of the 966 LOF alterations uncovered, only 229 were annotated in the OncoKB^17^ database. The algorithm correctly categorized 207 as LOF, while 22 were incorrectly categorized as GOF, yielding an accuracy of 90% (**Figure 3B**). Further, 76% of the candidate LOF alterations annotated by the algorithm had no information available in OncoKB^17^; most of these variants were whole gene deletions or premature truncations (**Supplemental Table 5)**. Overall, these findings demonstrate our algorithm’s effectiveness in accurately classifying known GOF and LOF mutations while also identifying novel variants that are not well described in existing databases.

### RIGATonI’s TCGA analysis is available for exploration online

Using the RIGATonI modules outlined above, we compiled and analyzed all genomic alterations (structural variations, gene fusions, point mutations, and copy number alterations) in TCGA. In total, RIGATonI identified 7,410 genomic alterations with possible immune effects among 5,746 genes. To determine the number of novel results among the RIGATonI output, we performed text mining on 226,093 abstracts mentioning “cancer” and “immunity” published between June 22nd, 2010, and June 22nd, 2023. We discovered that 2,773 (48%) of the RIGATonI output genes had not been previously connected to cancer immunity (**Figure 4A**). Only 72 genes (1%) had been mentioned in cancer immunity abstracts more than 100 times (**Figure 4A**). All results are available online at https://rigatoni.osc.edu/. Users select a gene of interest to explore and can subset output with alterations or cancer types of interest on the home page (**Supplemental Figure 3**). In the Transcriptomics page, the user can explore the expression levels of different genes across patient groups (**Supplemental Figure 4**). Finally, we provide cellular deconvolution results from quantiseqR^7^ on the Immunity page (**Supplemental Figure 5**). This website is intended to assist researchers in understanding how the function of their gene of interest impacts immunity in an unbiased and rapid manner.

**Figure 4.**
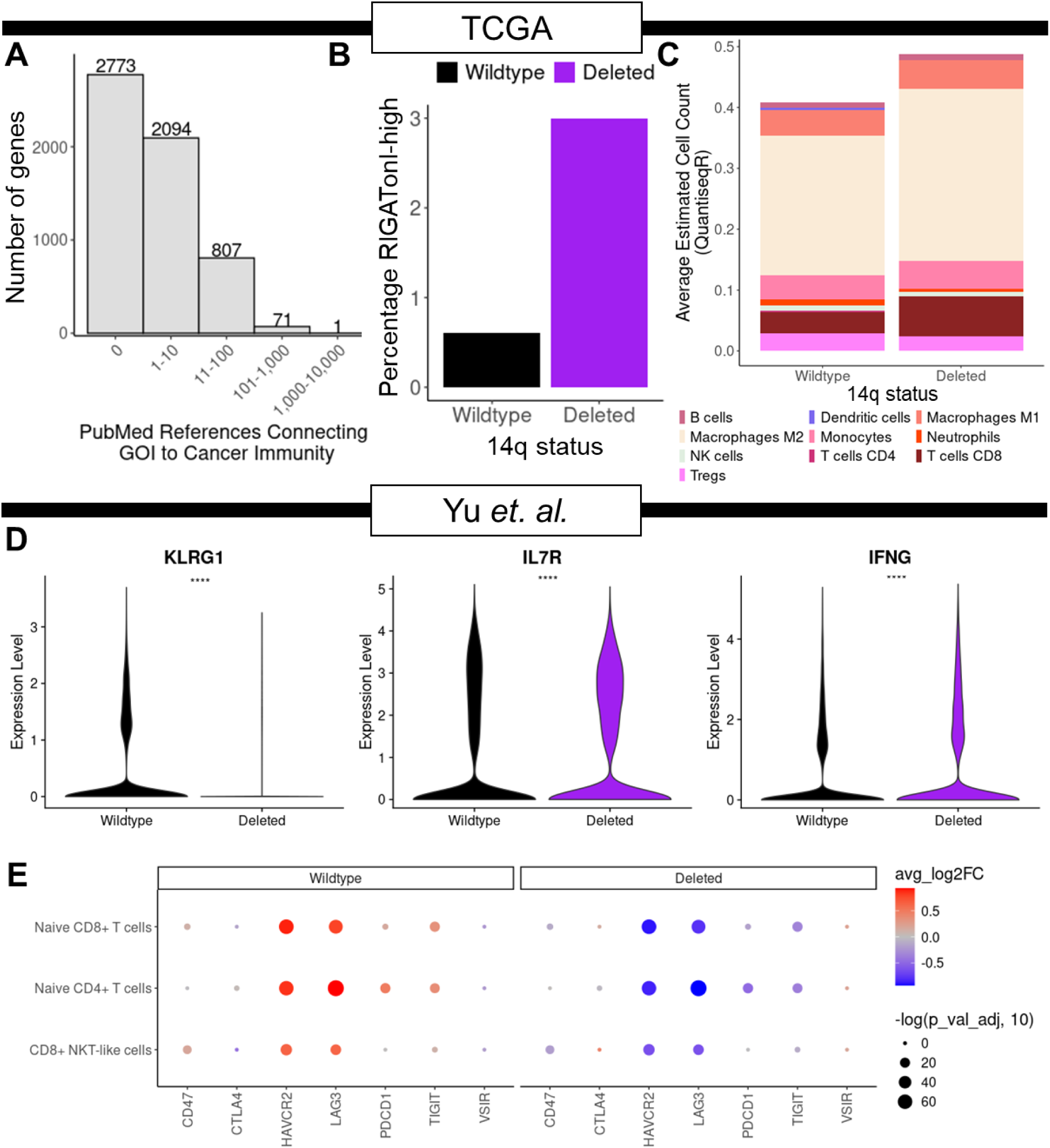
RIGATonI identifies novel genomic alterations of interest including 14q deletion associated with an increased immune infiltrate and effector CD8+ T cells in renal cell carcinoma. **A.** Through analysis of TCGA and text mining of PubMed, 48% (n = 2773) of genes harboring RIGATonI-identified genomic alterations (n = 5746) have never been associated with tumor immunity. **B.** As an example, RIGATonI identified 14q deletion in renal cell carcinoma (RCC) samples which corresponded to an increased immune infiltration compared to wildtype tumors. **C.** Using quanitseqR^7^, we corroborate our finding that there is a broad increase in immune cells in 14q-deleted RCC tumors**. D.** With 19 scRNAseq RCC experiments, we investigated markers of CD8+ T cell exhaustion and a “cold” immune microenvironment. CD8+ T cells from 14q-deleted tumors displayed decreased exhaustion marker *KLRG1* and increased anti-tumor immunity markers *IL7R* and *IFNG*^22–24^ **E.** Immune checkpoints, which are thought to promote tumor growth, are downregulated across all T cells studied in 14q-deleted tumors compared to wildtype tumors. These checkpoint receptors include *CD47*, *CTLA4*, *HAVCR2*, *LAG3*, *PDCD1*, *TIGIT*, and *VSIR* ^25, 26^. Significance values: p≤0.05: *, p≤0.01: **, p≤0.001: ***, p≤0.0001: ****

### RIGATonI identifies genomic alteration 14q deletion’s association with altered immune infiltration in renal cell carcinoma

Exploring novel results from the TCGA analysis, we uncovered a group of genes (n=207) all located on chromosome 14q which consistently showed the same patterns of deletion, in the same patients, and were RIGATonI-high. Further investigation led us to discover that these alterations were part of a larger chromosomal arm deletion of 14q. We observed this pattern in many cancer types, but one of the strongest relationships was in renal cell carcinoma (RCC). 3% of RCC patients harboring a 14q deletion were highly infiltrated compared to 0.6% of 14q wildtype RCC patients (**Figure 4B**). We also corroborated these results using quantiseqR^7^ where 14q-deleted tumors displayed an increased immune infiltration compared to wildtype (**Figure 4C**). Although these two bulk RNAseq methods show the same pattern, we also evaluated 14q deletion in an orthogonal dataset with 19 RCC tumors with single cell RNAseq^18^. Using copyKat^27^, we discovered 8/19 patients harbored a 14q deletion. We evaluated the immune cell compartment of these samples and observed that CD8+ T cells from 14q-deleted tumors displayed decreased *KLRG1*, increased *IL7R*, and increased *IFNG* expression (**Figure 4D**). This pattern is indicative of increased T cell proliferation, cytotoxicity, and T cell-mediated anti-tumor immunity among CD8+ T cells from 14q-deleted tumors^22–24^. We also investigated the expression of immune checkpoint genes expressed on T cells (*HAVCR2*, *TIGIT*, *LAG3*, *PDCD1*, *VSIR*, and *CD47*) (**Figure 4E**)^25, 26^. Across all T cells studied, expression of pro-tumor checkpoint genes is decreased in 14q-deleted tumors (**Figure 4E**). The largest differences were in *LAG3* (LAG-3) and *HAVCR2* (TIM-3) (**Figure 4E**). CD4+ T cells were the only cell type to have a significant decrease in *PDCD1* (PD-1) within 14q-deleted tumors (**Figure 4E**). Together these results indicate that 14q-deleted tumors contain more activated CD8+ T cells than wildtype as well as fewer features of T cell exhaustion and pro-tumor immune activity.

## DISCUSSION

RIGATonI is a novel tool to discover precision immuno-oncology targets using bulk RNAseq data in two distinct modules: the Immunity Module (**Figure 2A**) and the Function Module (**Figure 3A**). The Immunity Module addresses two key gaps not considered by current approaches 1) degree of immune cell dispersion alongside infiltration, and 2) immune phenotyping across big data resources without user interpretation. The Immunity Module uses a machine learning approach built with paired bulk RNAseq from pathologist reviewed histology slides (**Figure 2A)**. To our knowledge, this strategy has never been implemented before. The Function Module classifies genomic alterations as LOF or GOF based on bulk RNAseq features (**Figure 3A)**. RIGATonI is the first approach we are aware of for estimating immunity which also estimates gene-level function. RIGATonI was employed across the TCGA database and interesting results are visualized on our website (https://rigatoni.osc.edu/) for public use. Finally, we demonstrated that these two modules can be combined to discover novel connections between genomic alterations and immunity through further investigation of 14q deletion in renal cell carcinoma (RCC) scRNAseq datasets. In summary, RIGATonI is a unique machine learning approach specifically designed for precision immuno-oncology.

The Immunity Module considers features of cancer immunity not considered by available software tools. Currently, gene set enrichment, cellular deconvolution, and ensemble approaches for estimating immunity from bulk RNAseq have not been compared to benchmark pathologist review of tumor histology for degree and quality of lymphocyte infiltration (**Figure 1A**)^4–8^. To build RIGATonI’s Immunity Module, we asked pathologists to review tumor histology slides for degree of infiltration considering not only the absolute number of lymphocytes, but also the presence or absence of tertiary lymphoid structures (TLS), signs of cytotoxicity, and the general distribution across the tumor slide (deeply penetrating the tumor or on the periphery). By incorporating these nuanced features into our algorithm training and validation, RIGATonI benchmarks aspects of immunity not evaluable with either gene set enrichment or cellular deconvolution tools^4–8^. Furthermore, both cellular deconvolution and gene set enrichment utilize genes with known connections to cancer immunity (**Figure 1A**). These genes are selected either through literature review or single cell atlases^5^. Unfortunately, this approach does not allow for evaluation of *de novo* mechanisms that impact tumor immunity. To address this gap, RIGATonI’s Immunity Module uses expression of just 114 genes selected in an unbiased manner (**Supplemental Table 1**). These genes consistently predict immune phenotypes across cancer types and databases (**Figure 2B-H**). In the future, we will explore why this novel gene list can reproducibly predict cancer immune infiltration.

The Function Module’s prediction of protein-level effects of genomic alterations from bulk RNAseq expression data is also novel. Most genomic alterations in tumors are variants of unknown significance^28^. Therefore, a functional measure of associated gene activity can help to prioritize alterations that are most likely to have a biological or immunological impact. eVIP2 is the only other method designed to determine the function of genomic variants using bulk RNAseq^29^. Importantly, eVIP2 was validated using only two alterations in the same gene of interest, whereas our validation set demonstrated remarkable accuracy involving 291 different alterations across 207 different genes (**Figure 3B**)^29^. VIPER, another R package, utilizes cell specific “regulons” for protein activity prediction and did assess scores for some genomic variants of interest; however, it was not designed to analyze the functional impact of genomic alterations directly^30^. RIGATonI is the only available bulk RNAseq method to assess protein function from gene expression validated with several hundred genomic alterations across many different genes. In contrast to DNA-based annotation of mutations, the Function Module has several benefits. First, although we applied the Function Module to groups with differing genomic alteration status, the module assesses the function of a gene of interest in a testing group compared to a control group. Thus, RIGATonI could be applied to any subsets of any other feature (e.g. methylation, alternative splicing, treatment, etc.) (**Figure 3A**). Second, when evaluating genomic alterations, RIGATonI can assess novel variants and mechanisms of altered expression. Third, not all patients with the same genomic alteration experience identical molecular effects. The Function Module predicts the overall effects of alterations and makes specific predictions for each sample, considering its unique molecular characteristics (**Figure 3A**). These unique features make RIGATonI’s Function Module an effective tool for multi-omic research.

Like many computational approaches, RIGATonI is limited by sample size considerations. By using pathologist assessment of tumor histology rather than computer vision as a ground truth for the Immunity Module, we do limit the sample size of our training data. New approaches like Lunit SCOPE IO^31^ and that of Saltz *et. al.*^14^ use deep learning techniques to estimate immune cells on digital pathology slides. Lunit SCOPE IO was trained on >17,000 H&E images across 24 cancer types^31^. This scale of training data would be impractical using manual pathologist review. However, it is widely agreed that physician benchmarking is the gold standard against which machine learning approaches should be measured^32^. Therefore, we believe the quality of the training data classifications compensated for comparatively smaller sample size. Another sample size concern arose when validating the Function Module. Validation of RIGATonI’s Function Module only took place on alterations with ≥5 instances of a candidate alterations to ensure that statistical tests could be performed to determine the functional status. In the pan-TCGA analysis, we did not group together early terminations or different point mutations occurring at the same locus out of an abundance of caution (early terminations at different loci may have different effects; different point mutations at the same locus may have different effects). Without detailed knowledge of all 17,000 genes analyzed, we approached the first version of RIGATonI conservatively. Despite these sample size concerns, our robust validations give us confidence in our tool’s performance.

RIGATonI identified 14q deletion as a potential novel biomarker for increased anti-tumor immunity in renal cell carcinoma (RCC). 14q deletion has previously been identified as a negative prognostic indicator in RCC; however, these studies were done prior to the broad adoption of PD-1/PD-L1 immunotherapy in RCC^33, 34^. RIGATonI indicates 14q deletions are associated with a highly infiltrated immune microenvironment in TCGA (**Figure 4B**). These results were further supported using the cellular deconvolution tool quantiseqR^7^ which demonstrates enhanced immunity in 14q-deleted samples (**Figure 4C**). We were able to orthogonally assess 14q through analysis of scRNAseq data for 19 RCC patients^18^. We first explored the CD8+ T cells between 14q-deleted and wildtype tumors. CD8+ T cells from 14q-deleted tumors displayed evidence of superior cytotoxicity, increased release of interferon-gamma, and increased proliferation^22–24^. We also explored whether pro-tumor immune checkpoint receptors and ligands were more highly expressed in T cells from 14q-deleted vs wildtype tumors. Wholistically, we see that pro-tumor immune checkpoint receptors are decreased in 14q-deleted tumors compared to wildtype tumors across all cell types explored (**Figure 4E**)^25, 26^. Identification of 14q deletion in RCC demonstrates a successful application of RIGATonI to discover genomic alterations associated with altered tumor immunity.

In summary, RIGATonI is a powerful software leveraging tumor RNAseq in novel ways to accelerate discoveries in precision immuno-oncology research. RIGATonI can be applied across big data sources to understand tumor-driven mechanisms of immunity, identify novel biomarkers for immunotherapy treatment, and discover novel drug targets for future immunotherapies. In summary, we are pleased to introduce RIGATonI: an innovative approach for discovery novel genomic variants with associated altered immune phenotypes.

## ONLINE METHODS

### Tumor sequencing data

All data shown here were previously published or made available by a big-data source (described below). Secondary analyses were performed on the Pitzer cluster R studio v4.3.0 within Ohio Supercomputer Center (https://www.osc.edu/). Data sources include The Cancer Genome Atlas (TCGA) and the Oncology Research Information Exchange Network (ORIEN). RNAseq count data were downloaded from TCGA using GenomicDataCommons^35^ and batch corrected with ComBat-seq^36^ using institution of origin as the batch. Genomic variant calling data was downloaded in the form of combined .maf files. Copy number alteration data were downloaded in the form of gene based raw copy number. Finally, all whole genome .bam files were downloaded from TCGA and then processed with parliament2^37^ using Delly^38^, Manta^39^, breakdancer^40^, and breakseq^41^ to assess for structural variants. Results from parliament2^37^ were combined using SURVIVOR^42^ with default settings. RNAseq data was also obtained from ORIEN and batch corrected with ComBat-seq^36^ according to the RNAseq batch information made available through Aster Insights. Copy number alterations were downloaded from ORIEN in the form of gene based raw copy number. Demographic information was downloaded from ORIEN as well. ORIEN data is managed by Aster Insights, requests for this data should be sent to Aster Insights.

### RIGATONI immune phenotyping algorithm development and validation with pathologist review

Two pathologists independently reviewed 403 tumor slides assessing lymphocyte infiltration characteristics. Pertinent characteristics included percentage of space not occupied by tumors or stroma, which was occupied by lymphocytes, dispersion of lymphocytes within tumor, and presence or absence of tertiary lymphoid structures.

Taking into consideration all these characteristics, each pathologist annotated the slide either high, medium, or low infiltration. Paired RNA sequencing was collected and used along with annotations to build a series of models predicting immune phenotypes from bulk RNAseq counts. First, important features were selected using multinomial ElasticNet^19^ via the R package glmnet^43^. Next, these features were used as predictors for six different machine learning models. Three models were built via the R package XgBoost^20^ using 10-fold cross validation alongside Bayesian parameter optimization^21^ via the R package ParBayesianOptimization^44^. Three separate models were built with the R package ordinalForest^45^. Two models of the above six models were built considering both pathologists’ predictions, two considering just pathologist-one and other two considering only pathologist-two. The classification accuracy of each algorithm for each group is available in **Supplemental Table 6**. Each algorithm uses the same cohort of 334 samples for training and 69 samples for testing. The algorithm with the best performance (determined with caret^46^) on the testing data was selected to be used going forward. More specific information is provided in **Supplemental Table 6**. Accuracy of the testing data for the model selected were visualized using ggplot2^47^ and ggpubr^48^.

### Gastric cancer multiplex immunohistochemistry (mIHC) and flow cytometry

RNAseq count data was downloaded from Saito *et. al.*^16^ along with flow cytometry and IHC outputs. These include 32 patients with gastric cancer in Tokyo^16^. These results were processed as previously described in Saito *et al.*^16^ Using this resource, we used a MANOVA^49^ to compare the IHC counts to determine if there were any significant differences between groups. Next, we used a Wilcox test^50^ to assess the counts of each subset of IHC-marked cells to find significant differences. We performed pairwise comparisons of flow cytometry results using Wilcox tests^50^. Results were visualized using ggplot2^47^ and ggpubr^48^.

### Determine resulting immune phenotype of each genomic alteration using RNAseq data

An R function was created which converts RNAseq count data to TPM using the R packages DESeq2^51^ (to correct size factors) and DGE.obj.utils^52^ (to convert to TPM), and then filters the data down to only the genes selected by ElasticNet^19^. The immune phenotype of each sample was predicted using XgBoost^20^. The proportion of high and low tumors for each cancer type were calculated. To analyze a genomic alteration, all mutant samples provided to the function are compiled, and a 1-proportion z-test is performed where the null hypothesis is that the proportion of hot samples will be equal to the population proportion within that cancer type and the alternative hypothesis is that the proportion of high samples is greater than the population proportion. If the null hypothesis is rejected (p<0.05), the genomic alteration is annotated RIGATonI-high. If not, a second 1-proportion z-test is performed where the null hypothesis is that the proportion of low samples will be equal to the population proportion and the alternative hypothesis is that the proportion of low samples is greater than the population proportion within that cancer type. If this null hypothesis is rejected (p<0.05), the genomic alteration is annotated RIGATonI-low. If not, the alteration is annotated “Unknown”.

### Determining functional status of each genomic alteration using RNAseq data

The STRING^15^ database’s protein action version 10.5 was downloaded. The list of protein actions is subset to include only actions on or by the protein of interest (POI). These actions are further filtered into two lists: an upstream protein list with only proteins that act to affect the expression of the POI, and a downstream protein list including all genes the POI activates, inhibits, or alters expression. The upstream gene list is used to model the RNAseq counts of the POI using modulators of the POI’s expression. The downstream gene list is used to model the RNAseq counts of the POI using downstream genes as indicators of its activity.

To model typical expression patterns of the POI, all samples with no alteration (control samples) in the gene of interest (GOI) are collected, and two generalized linear models are created over a Poisson distribution to predict the RNA counts of the POI/GOI. One model uses the upstream protein list as predictors, and another uses the downstream protein list. Both models predict the RNAseq counts of the POI.

Next, the RNAseq counts of the GOI within samples harboring mutations (mutant sample) predicted separately with each regression model. If, in either model, the expression of the GOI is lower than the lower bound of the 95% prediction interval (created by ciTools^53^), the mutant sample is annotated loss of function (LOF). On the other hand, if, in either model, the expression of the GOI is higher than the upper bound of the 95% prediction interval, the mutant sample is annotated gain-of-function (GOF). Falling outside the bounds of these prediction intervals indicates that the mutant sample’s GOI expression or activity is more different than that of a control sample than we would expect from random chance. If the mutant sample’s GOI expression falls within the bounds of both these prediction intervals, we can conclude that there is not enough evidence to indicate the mutation is causing abnormal expression or activity in that sample.

Next, all mutant samples provided are compiled, and a 2-proportion z-test is performed (null hypothesis z=0.5) comparing the proportion of LOF and GOF annotations. If the null hypothesis is rejected (p<0.05), the genomic alteration is annotated with the more frequent sample level annotation. If not, the alteration is annotated “Unknown.”

### Function annotation algorithm validation

OncoKB^17^ provides a list of oncogenes with either targeting drugs or known oncogenic mutations. All alterations in these 1008 genes were analyzed in parallel. The results were filtered to only include GOF and LOF calls from the algorithm. Various alterations are called Unknown due to low sample count, confounding variables, or lack of known connections in STRING^15^. We did not include these samples in the pan-cancer analysis of TCGA, however users can elect to include them in their own analysis. Additionally, gene fusions were removed due to complexity of their calling. This left 1366 genomic alterations to investigate. We manually searched OncoKB^17^ for information about each alteration and, if available, recorded the true function of the variant. Results were visualized with ggplot2^47^ and ggpubr^48^.

### Building the R package

Functions were created with the R package devtools^54^ which create an upstream and downstream gene list from STRING^17^, determine the function of a group of samples using RNA expression data from bulk RNAseq, and the sample level immune phenotype using RNA expression data from bulk RNAseq. These functions are described in detail above. The R package along with relevant documentation is available at https://github.com/OSU-SRLab/RIGATONI.

### Comprehensive analysis of genomic alterations in TCGA

All mutation information in TCGA was downloaded and compiled. A sample is said to have a copy number variation (CNV) if the total number of copies is ≥6 or <2. We also considered the sex of the patient in question if the gene of interest was on the X or Y chromosome, and we were considering a copy number loss. For male patients, genes on the X chromosome were said to be deleted if there were zero copies. For female patients, no genes on the Y chromosome were considered deleted. For Identification of genomic variants and altered immune phenotypes in cancer any patients without biological sex available, we excluded them from the copy number analysis of genes on the X or Y chromosome. Next, all genomic alterations were analyzed in parallel through the RIGATonI R package described above. We ensured to analyze each alteration within each cancer type for both functional annotations and immune phenotyping. The population proportions used for immune phenotype were those of the cancer type being analyzed. Finally, we stored significant results from TCGA to be visualized by our online tool. The RIGATonI website is located at https://rigatoni.osc.edu/ and is managed by the Roychowdhury lab group and the Ohio Supercomputer Center. The website is built with the R package shiny^55^, all graphs are visualized with ggplot2^47^ and ggpubr^48^. The website input is a user provided GOI, and the website compiles gene level copy number, fusion, and simple single nucleotide variation results from the pan TCGA analysis. Next, we provide various visualizations to the user (quantiseqR^7^ cell type proportions, RNAseq count data, and primary site prevalence) along with a table describing the different genomic alterations which were both functional and immunogenic within the GOI.

### Text mining of PubMed abstracts to estimate novelty of RIGATONI TCGA output

PubMed abstracts containing the words “cancer” and “immunity” or “immunology” or “immune” since June 12^th^, 2010, were downloaded using the R package pubmed.mineR^56^. The function gene_atomization was used to perform text mining annotation of each gene mentioned. The RIGATonI results were extracted, and each gene was annotated with their frequency of appearance. Preprint publications were excluded from this analysis. 226,093 abstracts were analyzed. Results were visualized with ggplot2^47^ and ggpubr^48^.

### ScRNAseq analysis of renal cell carcinoma

The data was analyzed using Seurat^57–60^ Quality control measures were performed as follows: remove cells with <5x the standard deviation below median feature count, >5x the standard deviation above median feature count, <5x median total count, and <10% mitochondrial gene expression. We performed quality control steps for each sample individually. To mitigate experimental batch effect, we used harmony^61^ and clustered using clustree^62^. To perform cell typing, we clustered all experiments together using UMAP with the Seurat^57–60^ package. We then cell typed using the “kidney” tissue designation from ScType^63^. Any cells which were not able to be typed using the “kidney” designation were separated, clustered again using UMAP and Seurat^57–60^, and re-typed using “Immune system” as the tissue of origin. Using copyKat^27^, a scRNA-seq method to detect copy number alterations, we have identified 14q deleted tumors by using hematopoietic cells as a somatic control. We choose 14q deletion based on average copyKat score across chromosome 14q. If the score was less than zero on average, we determined that the sample harbored a 14q deletion. Additionally, we confirmed this by comparing the expression of all suspected 14q deleted tumor cells to all suspected wildtype tumor cells for each gene on chromosome 14q. For each gene, the 14q deleted tumors displayed significantly lower expression, Finally, using the FindAllMarkers function from Seurat^57–60^, we investigated cell markers which were differentially expressed between 14q deleted and wildtype samples. These results were visualized using base Seurat^57–60^ functions, ggplot2^47^, and ggpubr^48^.

## Acknowledgements

The results published here are in whole or part based upon data generated by the TCGA Research Network: https://www.cancer.gov/tcga. All computational analyses were done on the Pitzer cluster at the Ohio Supercomputer Center (OSC) (https://www.osc.edu/). OSC also assisted in developing and hosting the RIGATonI website.

## Author Contributions

RV conceived the idea for RIGATonI, developed the algorithms, wrote all R code for the project, wrote all Linux code along with ELH and ES, managed data, reviewed data analyses, and wrote/revised/edited the manuscript. ELH assisted with RIGATonI algorithm development, assisted with Linux coding, assisted with data management, and revised/edited the manuscript. LY assisted with RIGATonI algorithm development, assisted with data management, and revised/edited the manuscript. JWR, MRW, LS, and AP reviewed data analyses, revised/edited the manuscript. ES assisted with Linux coding, wrote all python code, and revised/edited the manuscript. EGB reviewed OncoKB and annotated the RIGATonI function algorithm’s validation output. MC and NS enabled access and sequencing of the OSU-ORIEN dataset’s RNAseq. WC and AF performed pathologist review of the OSU-ORIEN digital pathology images. SR conceived the idea for RIGATonI, reviewed data analyses, and wrote/revised/edited the manuscript.

## Libraries

Load necessary libraries:

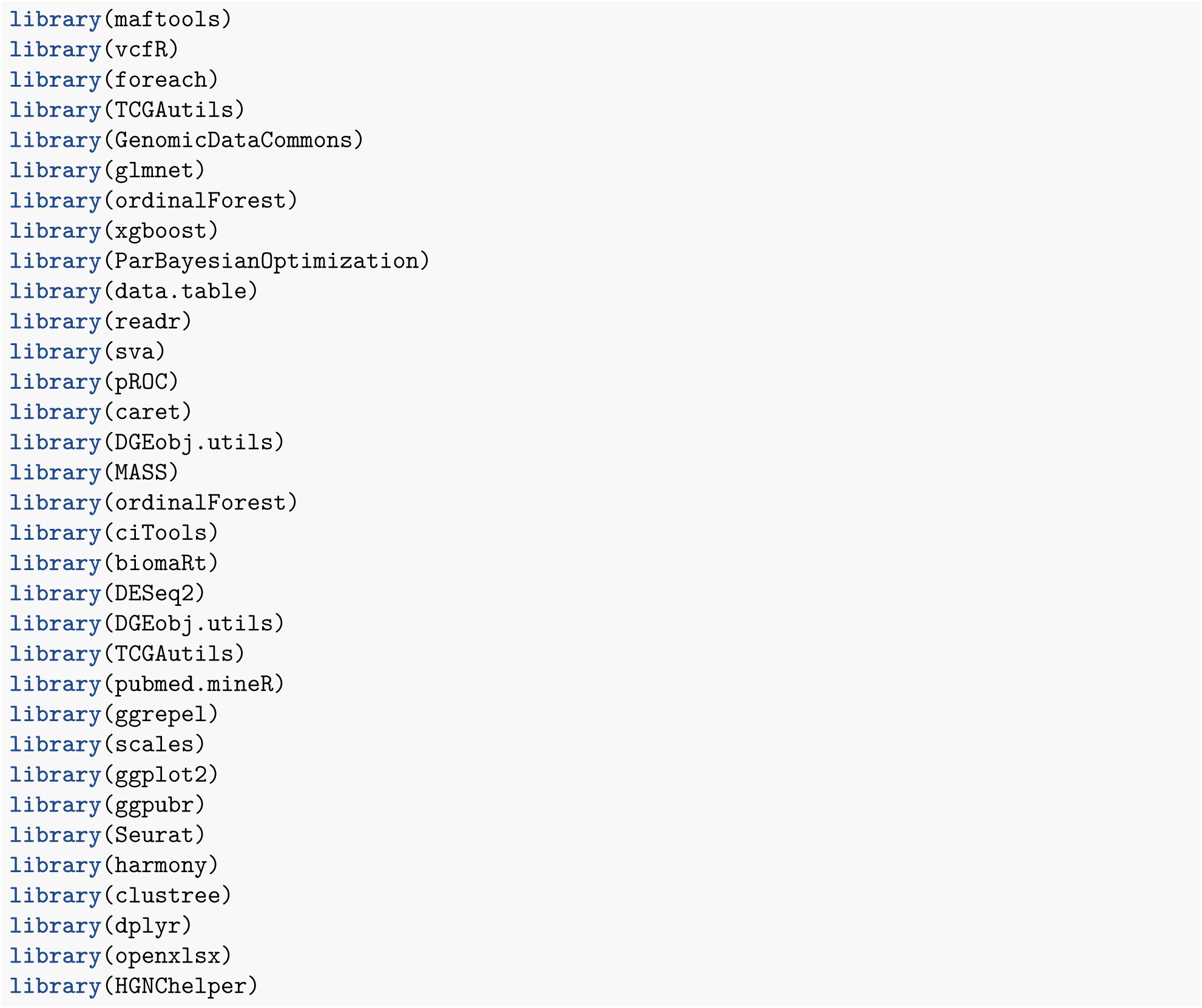

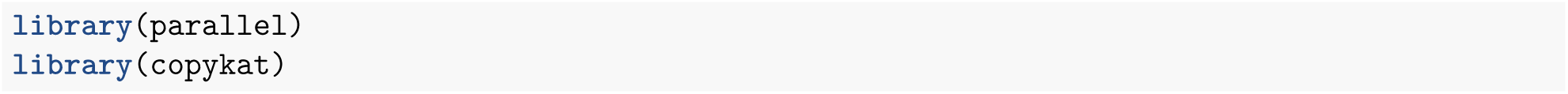

## Part 1: Process simple nucleotide variations (SNVs), copy number variations (CNVs), Structural Variants (SVs), and gene fusion outputs from TCGA

In order to run RIGATONI on an entire database, you need to have all the fusion, SNV, CNV, and SV outputs from said database in list format. I used the outputs directly from TCGA for the SNV, CNV, and Fusions callers. I processed the whole genome samples using parliament2 for SVs myself. Shown below are the steps for collating and organizing the SNVs, CNVs, and fusions.

### 1.1: Download SNV, CNV, and fusion outputs from TCGA

First, I downloaded the SNV, CNV, and fusion outputs from TCGA. I created manifests using the GDC database and downloaded the files as shown below.

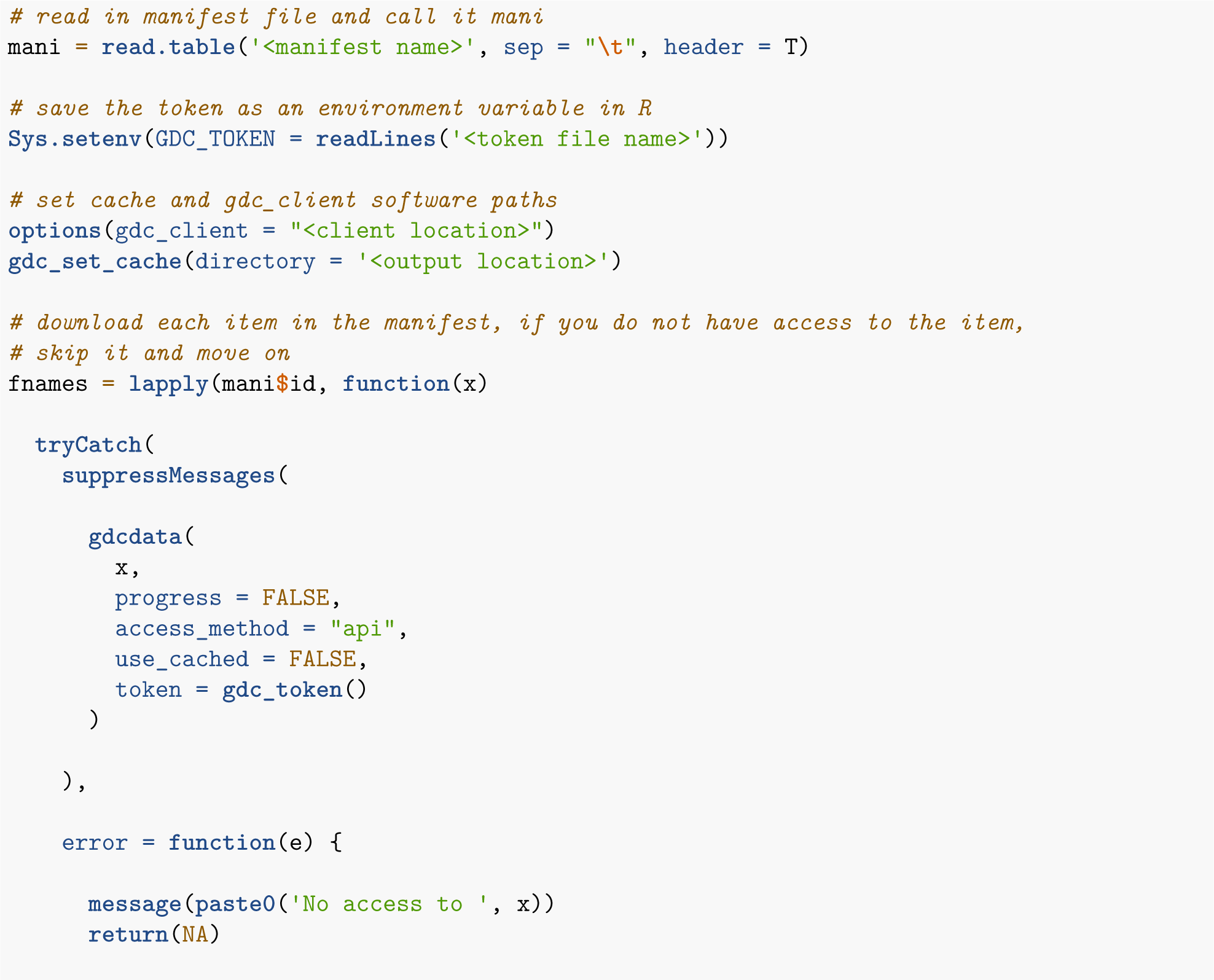

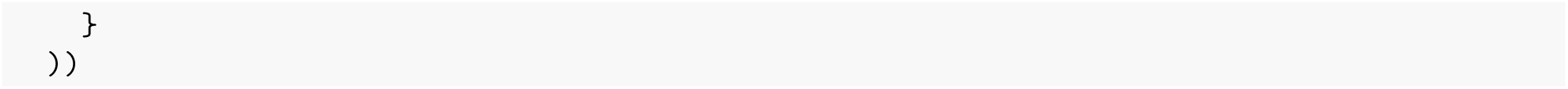

### 1.2: Process SNV outputs into a list

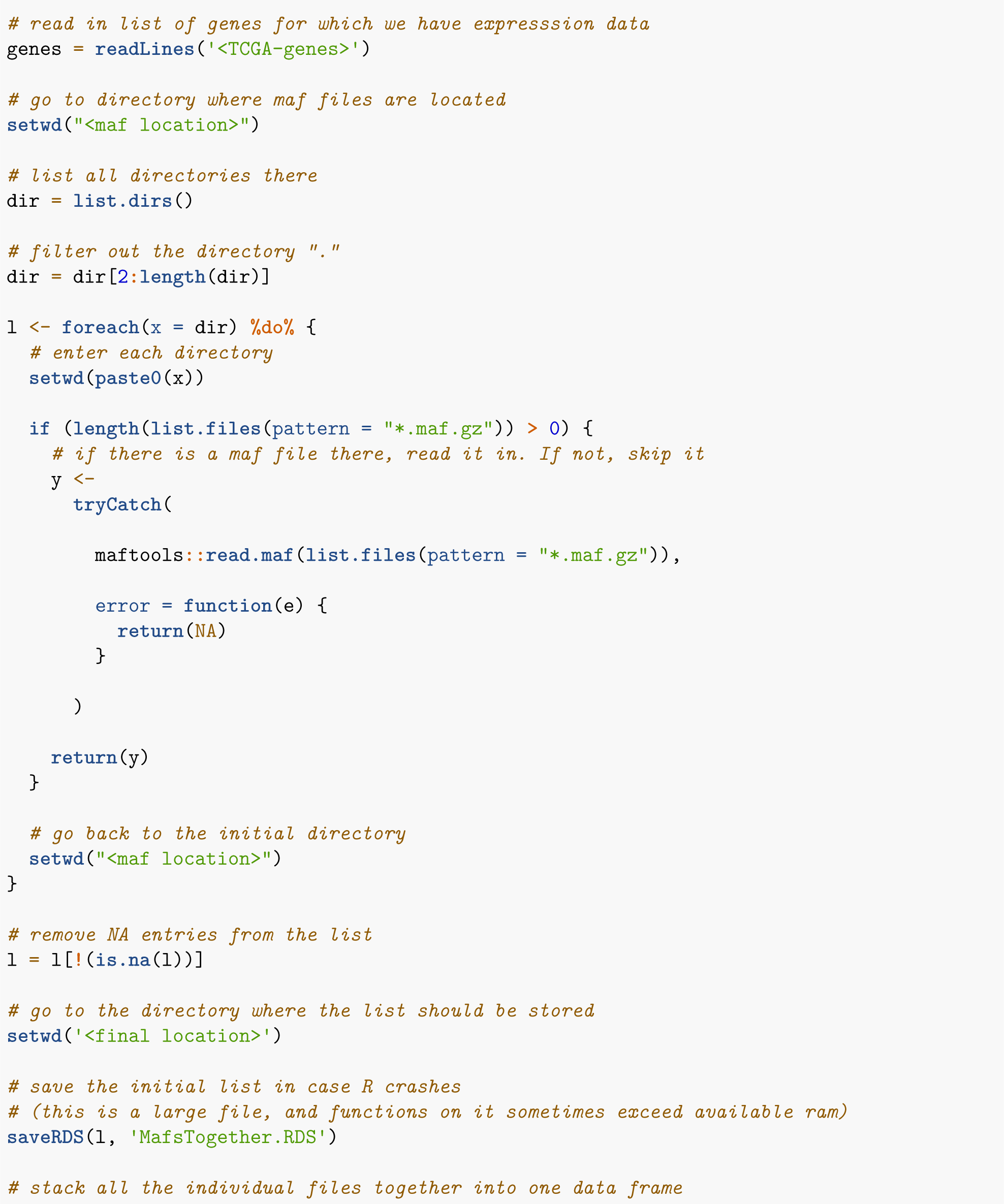

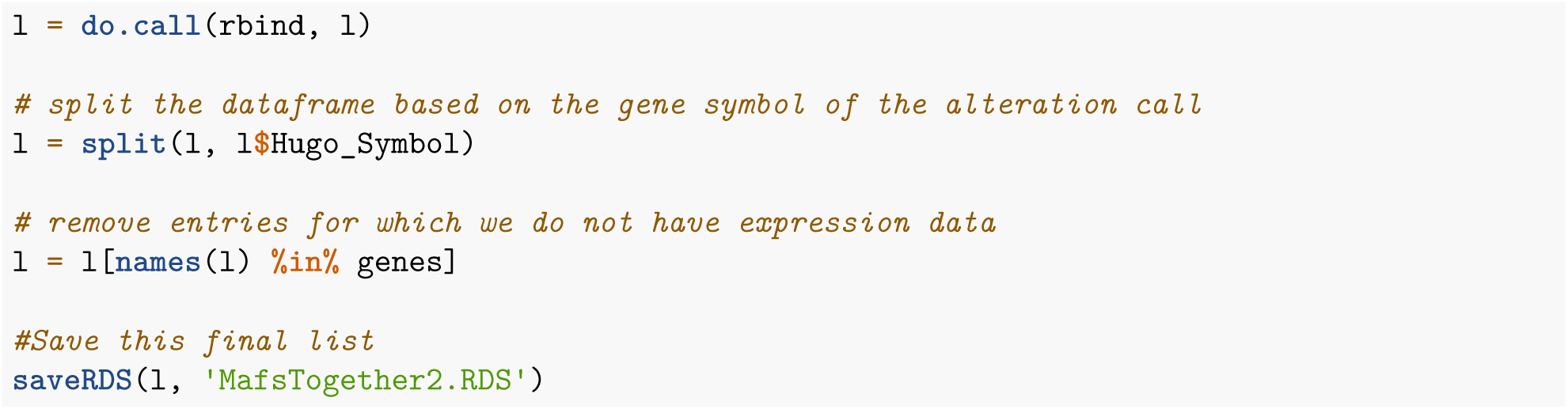

### 1.3: Process fusion outputs into a list

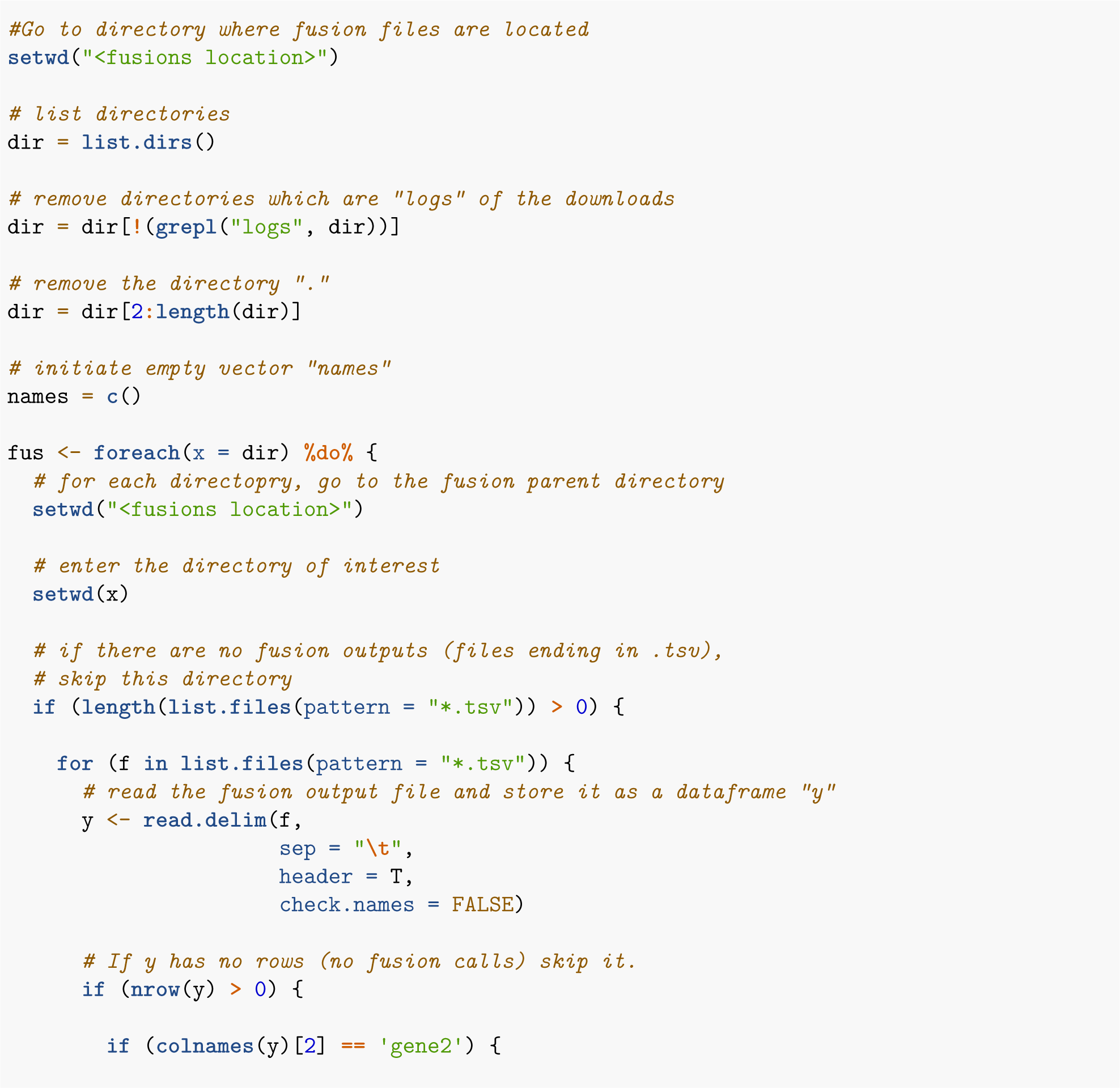

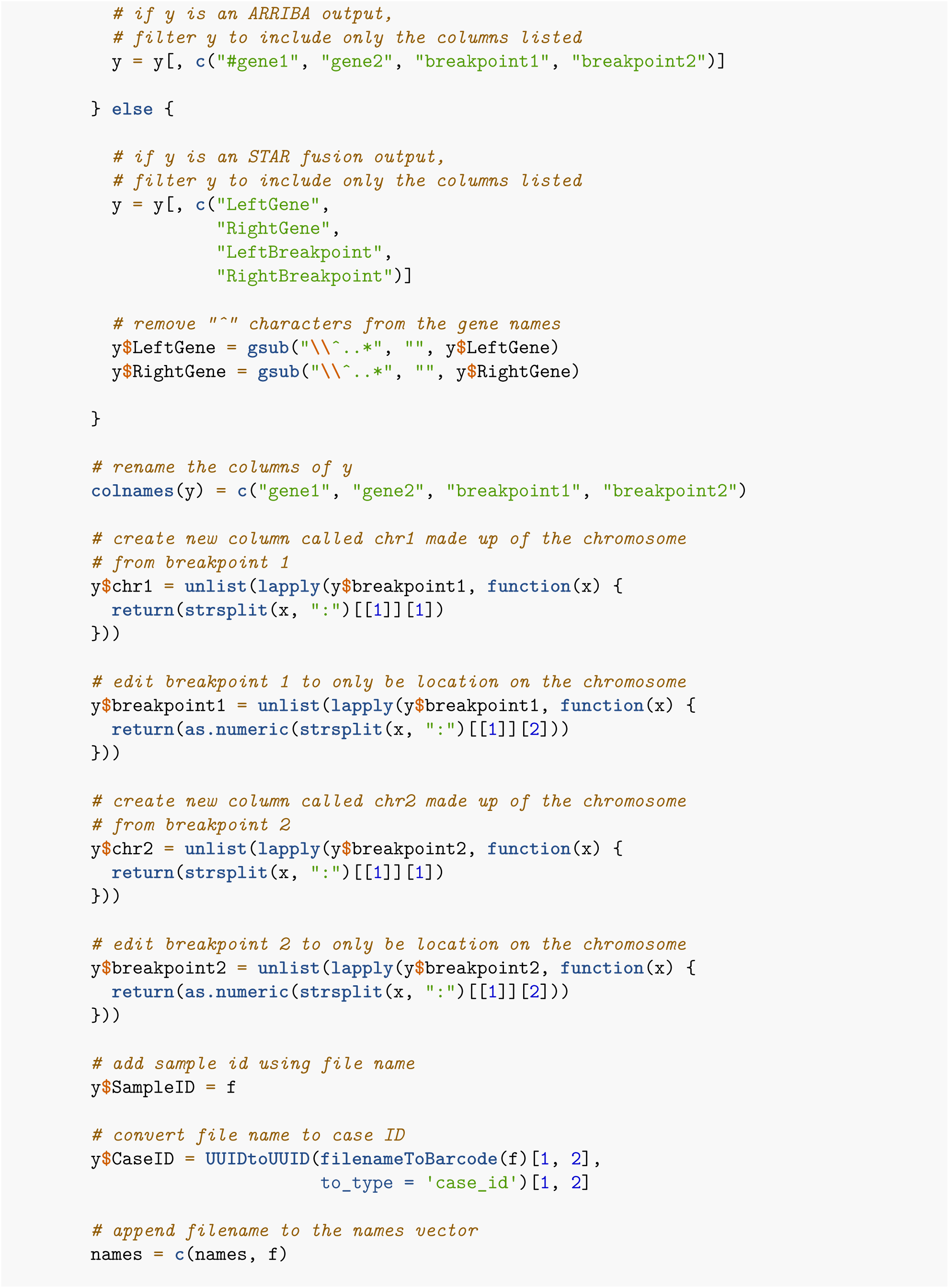

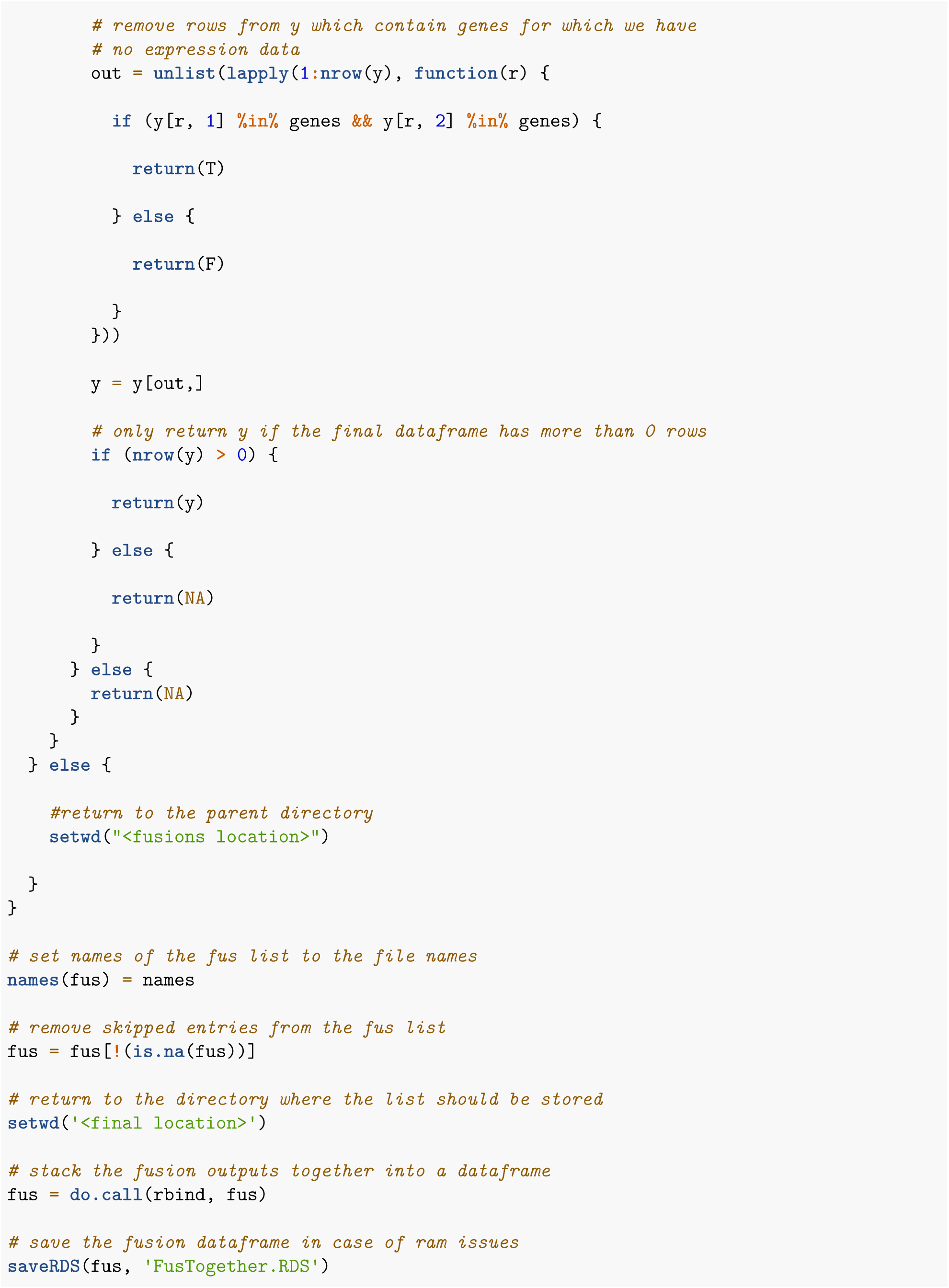

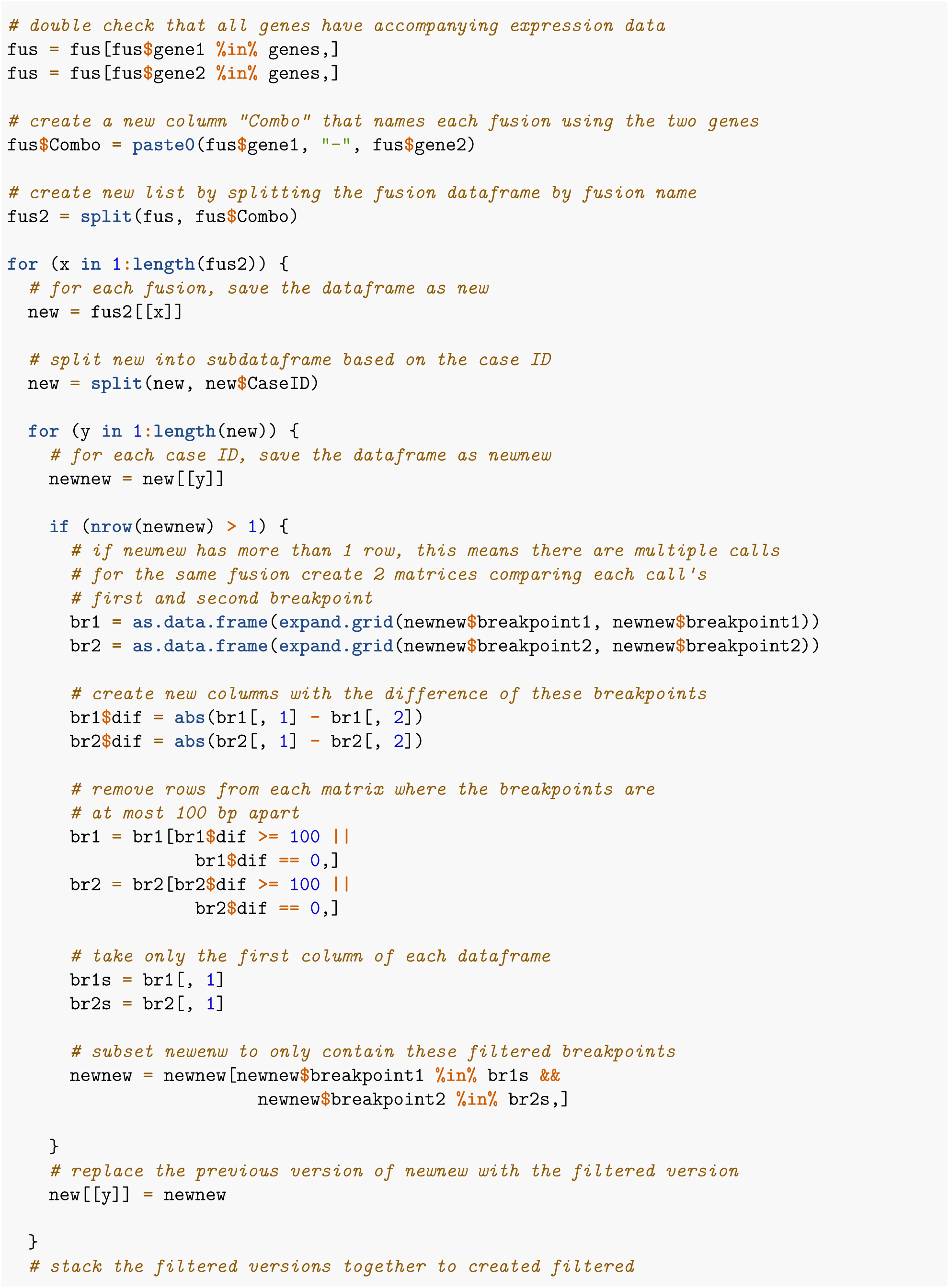

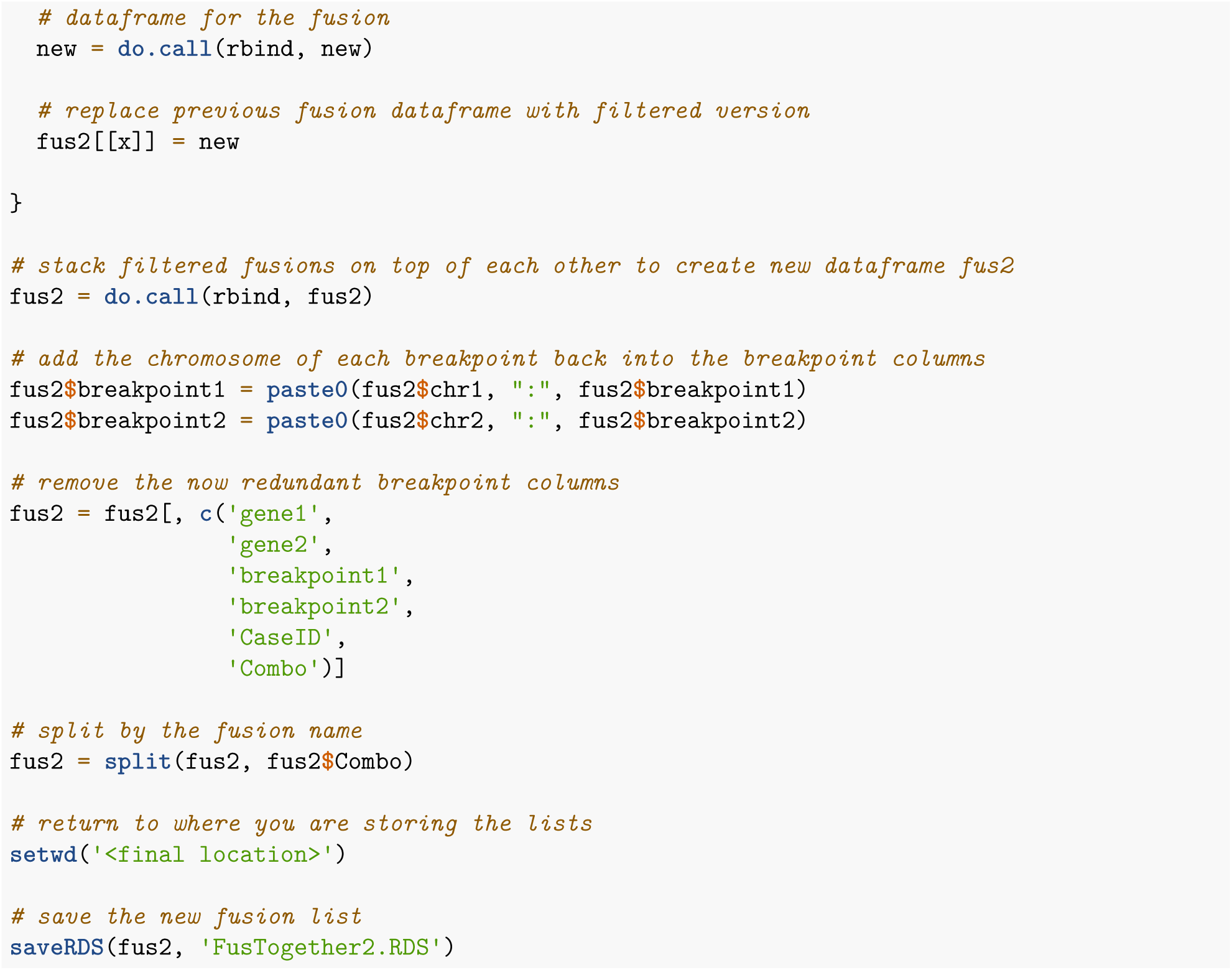

### 1.4: Process CNV outputs into a list

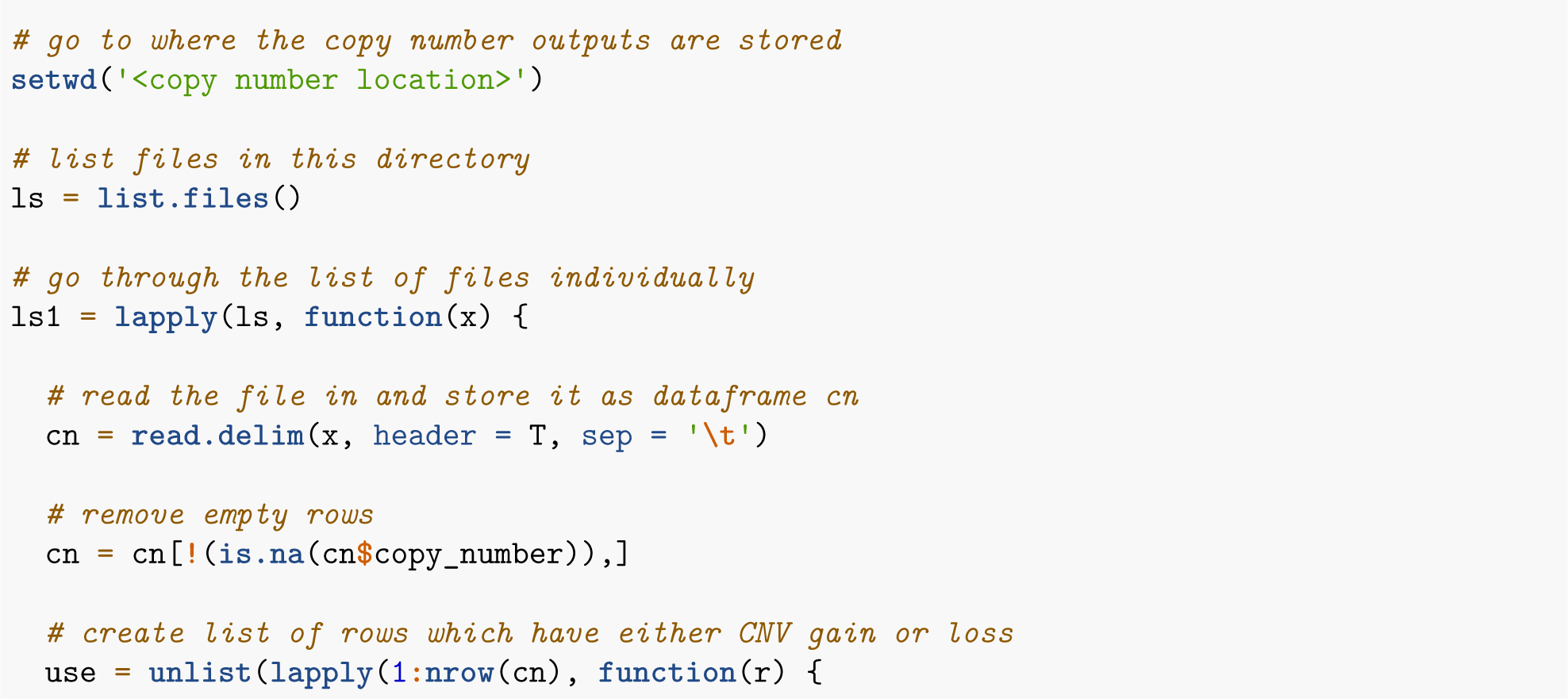

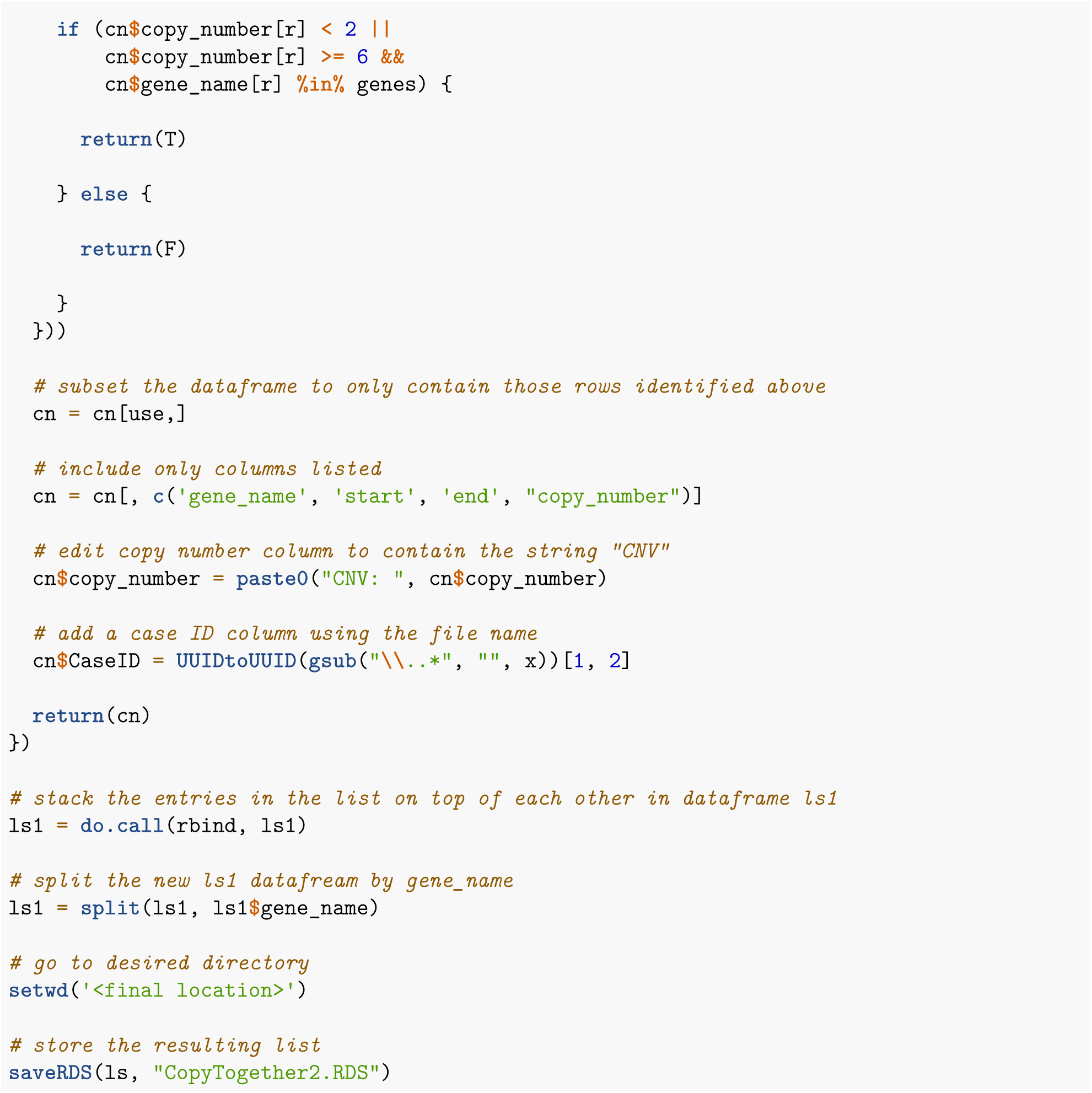

### 1.5: Get parliament2 results for whole genome samples

First you need to get the parliament2 image from dnanexus

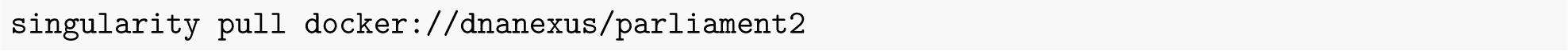

Next I wrote the following script to run parliament2 on all whole genome samples in parallel

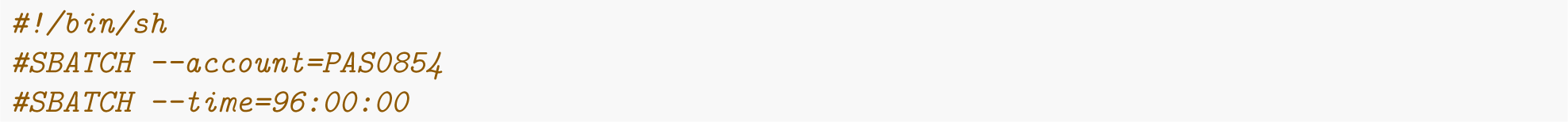

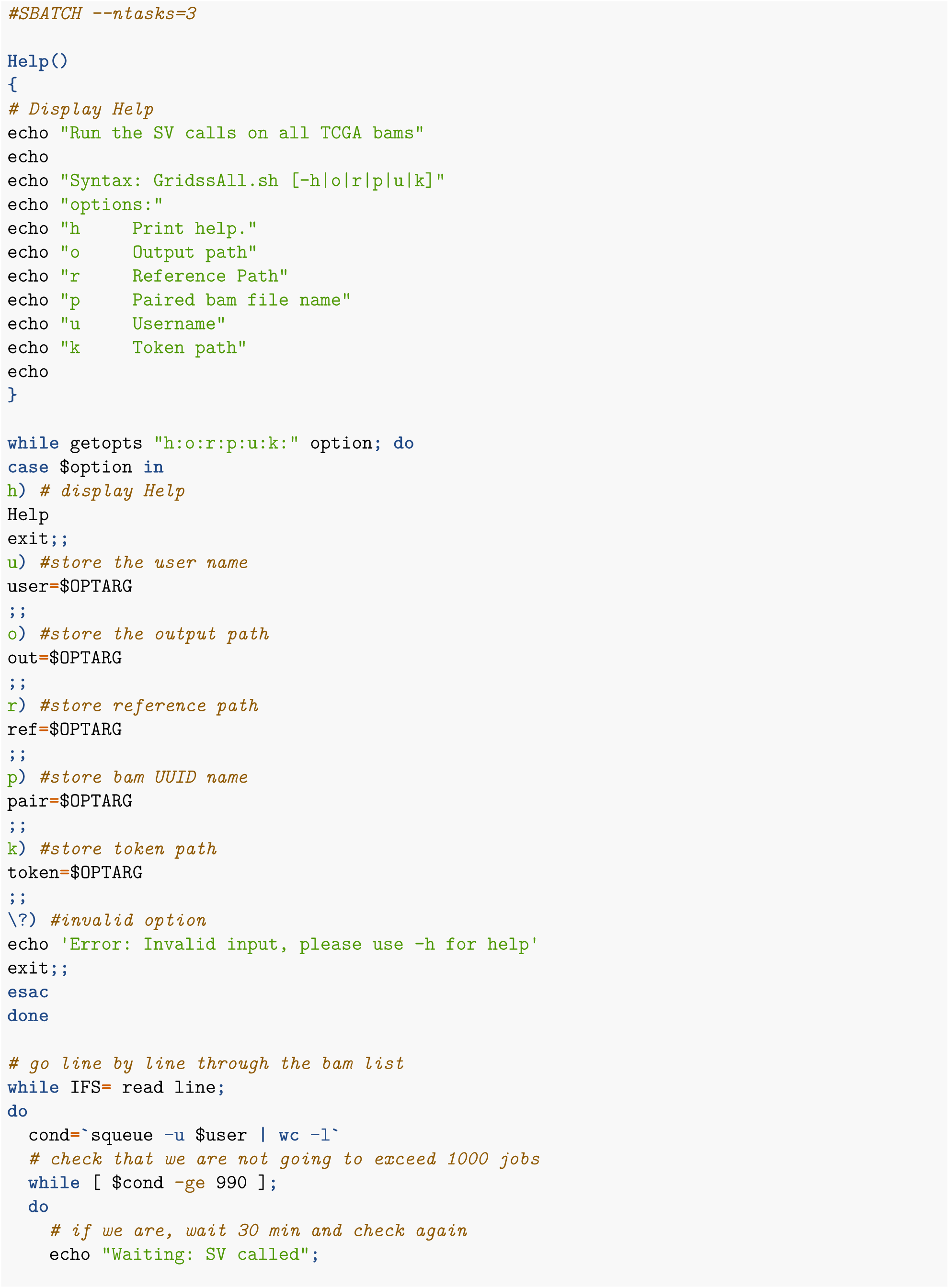

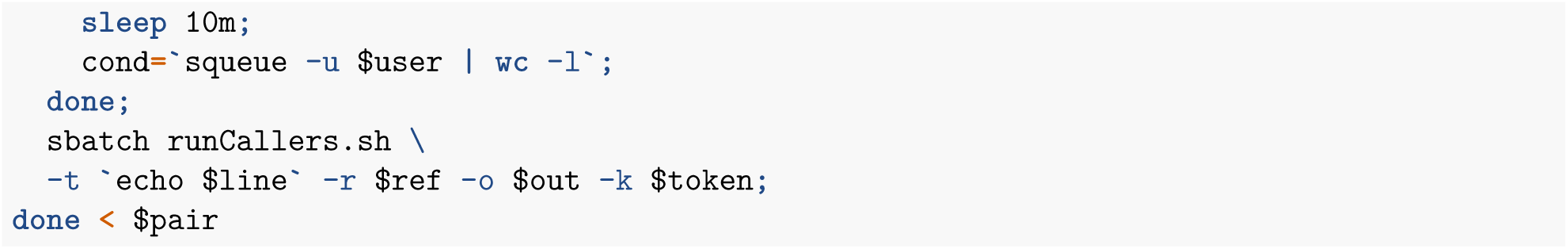

To run this script, I would use the line of code below (replaceing with the appropriate file paths).

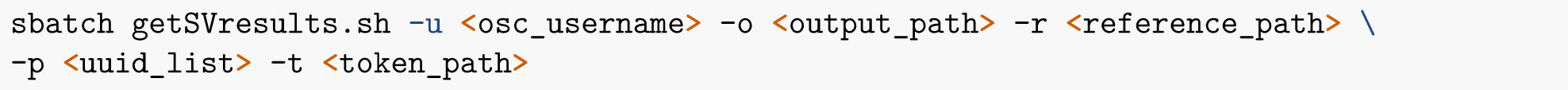

The runCallers.sh script is shown below.

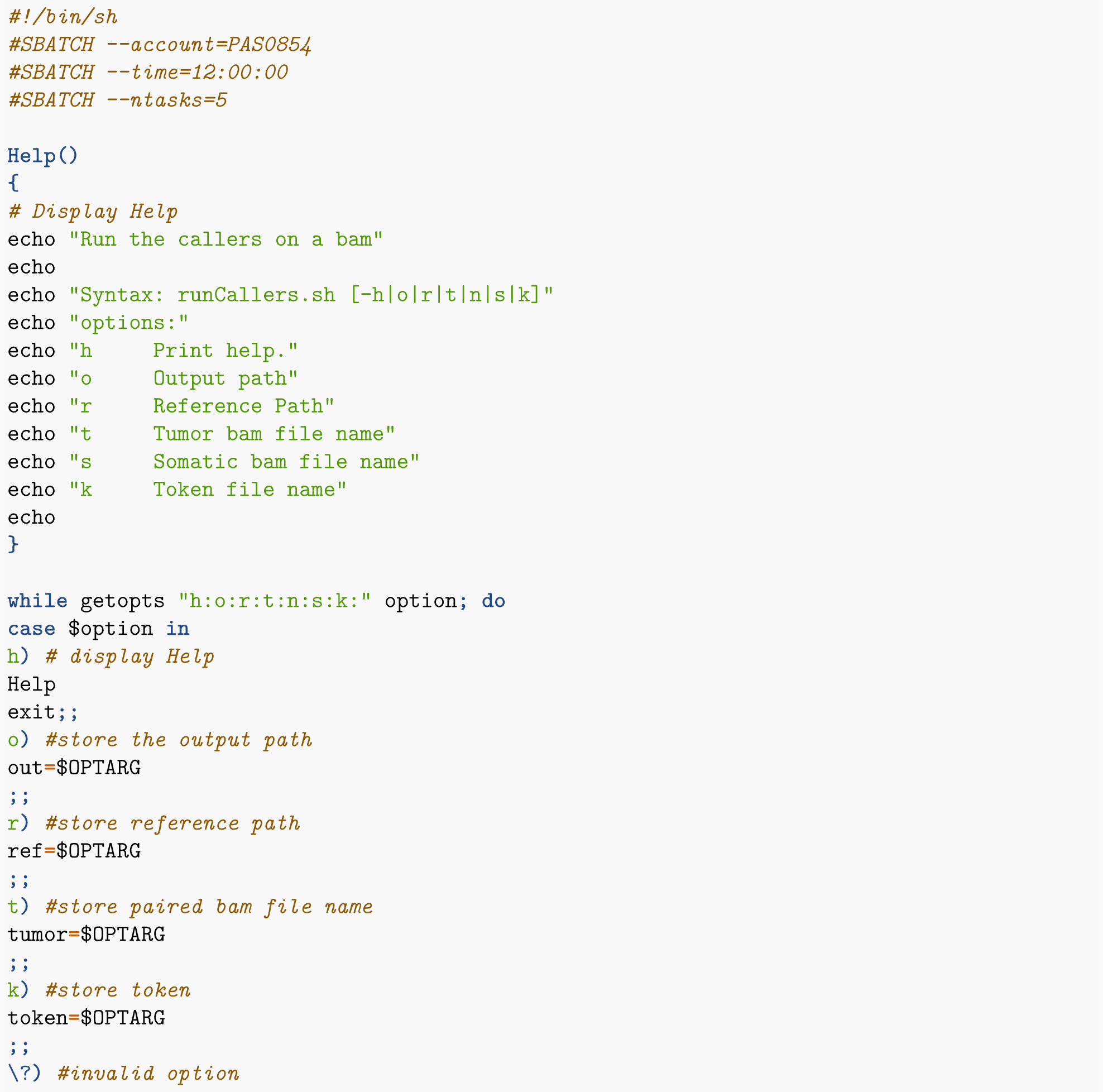

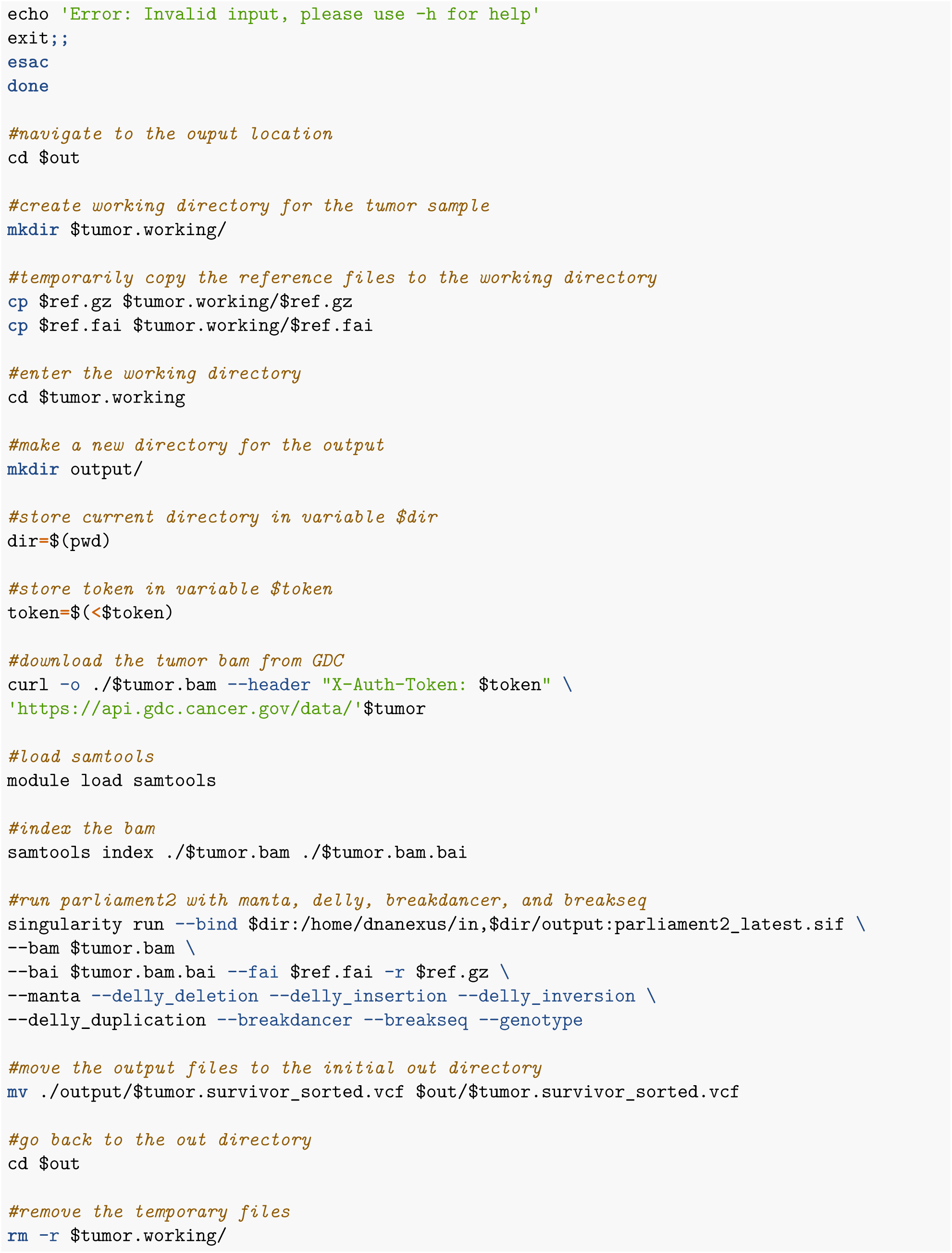

### 1.6 : Process SV outputs into a list

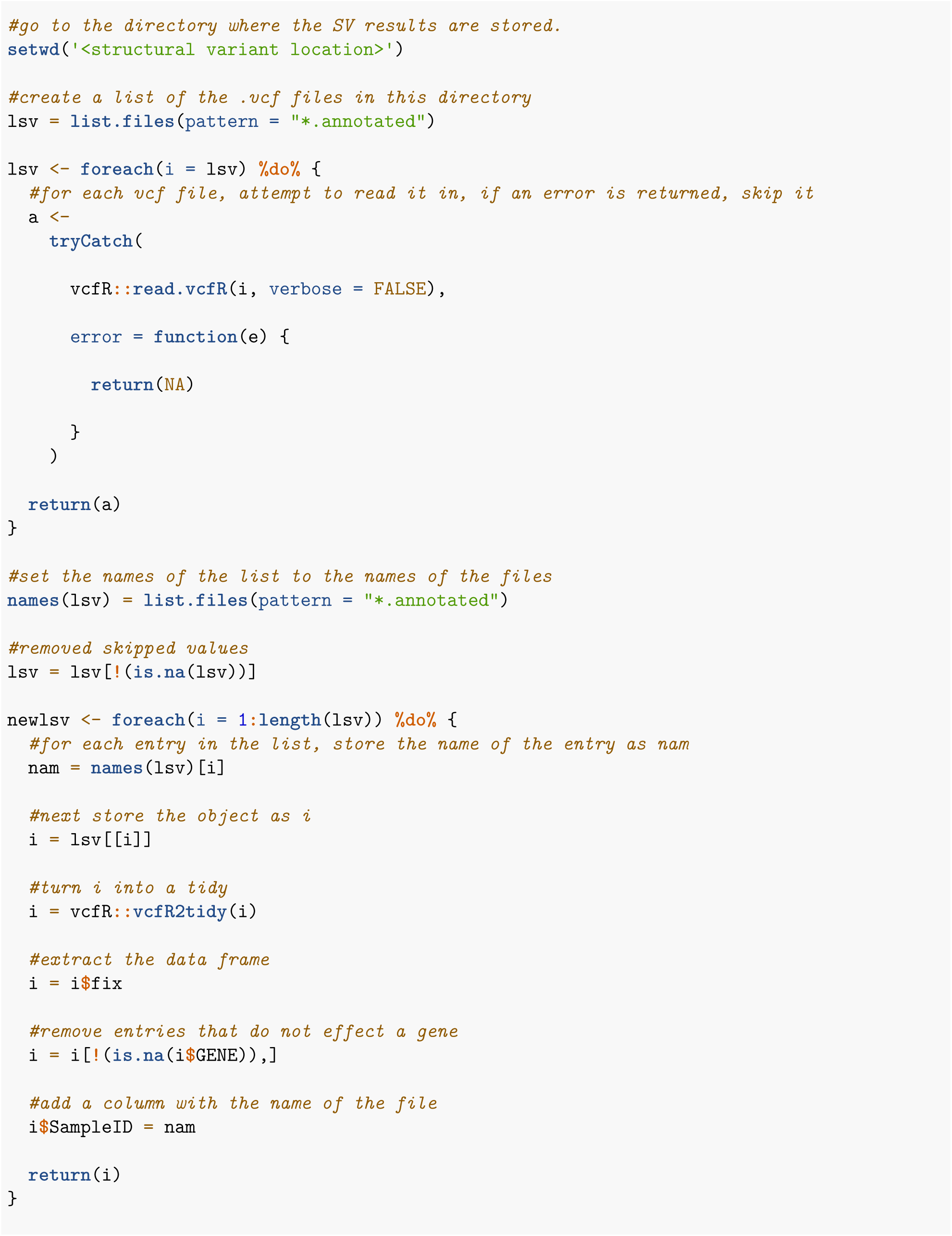

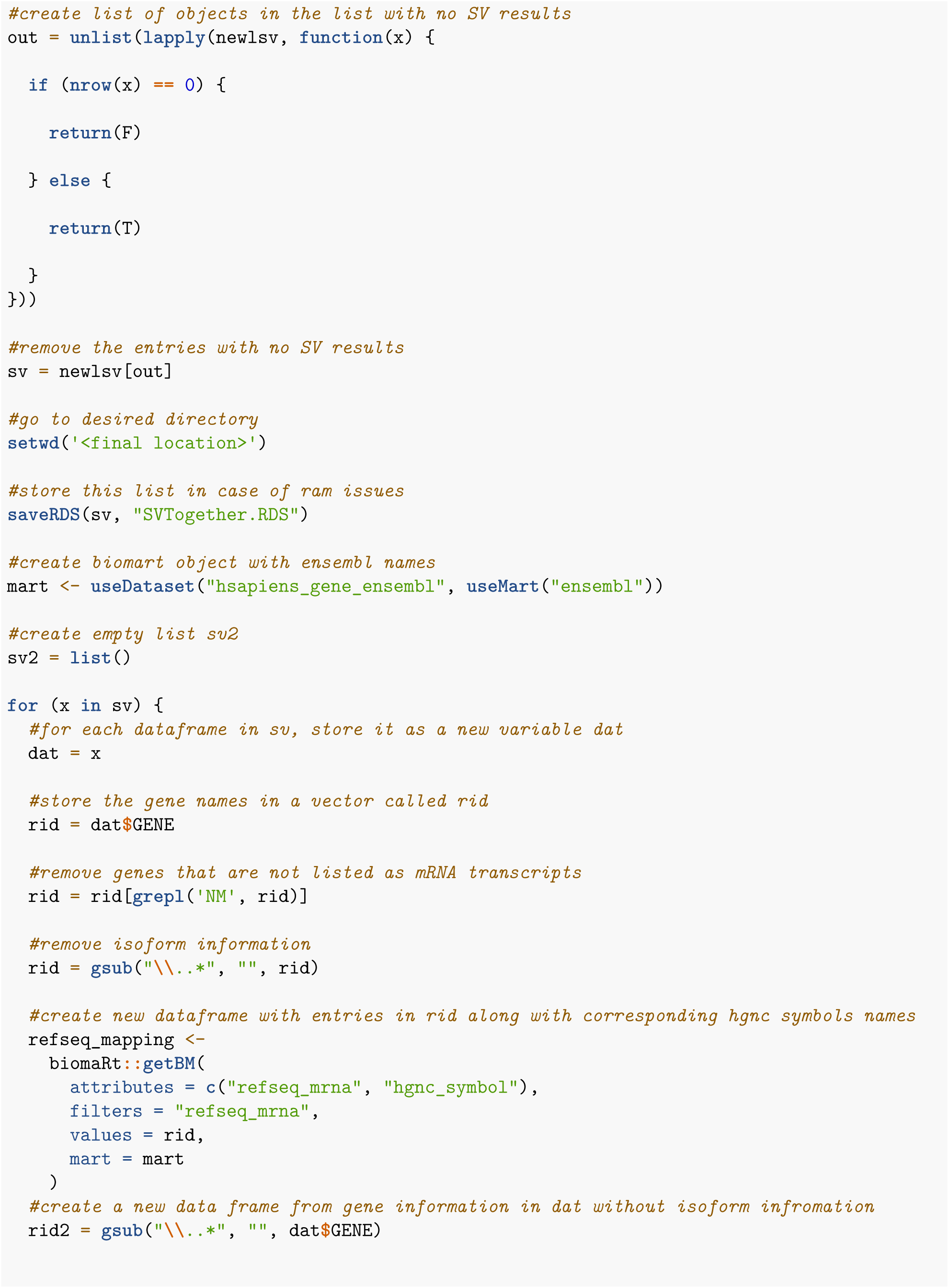

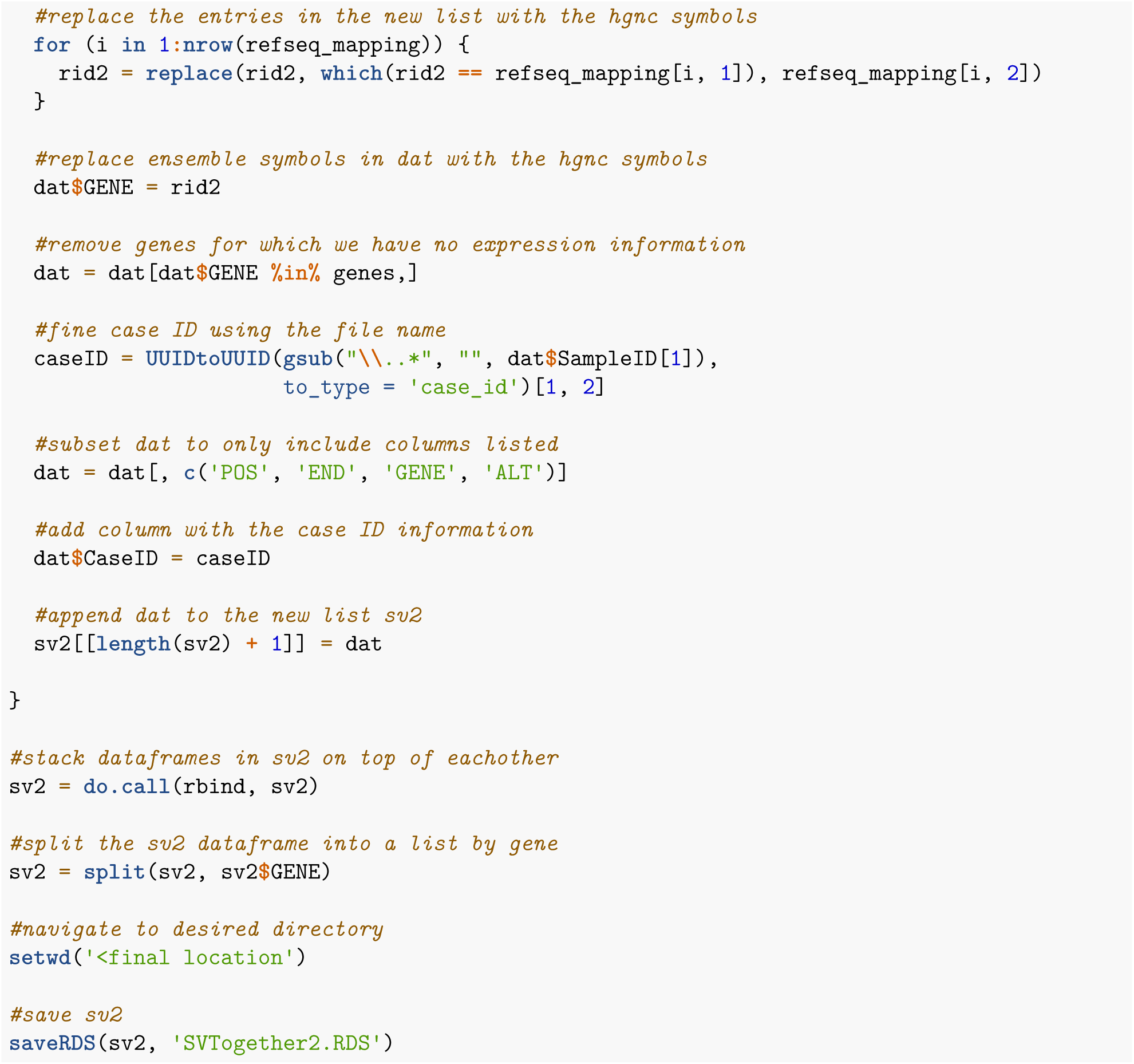

## Part 2: Batch Correction on TCGA expression files

In order to analyze the RNA seq data all together from TCGA, batch correction was performed by institution. First all RNA expression files must be downloaded. To do this, I went to GDC, created a manifest and downloaded them as shown below.

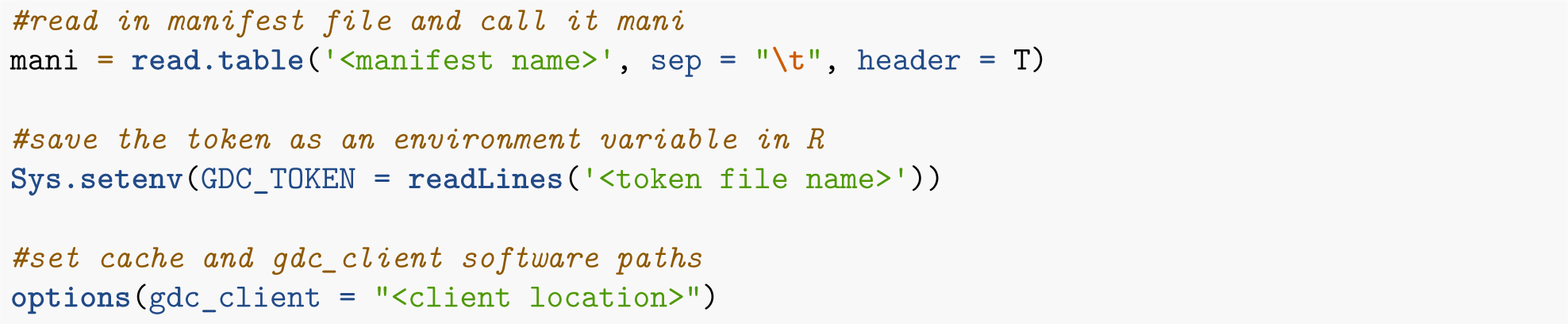

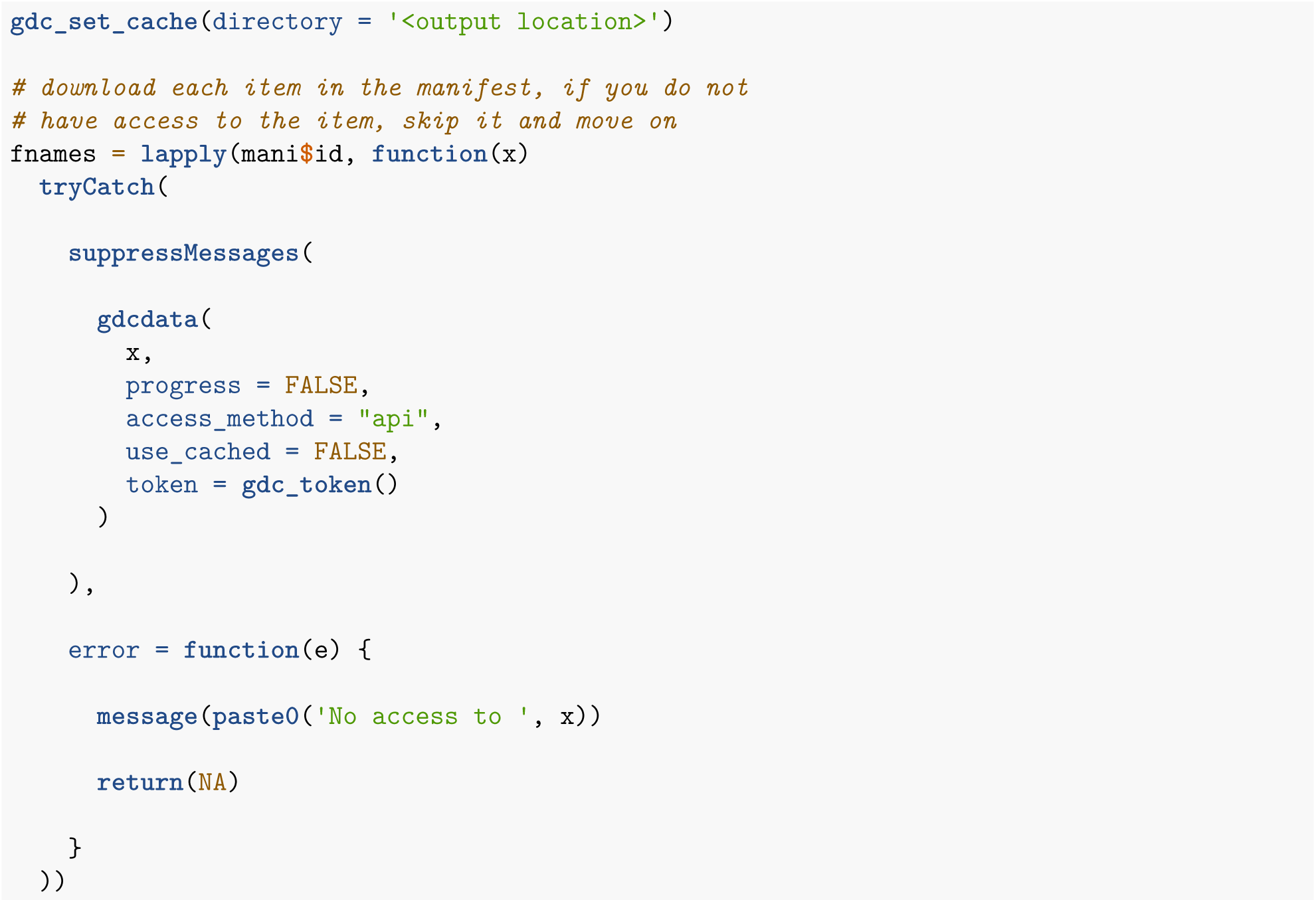

Next, you need to unpack all the directories.

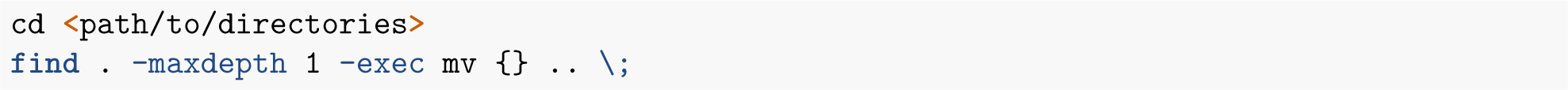

Next, I ran combat-seq from SVA as shown below.

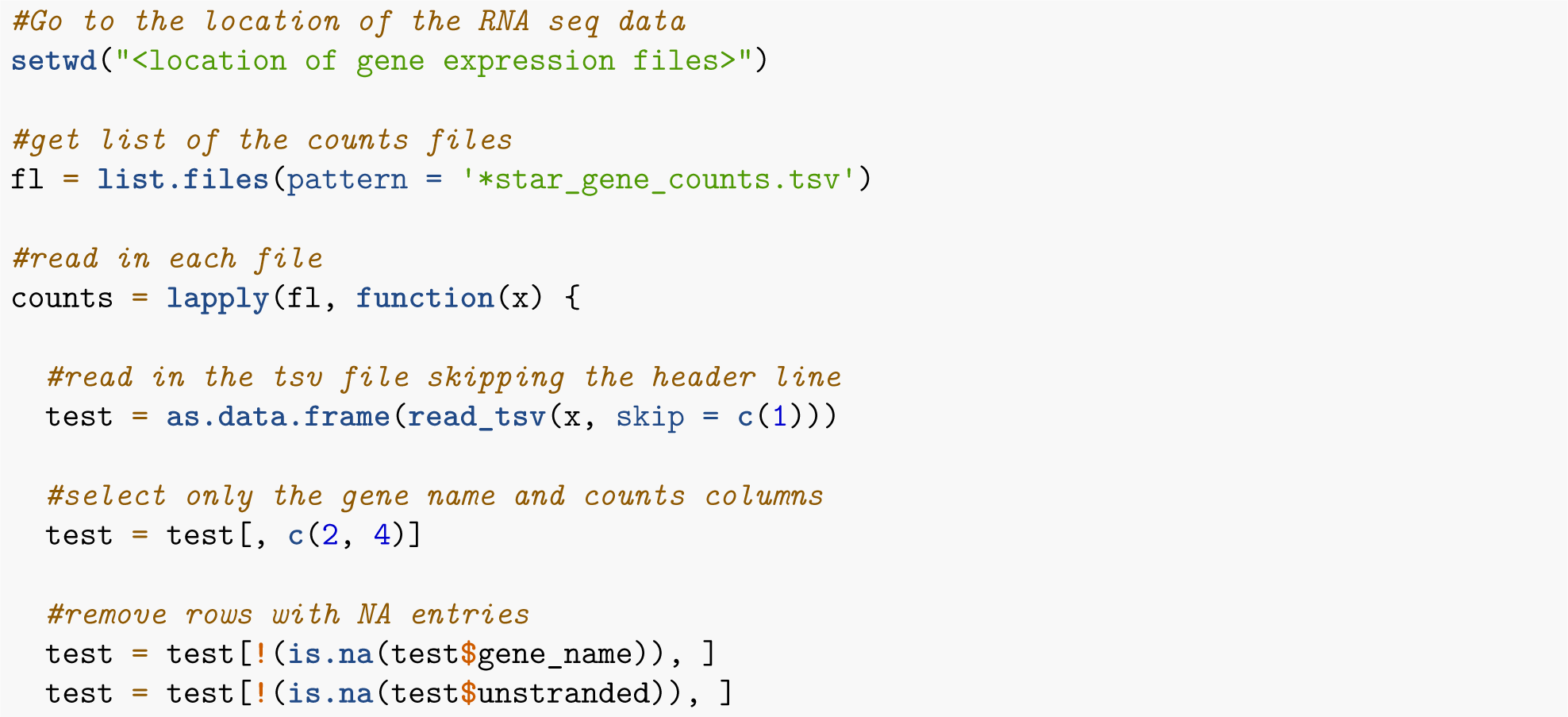

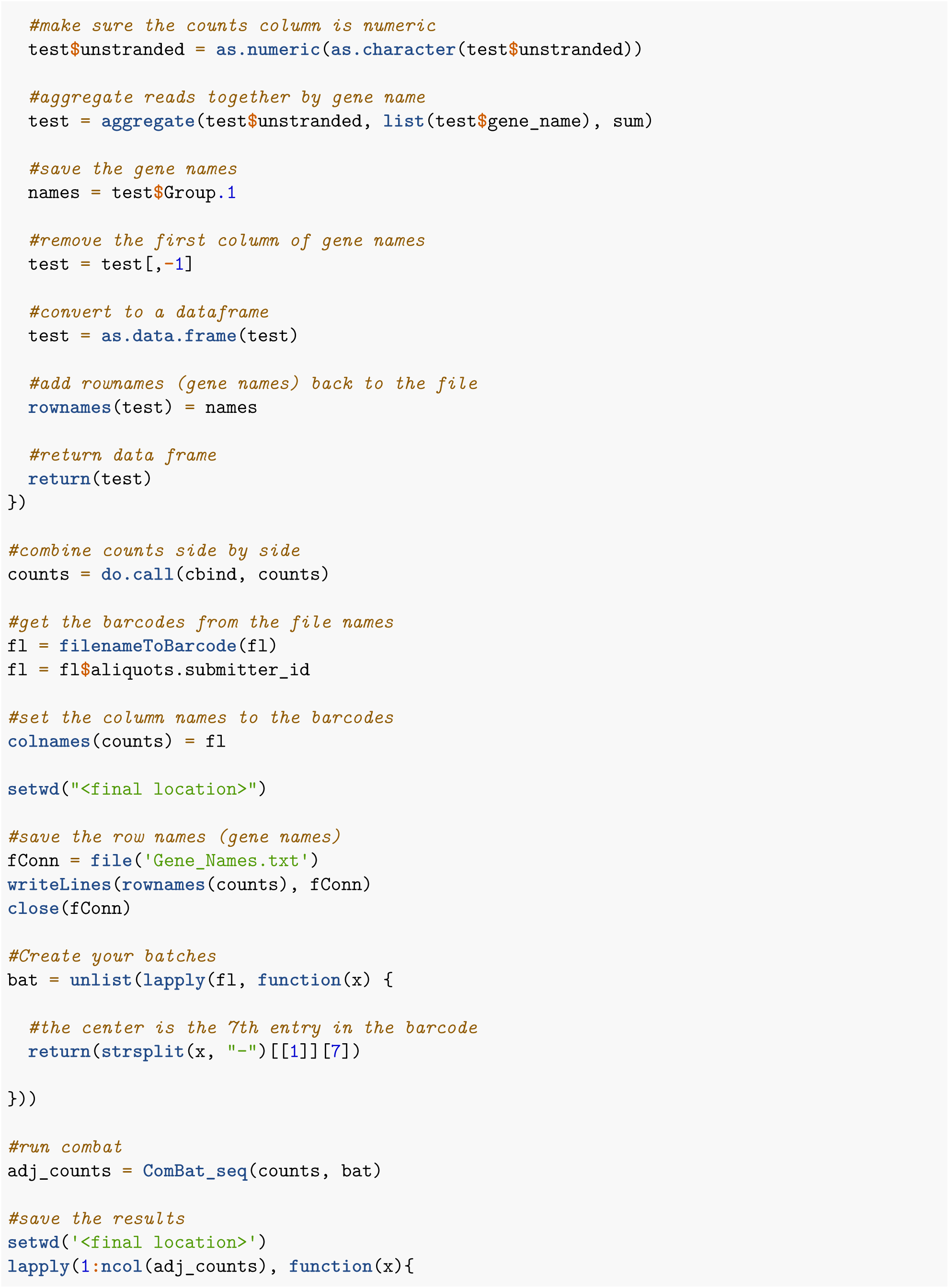

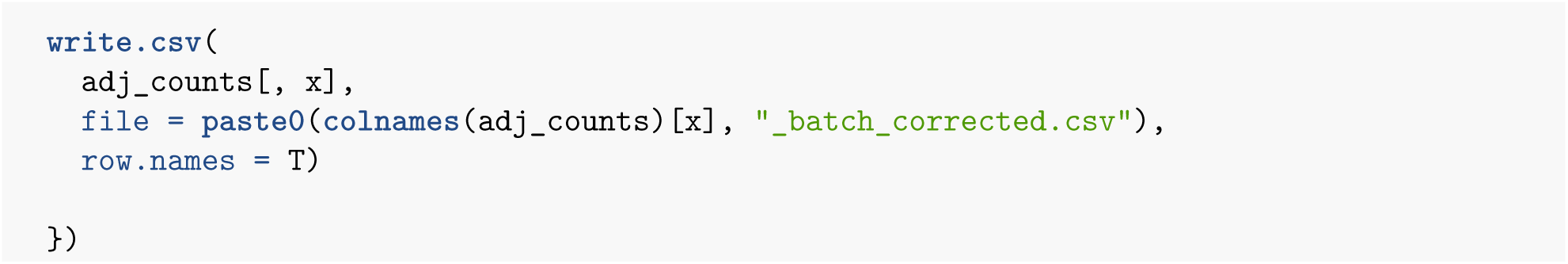

## Part 3: Build hot/cold detection algorithm

### 3.1: Preprocessing input data

First I combine the pathologist annotations and gather the paired RNA for each sample.

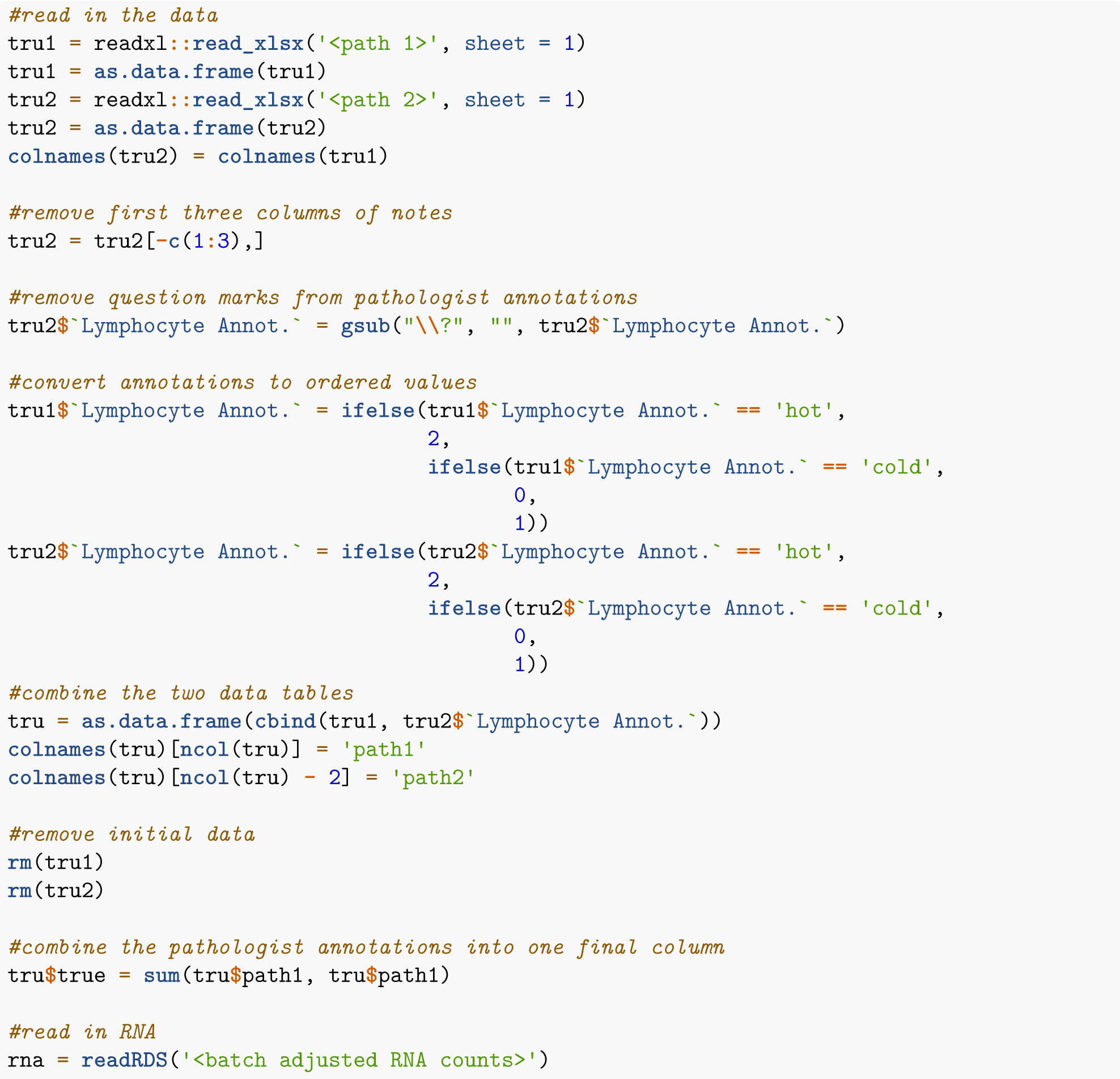

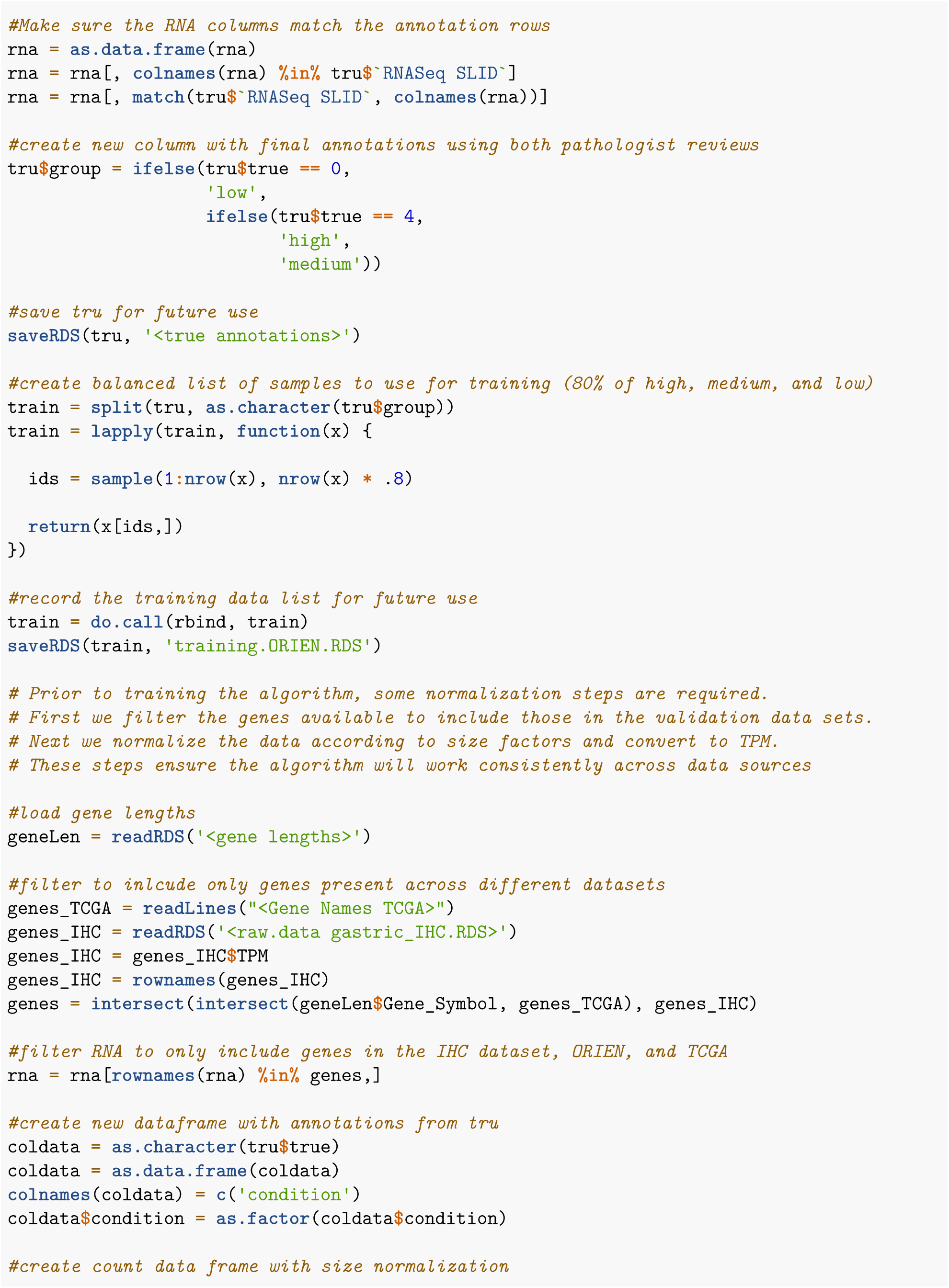

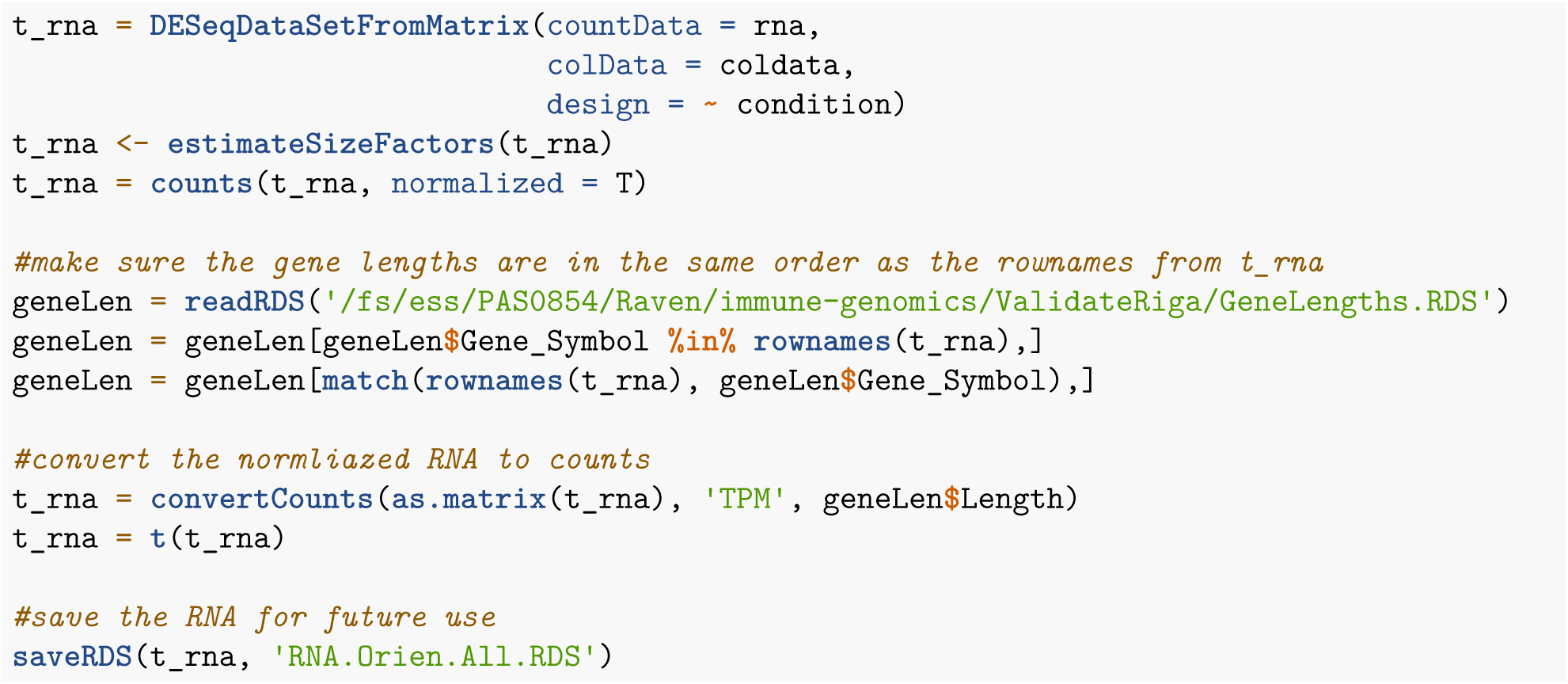

### 3.2 ElasticNet feature selection

Next we will select features for our machine learning algorithm using elastic net. This was done for all 3 types of annotations (pathologist 1, pathologist 2, and the combination.)

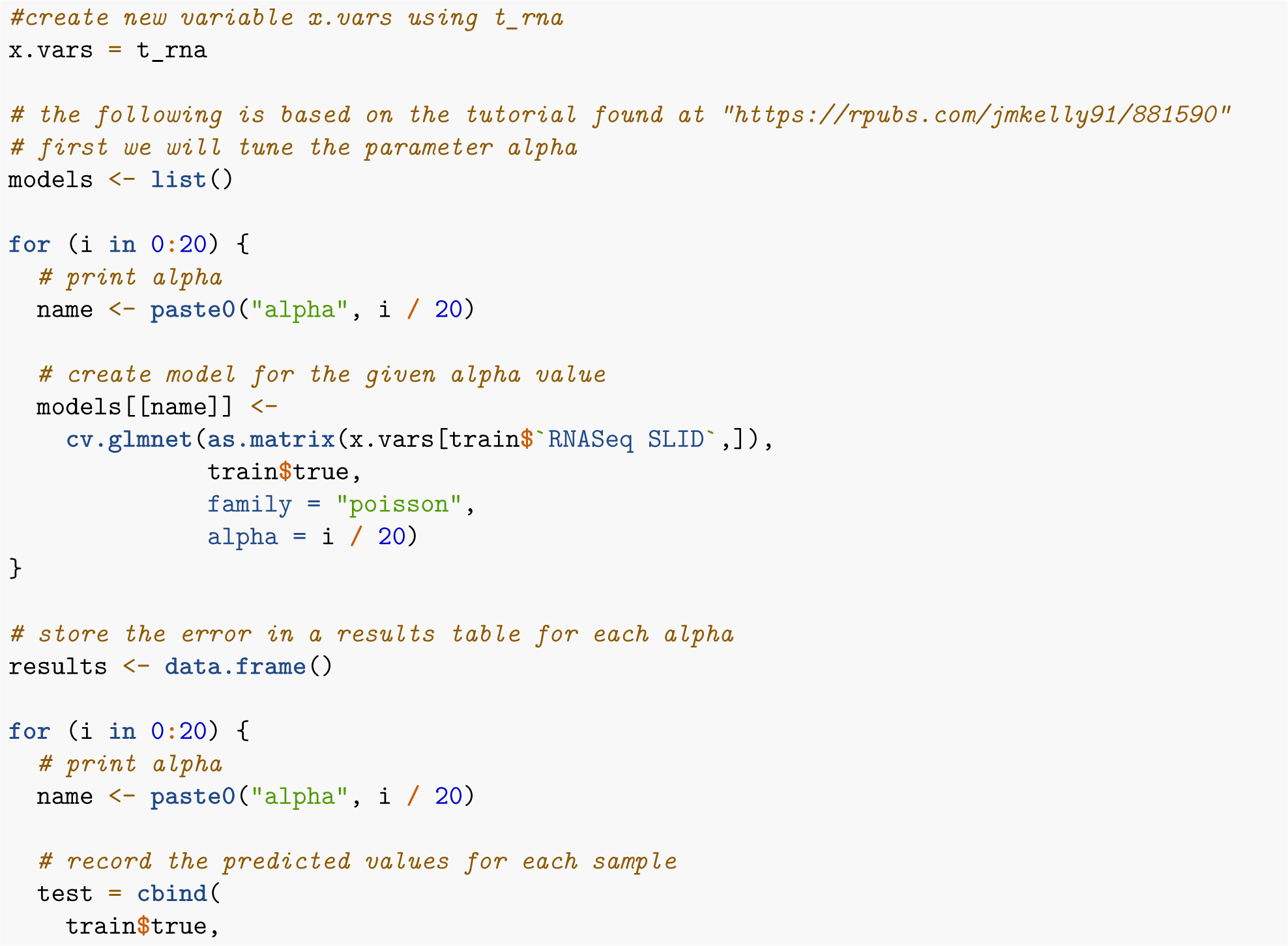

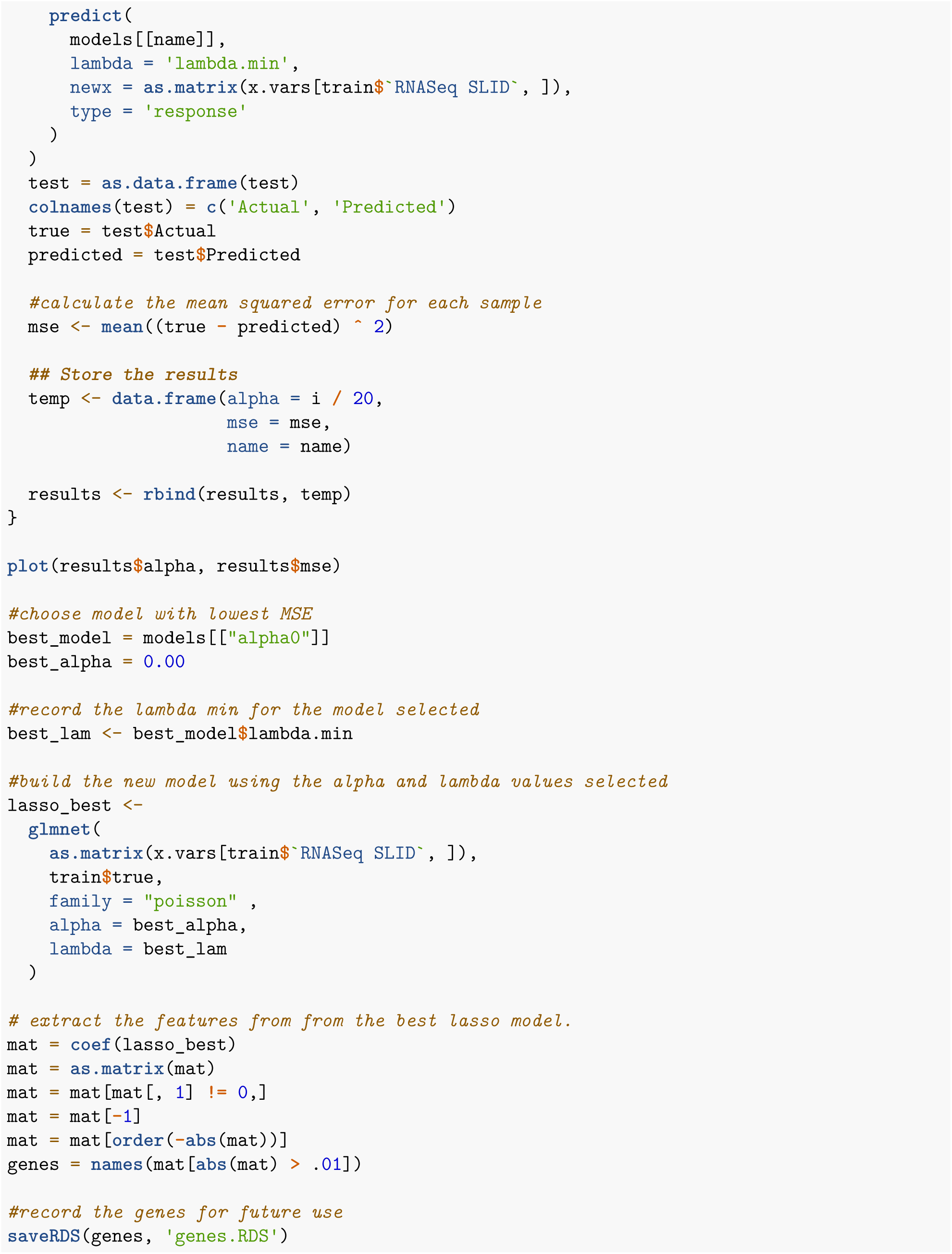

### 3.3 OrdinalForest Machine learning

After all the genes are collected, we built two kinds of machine learning models for each annotation: OrdinalForest and XgBoost. The process for OrdinalForest is shown below

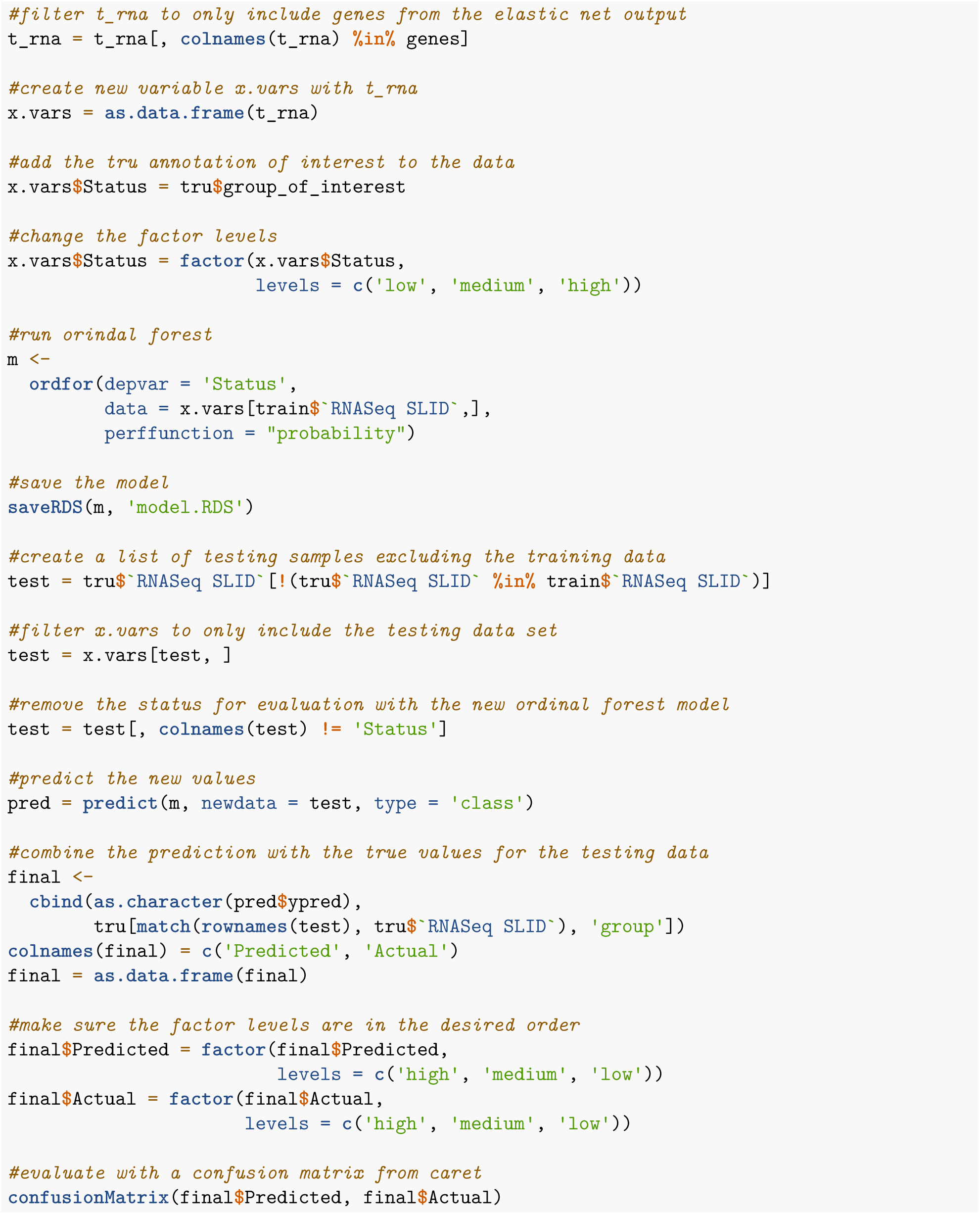

### 3.4 XgBoost machine learning

Below is shown the process for machine learning with XgBoost. This is based on a tutorial found at “https://www.r-bloggers.com/2022/01/using-bayesian-optimisation-to-tune-a-xgboost-model-in-r/”

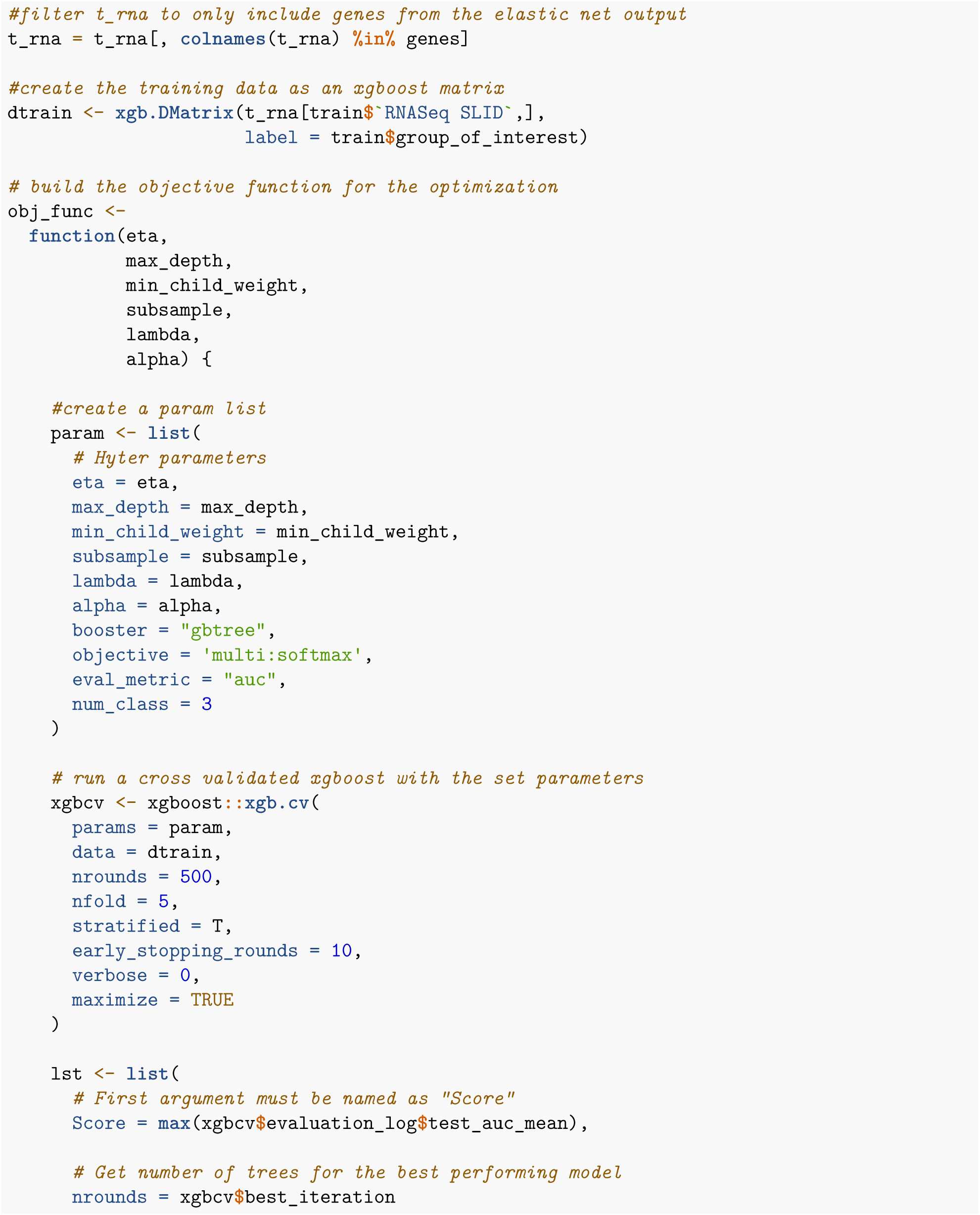

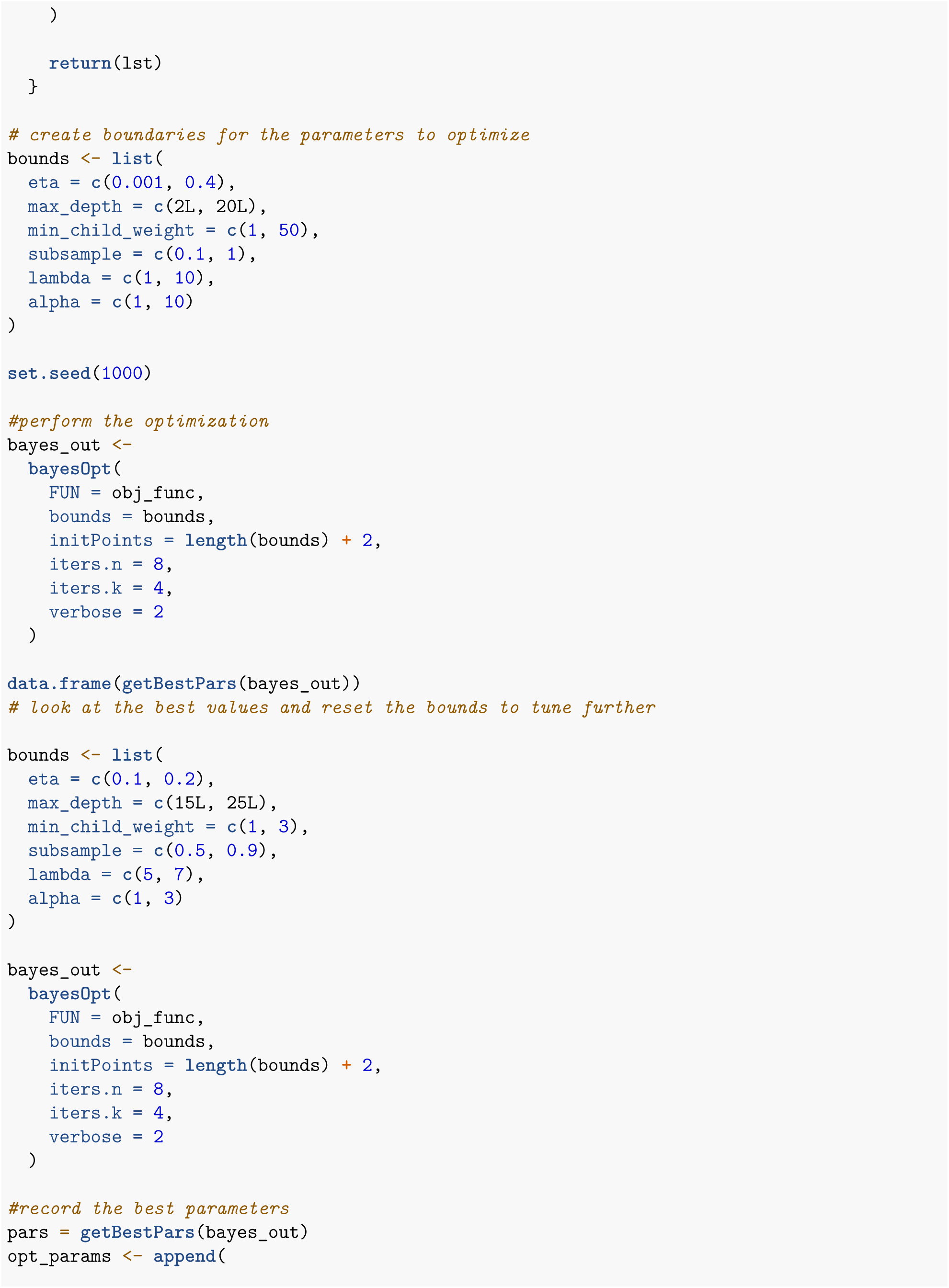

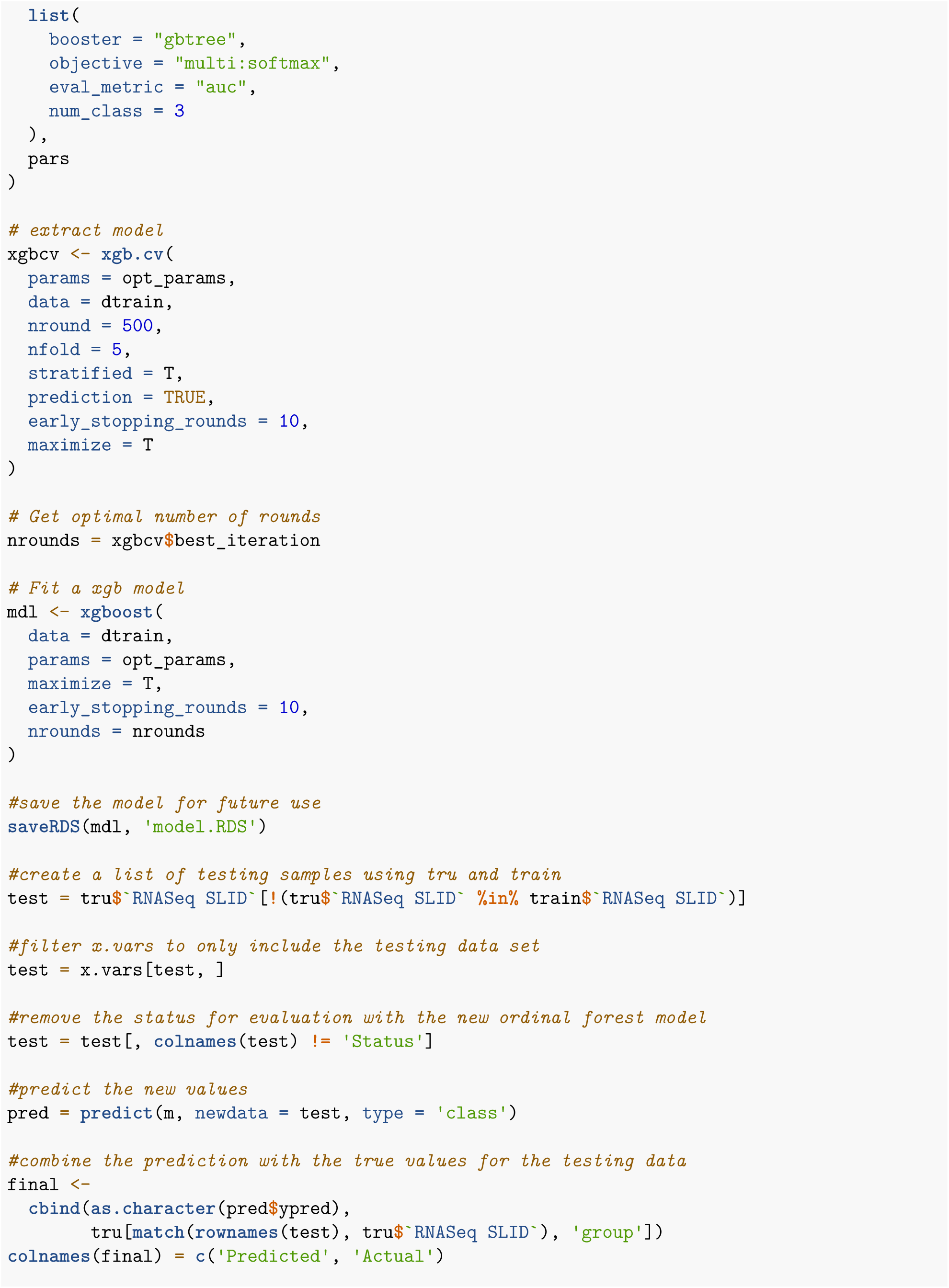

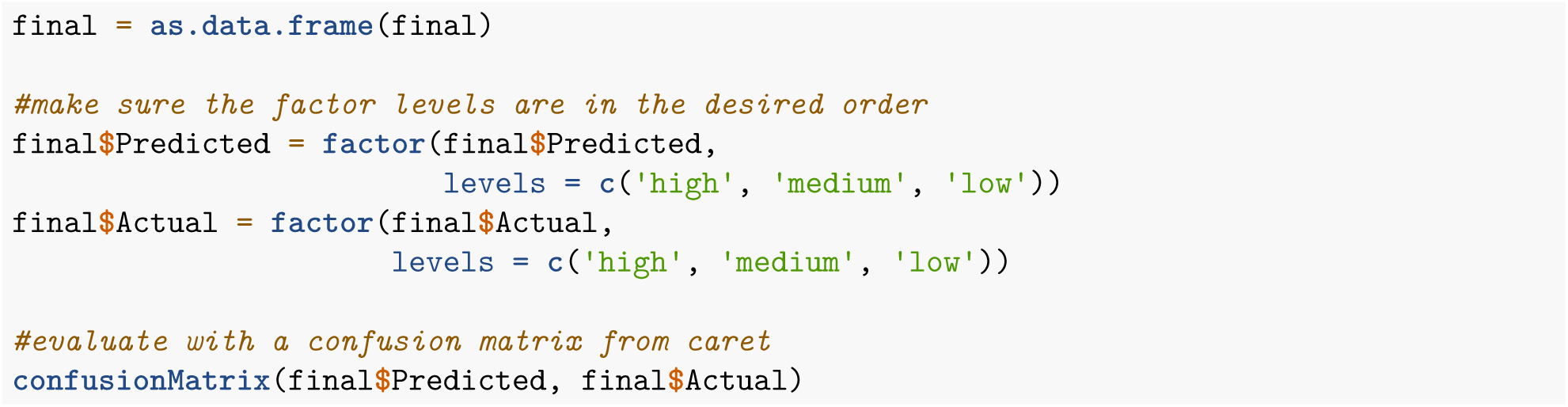

All 6 models were saved and ultimately model 6 was chosen because of its performance in the testing data.

## Part 4: Build RIGATONI package functions

We have all the elements needed to put together the RIGATONI r package. Now we must put them together to build functions that will be useful to our users.

### 4.1: Download STRING database connections

First we download the string protein-protein interaction database (https://string-db.org/) and change the gene names

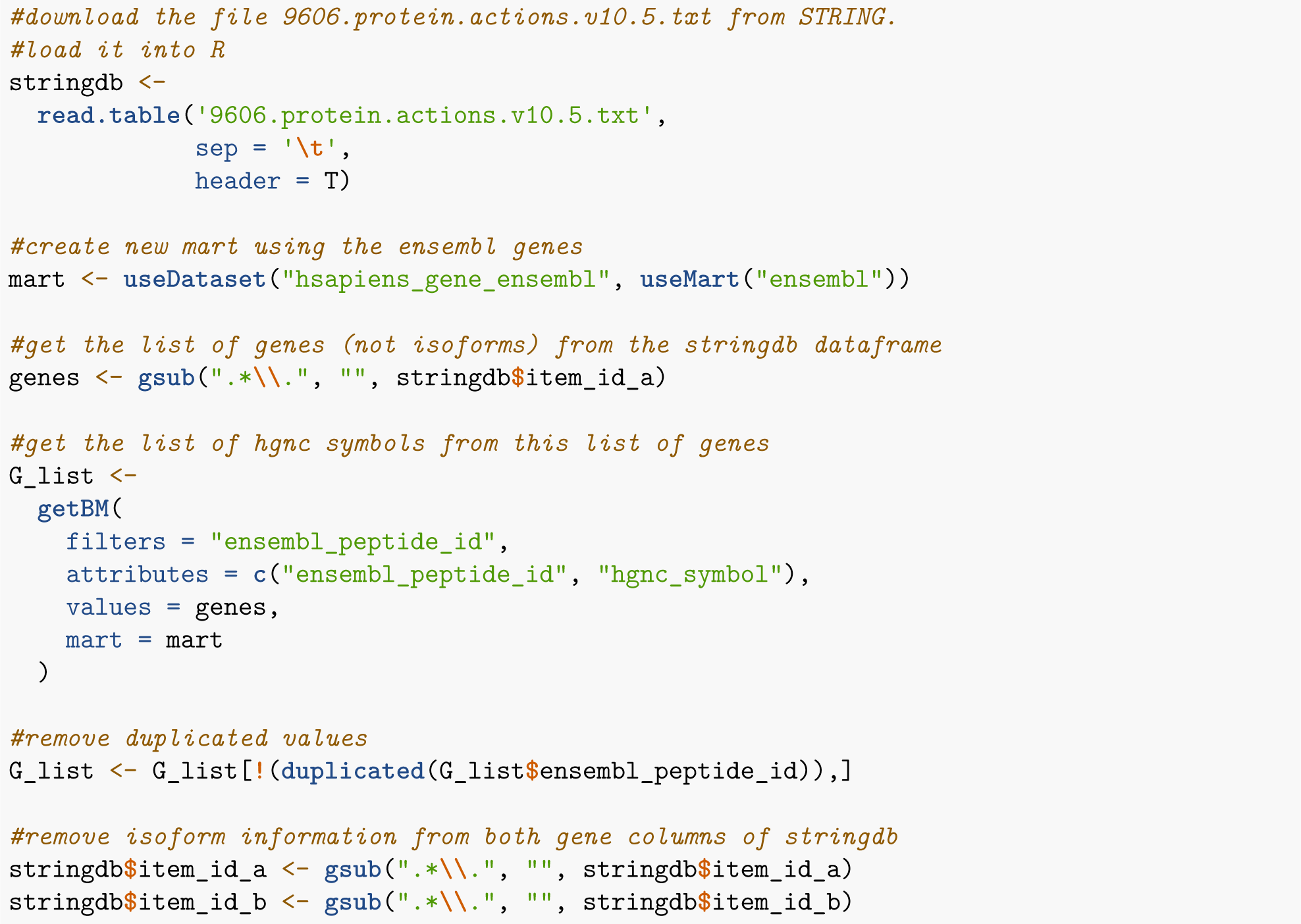

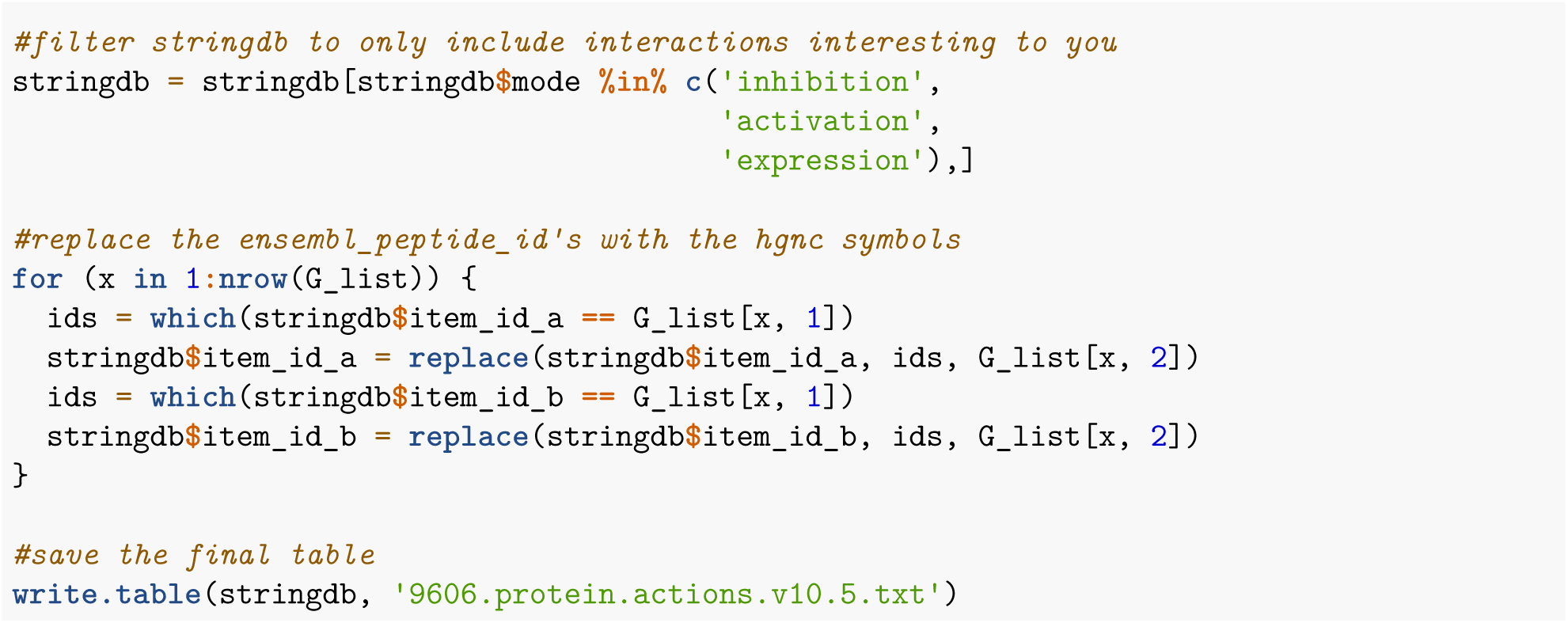

After this, we need a way to filter the STRING output for a particular gene.

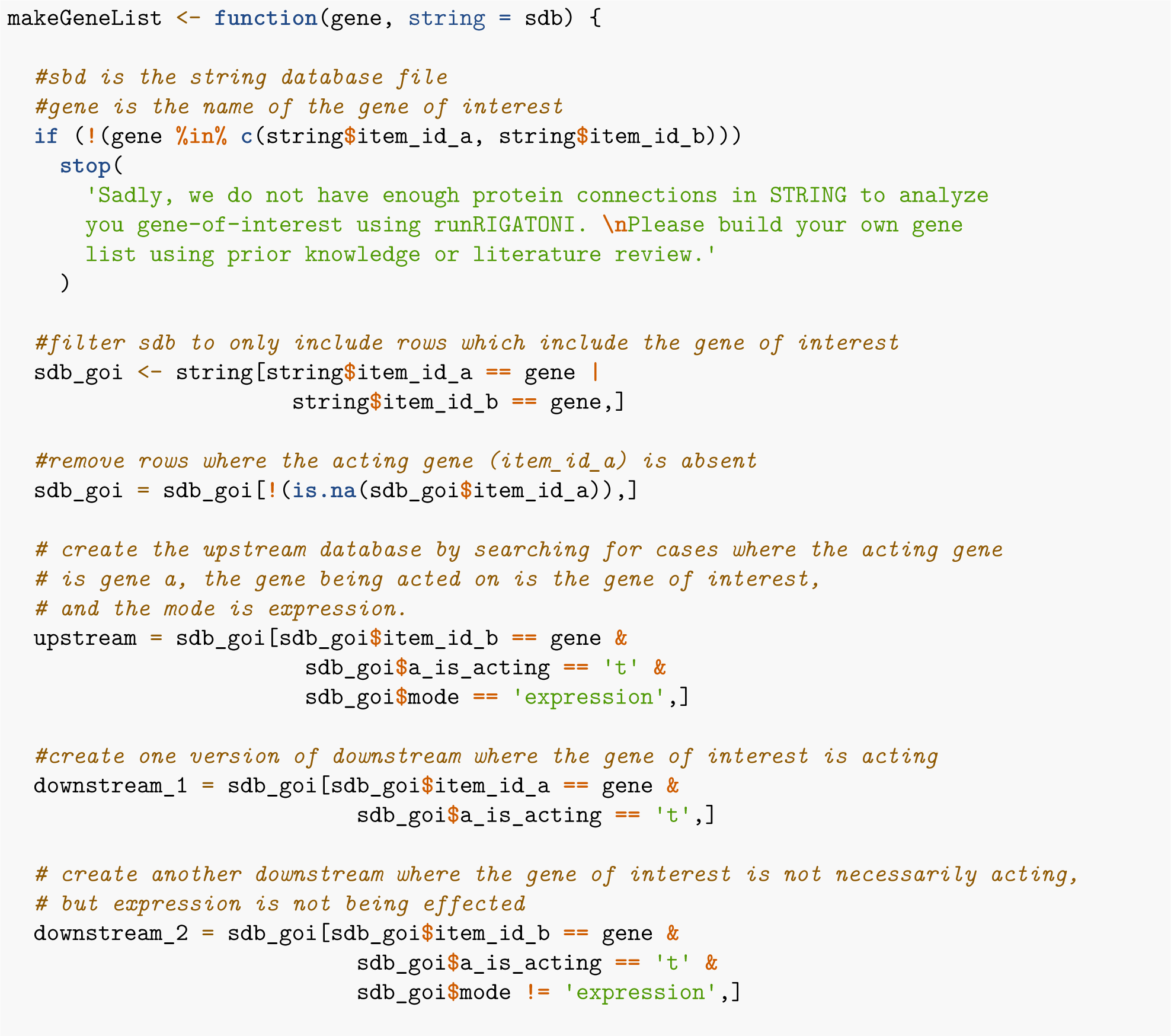

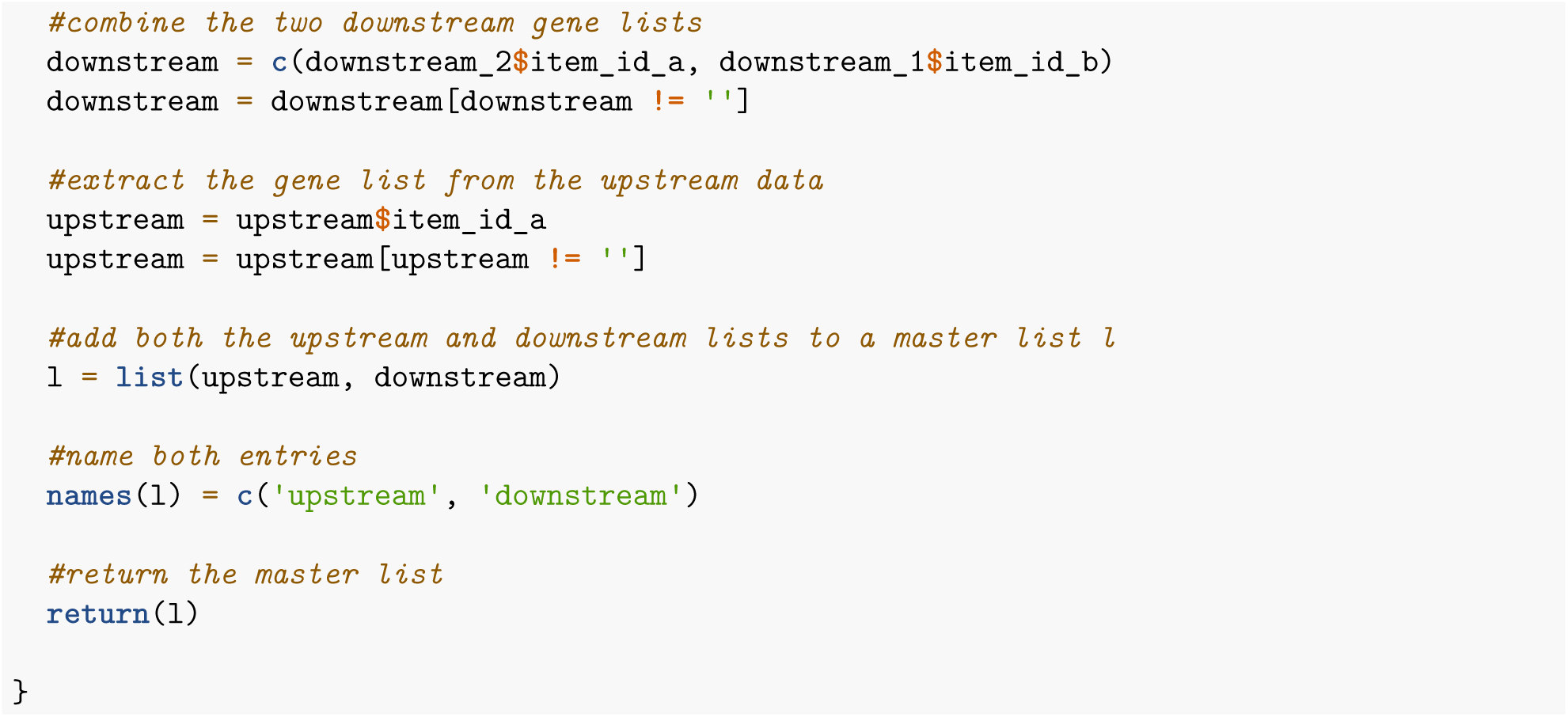

### 4.2: Create the regression models themselves

After this, we need to create the regression models for the gene of interest using the control data.

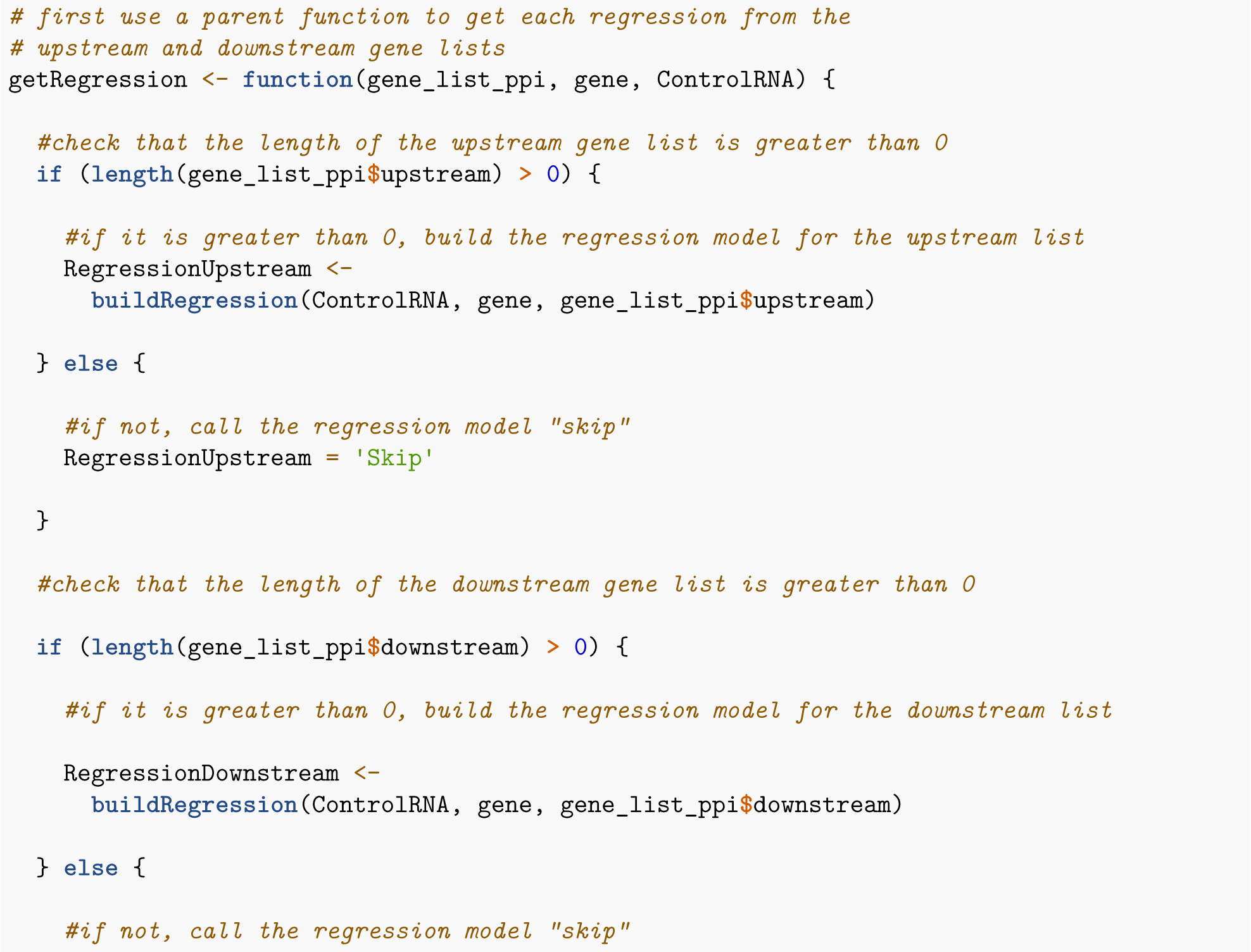

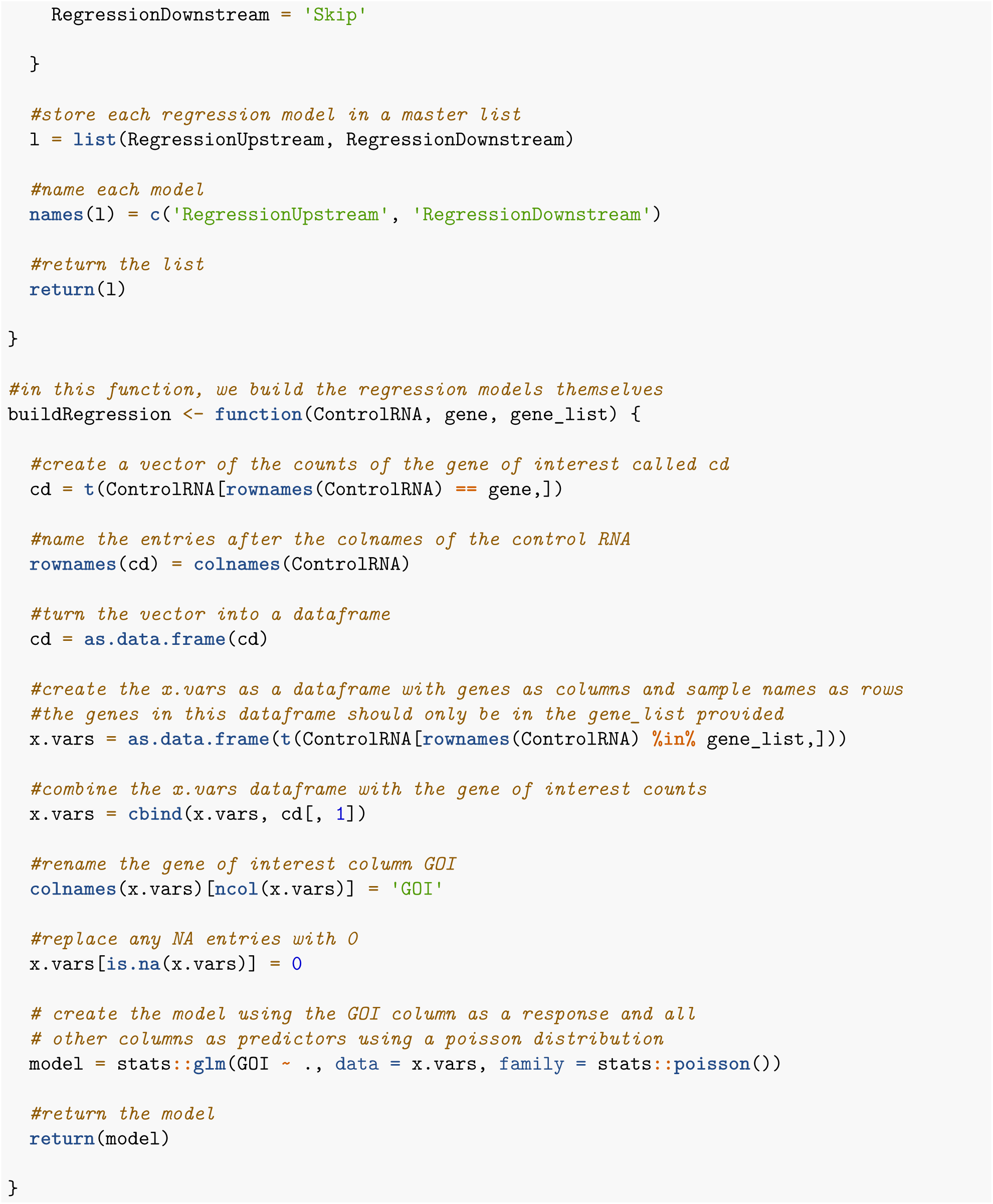

### 4.3: Predict the function of each mutant sample

After creating the models, we need to use the models to predict the functional status of each mutant sample

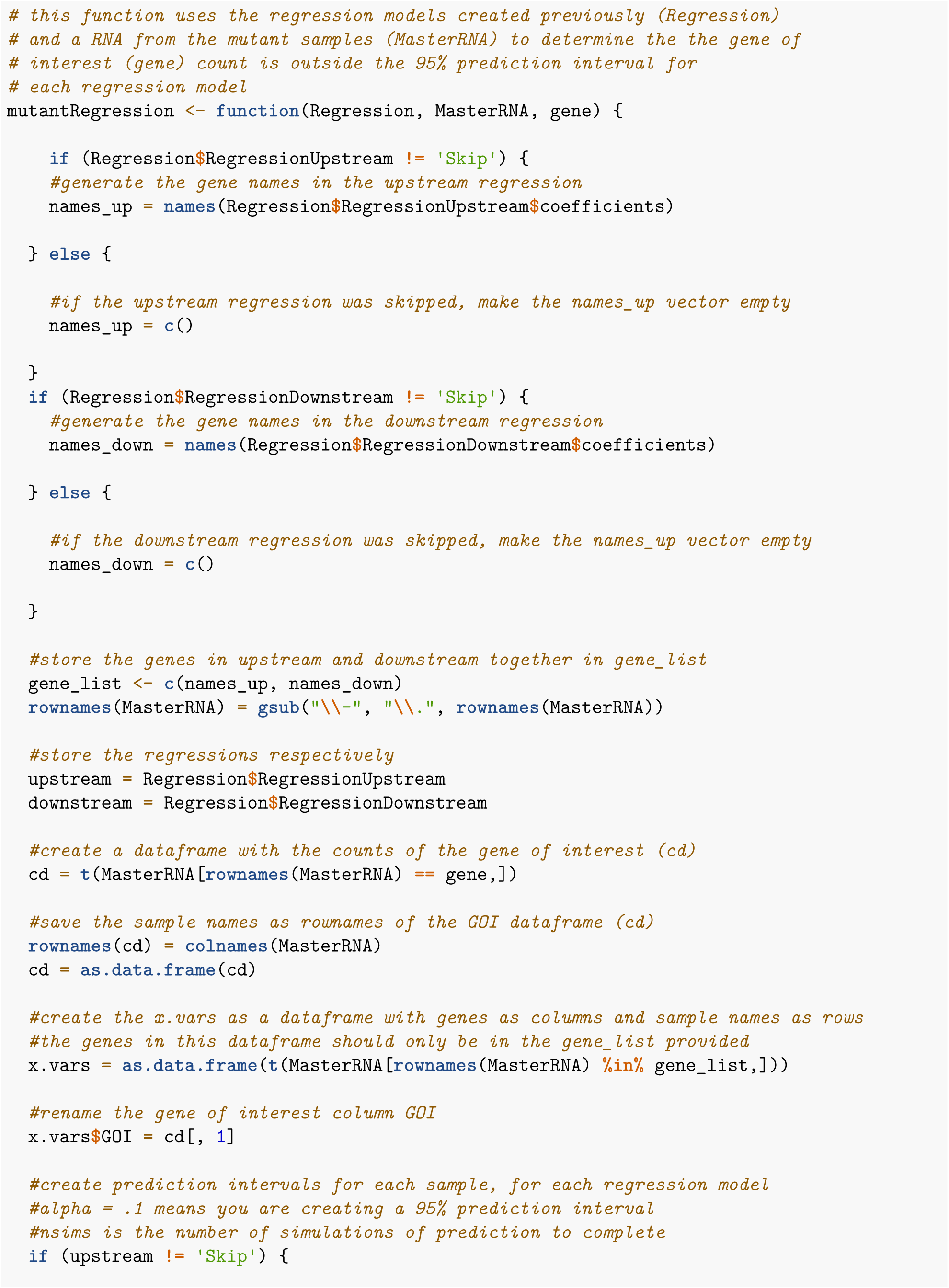

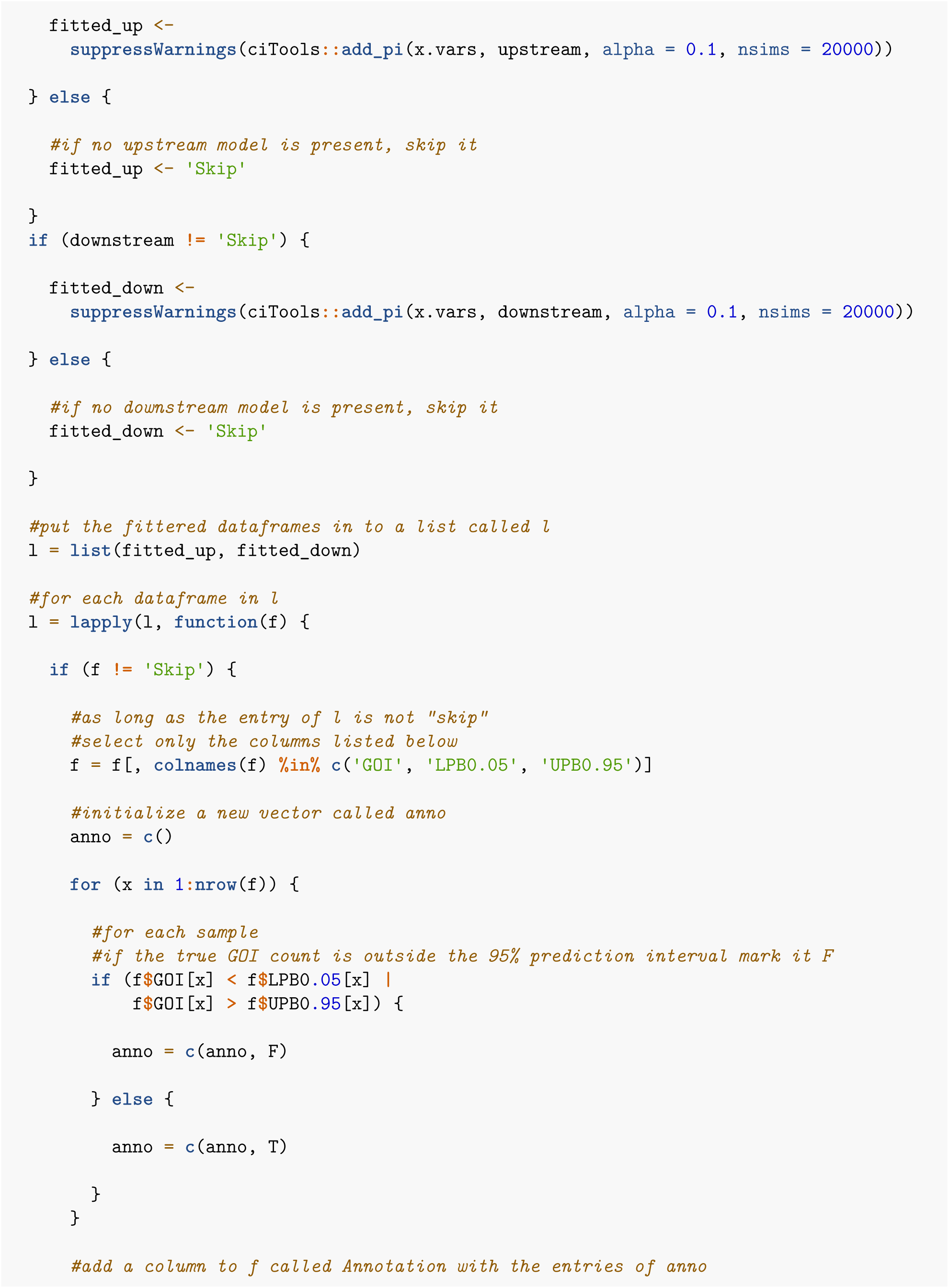

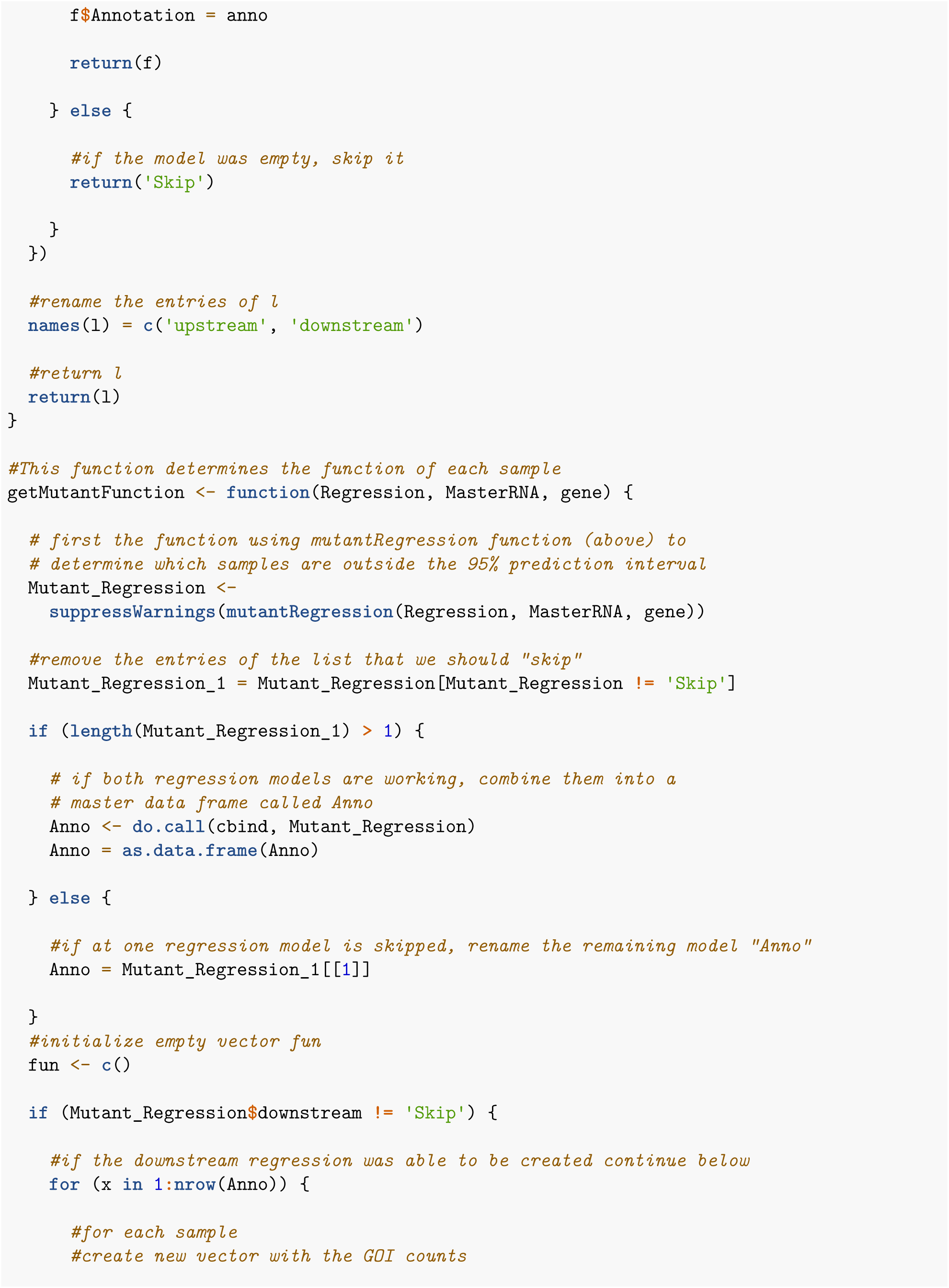

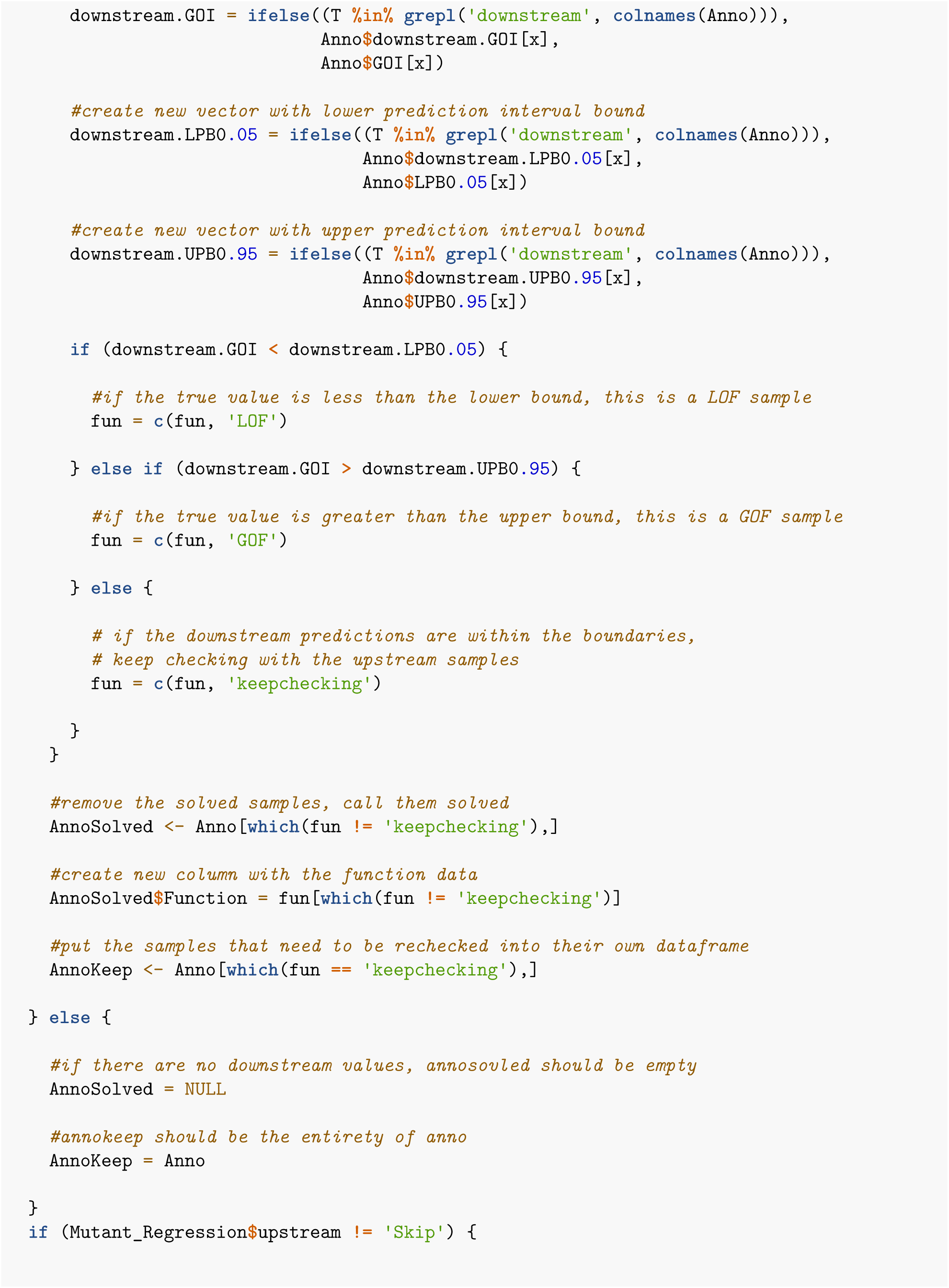

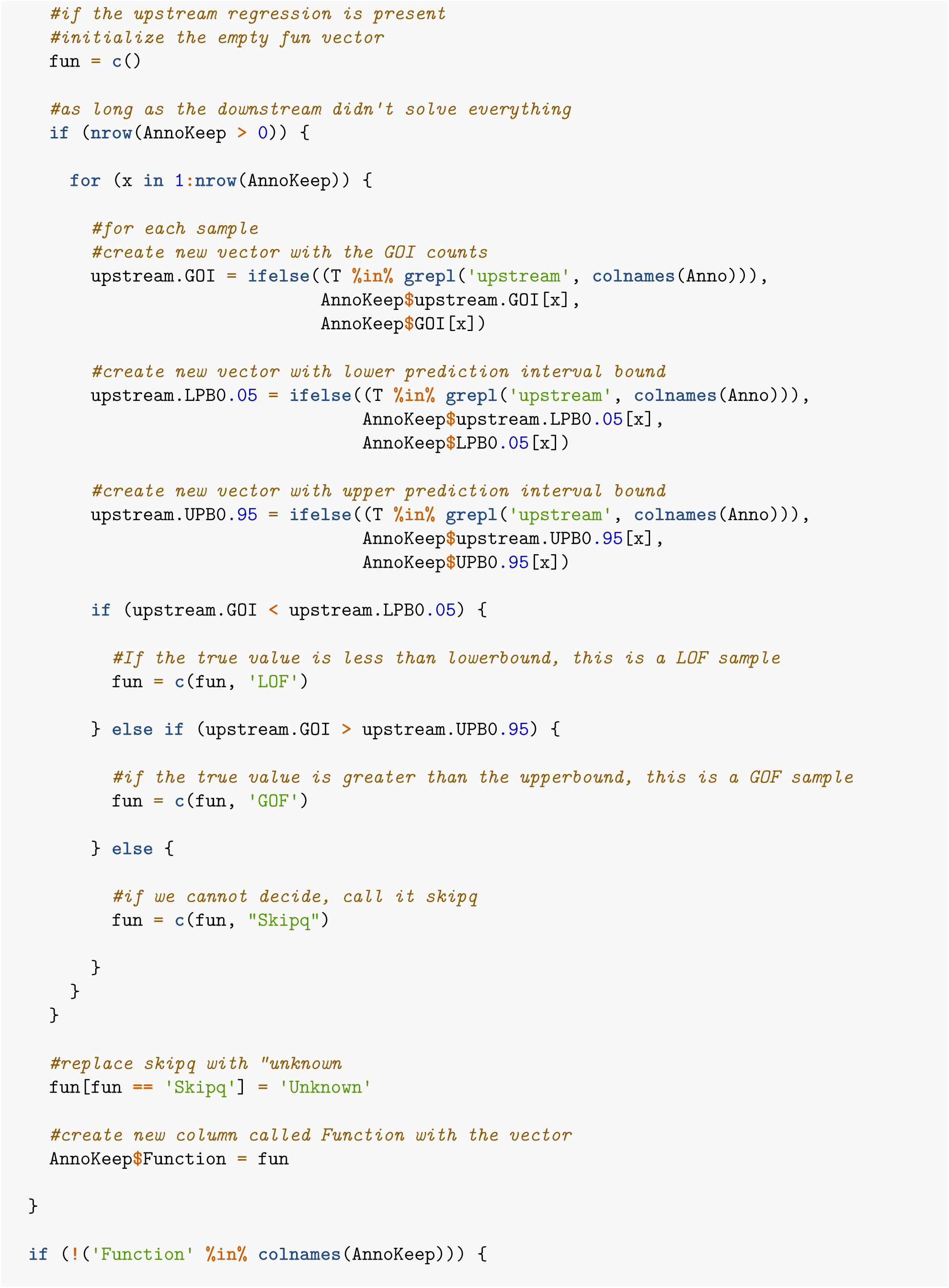

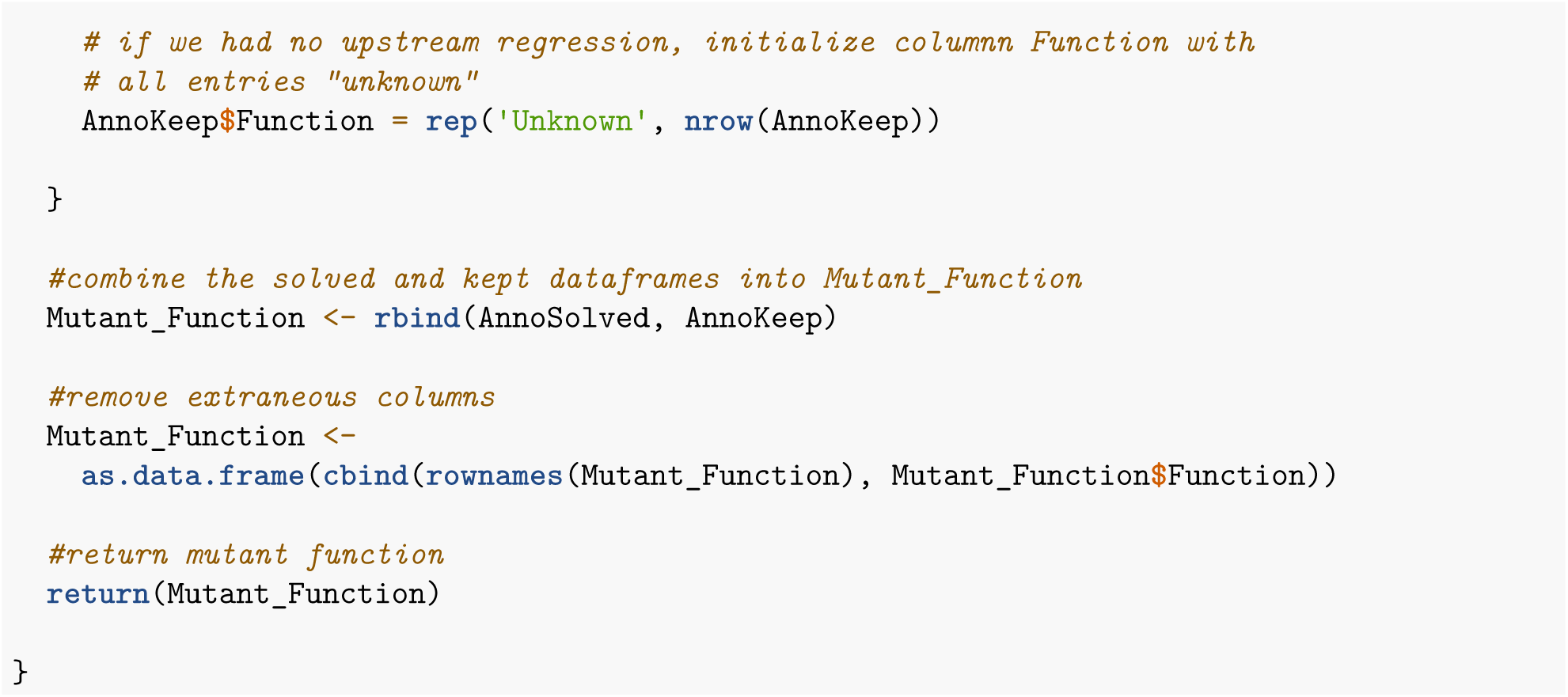

### 4.4: Predict the immune phenotype of each sample

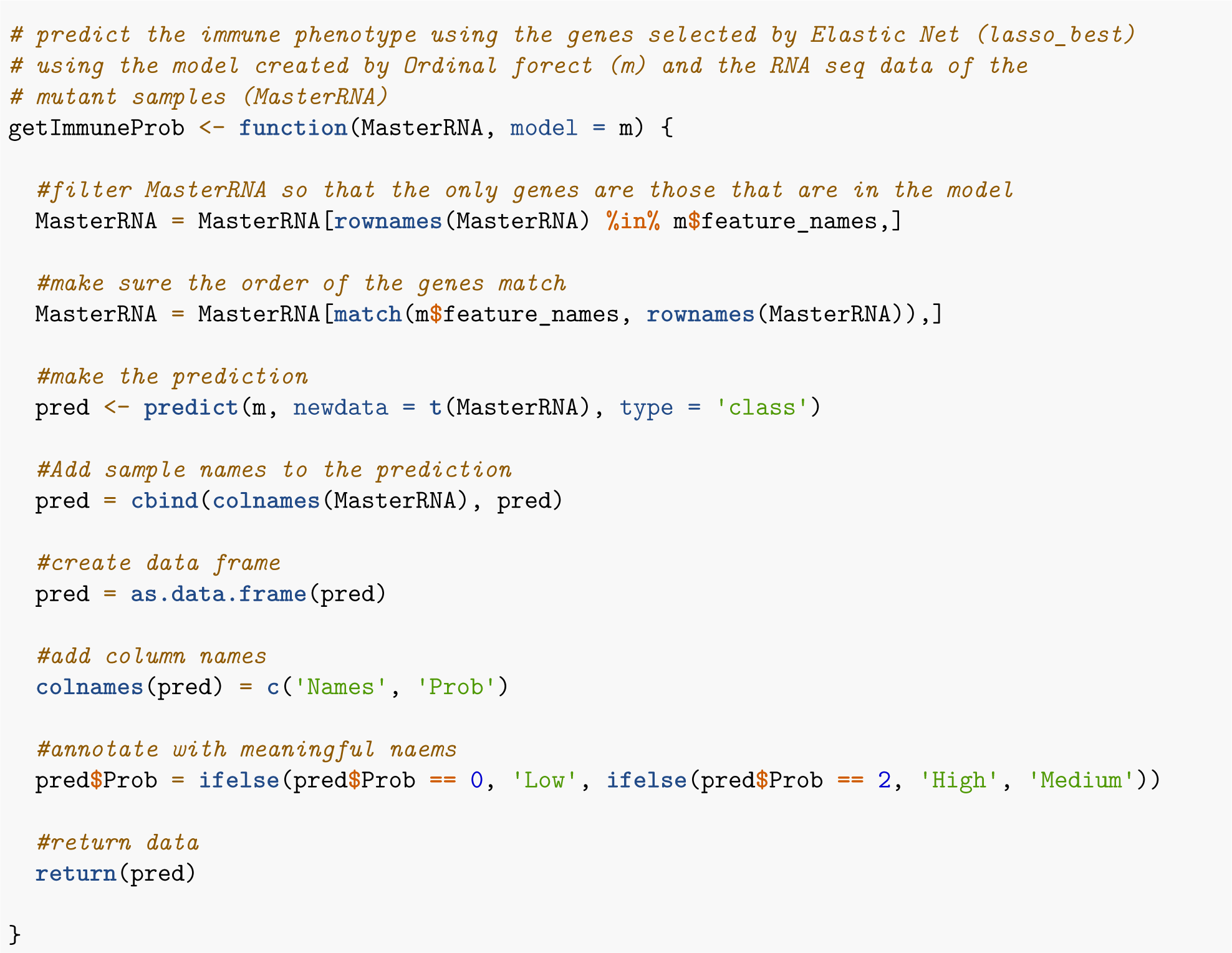

### 4.5: Evaluate mutants for users

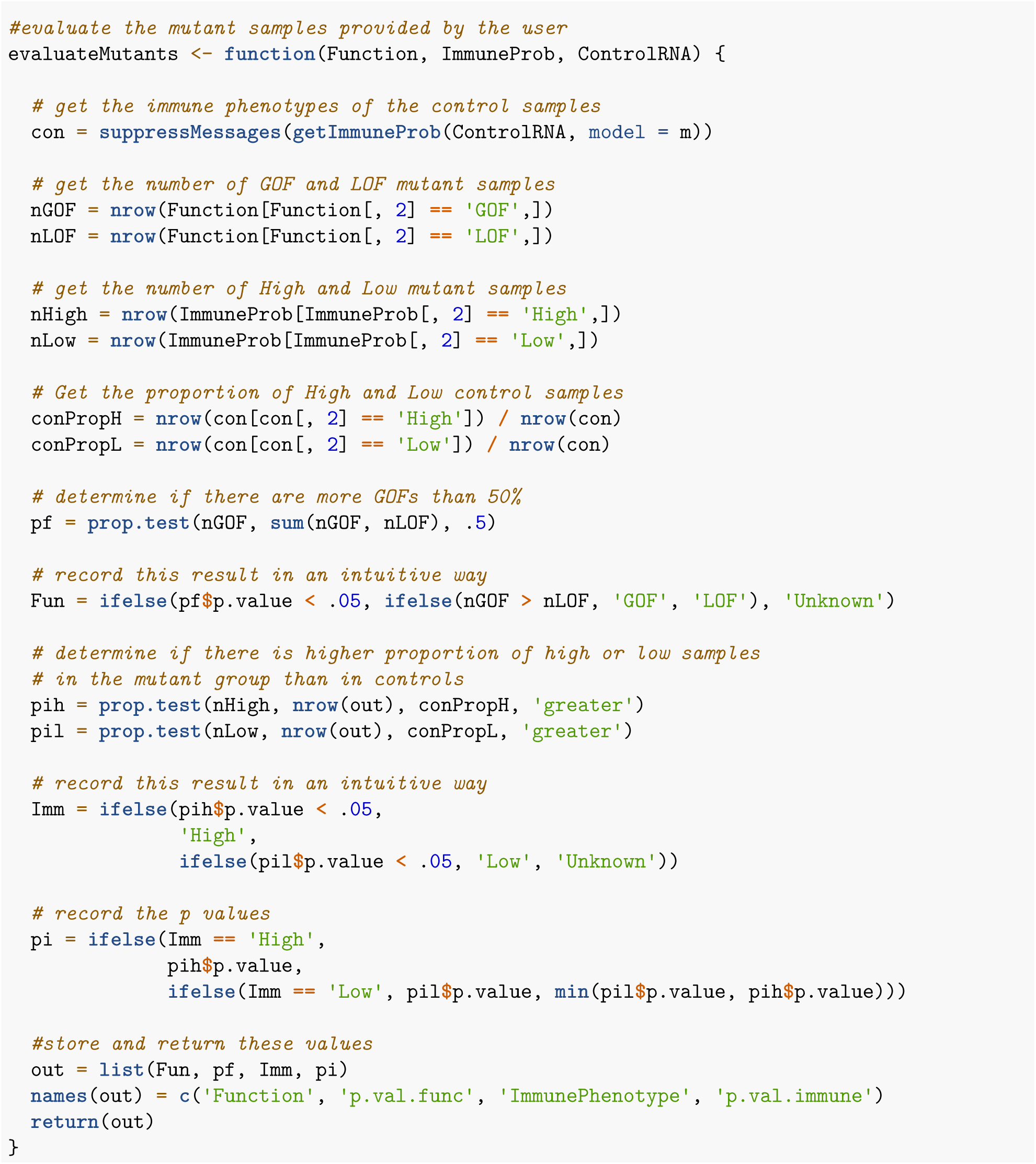

### 4.6. Run RIGATonI all together

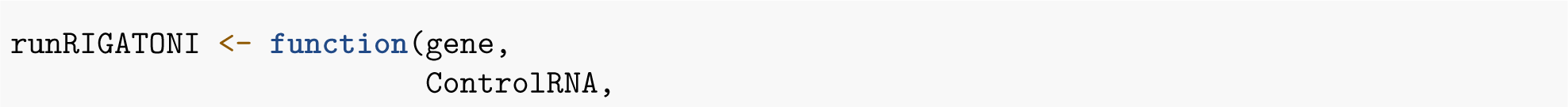

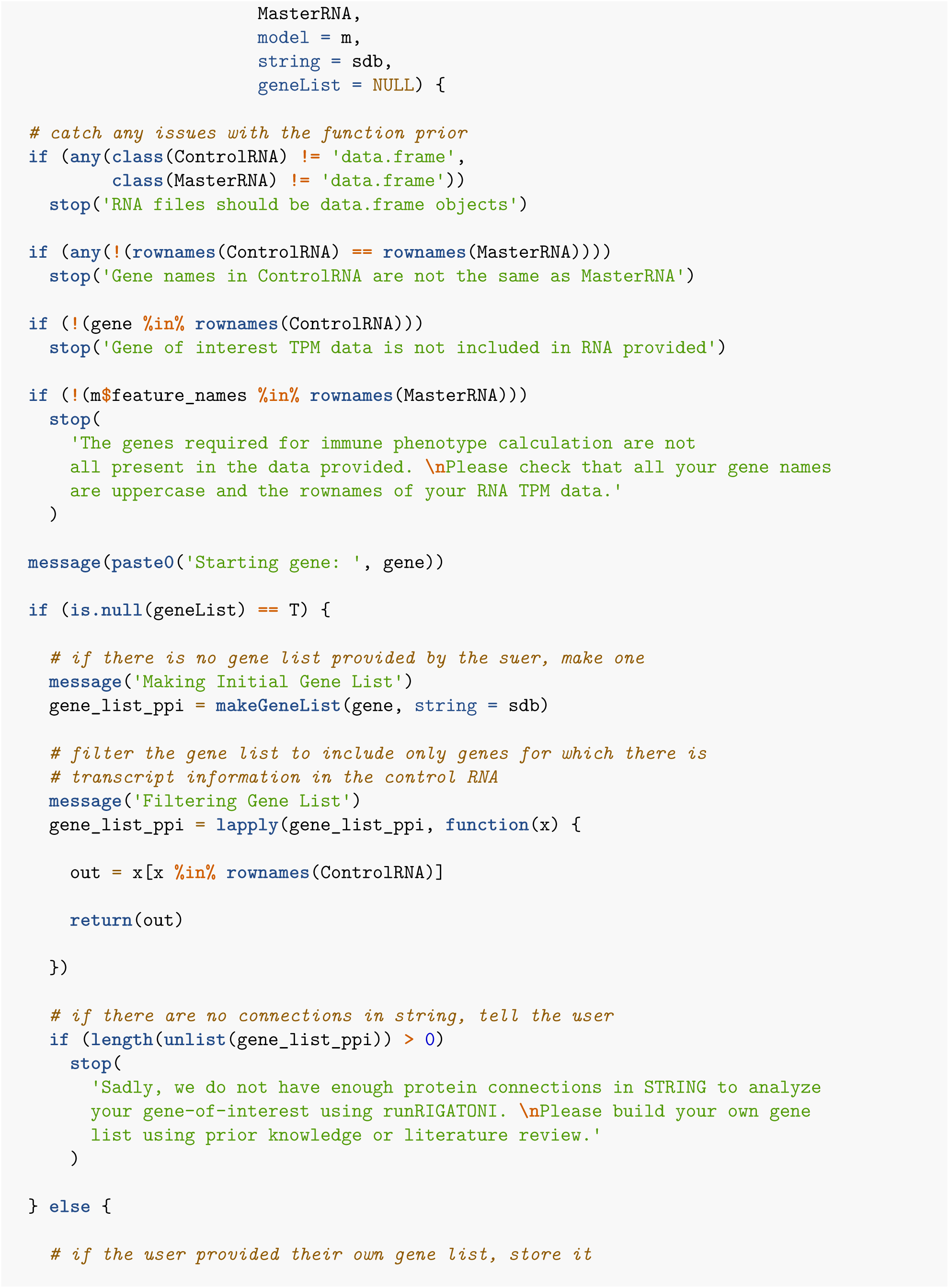

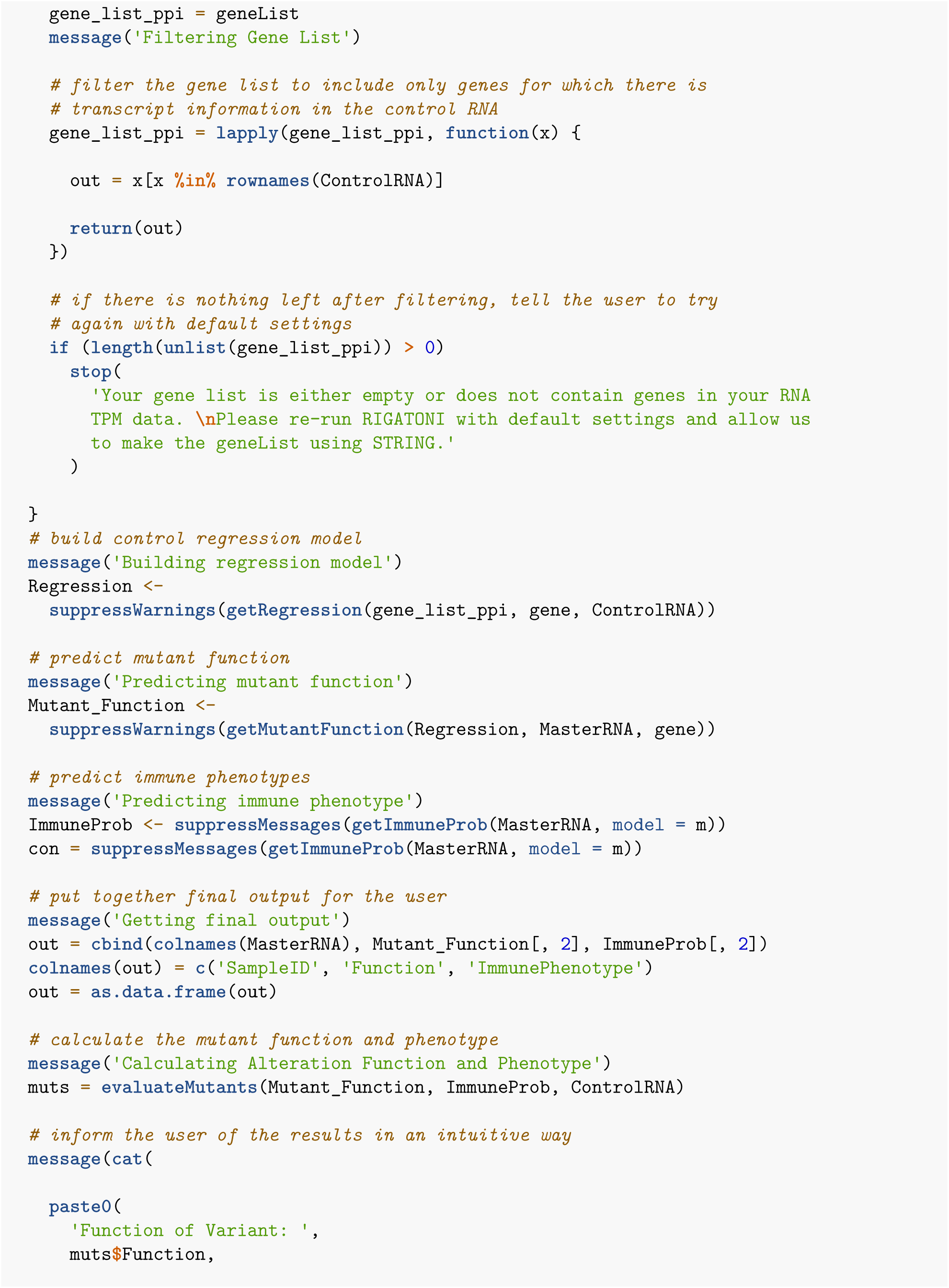

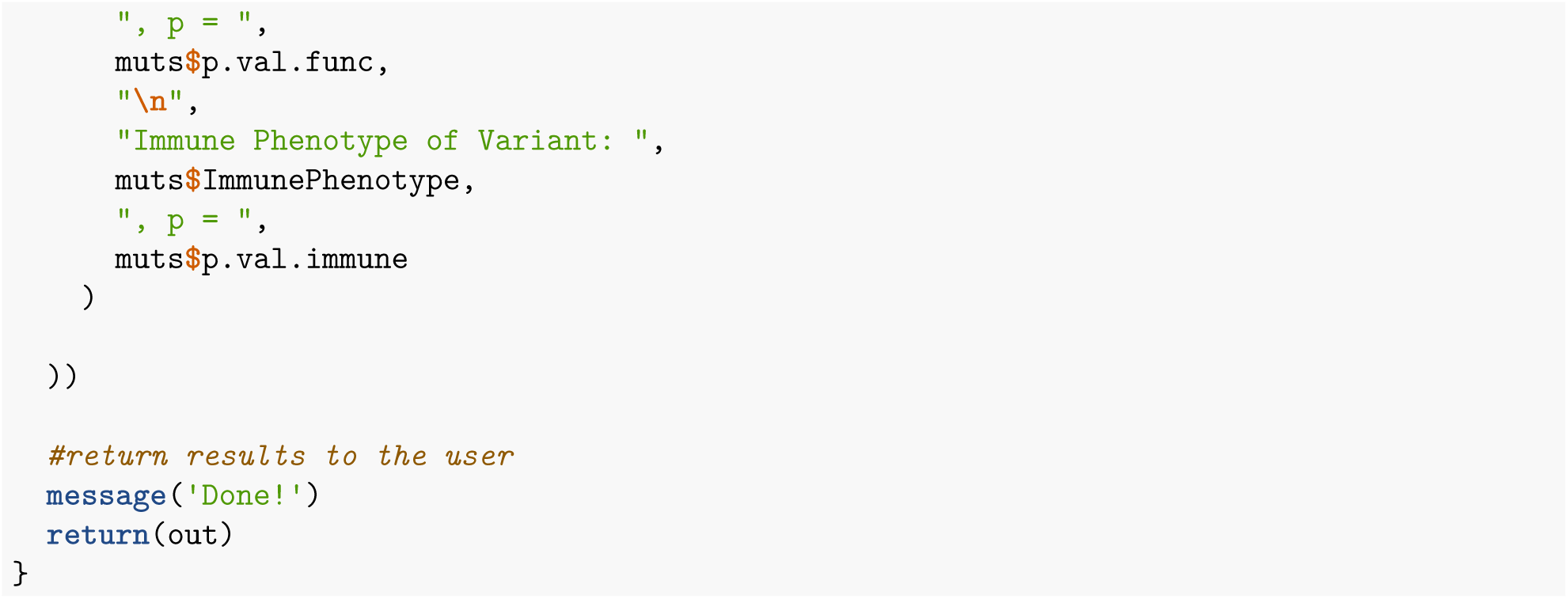

## Part 5: Performing RIGATonI validations (Fig 2 and 3)

### 5.1: Validation of Immunity Module with IHC and Flow (Fig 2)

All data shown here is from the paper “https://www.nature.com/articles/s41598-022-12610-w#Abs1” First we downloaded the raw fastqs from dbGaP (the accession number is provided under “Data Availability” in the paper). Next we ran Salmon to extract RNA TPM from the fastqs; code shown below.

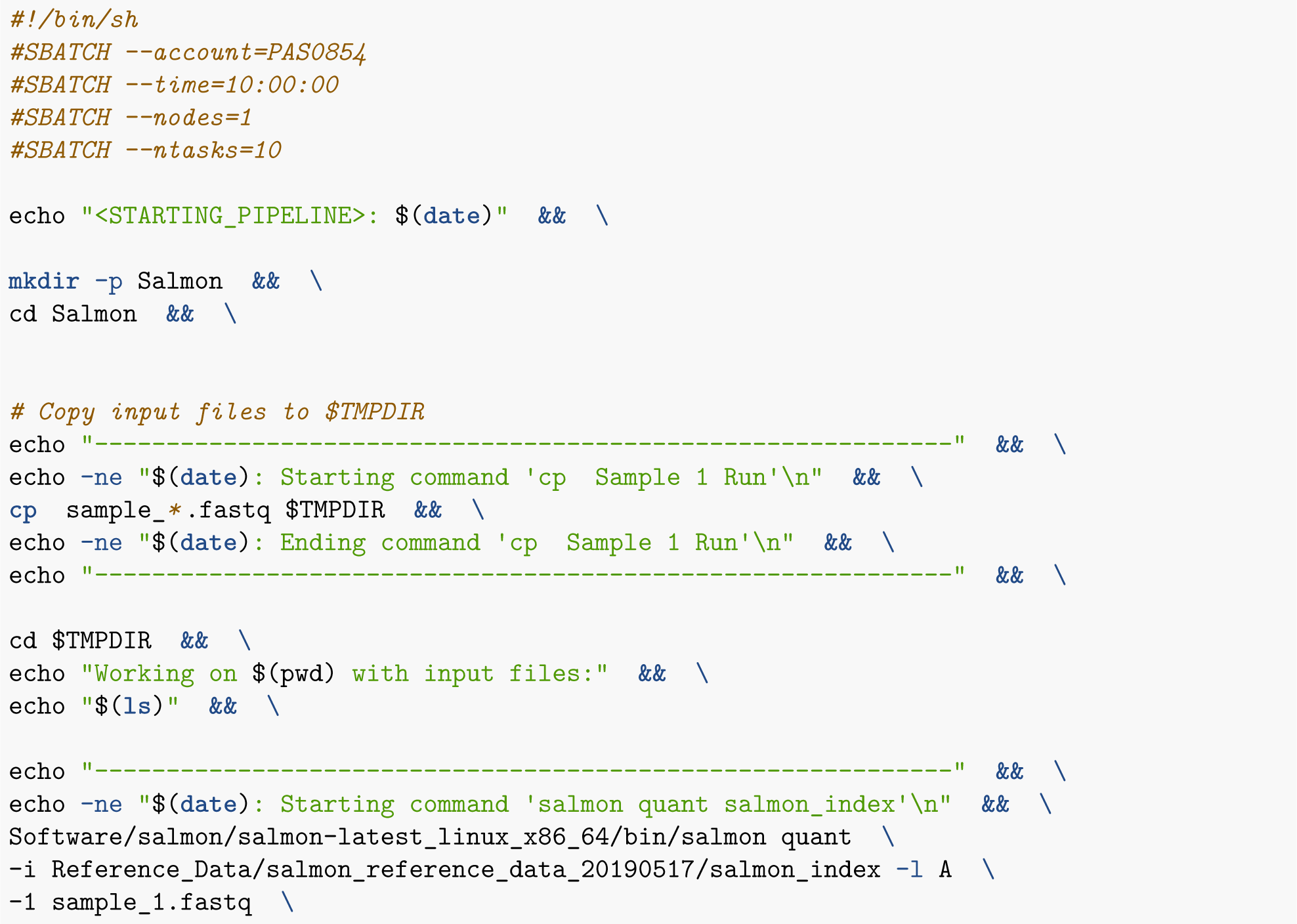

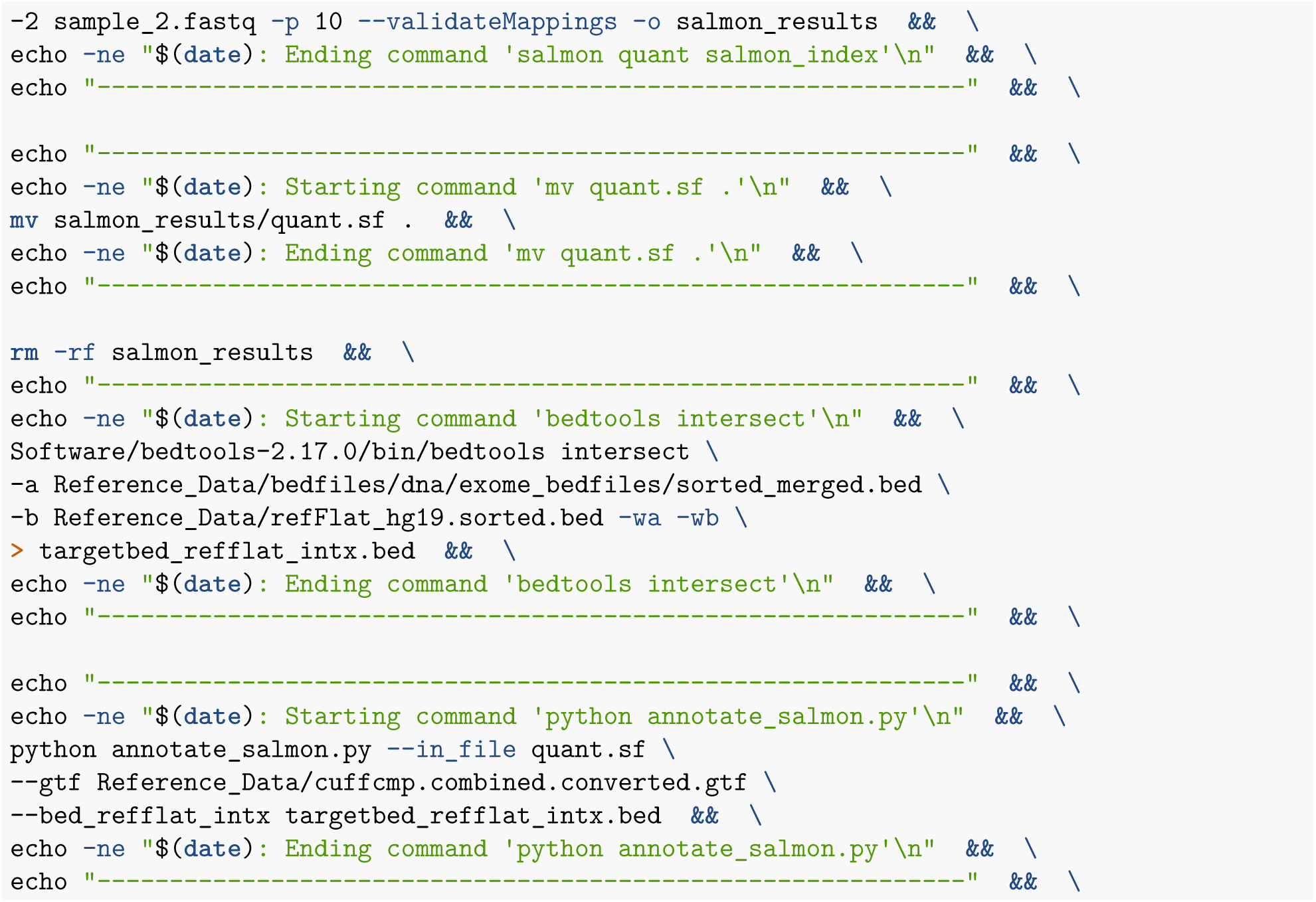

Once salmon has run, we use a python script (annotate_salmon.py) to transform RefSeq ID to gene names shown below

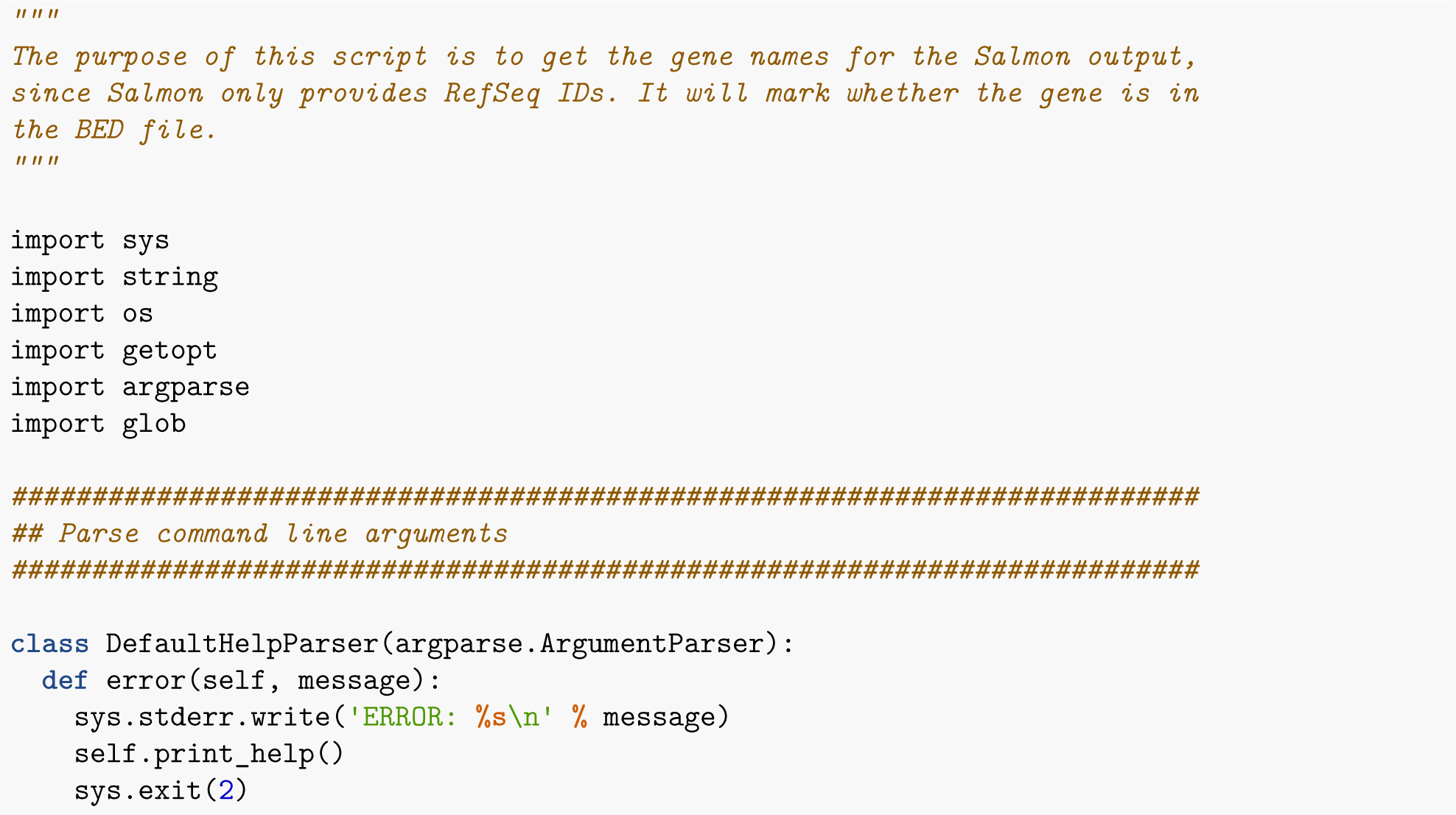

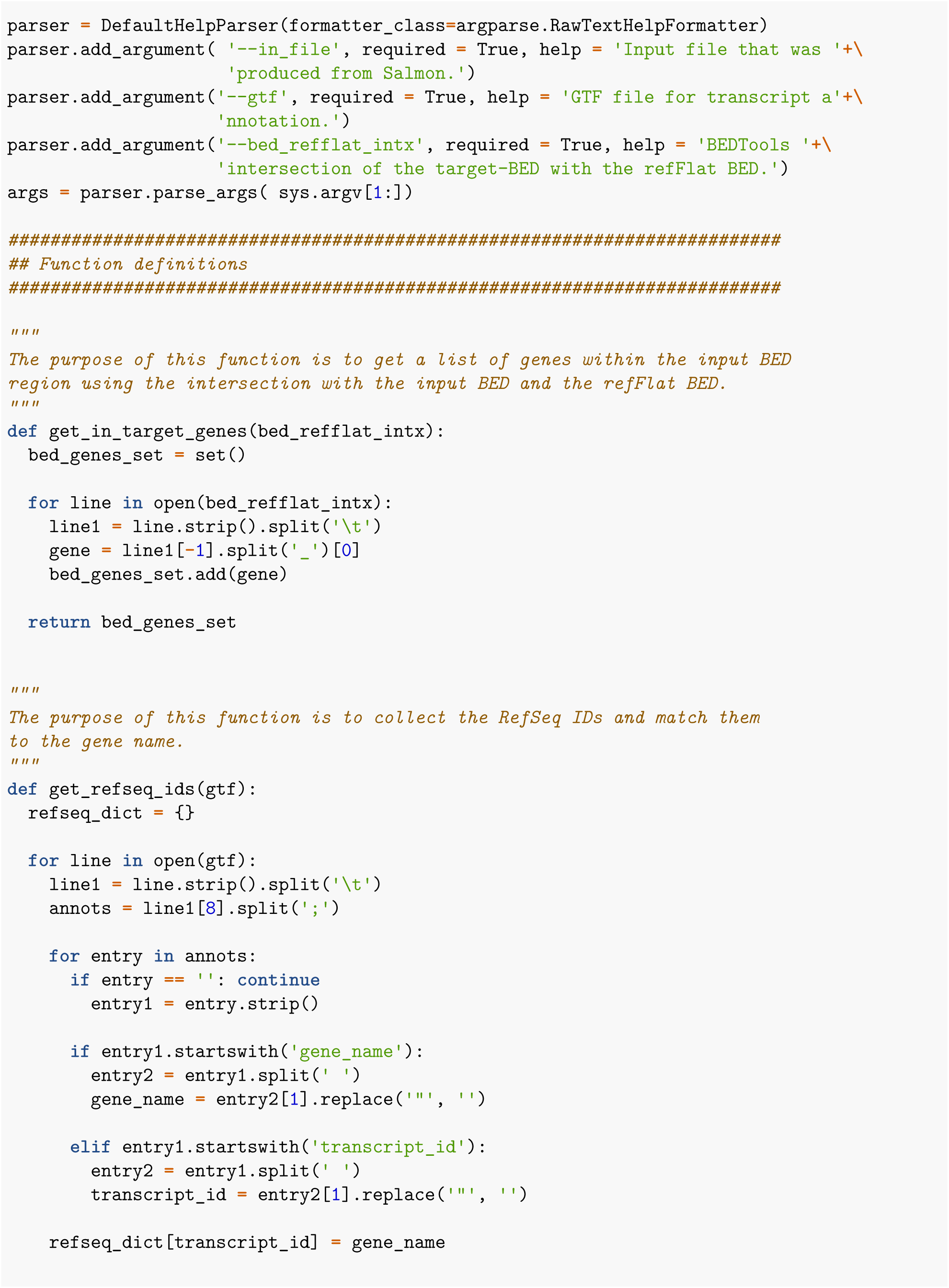

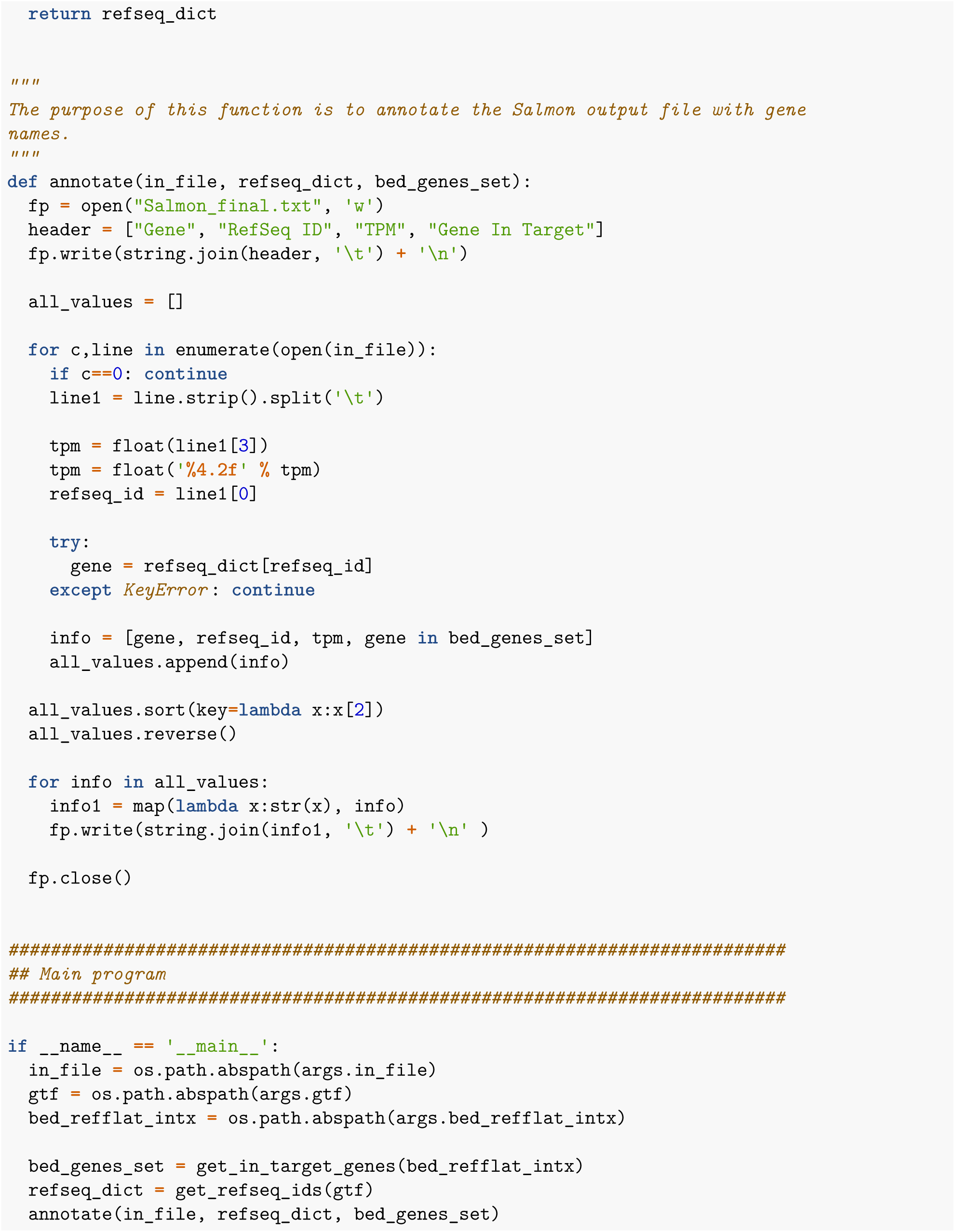

After downloading and performing salmon, I then combined and analyzed the IHC, flow, and RNAseq results.

Processed data was downlaoded from the supplemental data at this link https://static-content.springer.com/esm/art%3A10.1038%2Fs41598-022-12610-w/MediaObjects/41598_2022_12610_MOESM2_ESM.xlsx.

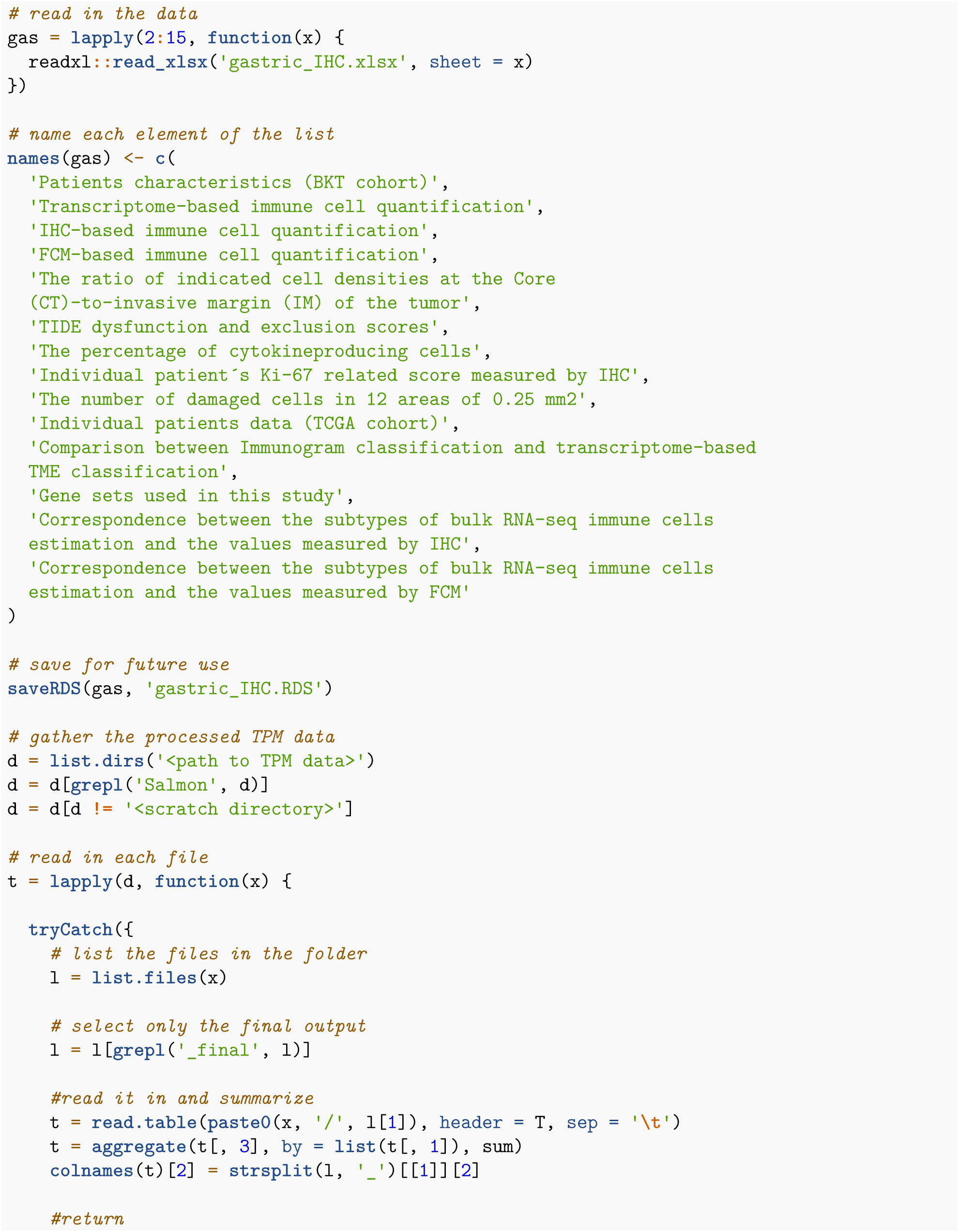

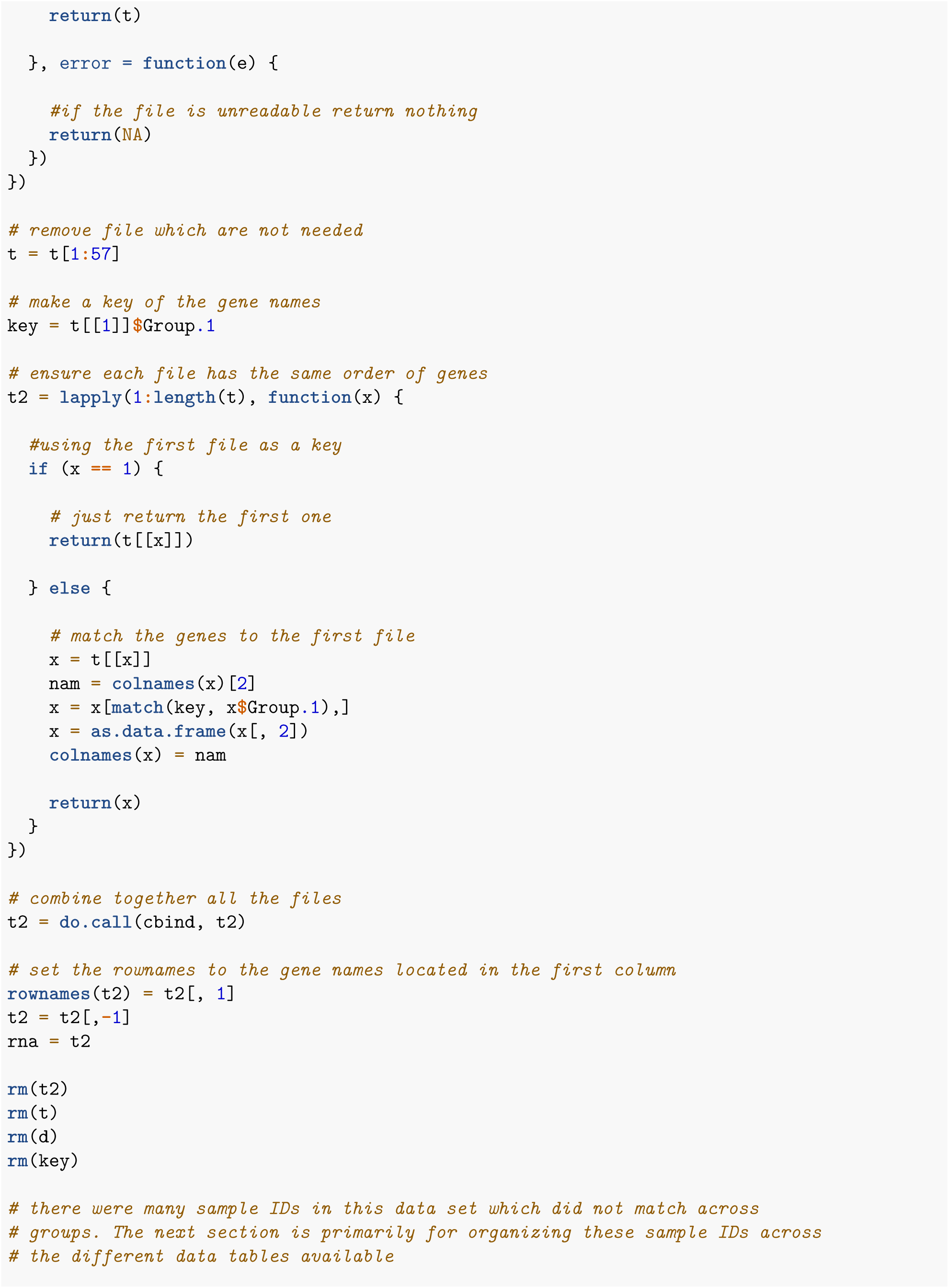

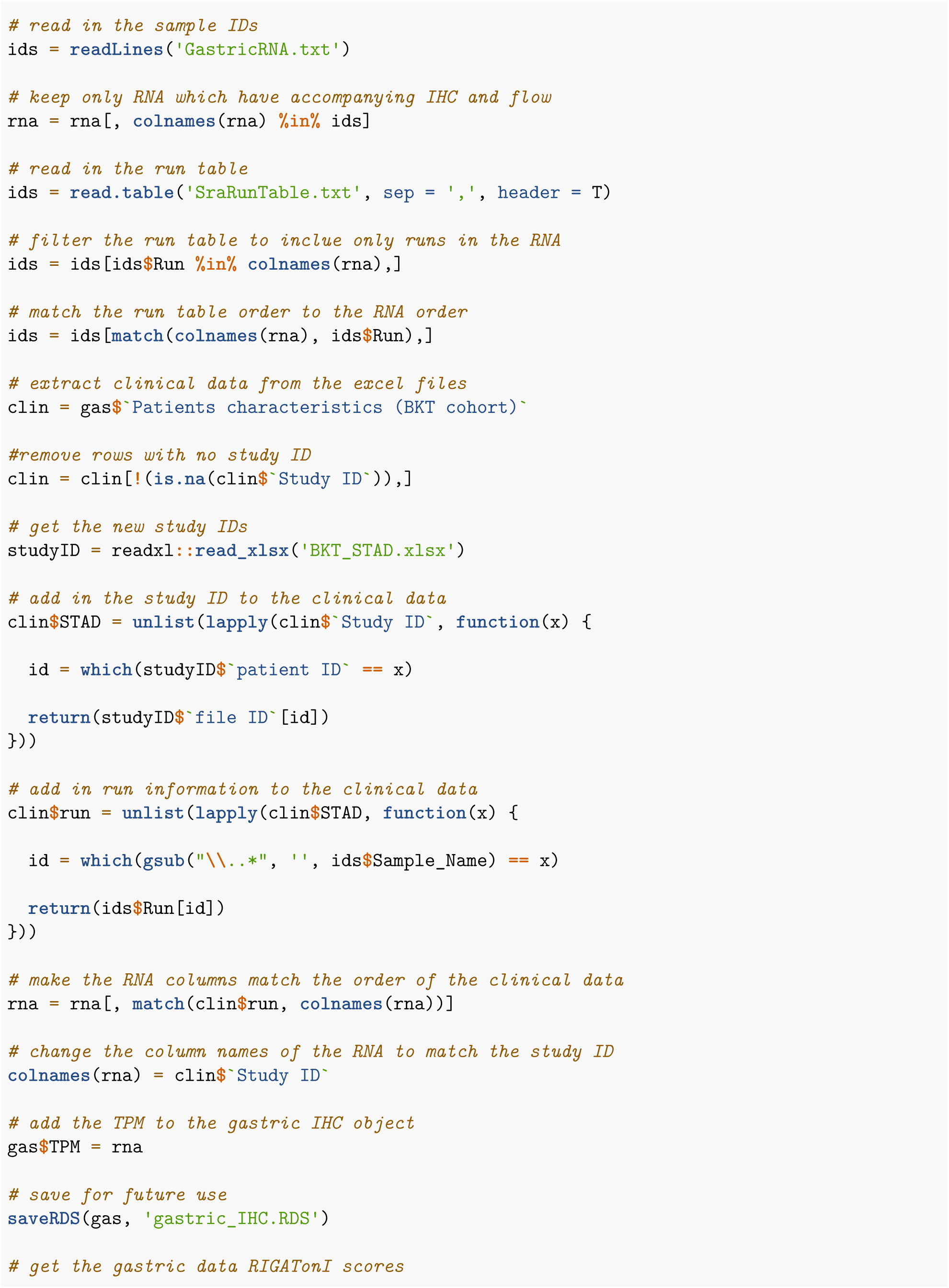

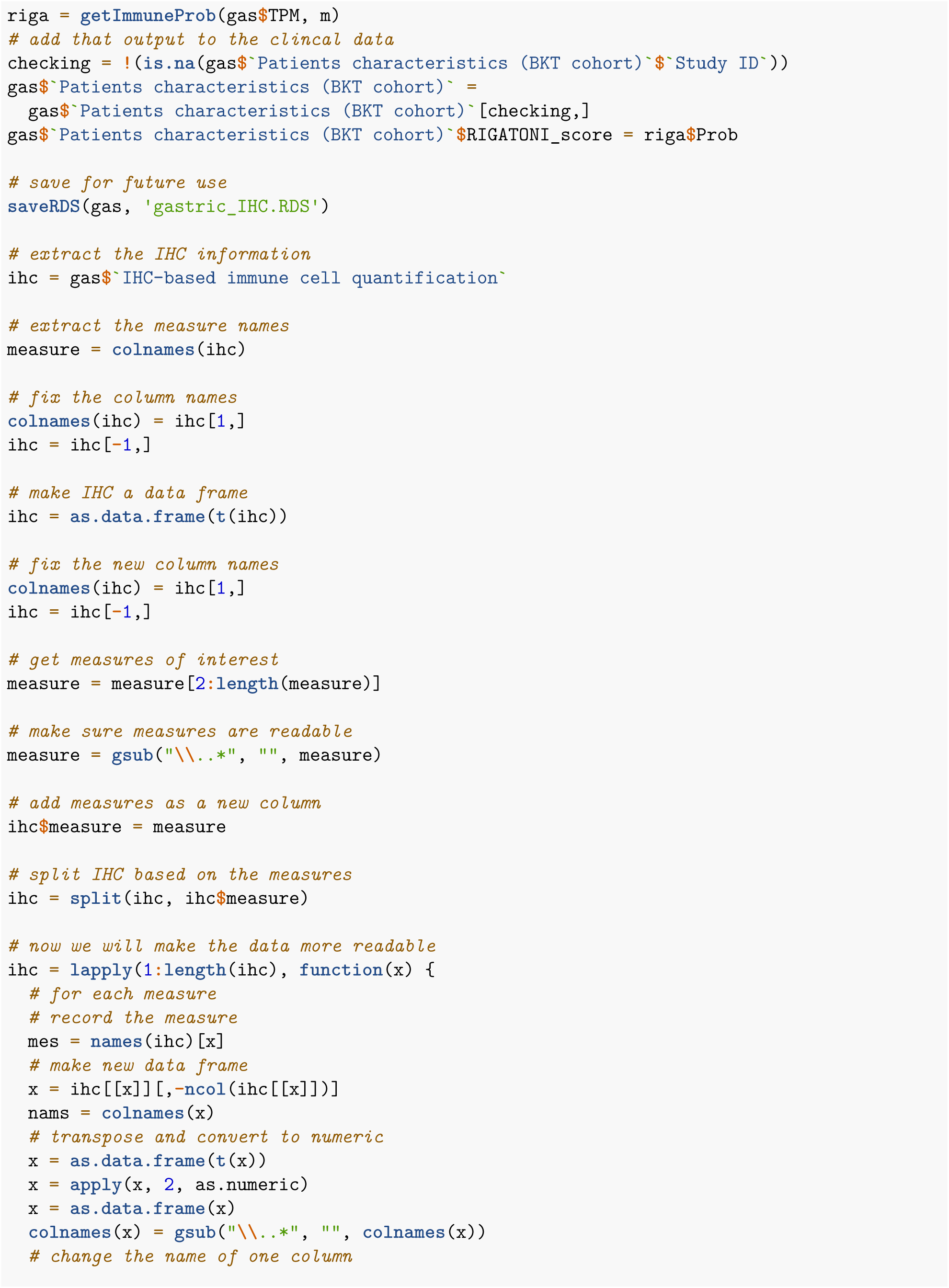

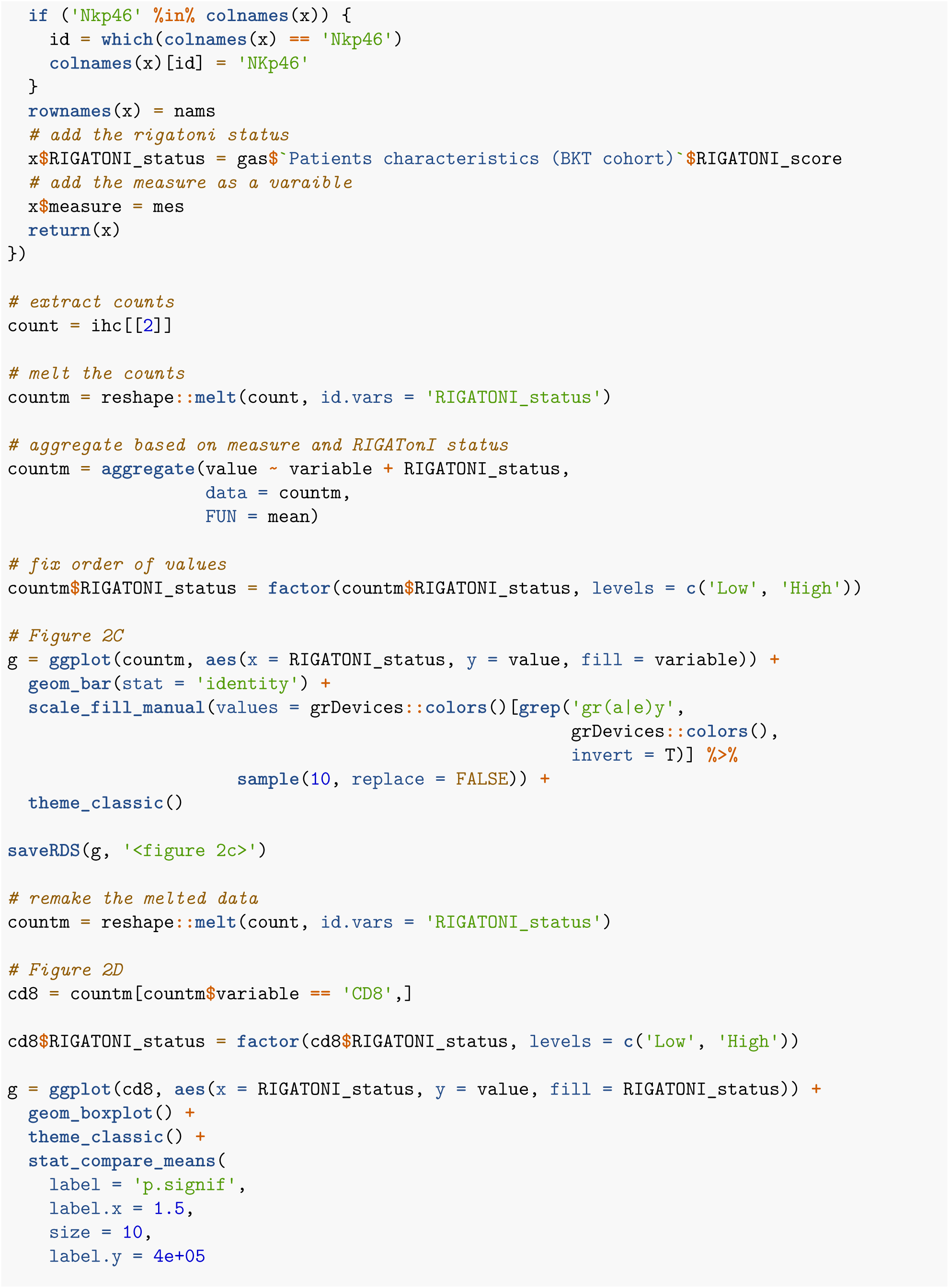

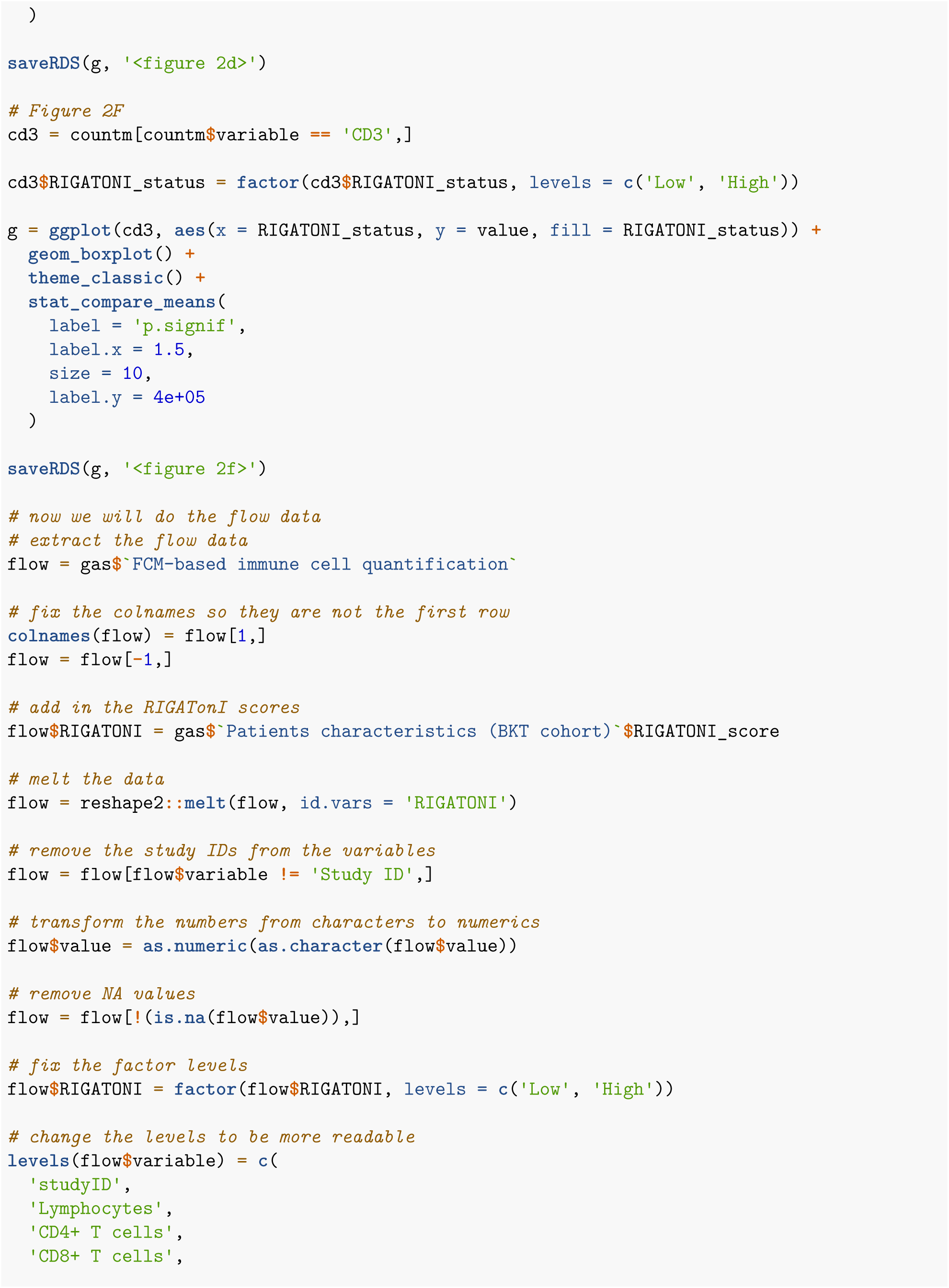

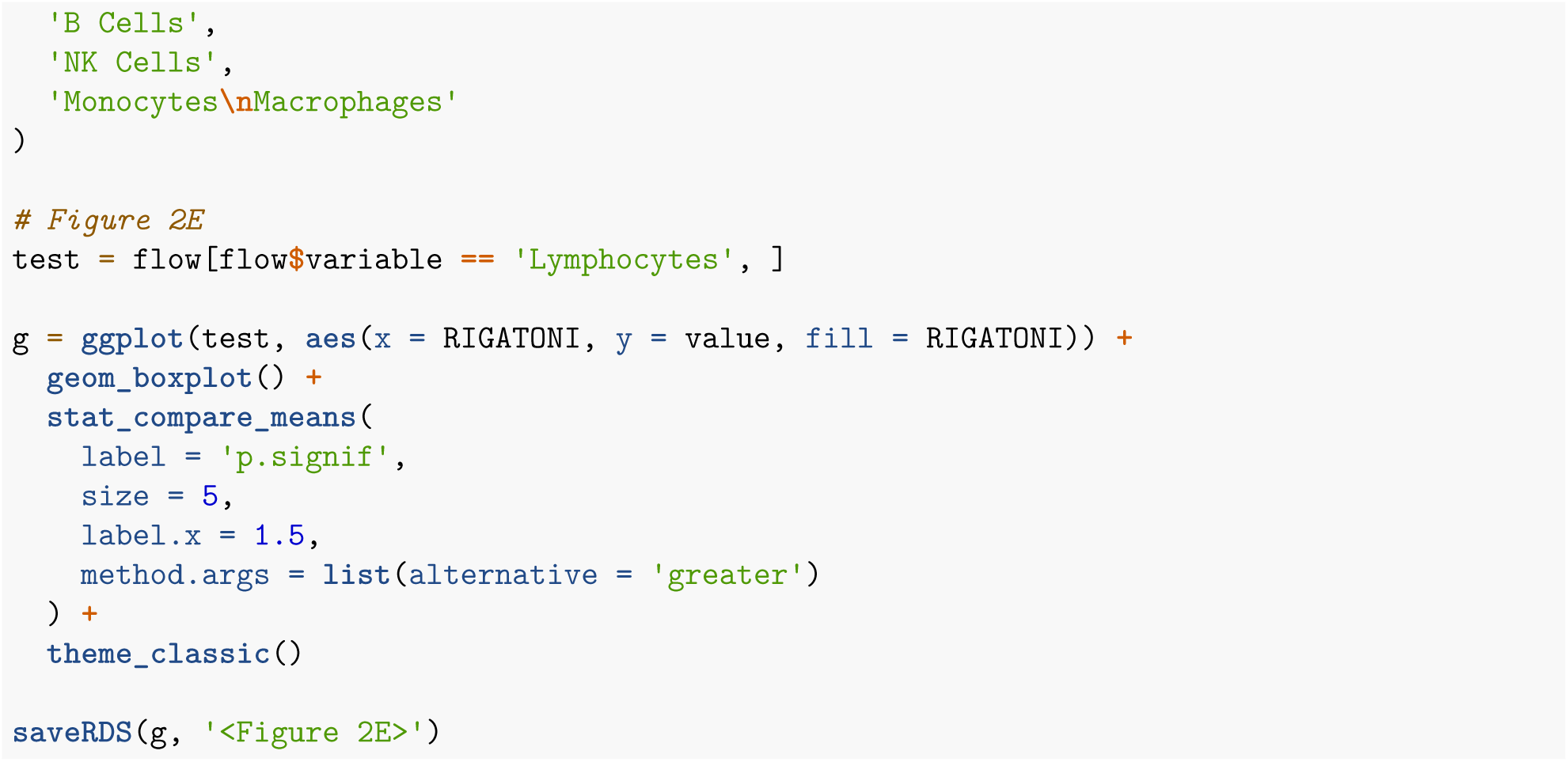

### 5.2: Validation of Immunity Module with Saltz Et Al output (Fig 2)

All data shown here is from the paper “https://www.sciencedirect.com/science/article/pii/S2211124718304479?via%3Dihub” First we downloaded the output from Saltz et al found at “https://ars.els-cdn.com/content/image/1-s2.0-S2211124718304479-mmc2.xlsx” Then the following code was run.

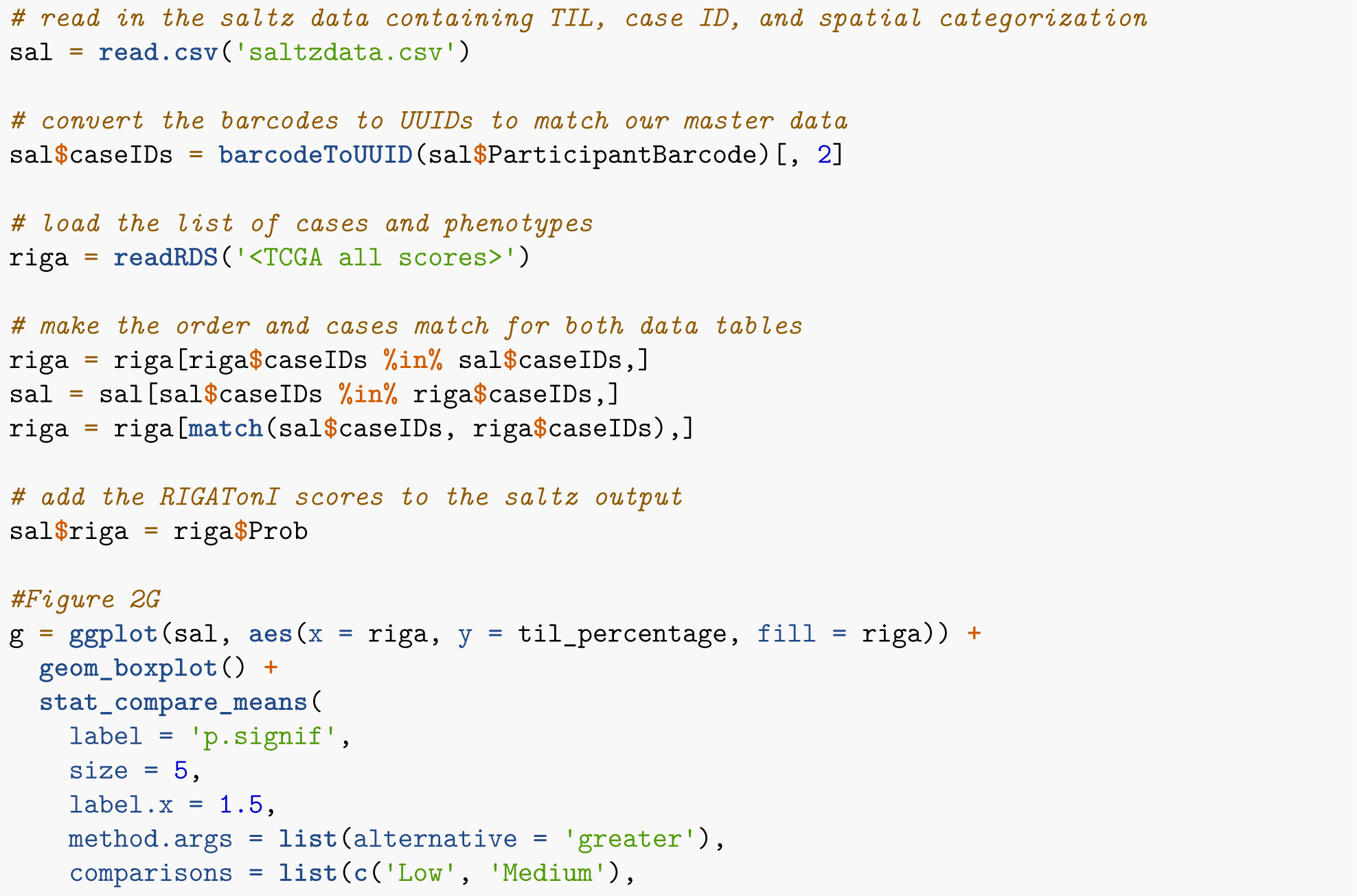

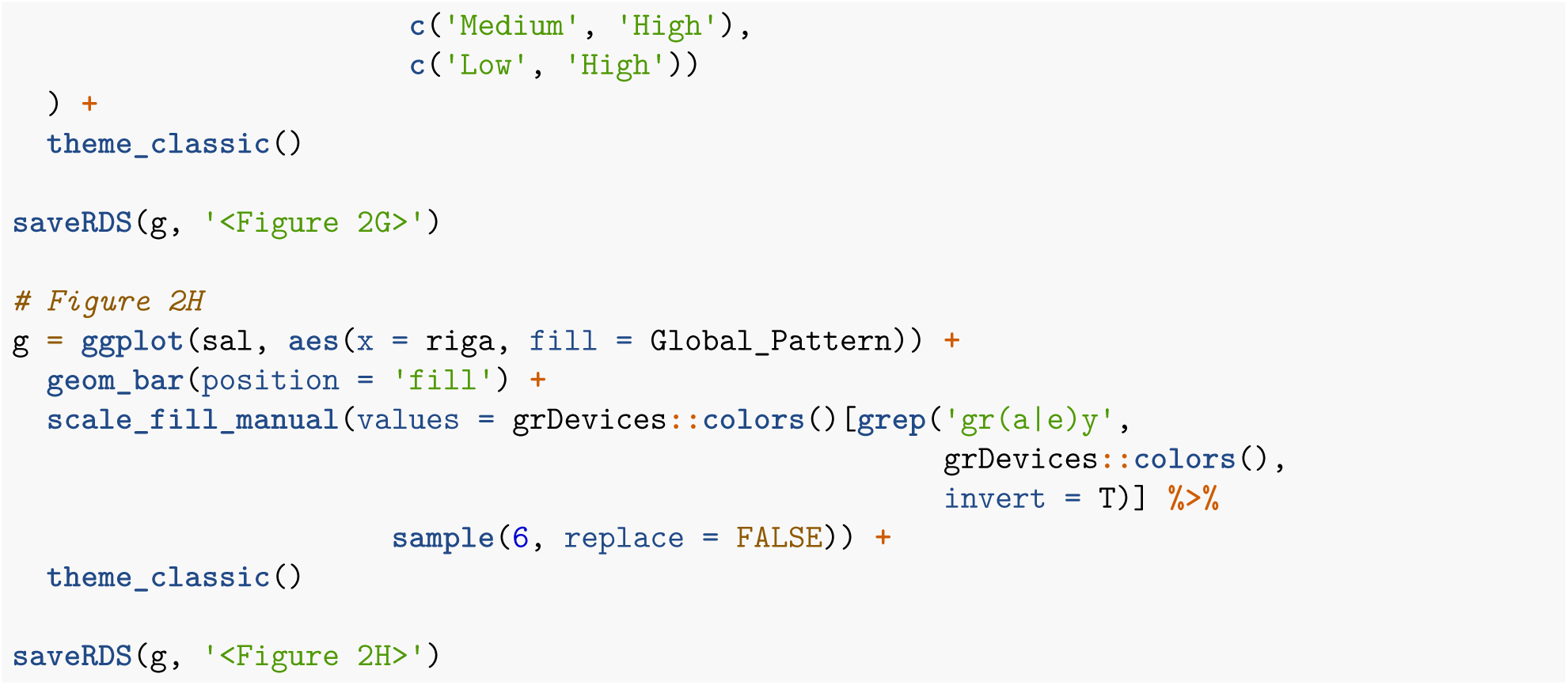

### 5.3: Validation of Function Module with OncoKB (Fig 3)

First, the Function Module was run using the functions described in section 4 on OncoKB genes. Results were manually annotated using the OncoKB database. The complete results can be found in Supplementary table 3.

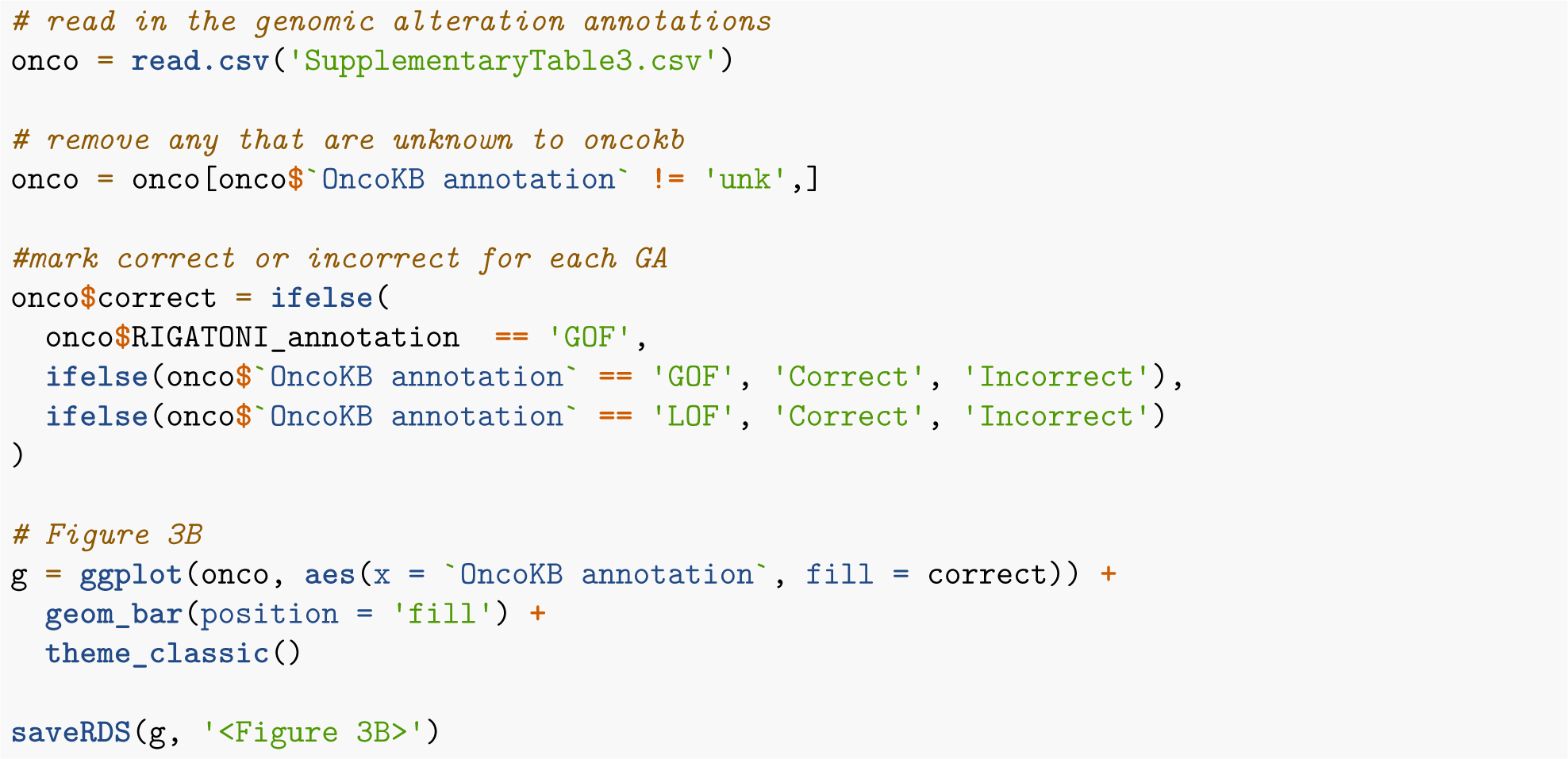

## Part 6: Analyze all of TCGA using RIGATONI

### 6.1: Create functions specific to the analysis of TCGA data

I created a series of functions to make analysis of TCGA data easier. They are shown below.

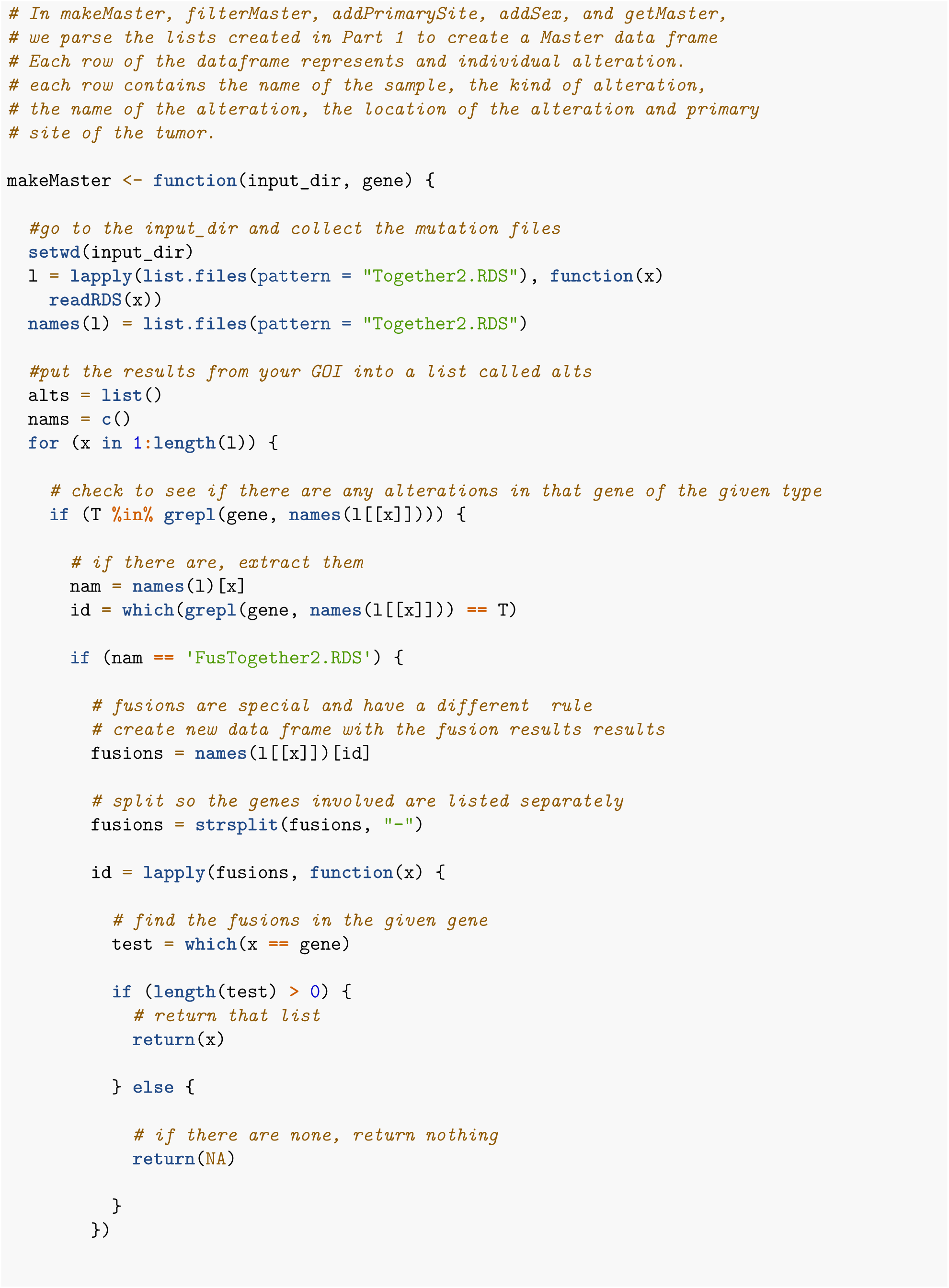

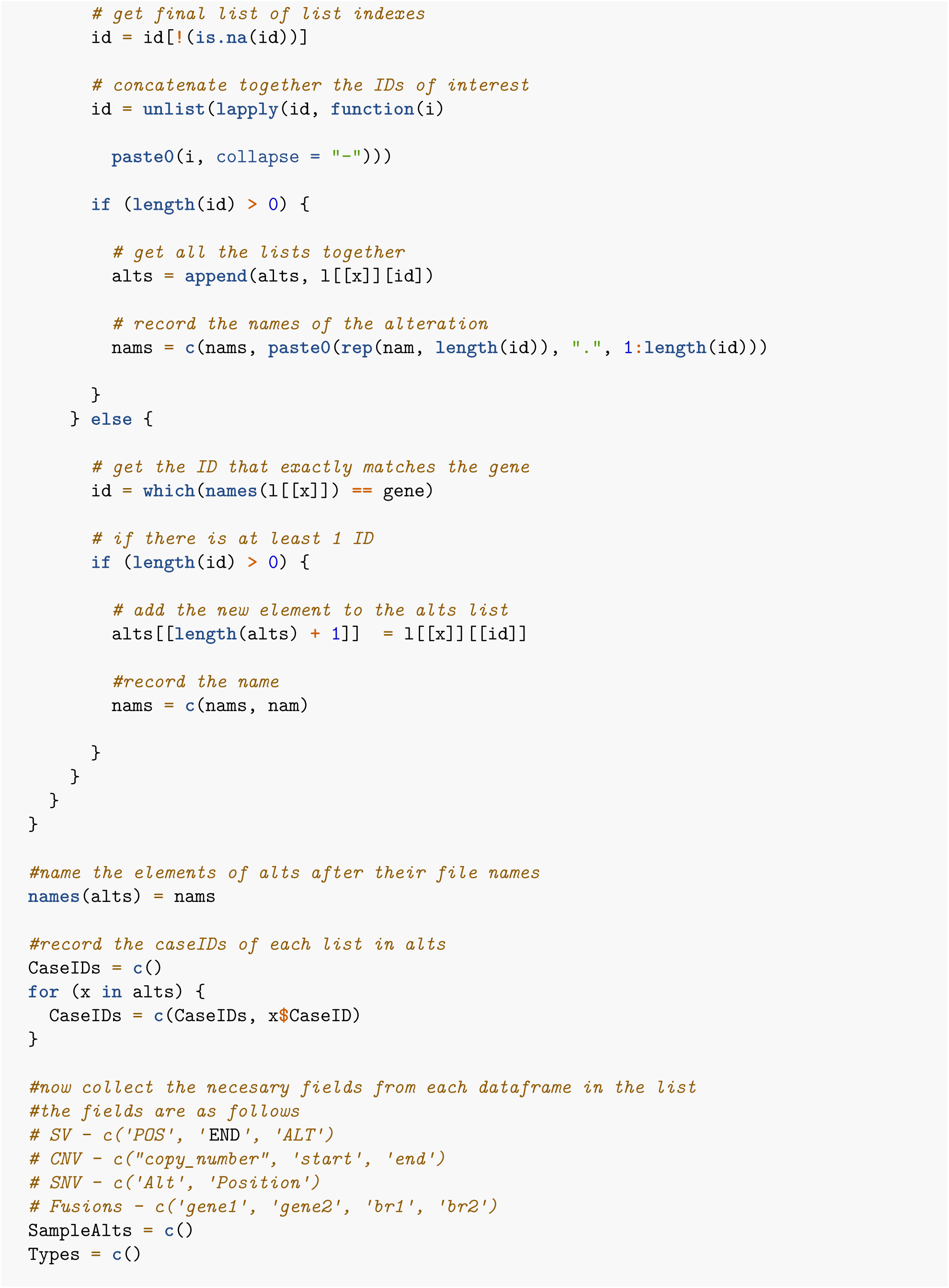

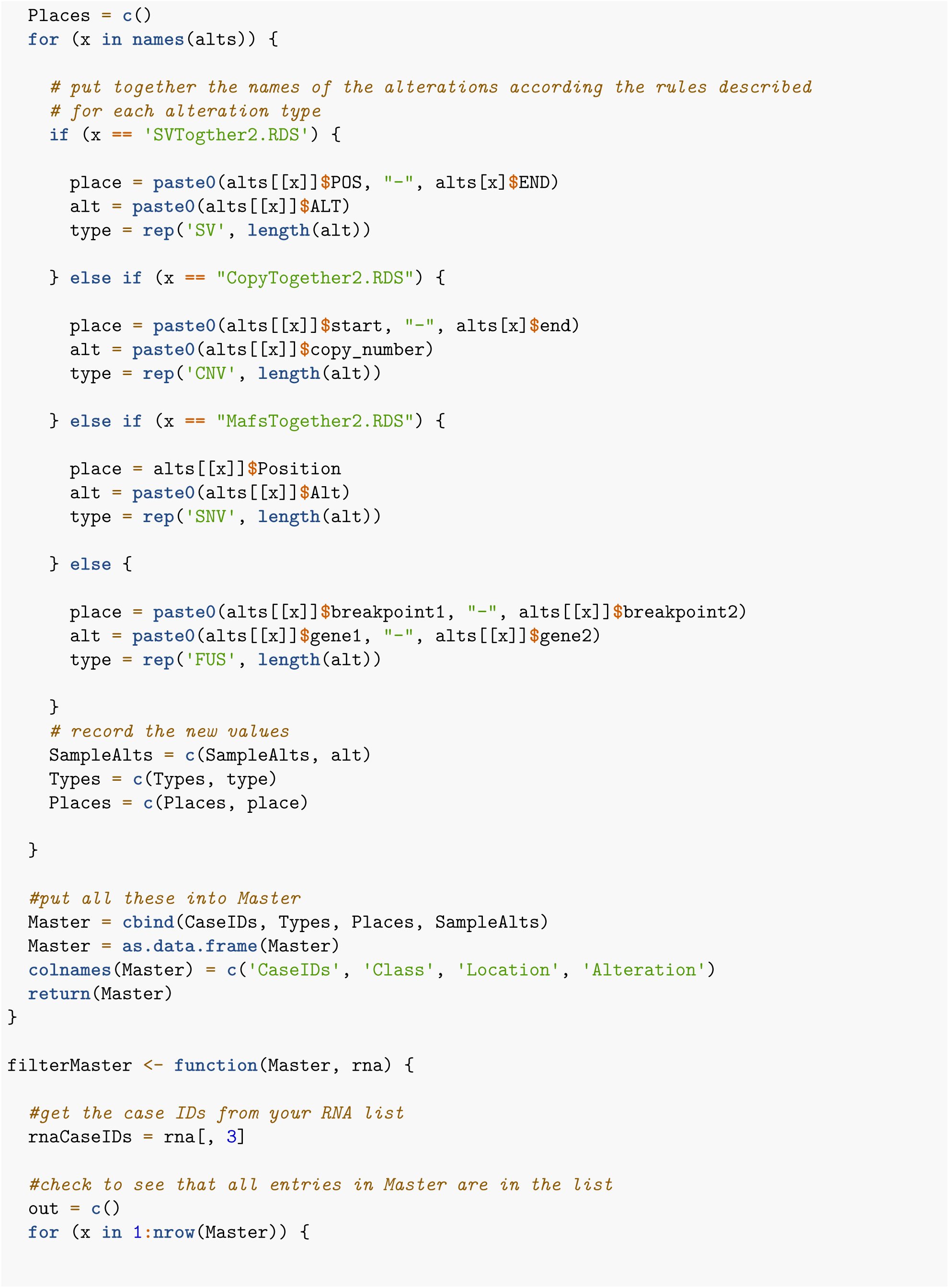

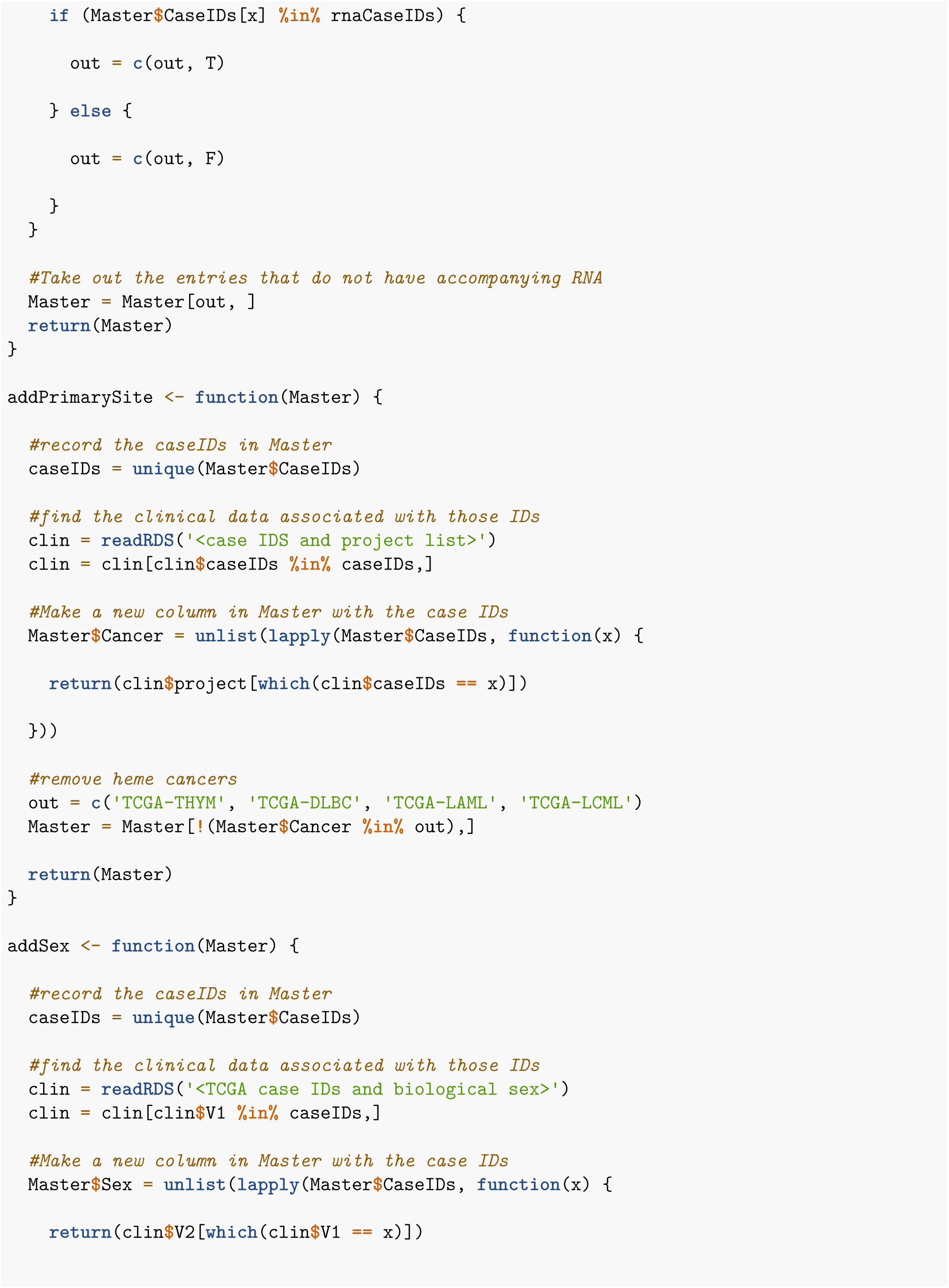

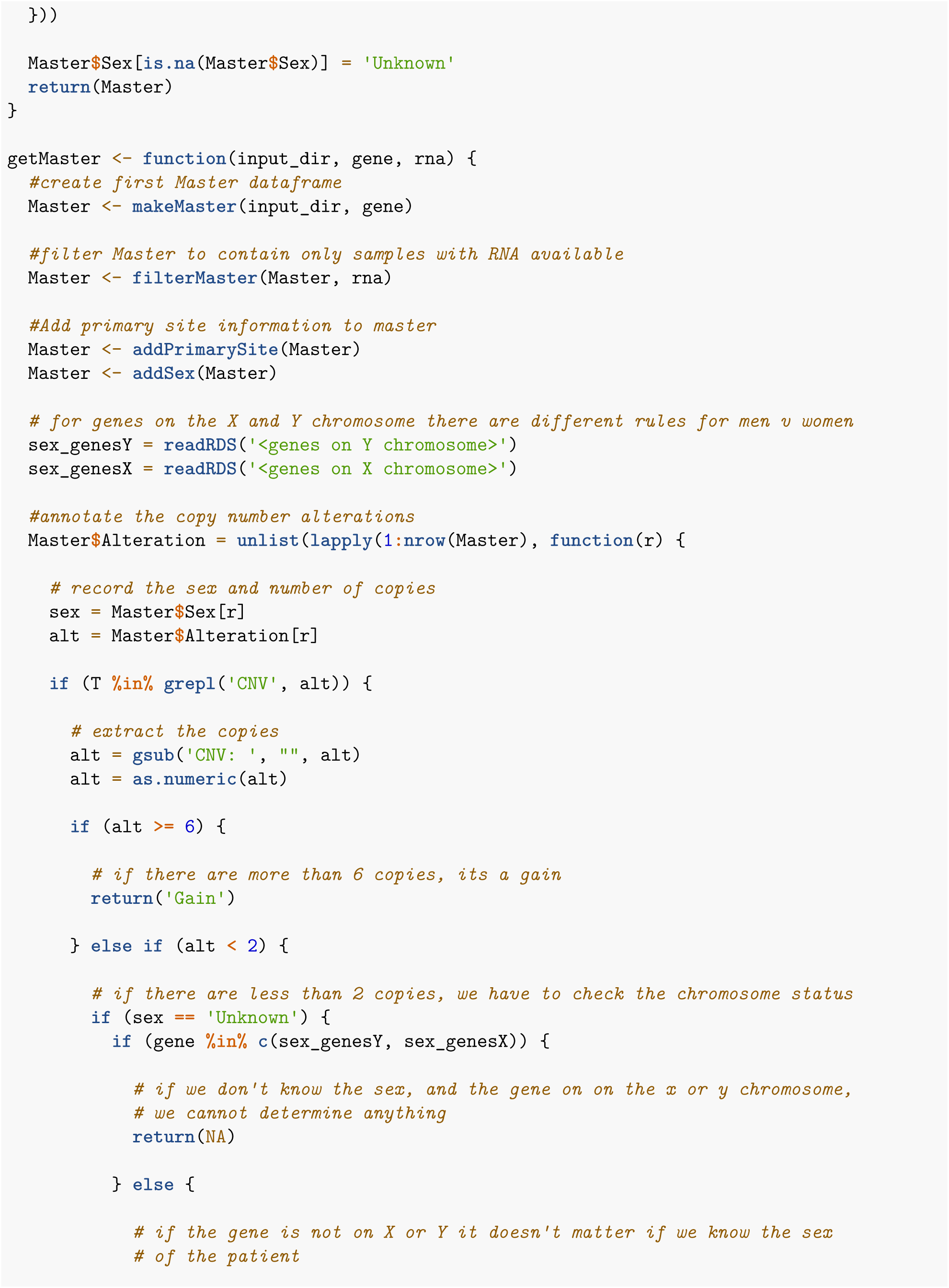

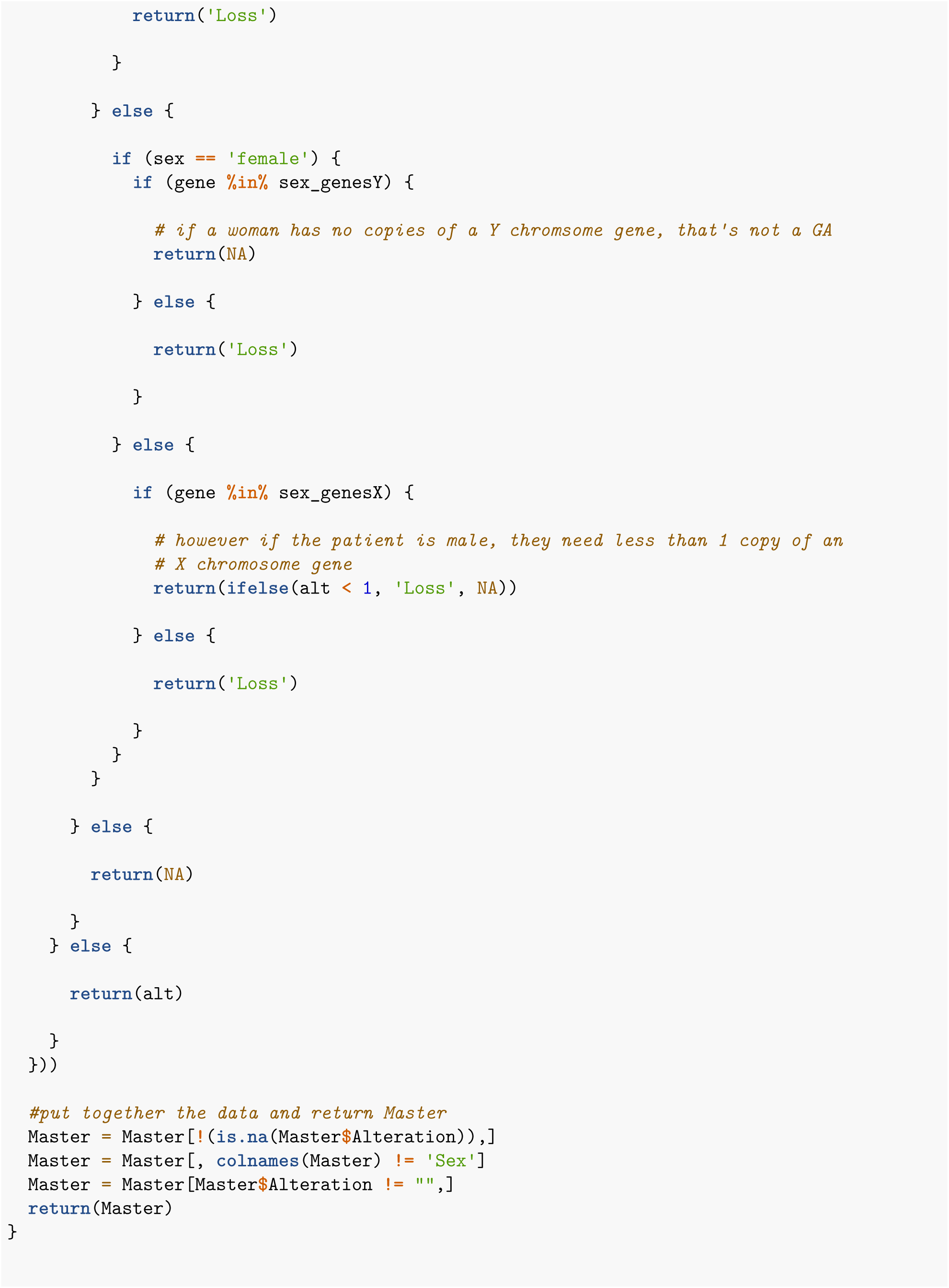

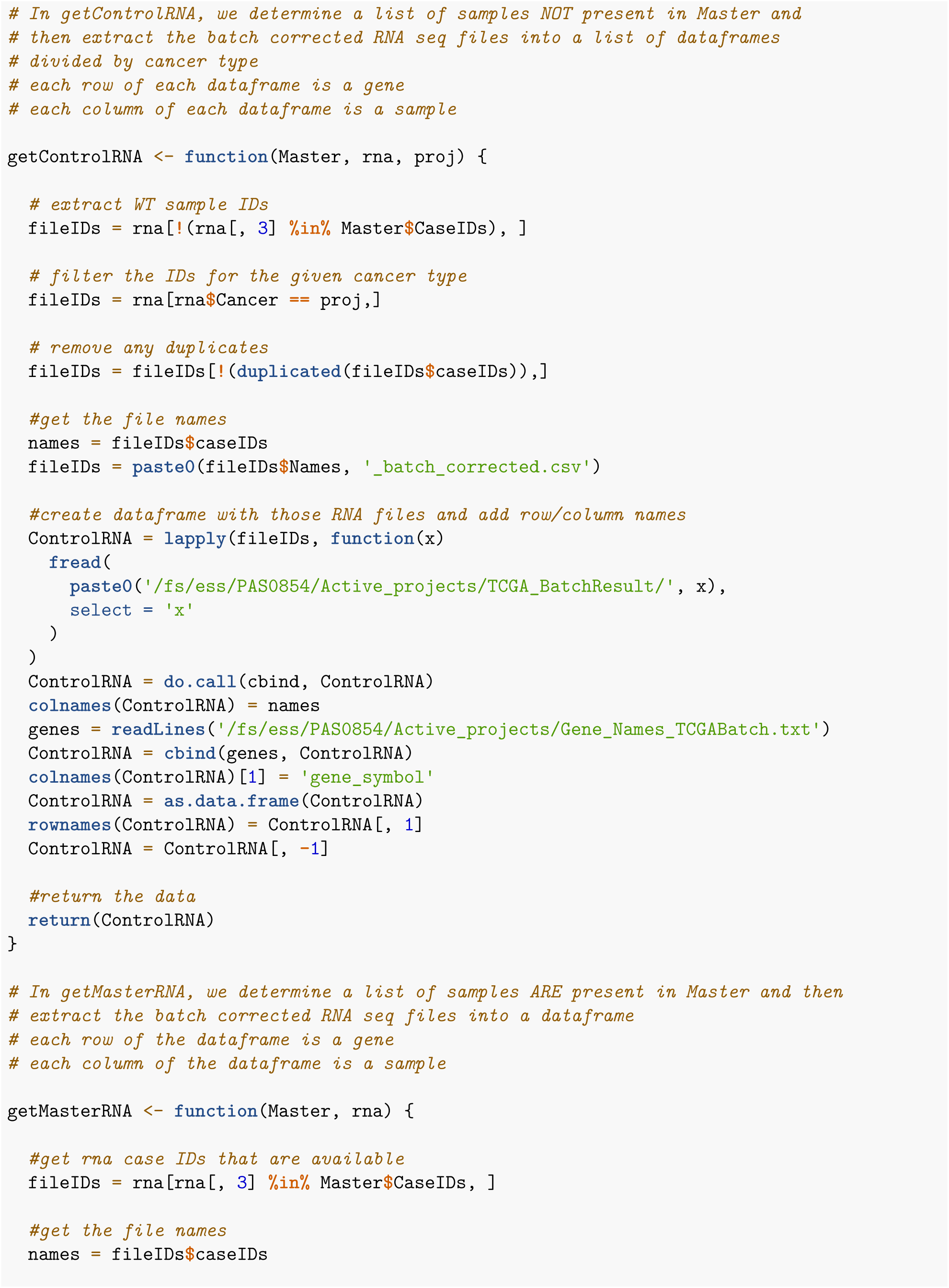

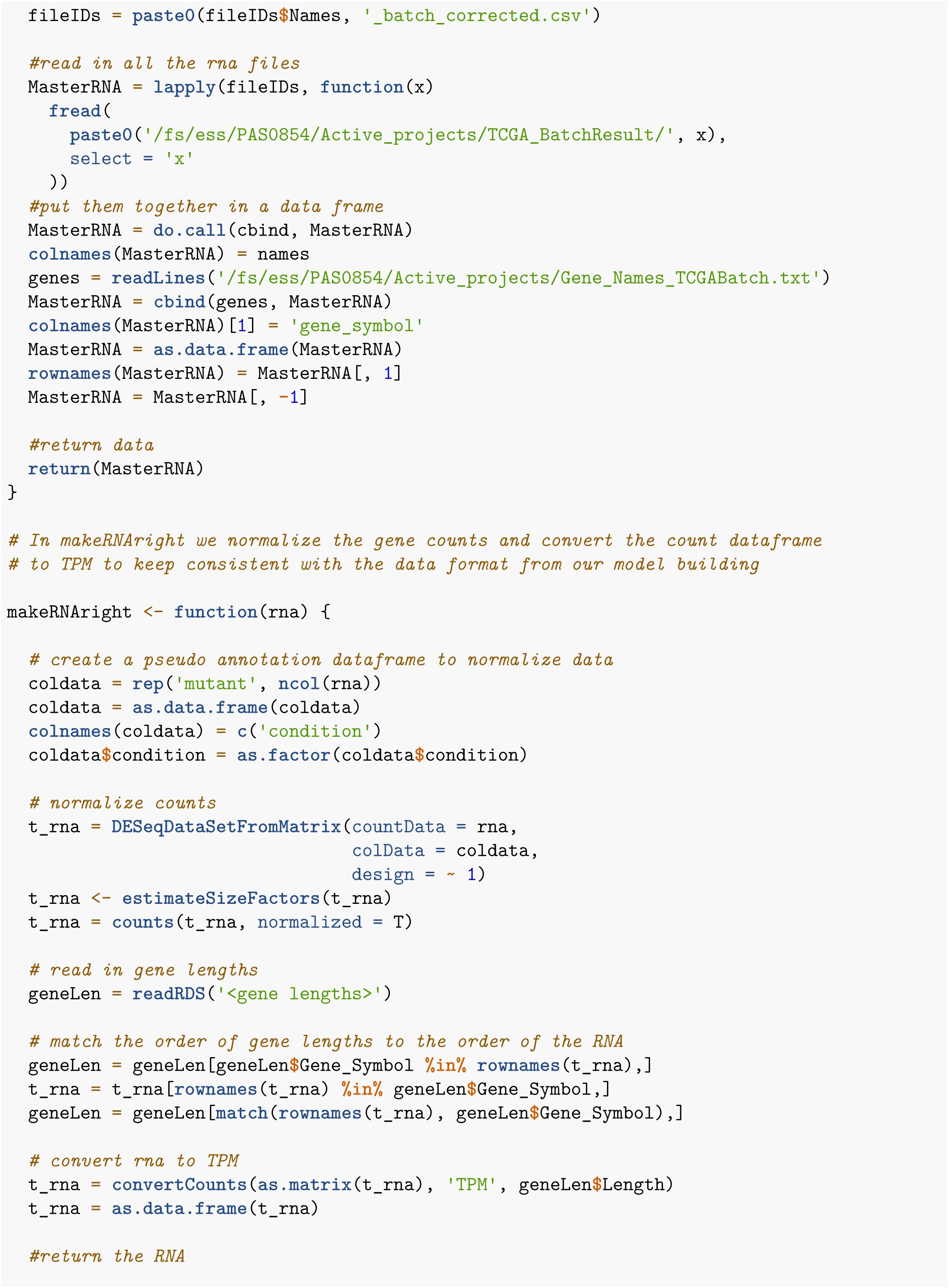

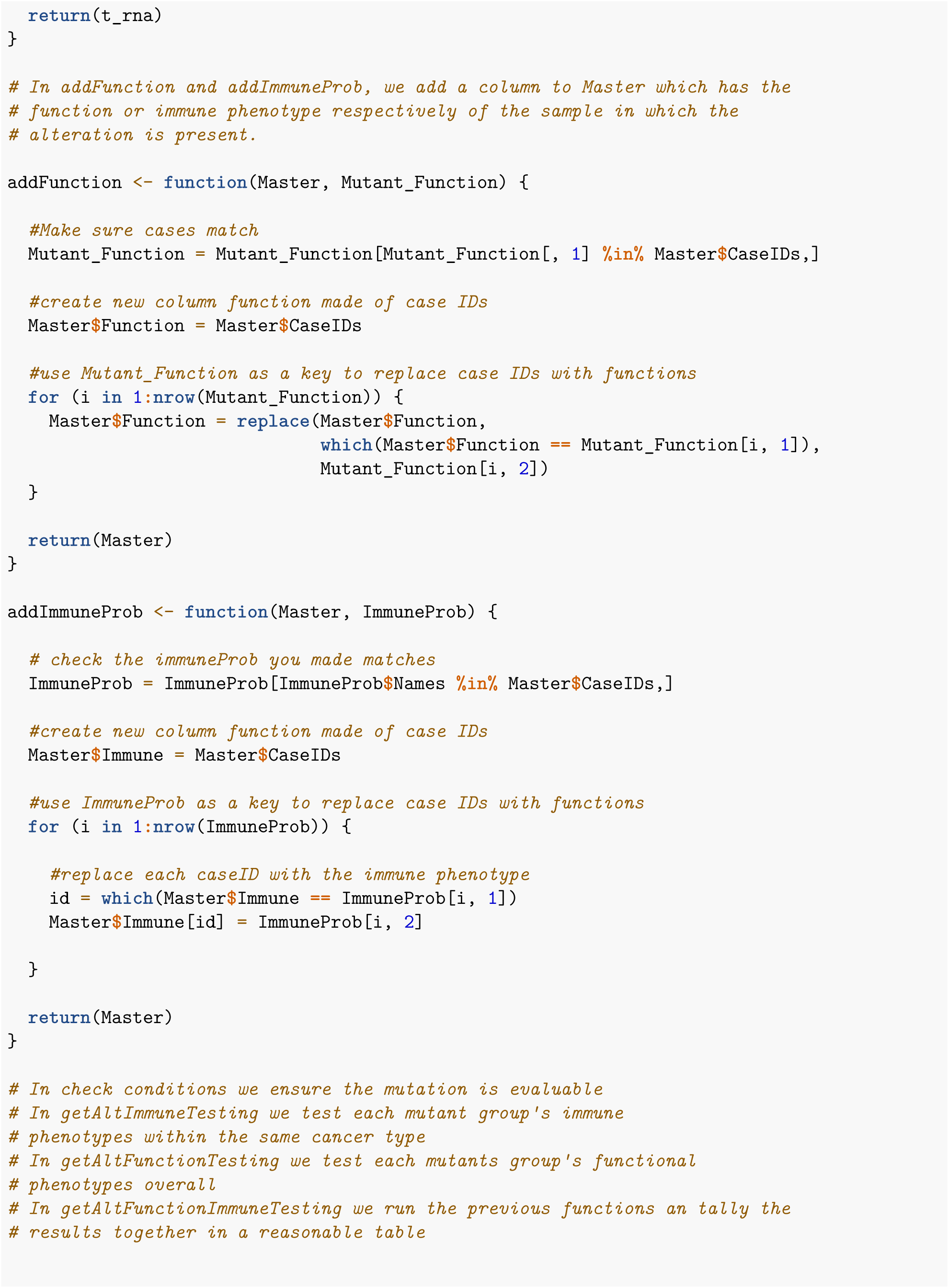

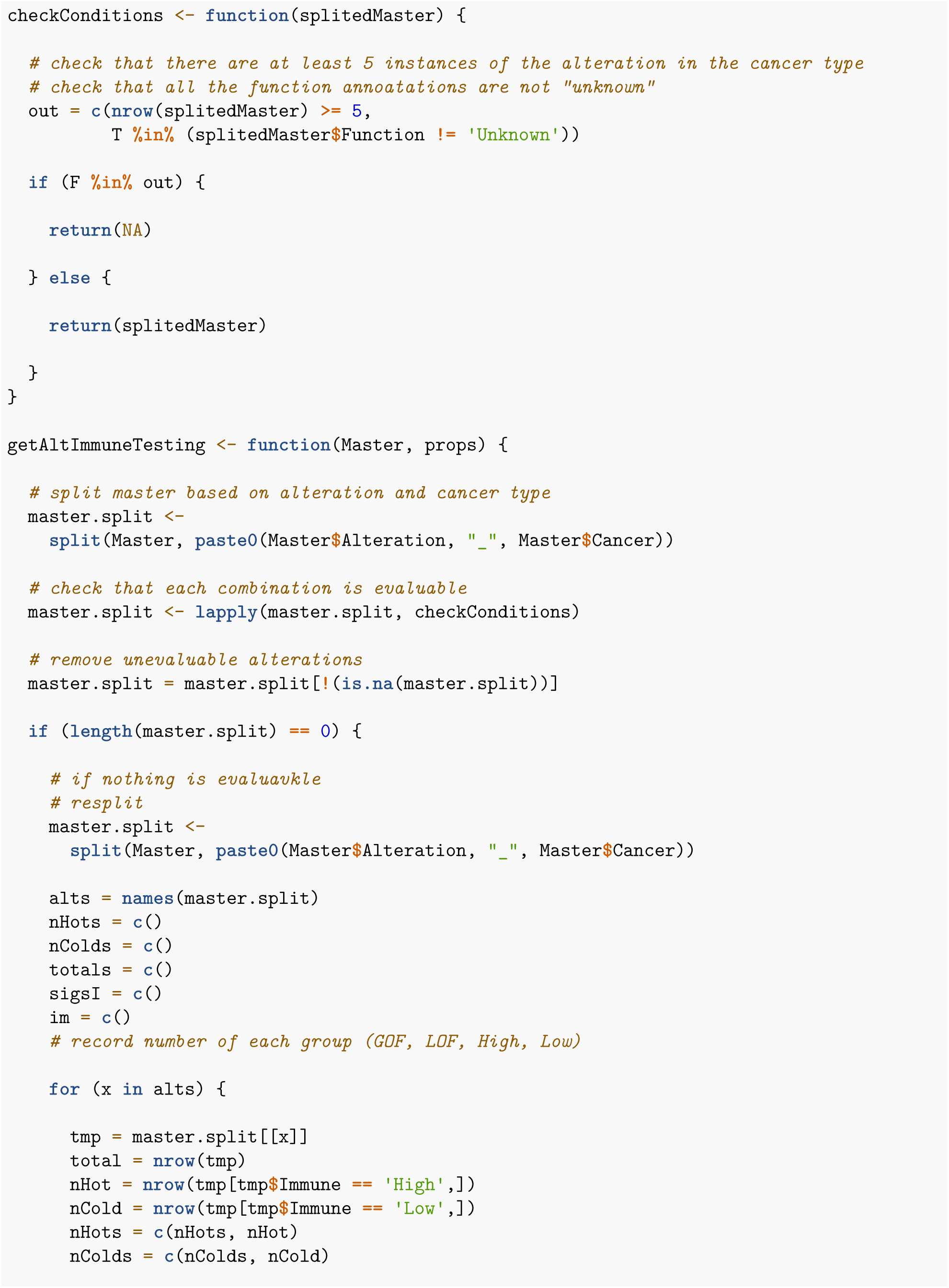

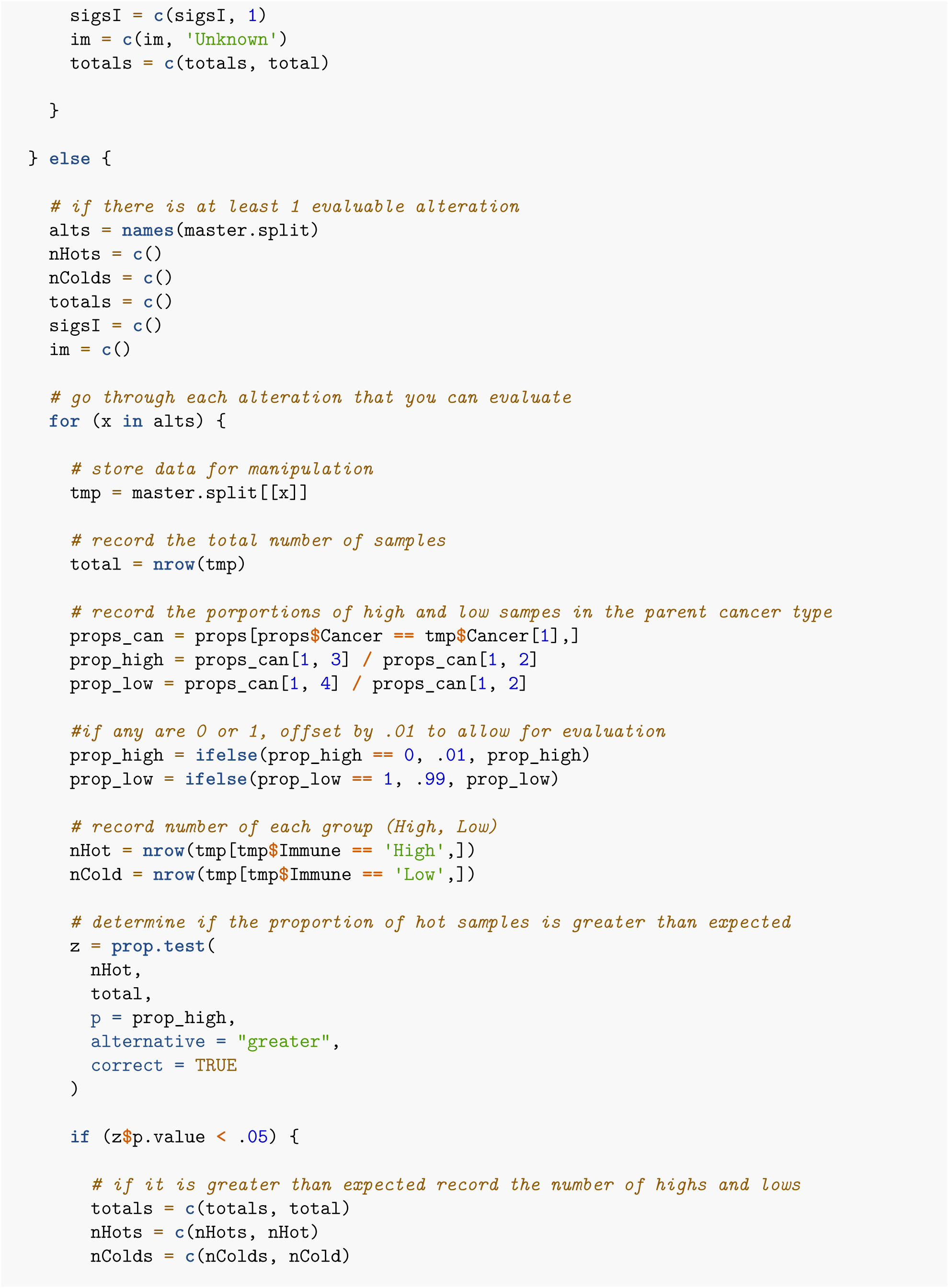

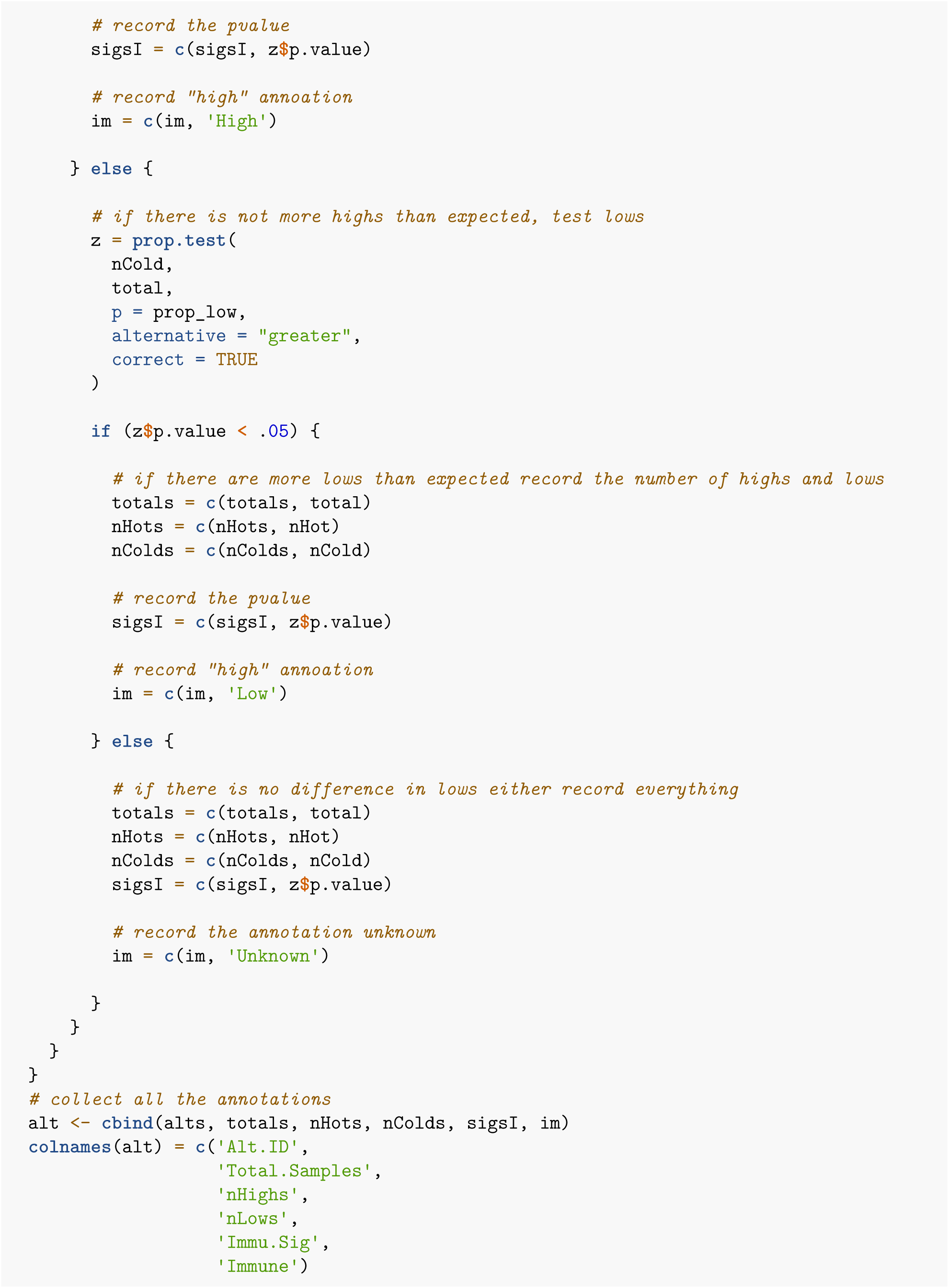

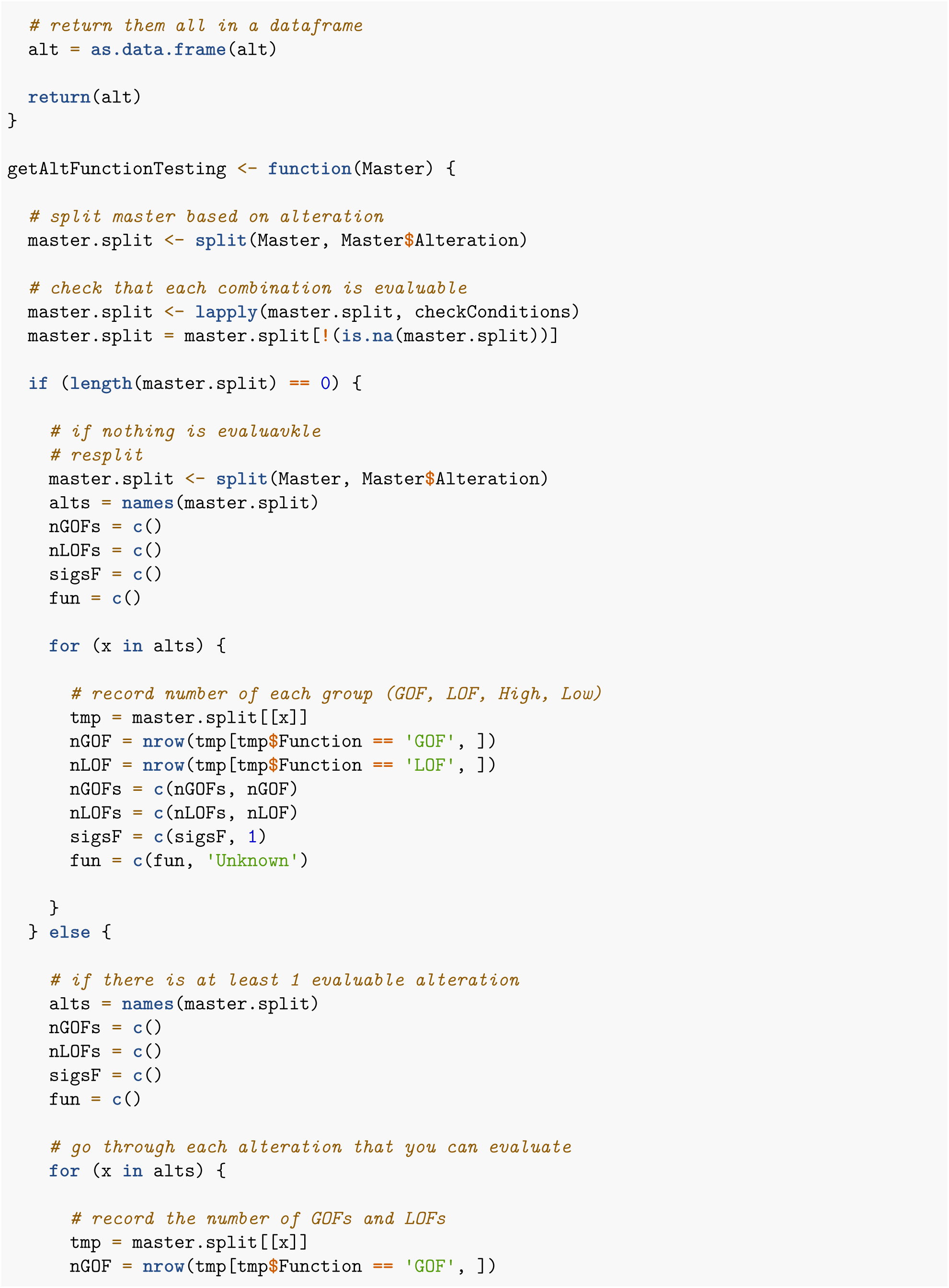

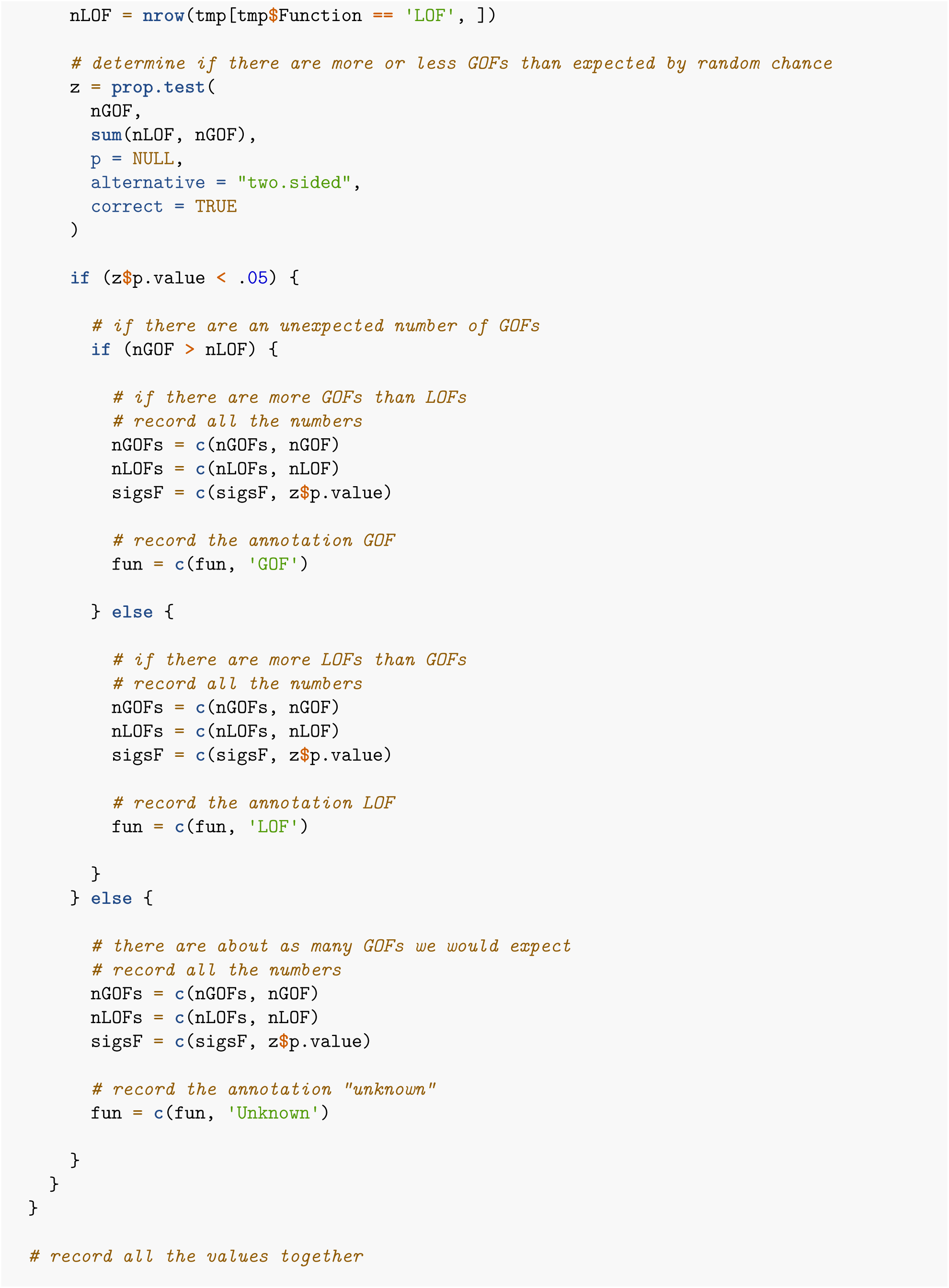

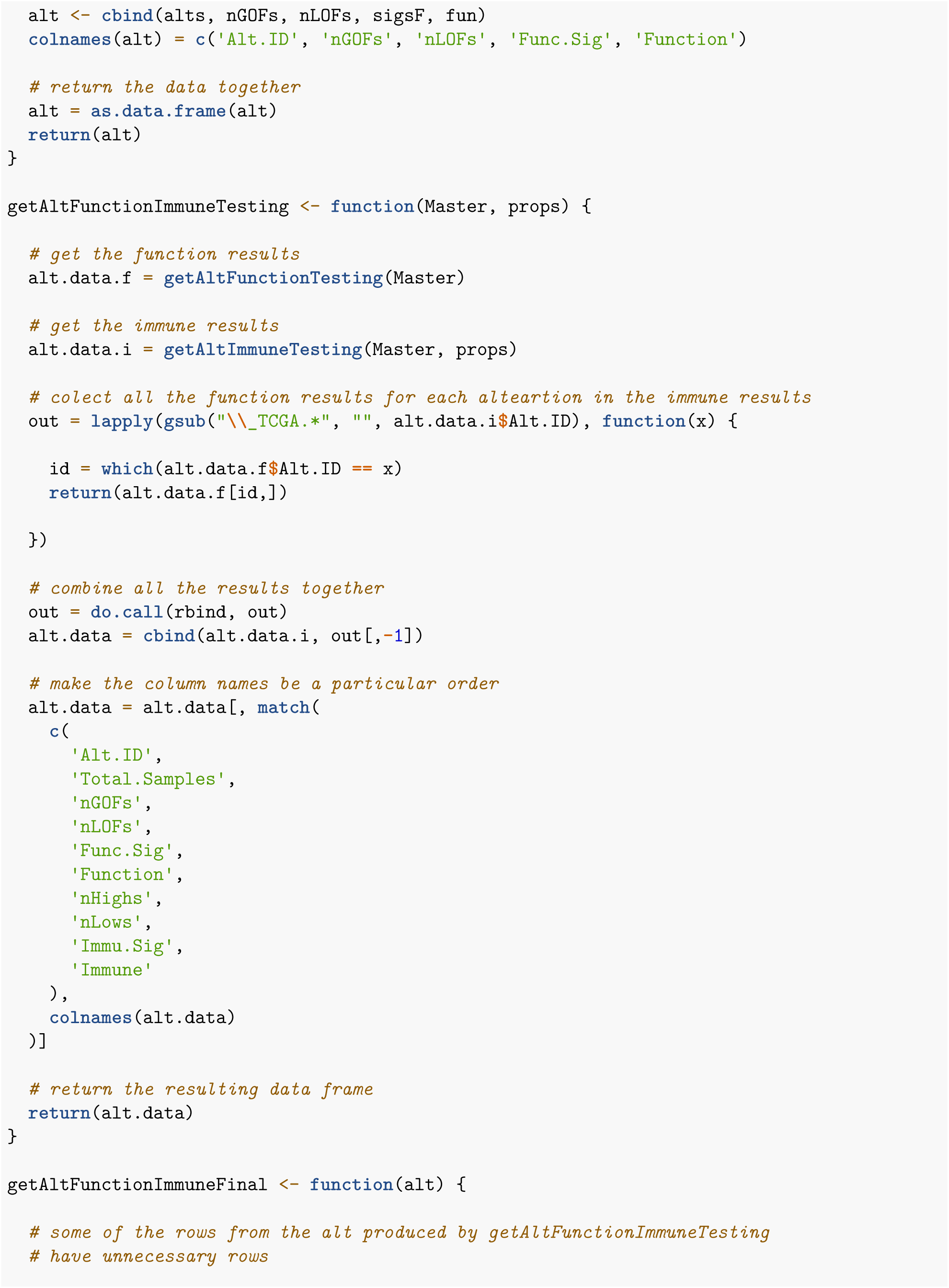

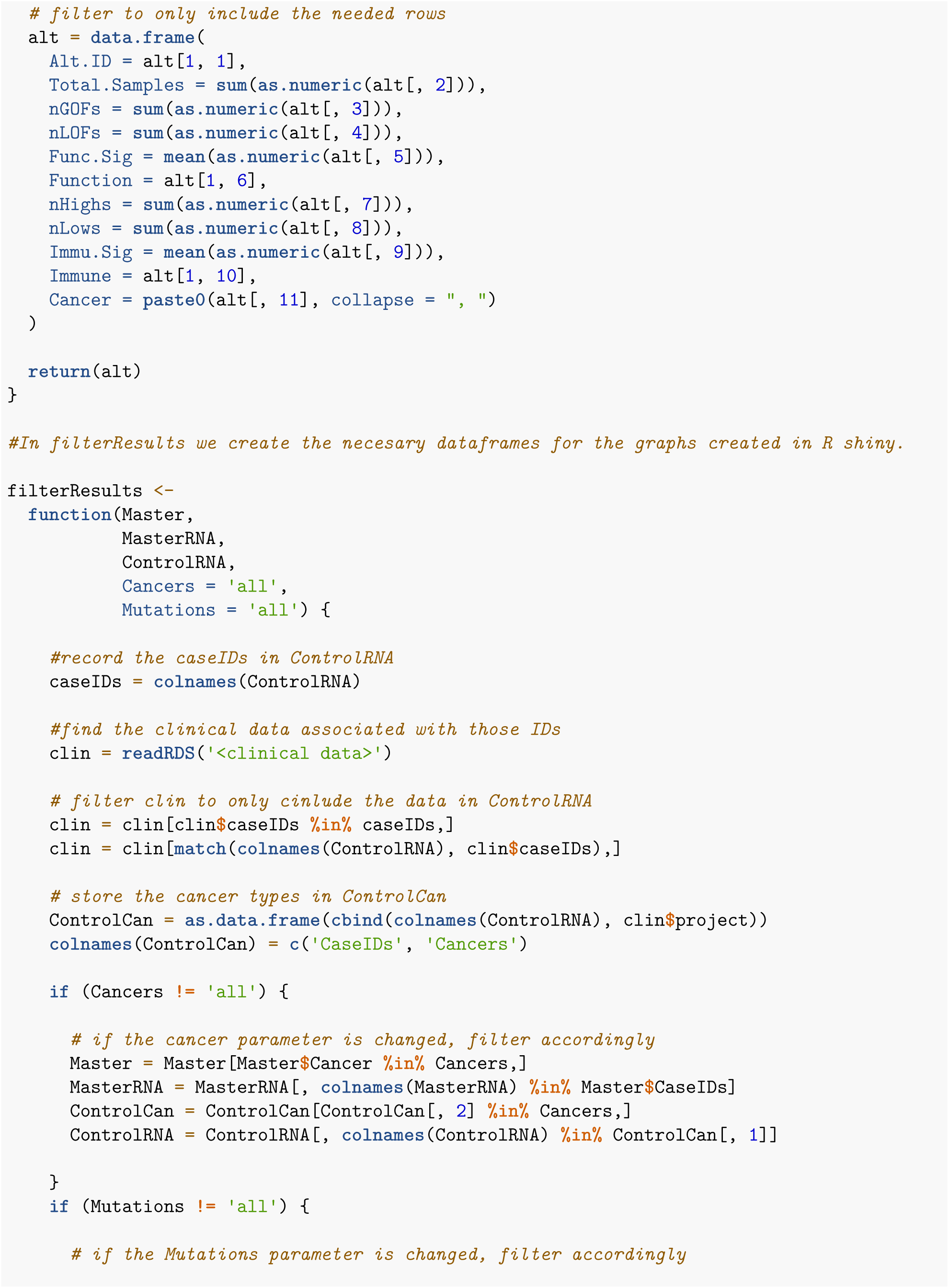

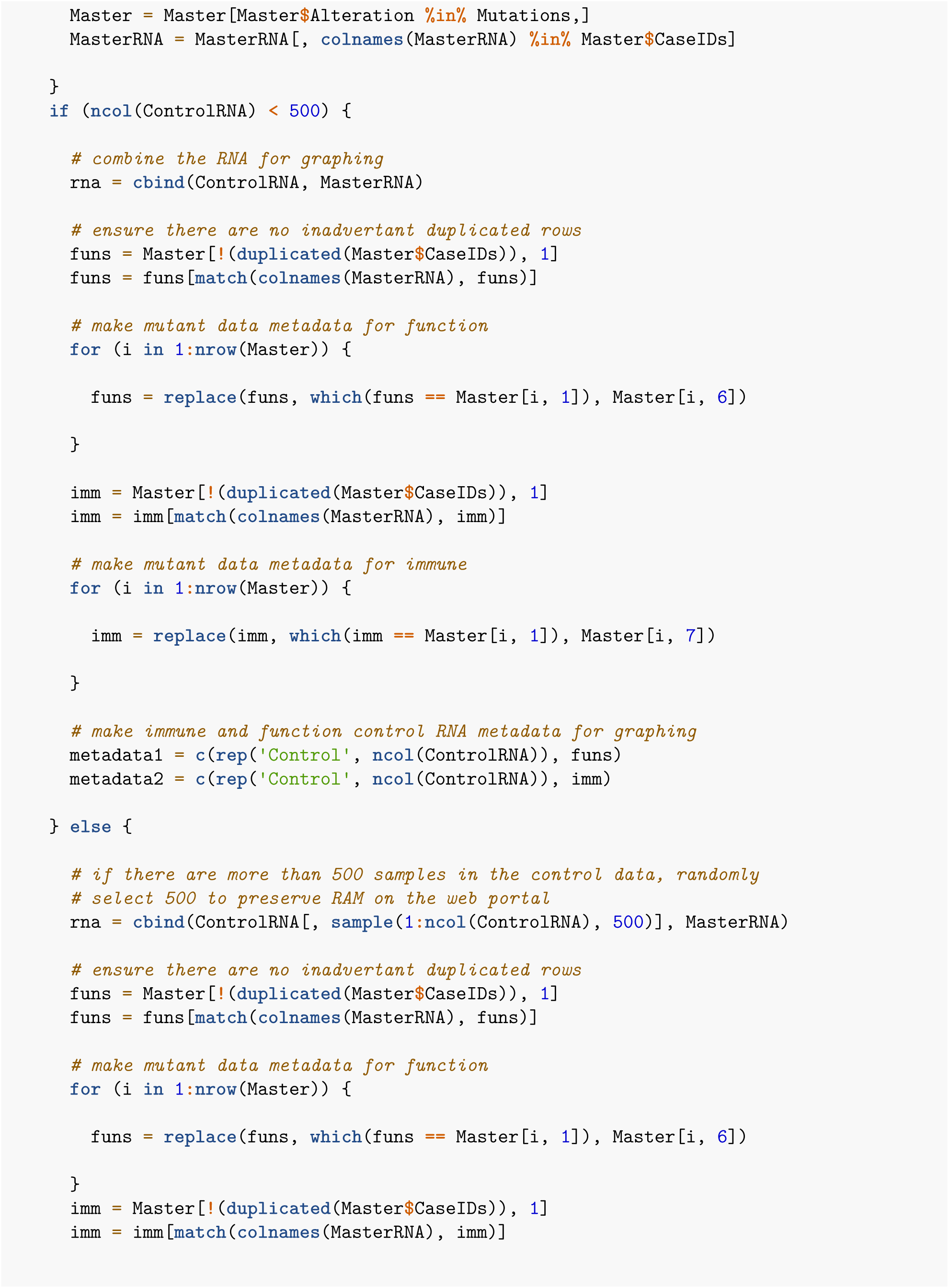

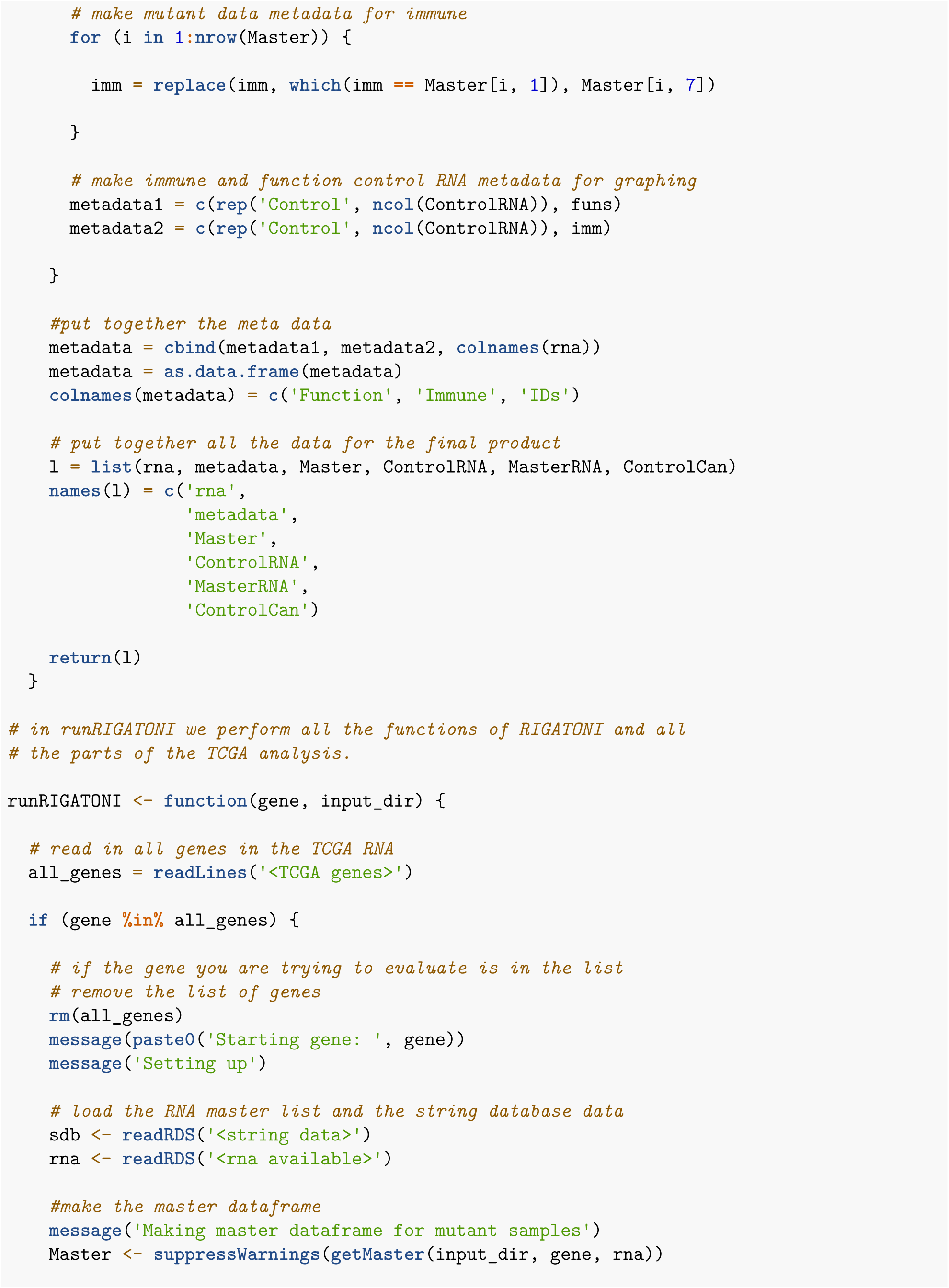

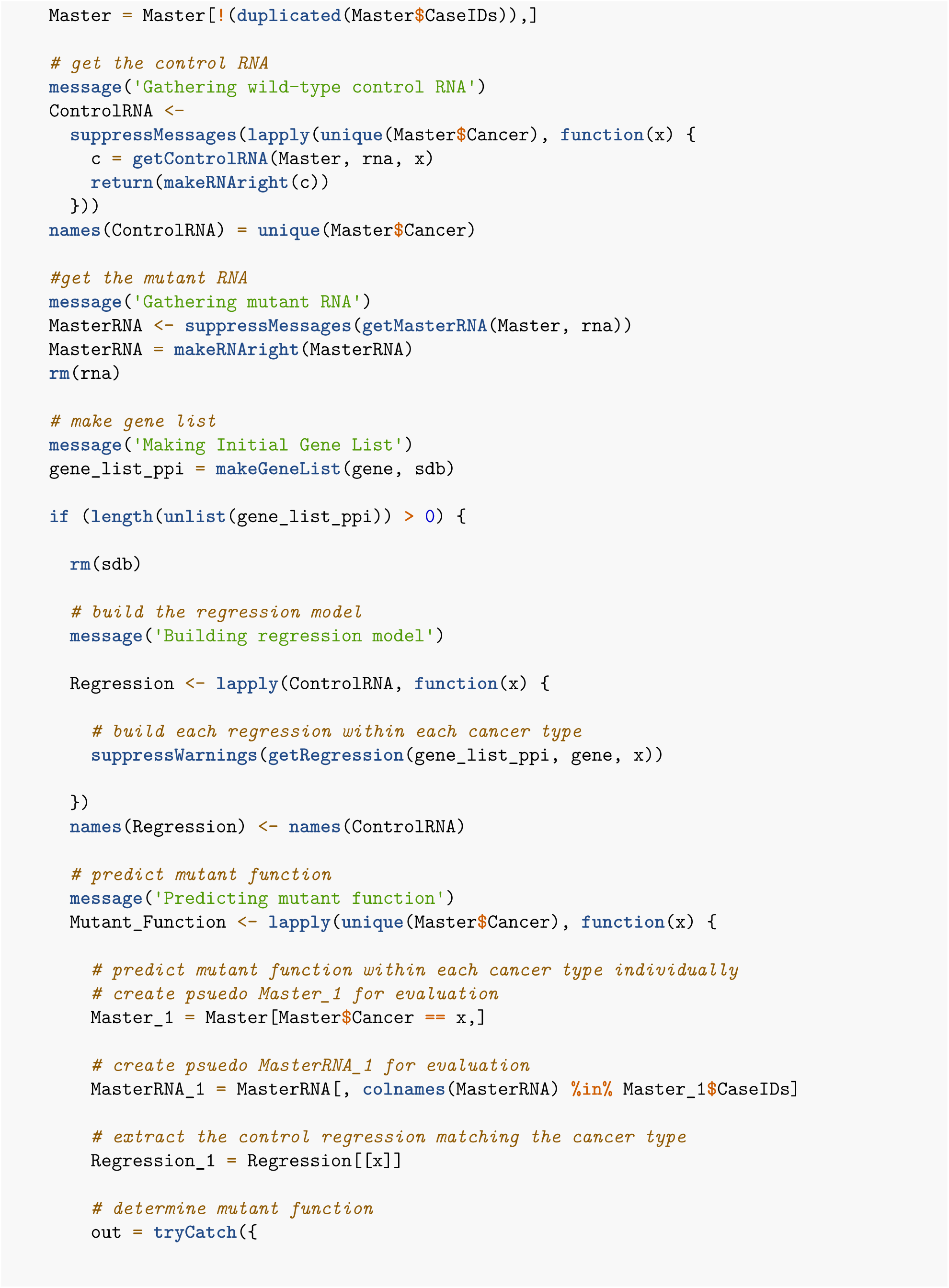

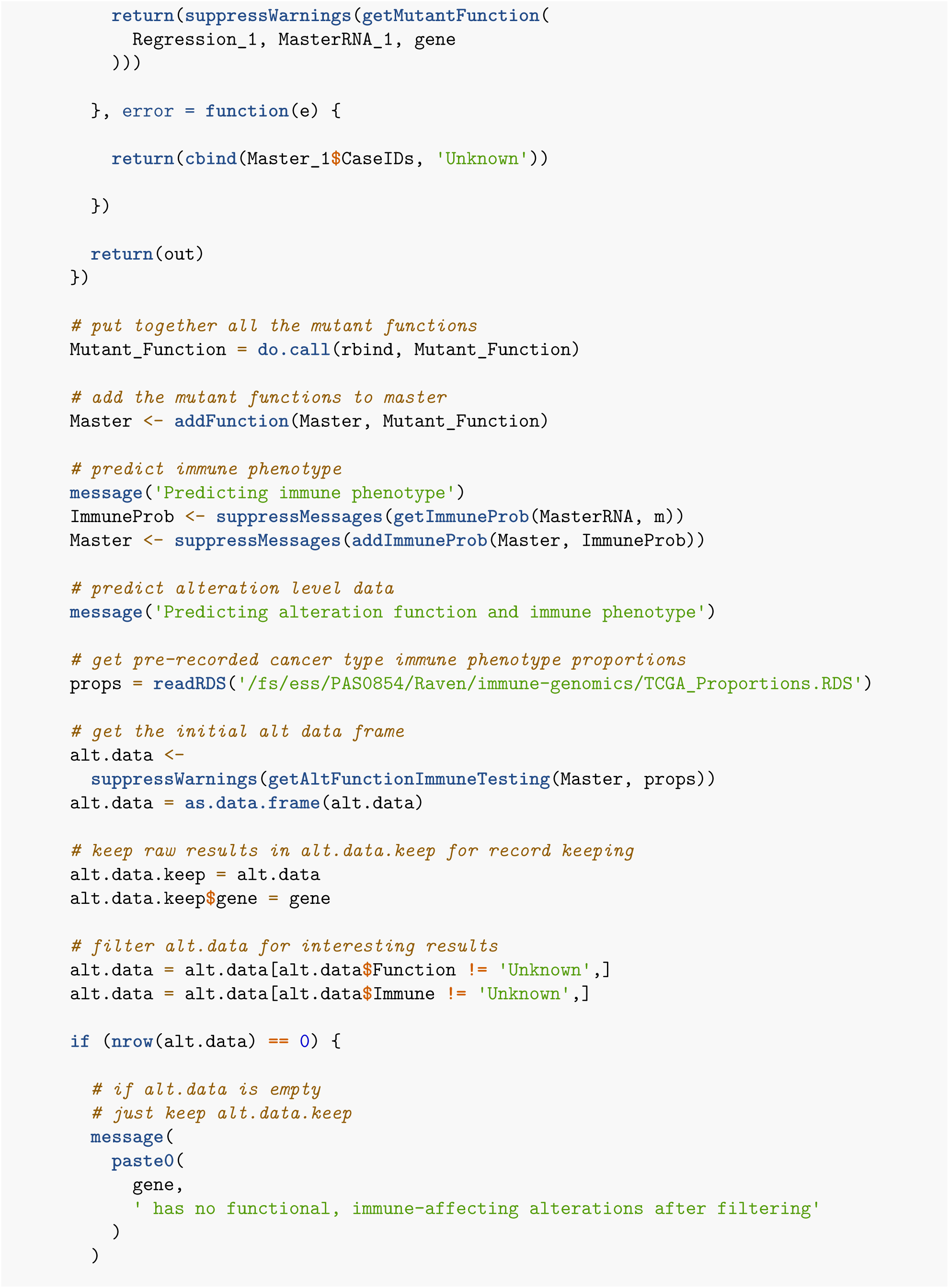

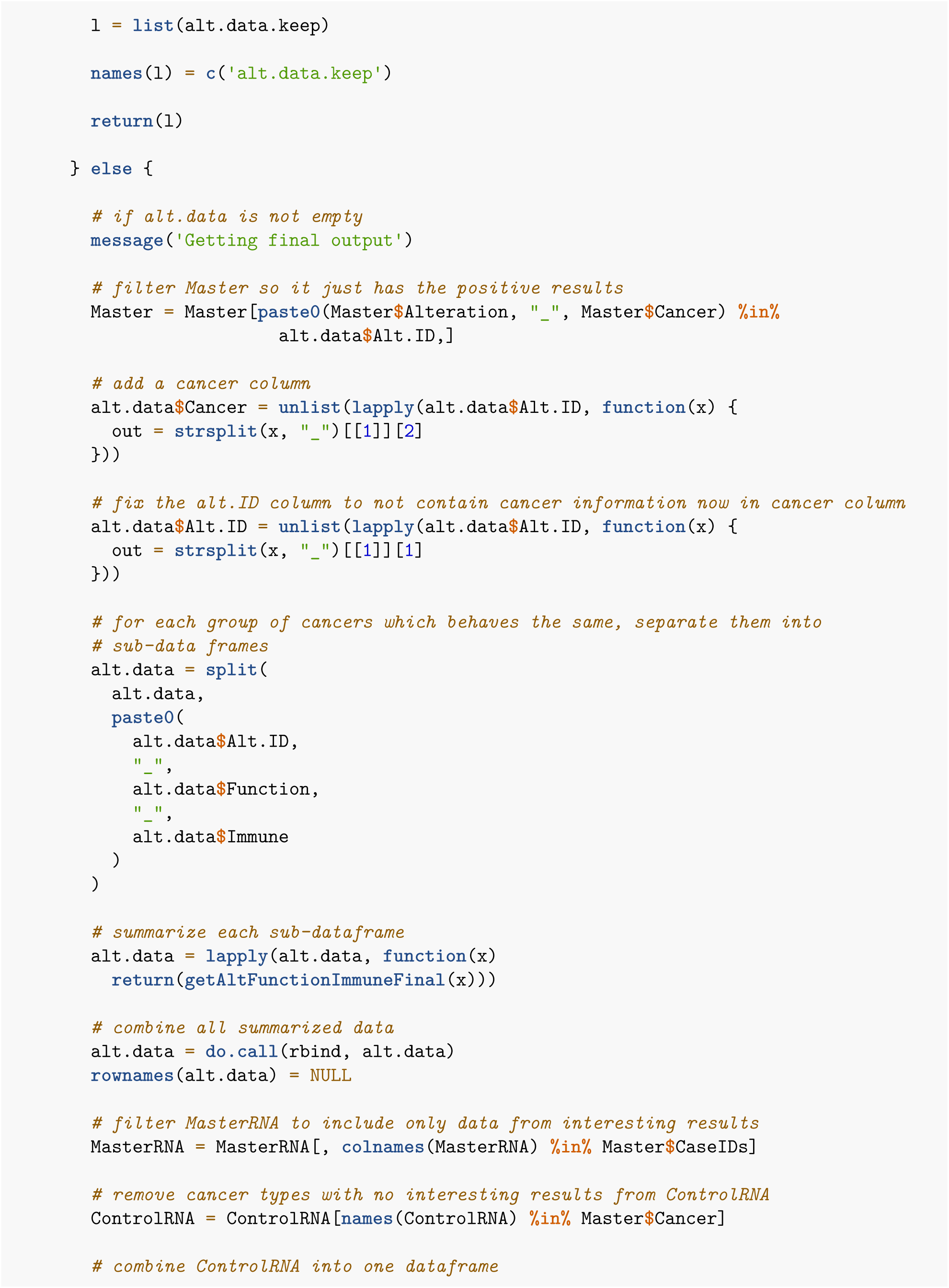

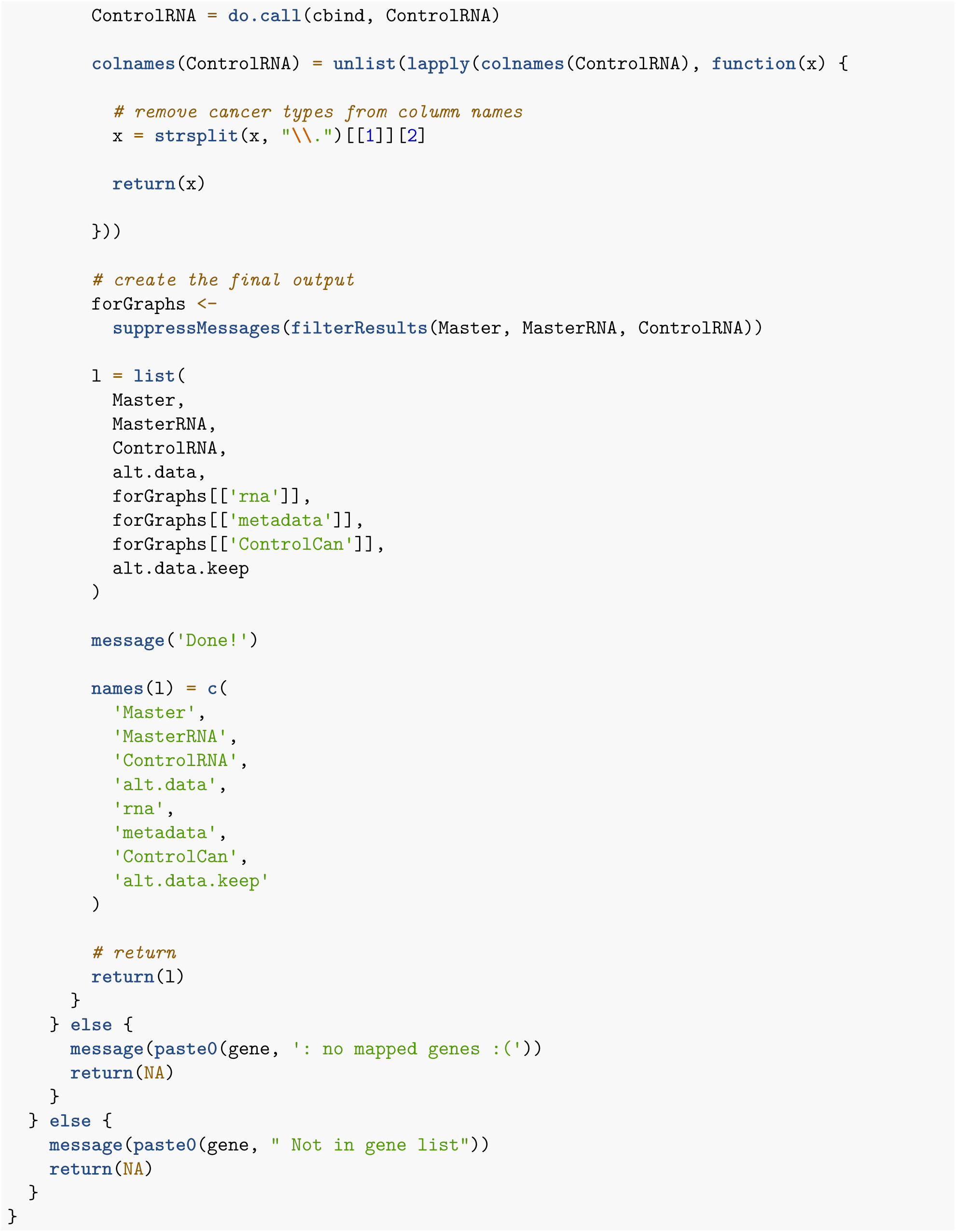

### 6.2: Run RIGATONI for all genes

In order to run RIGATONI for all genes, I created a plain text file with a new gene to run on each line of the file and two batch scripts to run each gene through my pipeline in parallel.

In the first batch script, I run each batch job individually and ensure I do not excede batch limits.

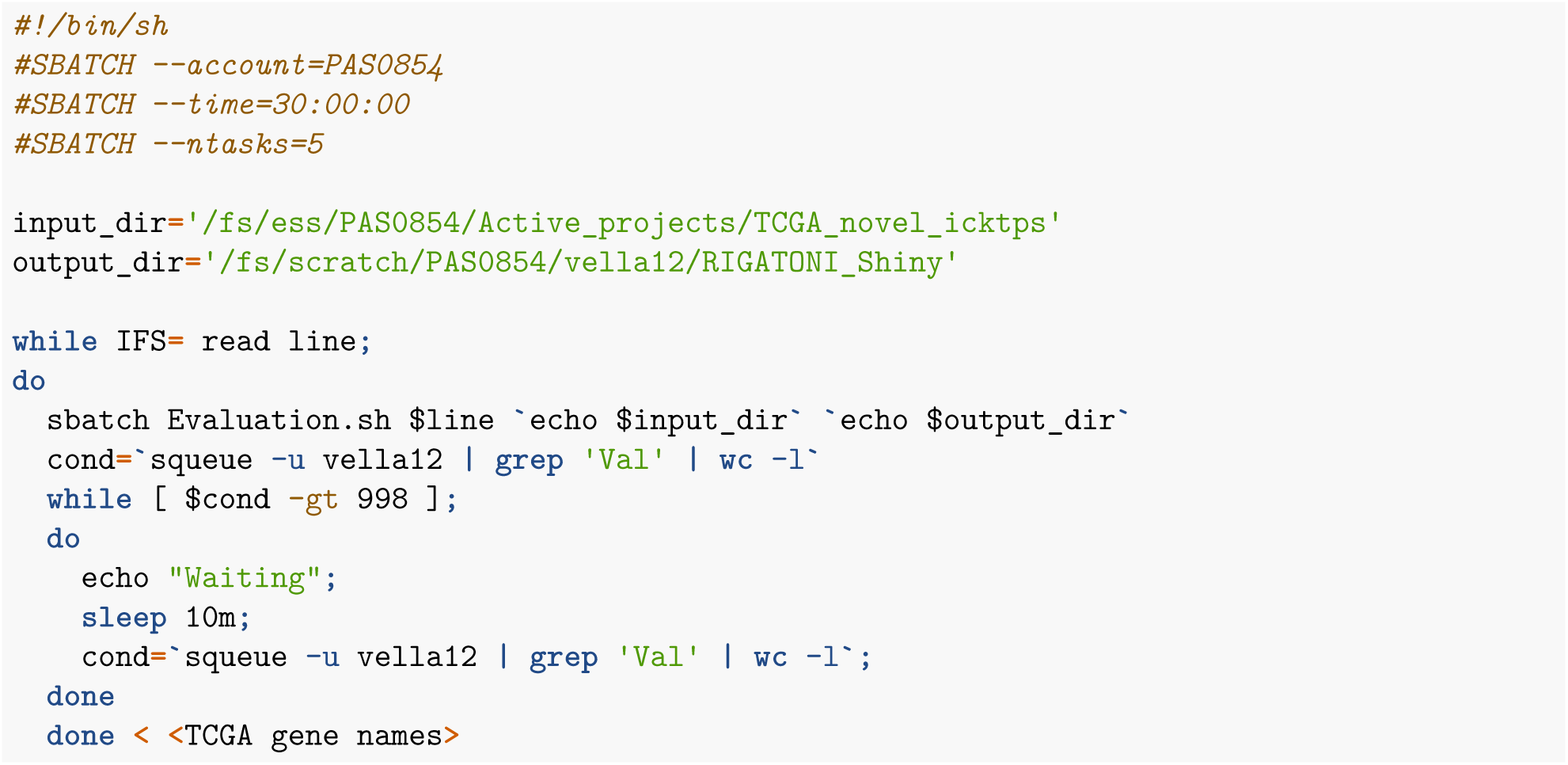

In the next batch script, I simply run the evaluation script in R described in detail in 5.1

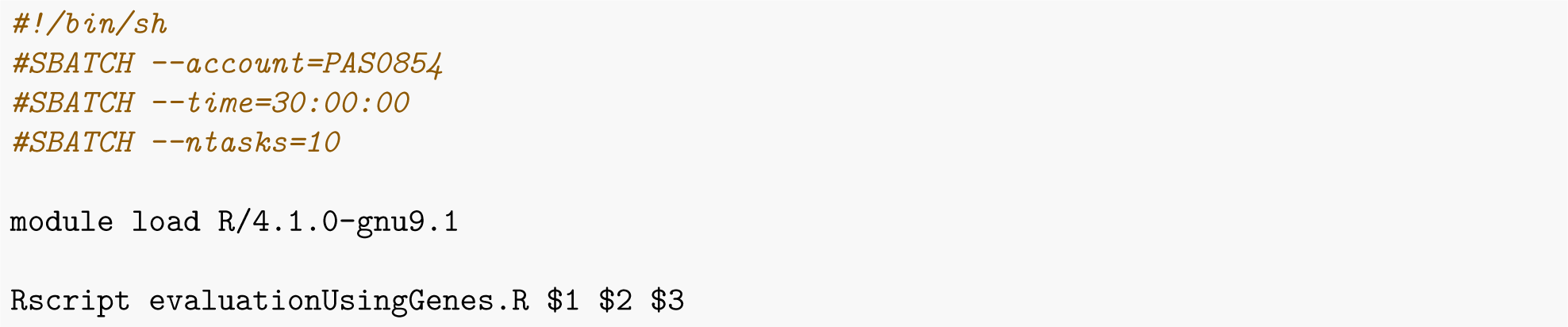

The evaluationUsingGenes.R script simply loads each function described in 5.1, and the R package described in 4 and then executes the following code:

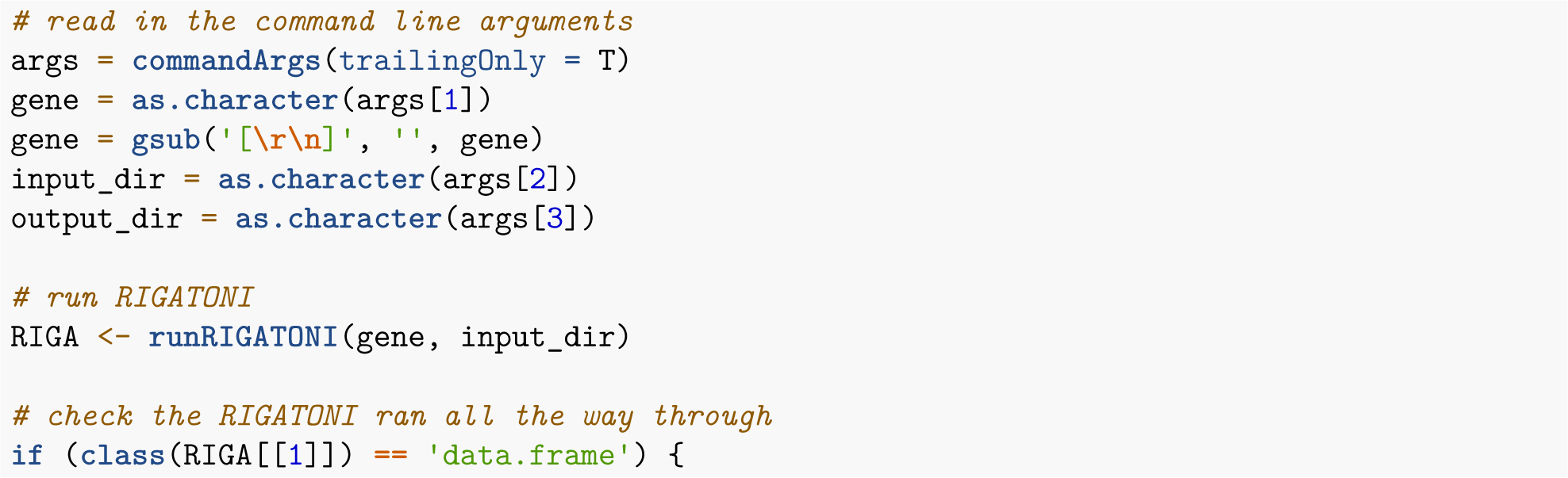

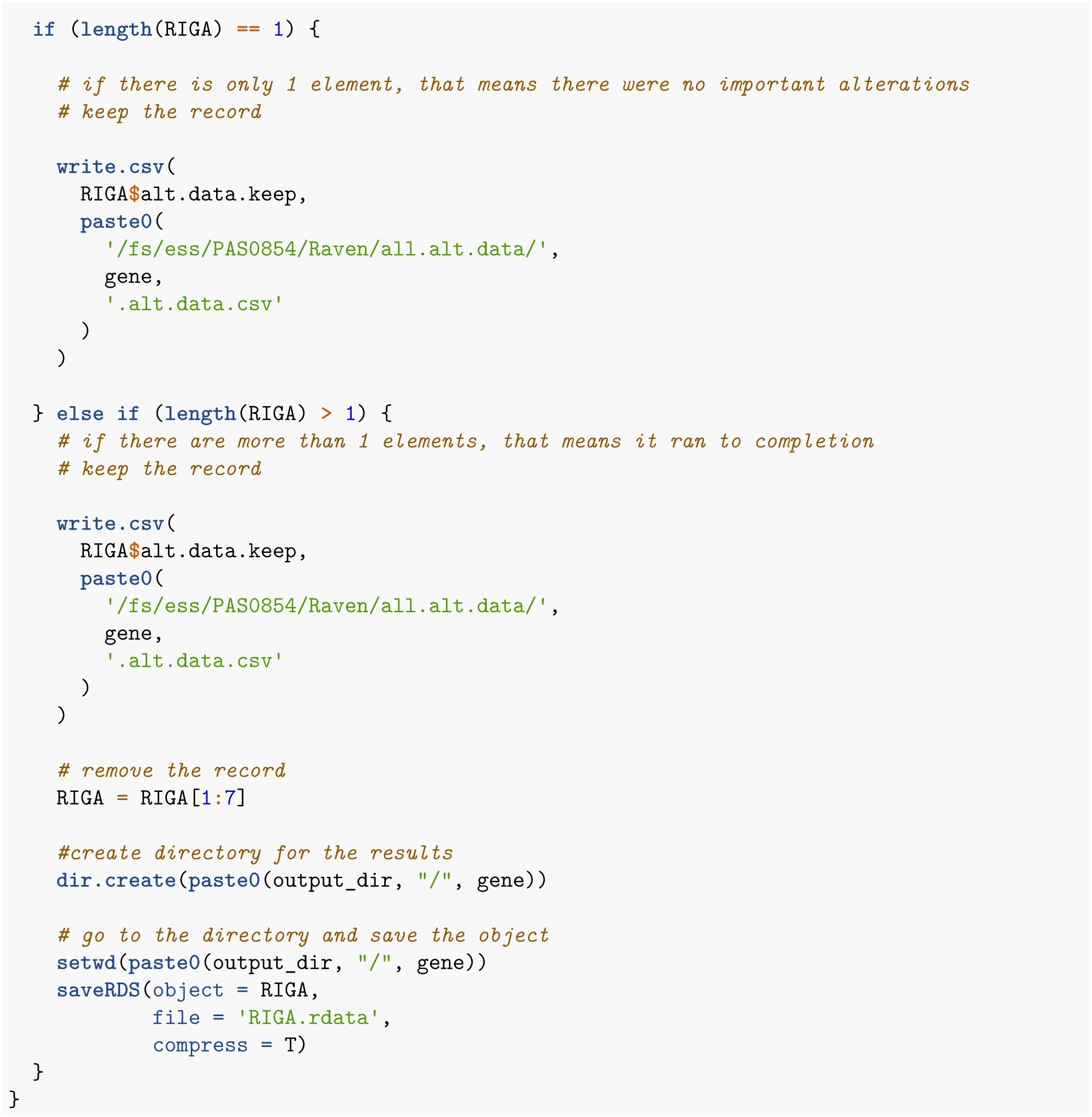

## Part 7: Text mining (Fig 4)

We extracted all the results from the Pan TCGA analysis and then performed text mining to understand how our results compare to existing literature. First we ran the following in linux.

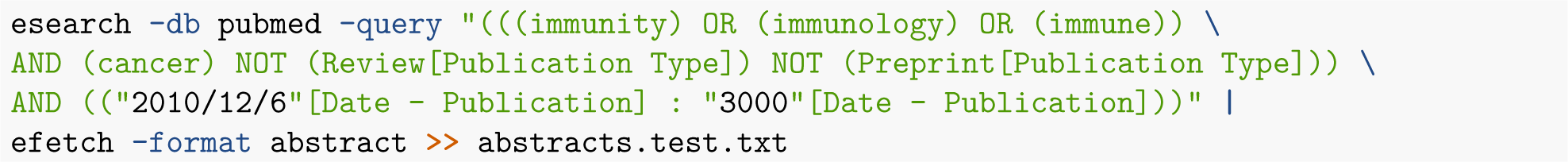

Next we did the following in R

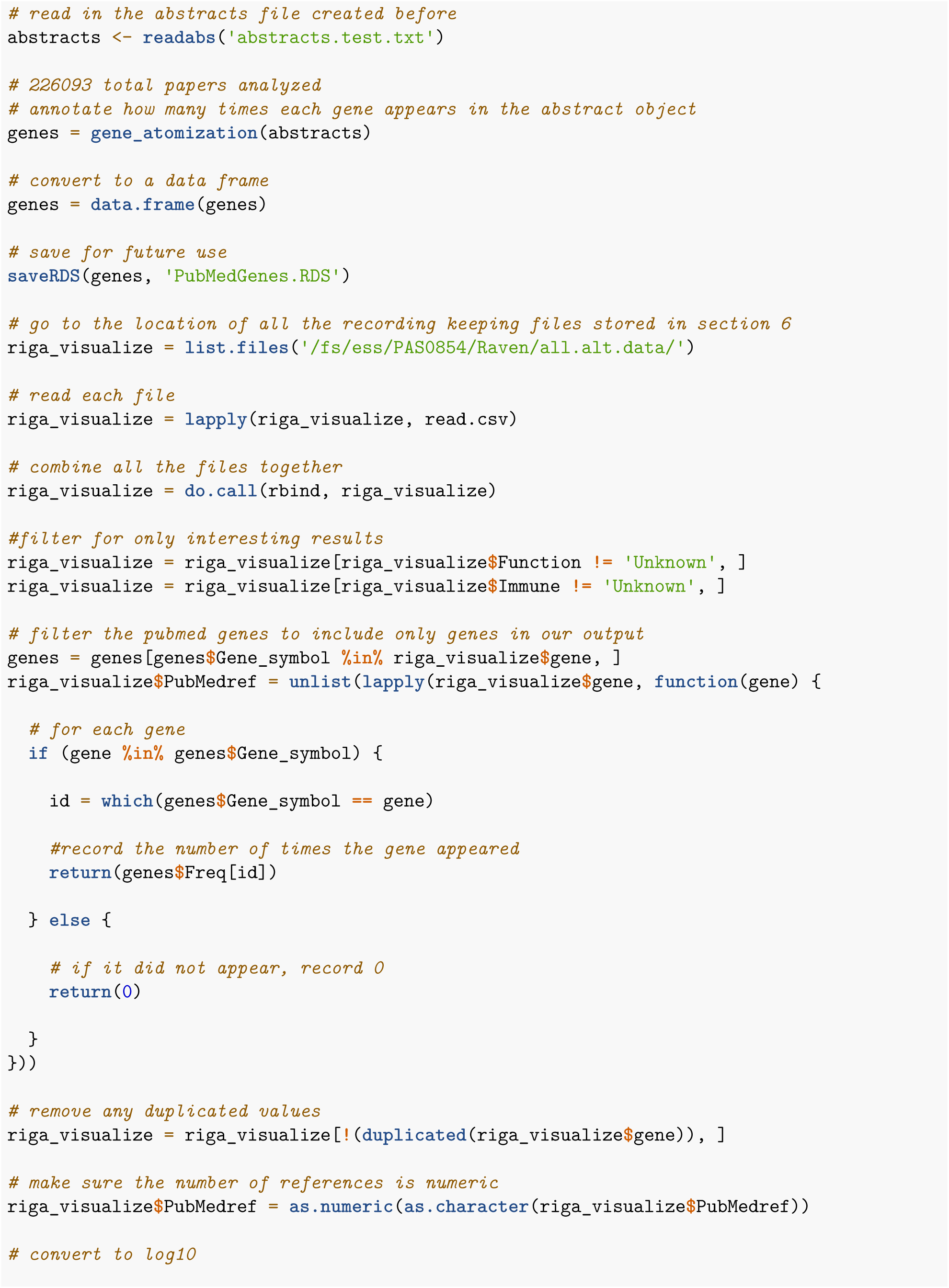

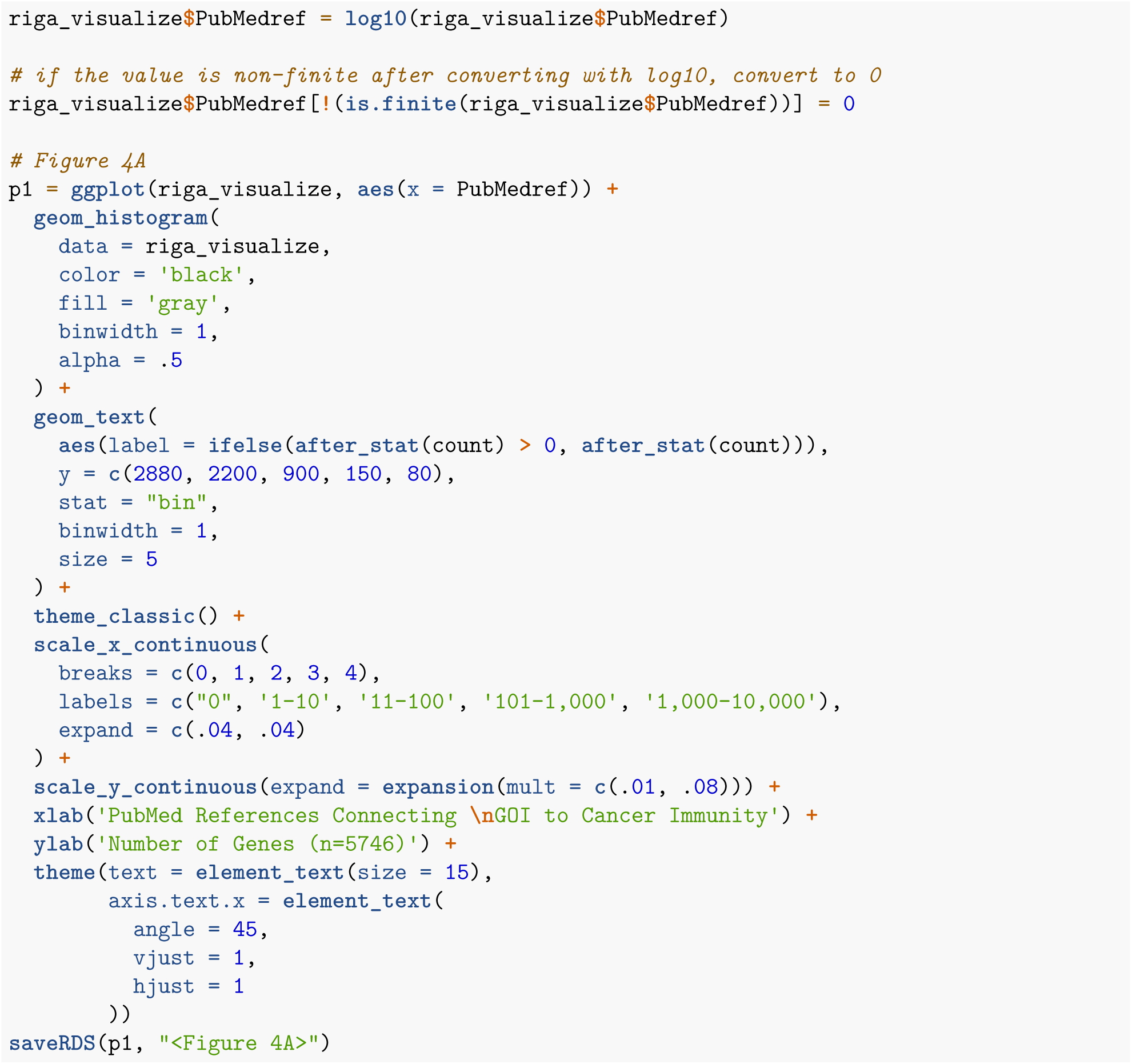

## Part 8: 14q deletion in renal cell carcinoma (Fig 4)

All data shown here is from the paper found at “https://aacrjournals.org/cancerres/article/83/5/700/716683/Integrative-Single-Cell-Analysis-Reveals” The bulk RNAseq results for 14q deletions shown in figures 4B and 4C are extracted from the pan-TCGA analysis described in section 6. The single cell analysis is shown here. All data was downloaded from “https://www.ncbi.nlm.nih.gov/geo/query/acc.cgi?acc=GSE207493”

### 8.1 Preprocessing

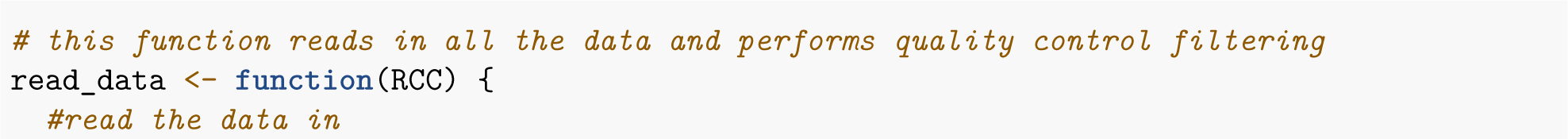

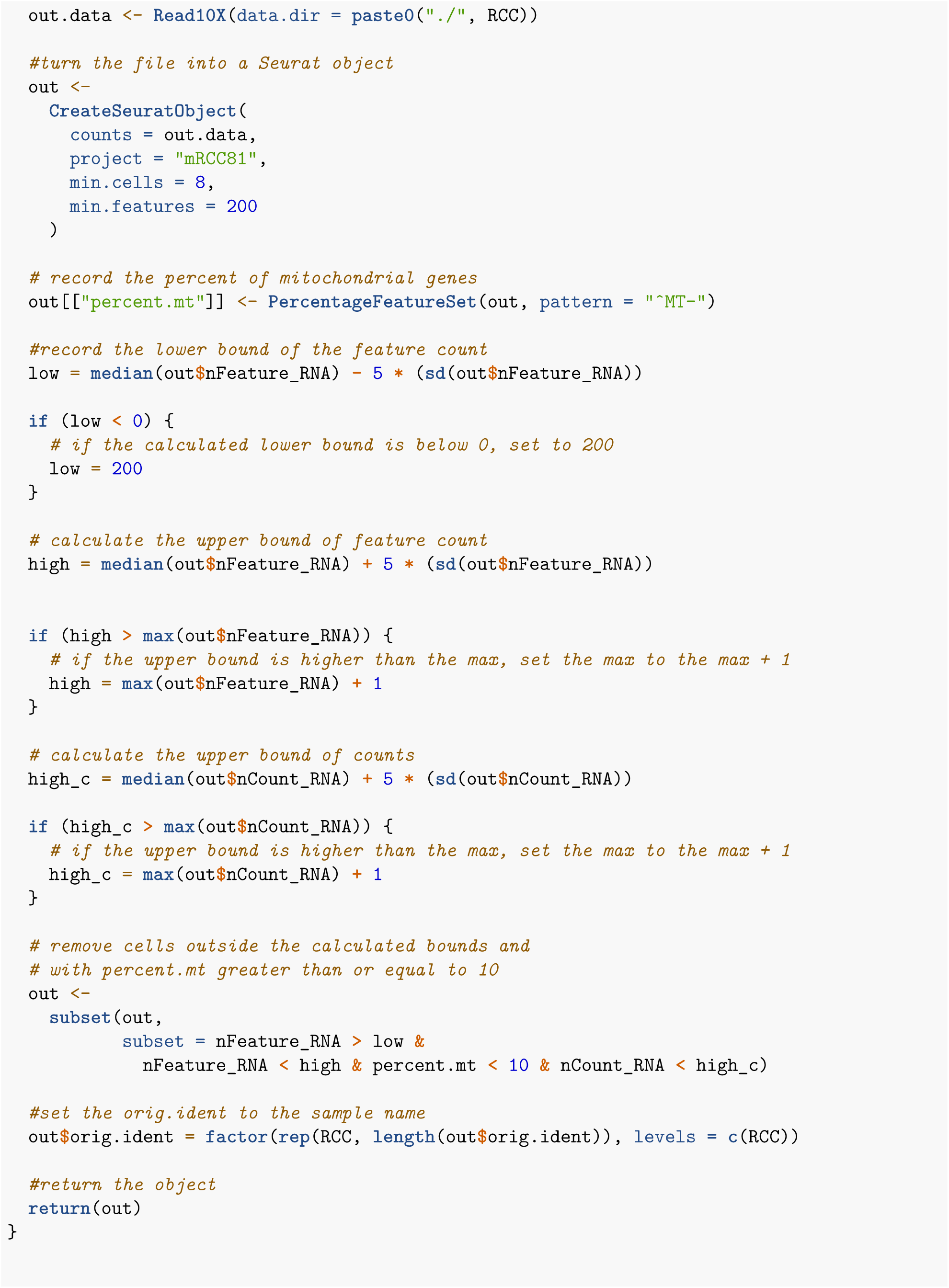

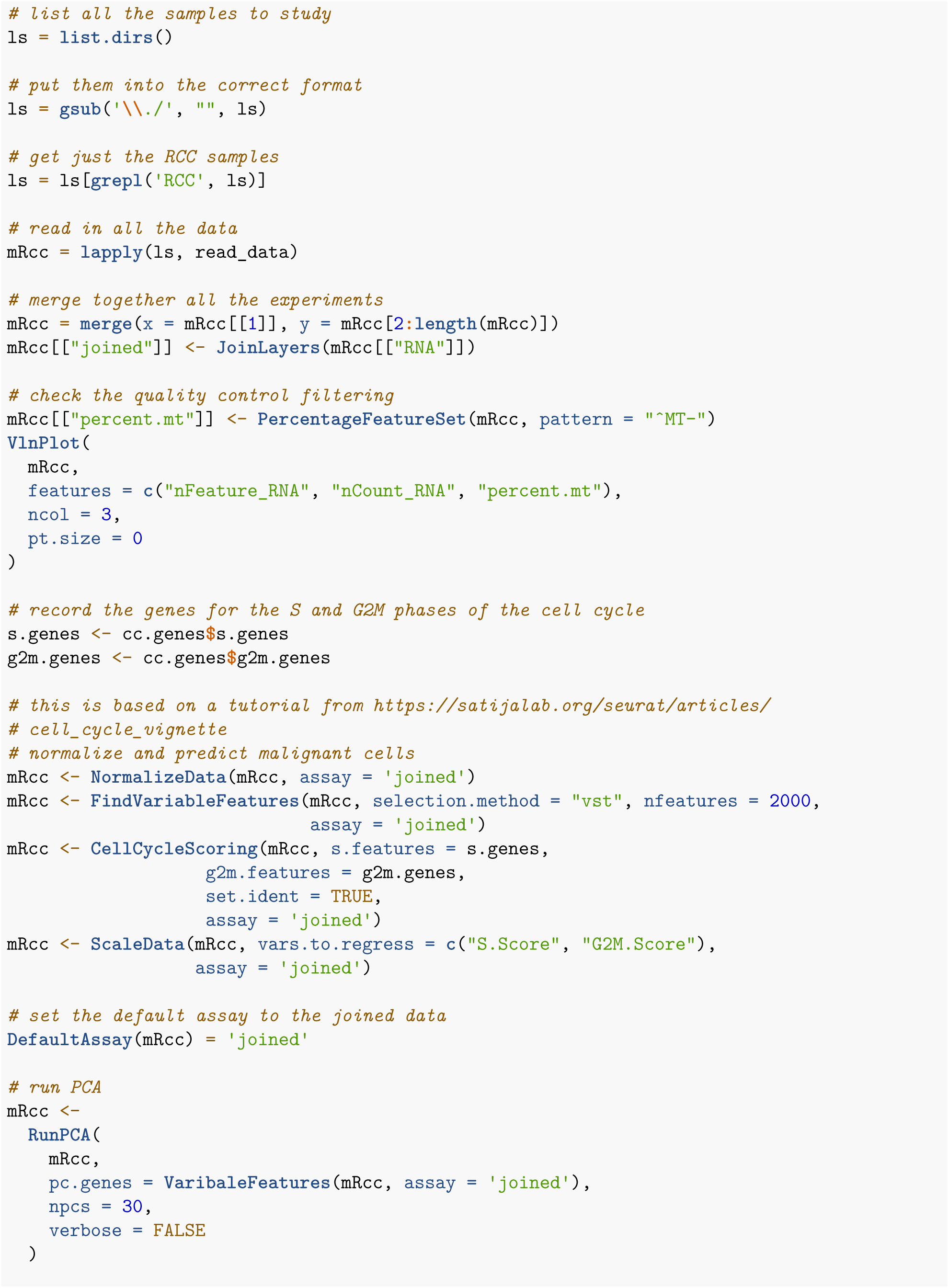

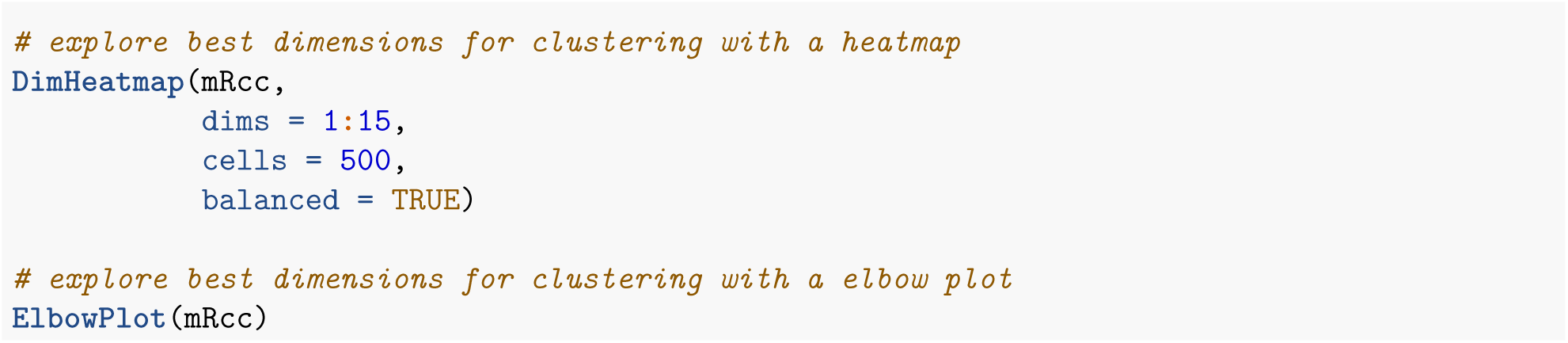

### 8.2 Cell typing for normal and tumor groups

The following is based on tutorials from “https://hbctraining.github.io/scRNA-seq_online/lessons/06a_integration_harmony.html” and “https://github.com/IanevskiAleksandr/sc-type”

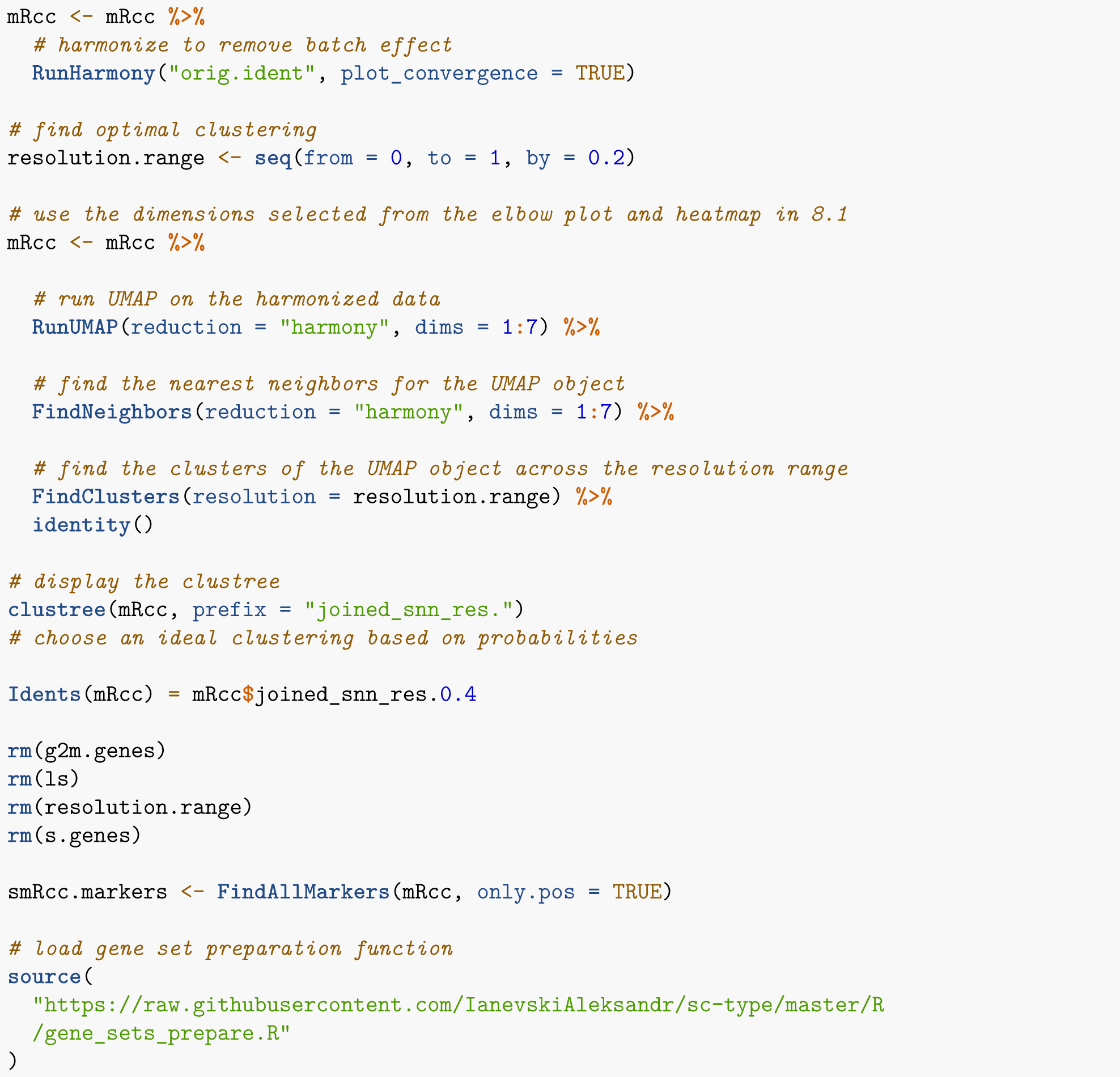

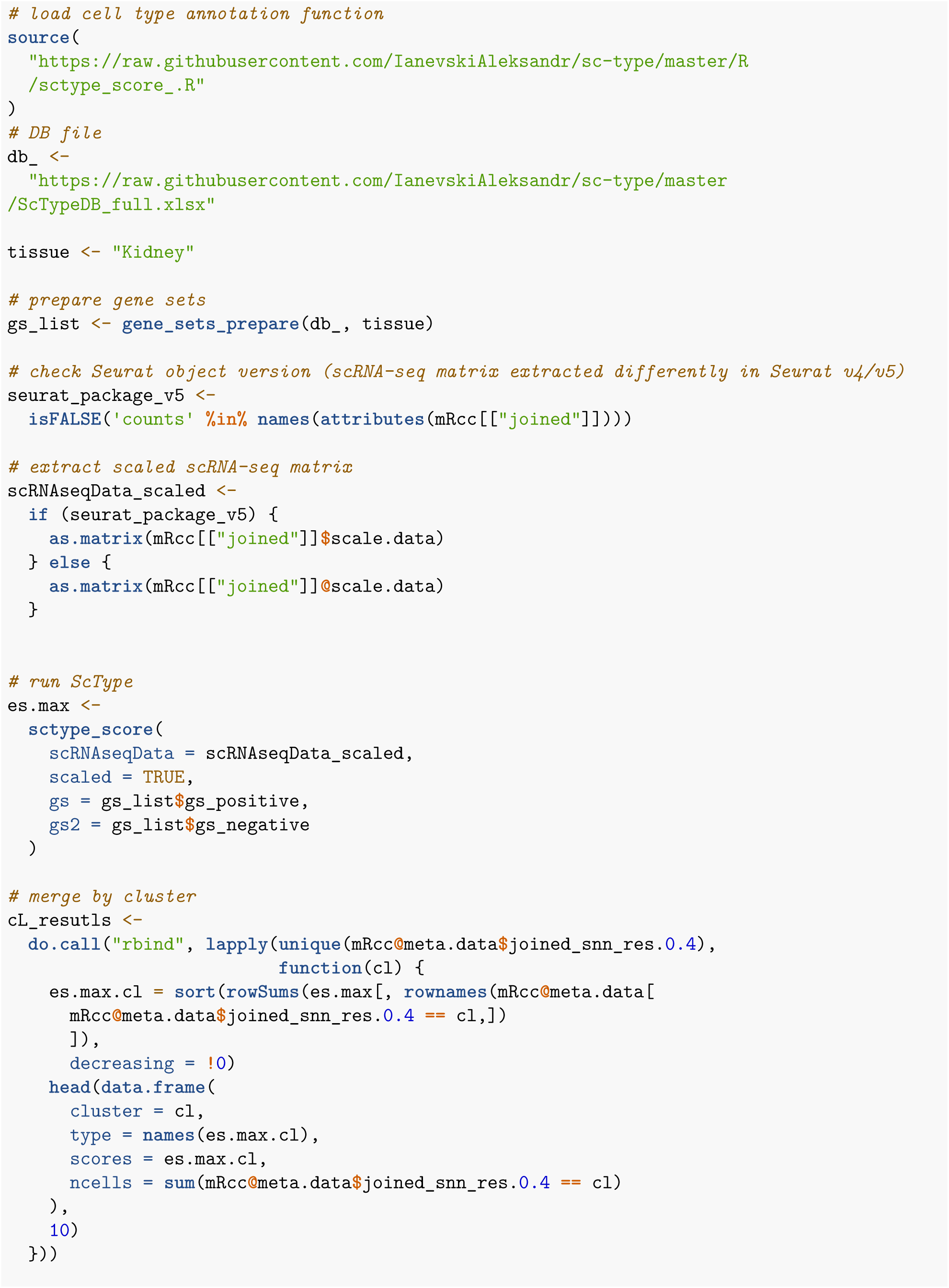

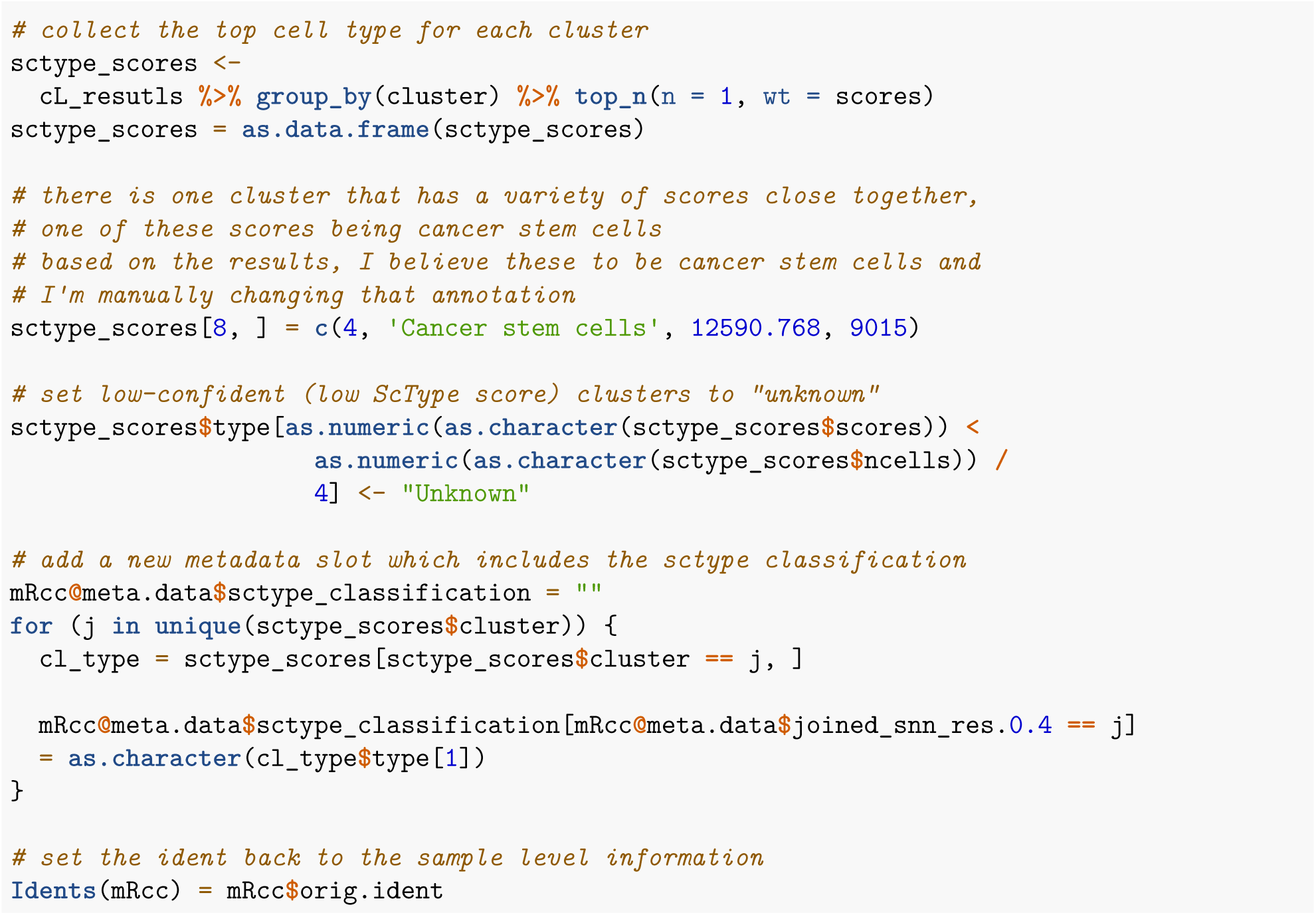

### 8.3 Copy number analysis

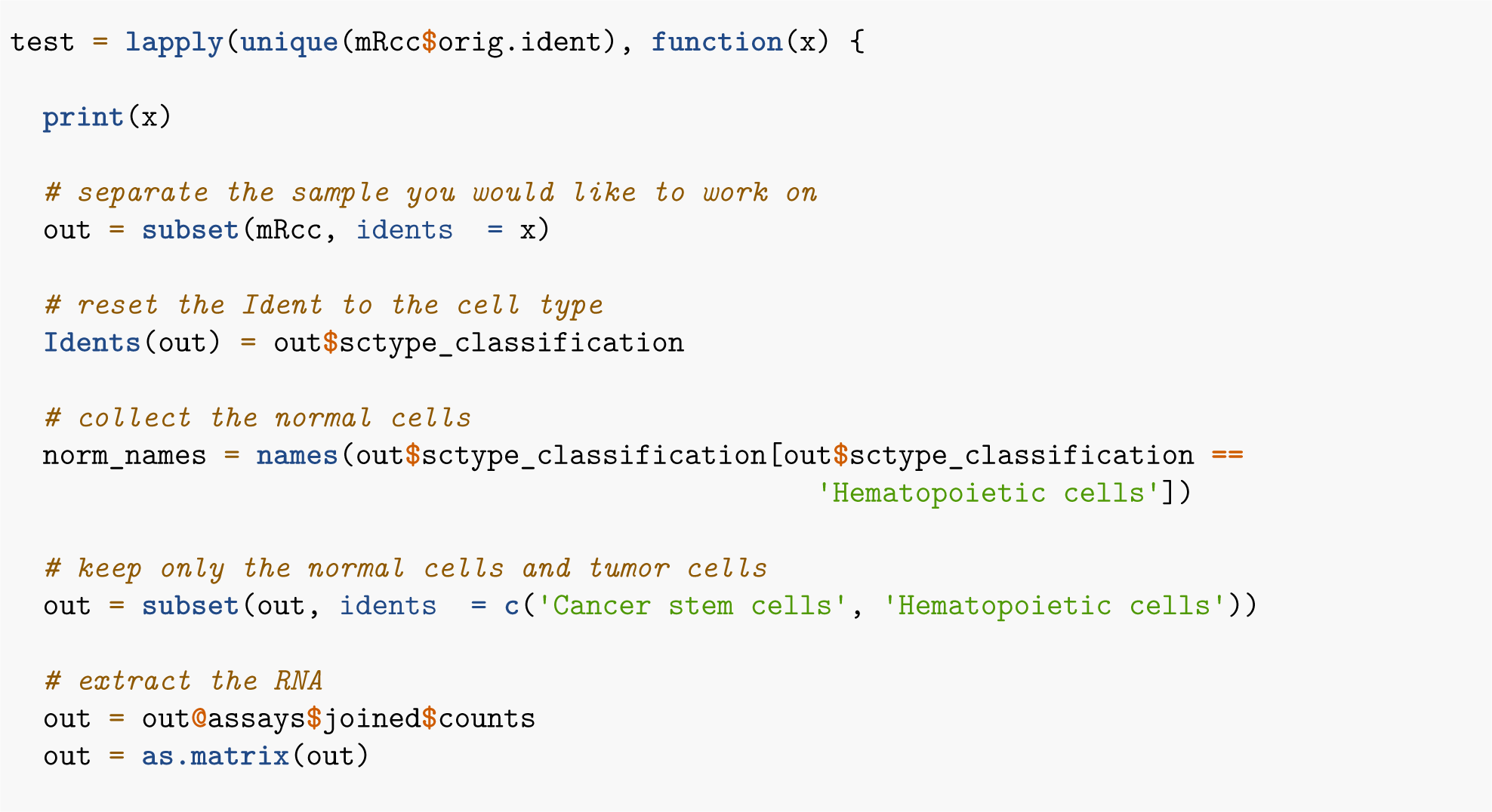

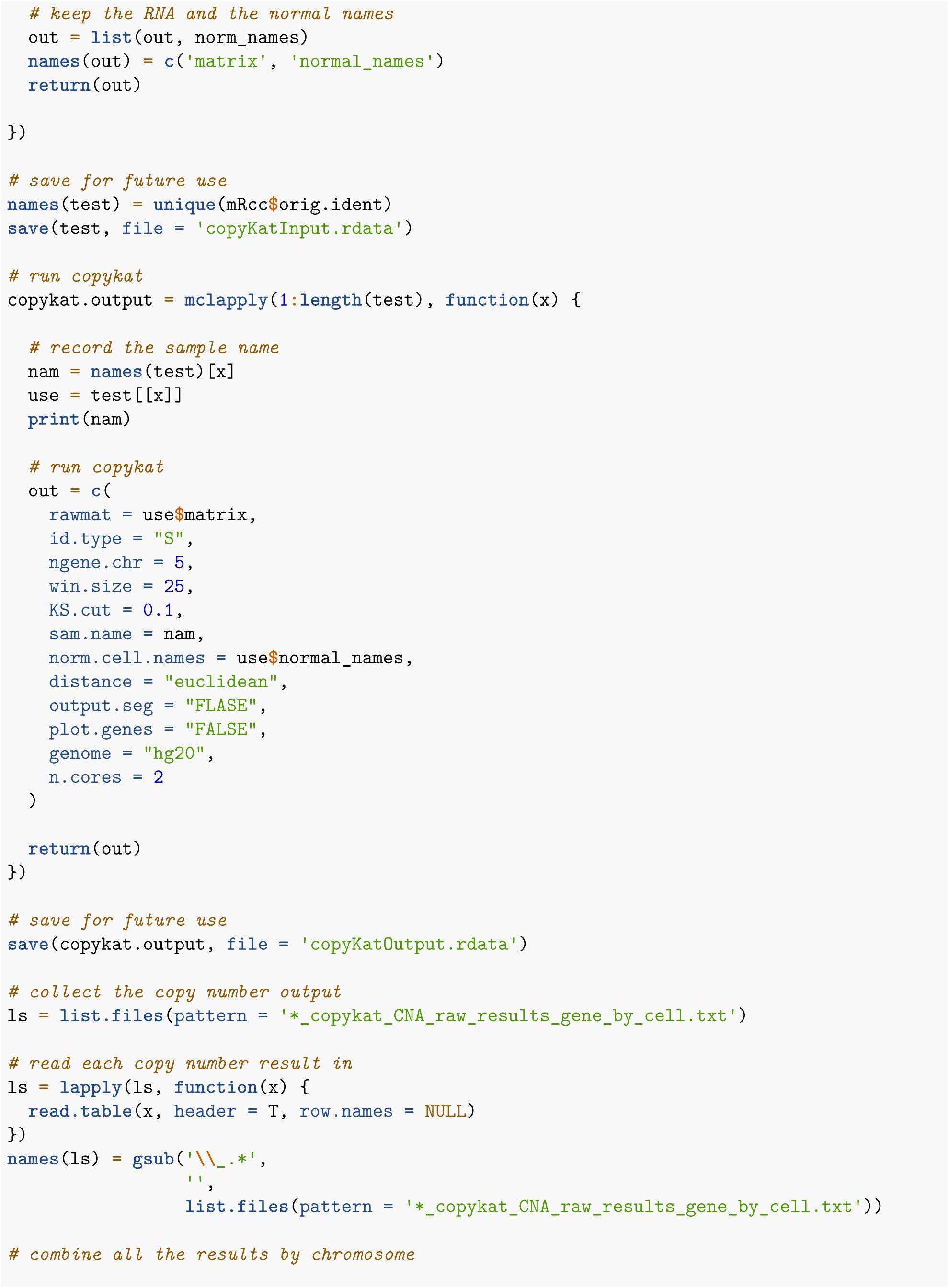

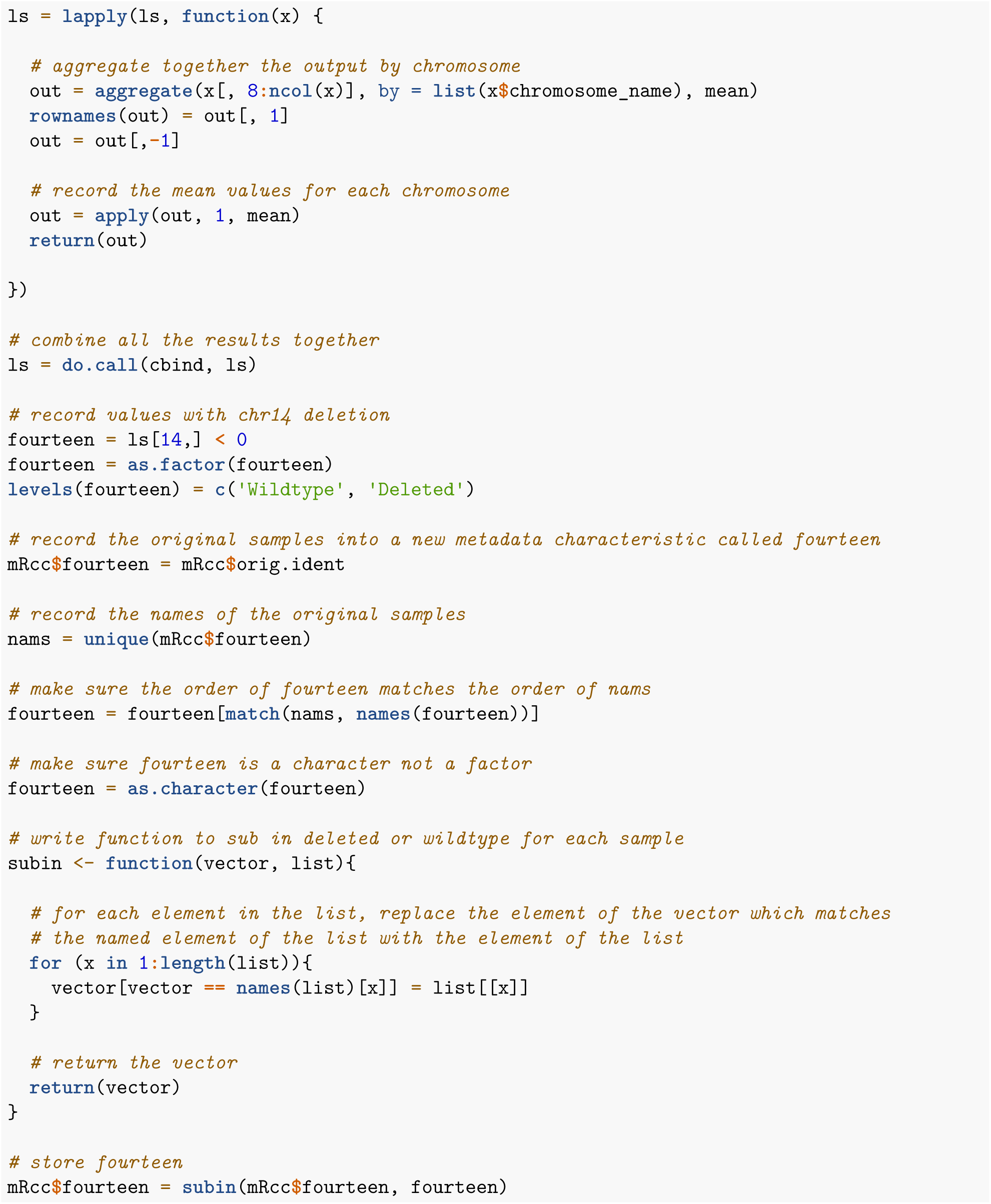

### 8.4: Immune cell analysis

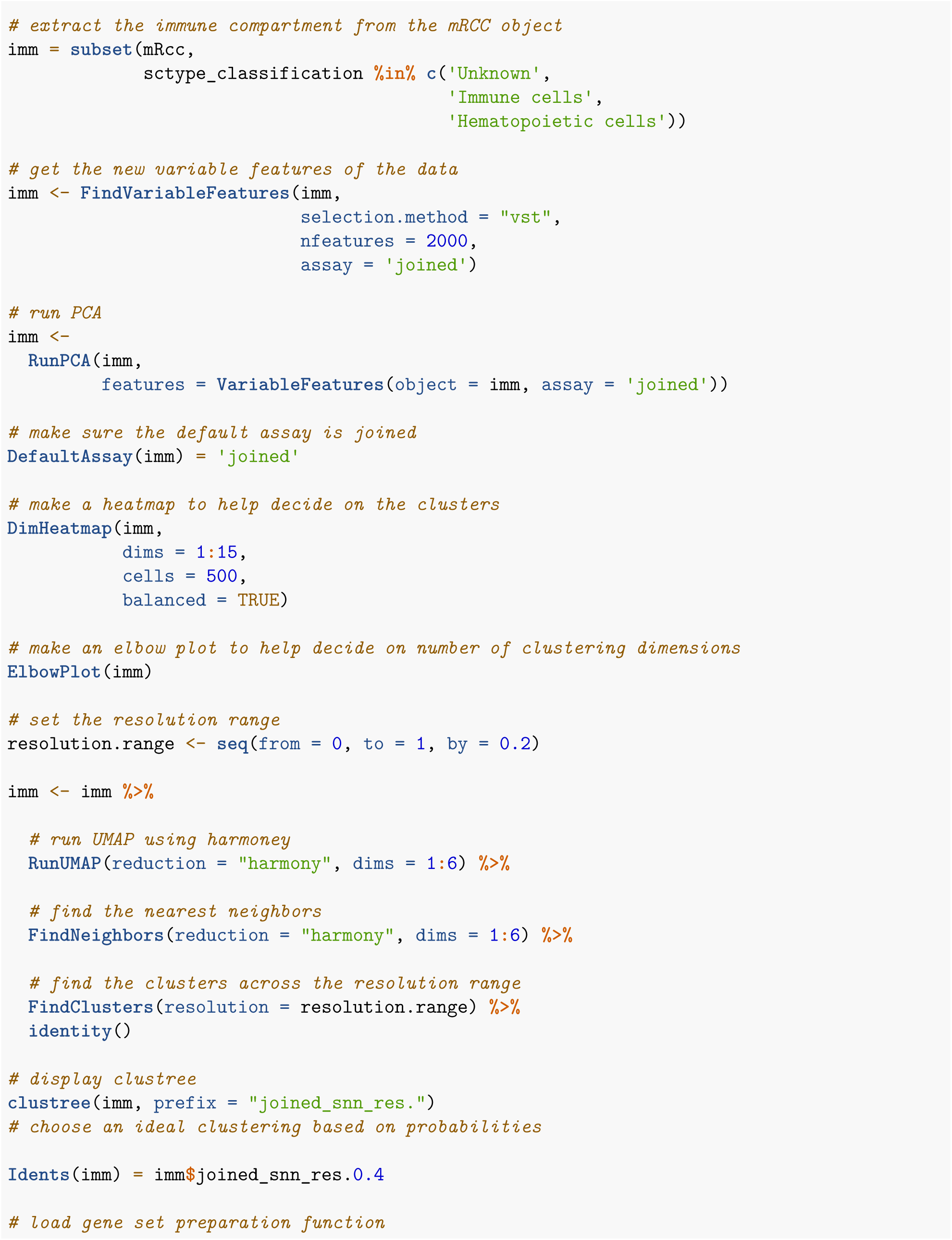

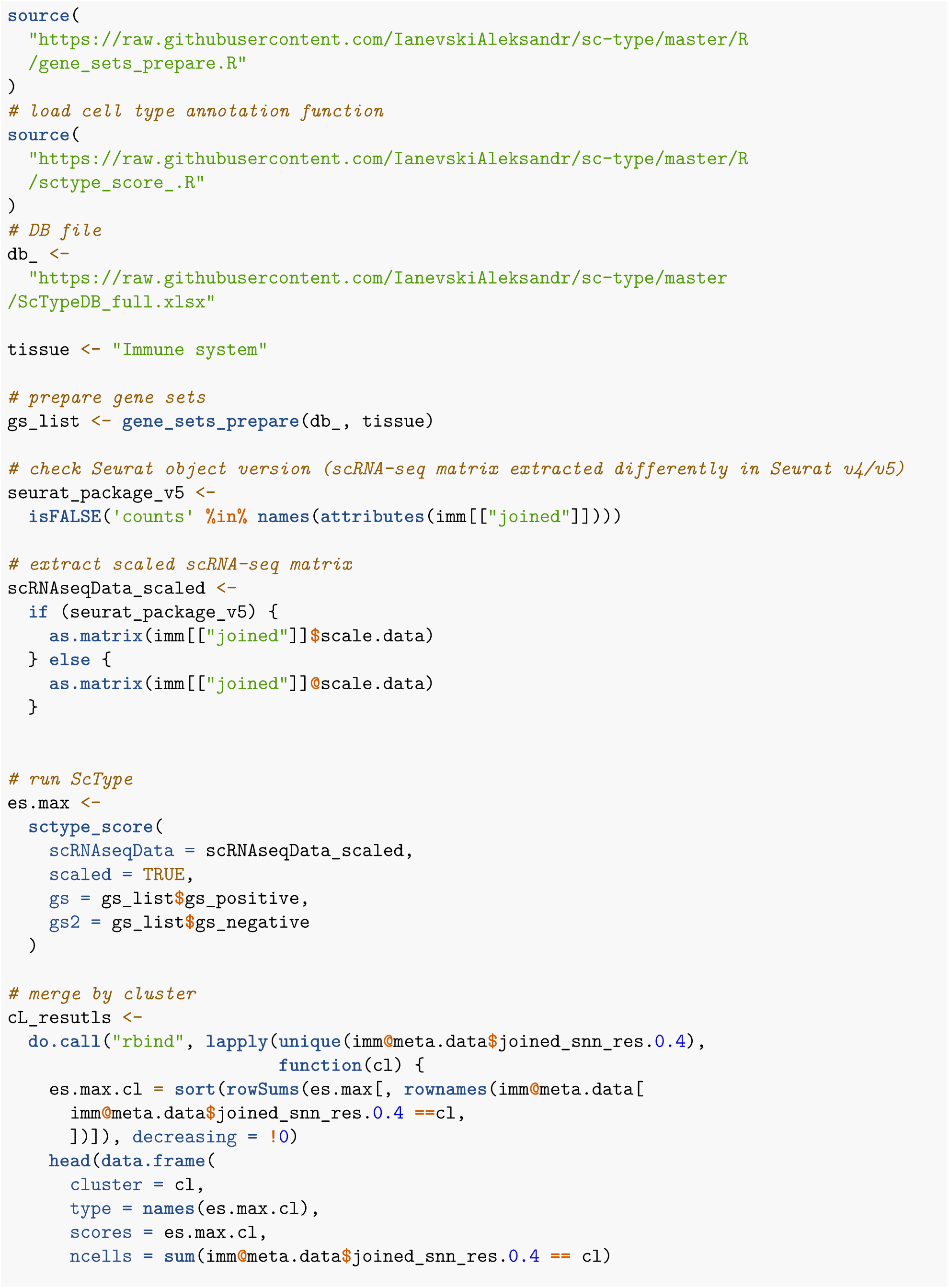

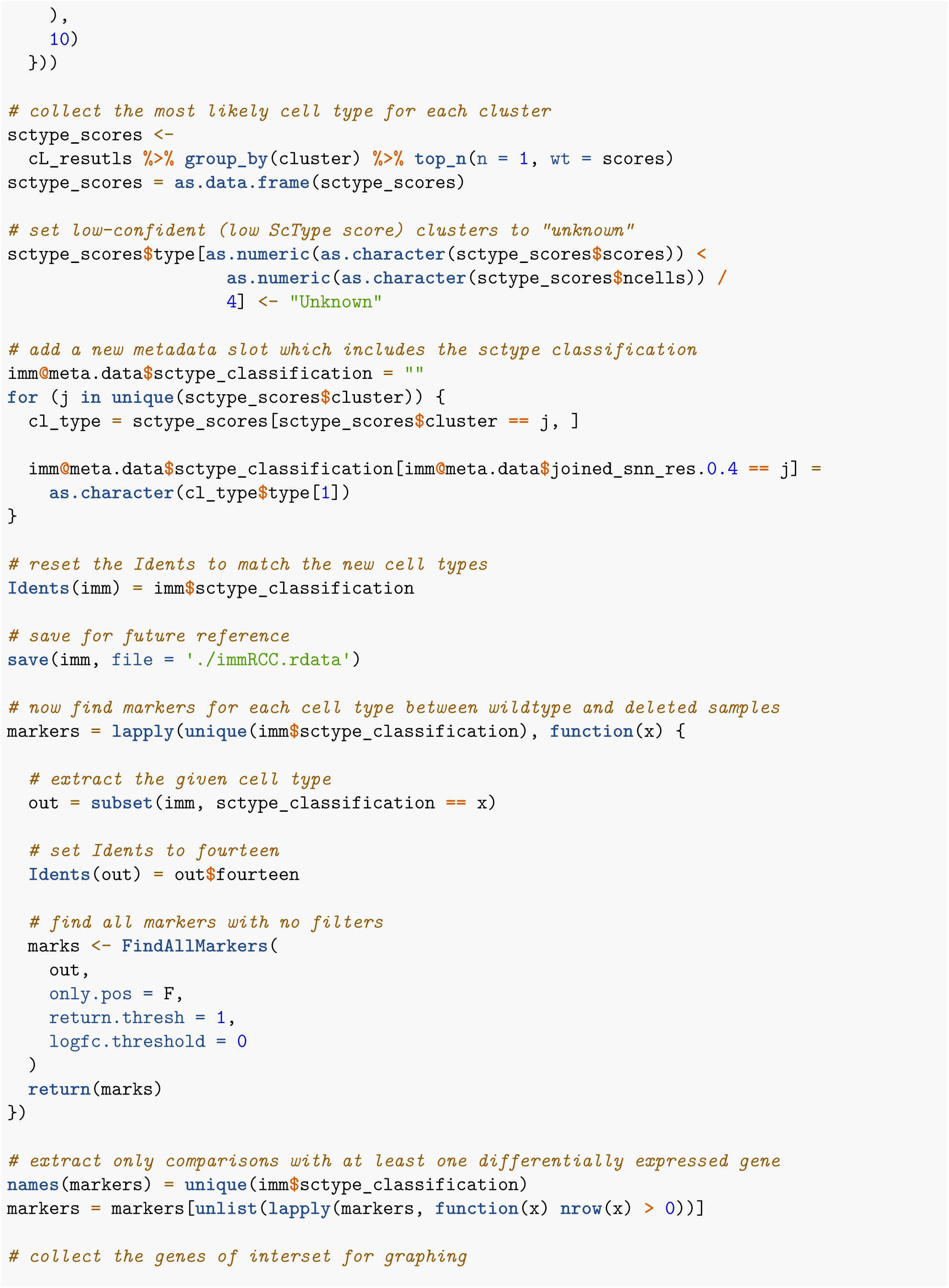

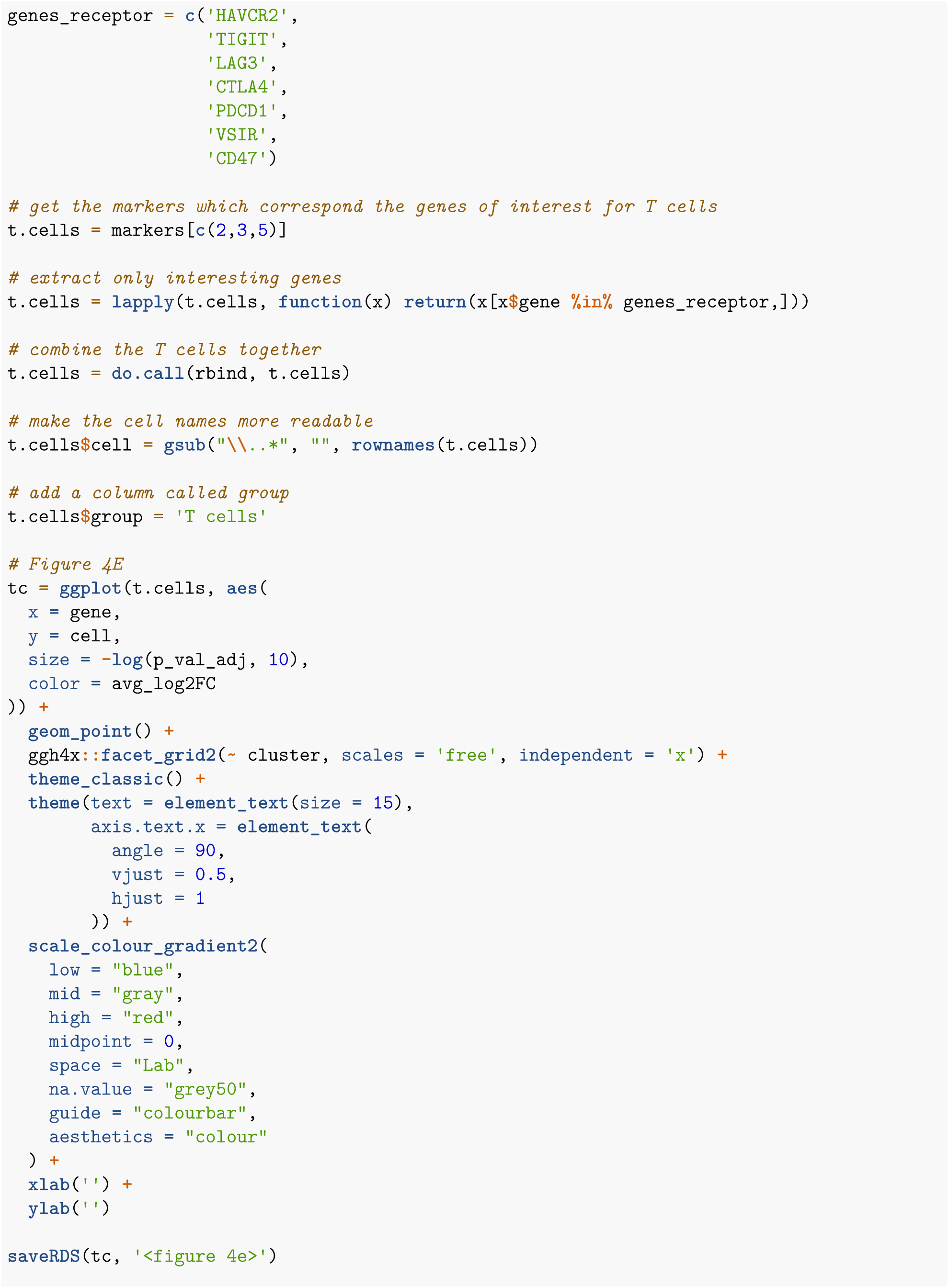

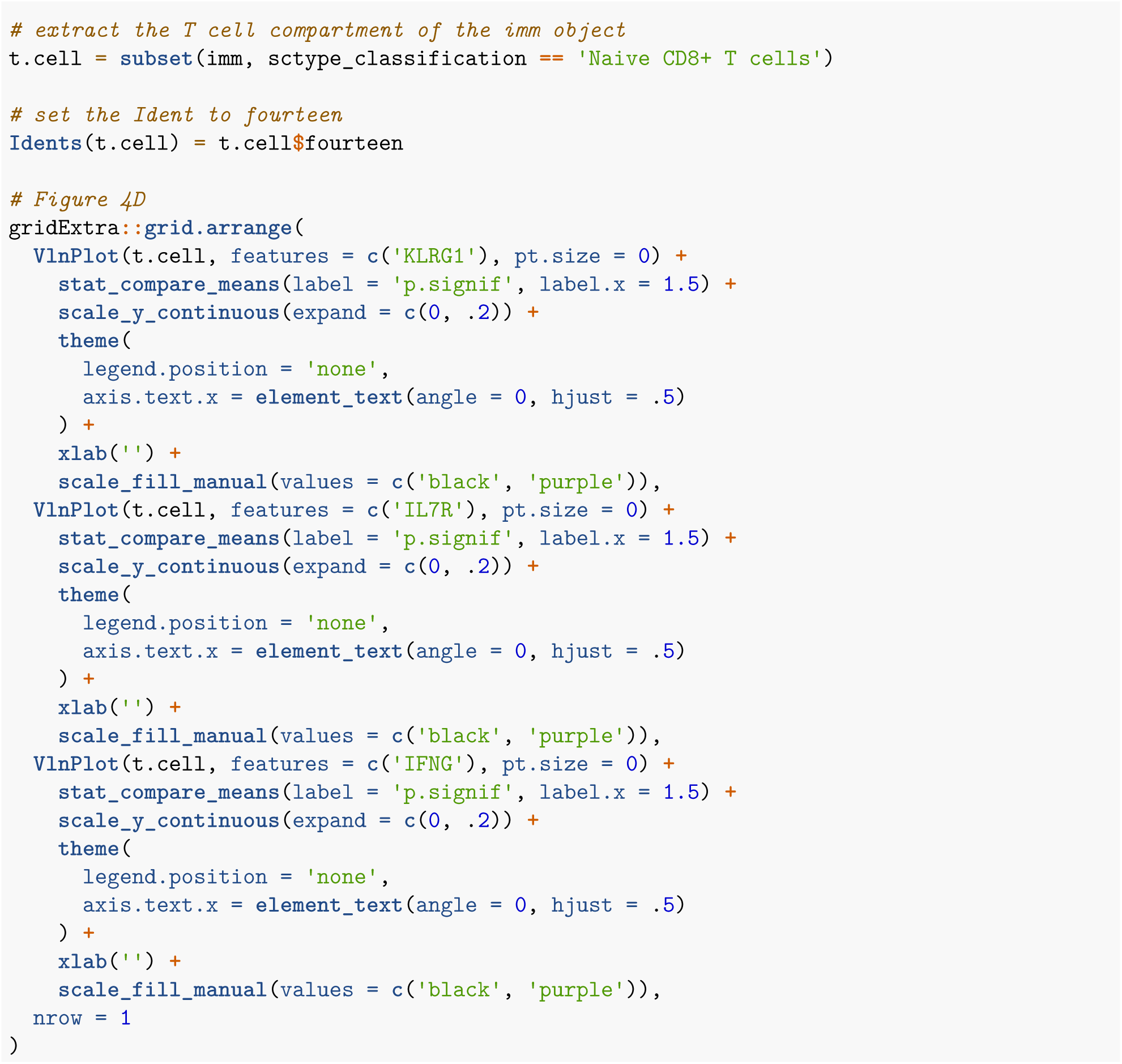

