## Supplementary Data for "RIGATonI: An R software for Rapid Identification of Genomic Alterations in Tumors affecting lymphocyte Infiltration"

### Table of Contents

**Supplemental Figure 1:** Training data cancer type distributions

**Supplemental Figure 2:** Complete summary of mIHC and flow cytometry results

**Supplemental Figure 3:** RIGATonI Website Homepage

**Supplemental Figure 4:** RIGATonI Website Transcriptomics page

**Supplemental Figure 5:** RIGATonI Website Immune page

**Supplemental Table 1:** List of gene names selected for Immunity Module (see in separate .txt)

**Supplemental Table 2:** Confusion matrix of pathologist annotations and RIGATonI Immunity Module predictions

**Supplemental Table 3:** Detailed statistics for RIGATonI's Immunity Module across all classes (low, medium, and high)

**Supplemental Table 4:** Detailed statistics for RIGATonI's Immunity Module within classes (low, medium, and high)

**Supplemental Table 5:** List of genomic alterations in OncoKB genes identified by RIGATonI with corresponding annotations and OncoKB descriptions (see in separate .txt)

**Supplemental Table 6:** Comparison of computational models developed and summary statistics

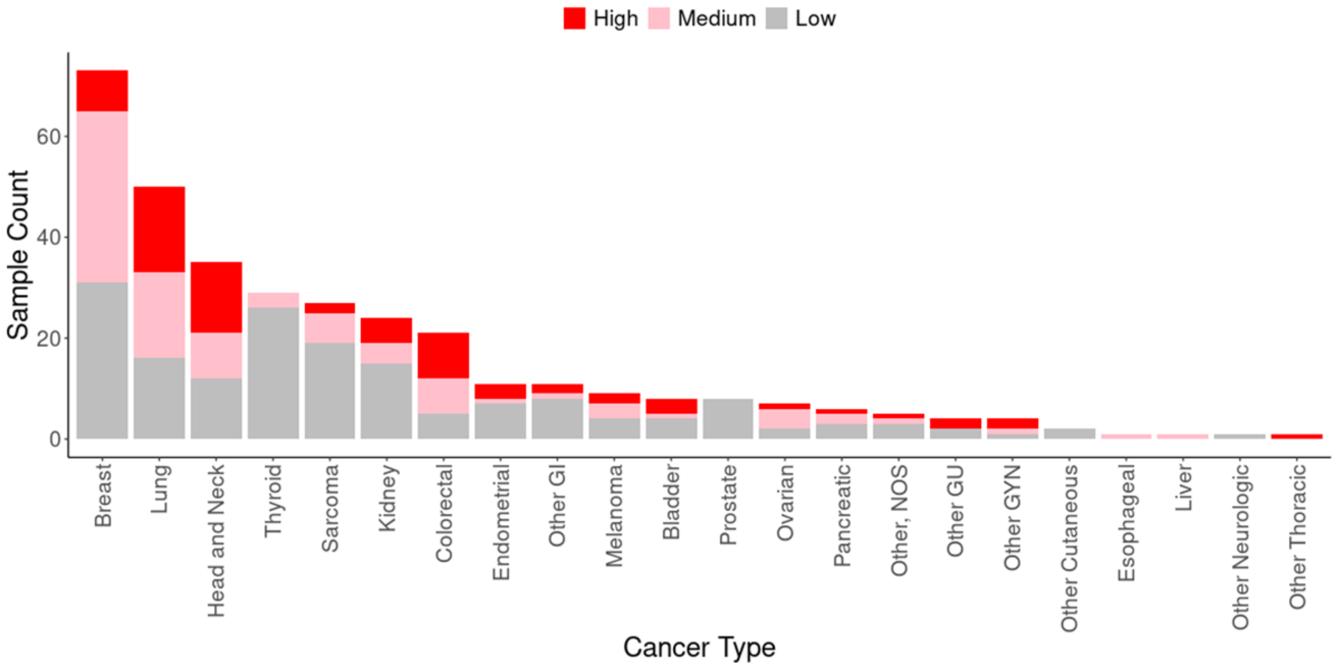


**Supplemental Figure 1: Training data from OSU-ORIEN contains a wide distribution of cancer types and variety of immune phenotypes.** 22 Cancer types from OSU-ORIEN and 403 unique tumors were included in the training data for the Immunity Module. Every effort was made to include tumors with high (red), medium (pink), and low (gray) degrees of immune infiltration from each cancer type. Cancer types are present in proportion with their availability in the OSU-ORIEN dataset.

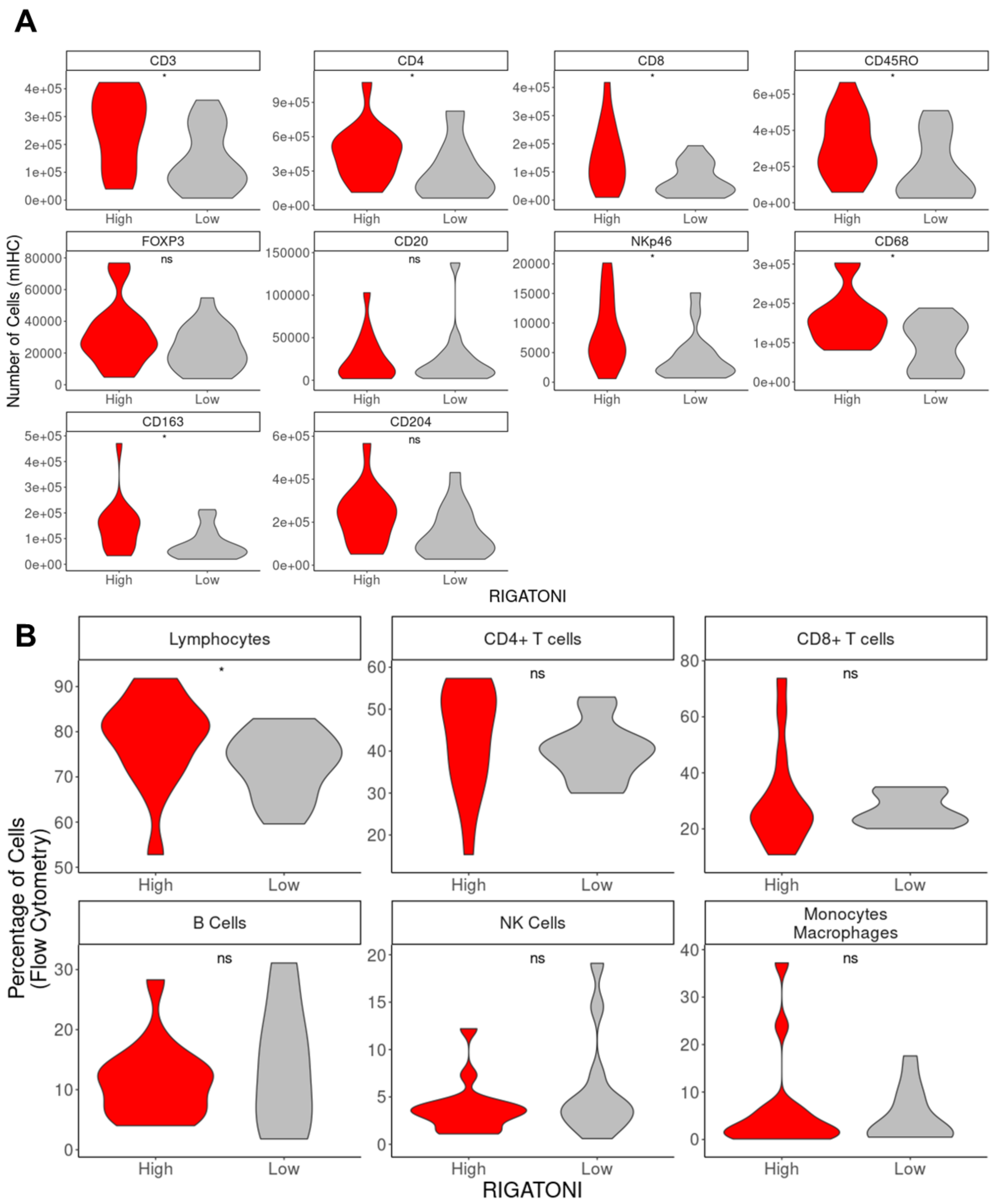


**Supplemental Figure 2: Multiplexed immunohistochemistry and flow cytometry from Saito *et. al.* 2022 reveals a distinct difference between RIGATonI-high and RIGATonI-low samples. A.** Multiplex immunohistochemistry results from 32 gastric cancer tumors. Ten immune markers were assessed for RIGATonI immune phenotypes. RIGATonI-high samples consistently demonstrate increased cells within many of the markers studied. **B.** Flow cytometry results from the same study. Six immune subsets by flow cytometry were assessed. Lymphocyte proportion is significantly increased in RIGATonI-high samples compared to low. No other significant differences are noted. **Significance values: p <=0 .05: *, p <=0.01: **, p <=.001: ***, p <= 0.0001: ******

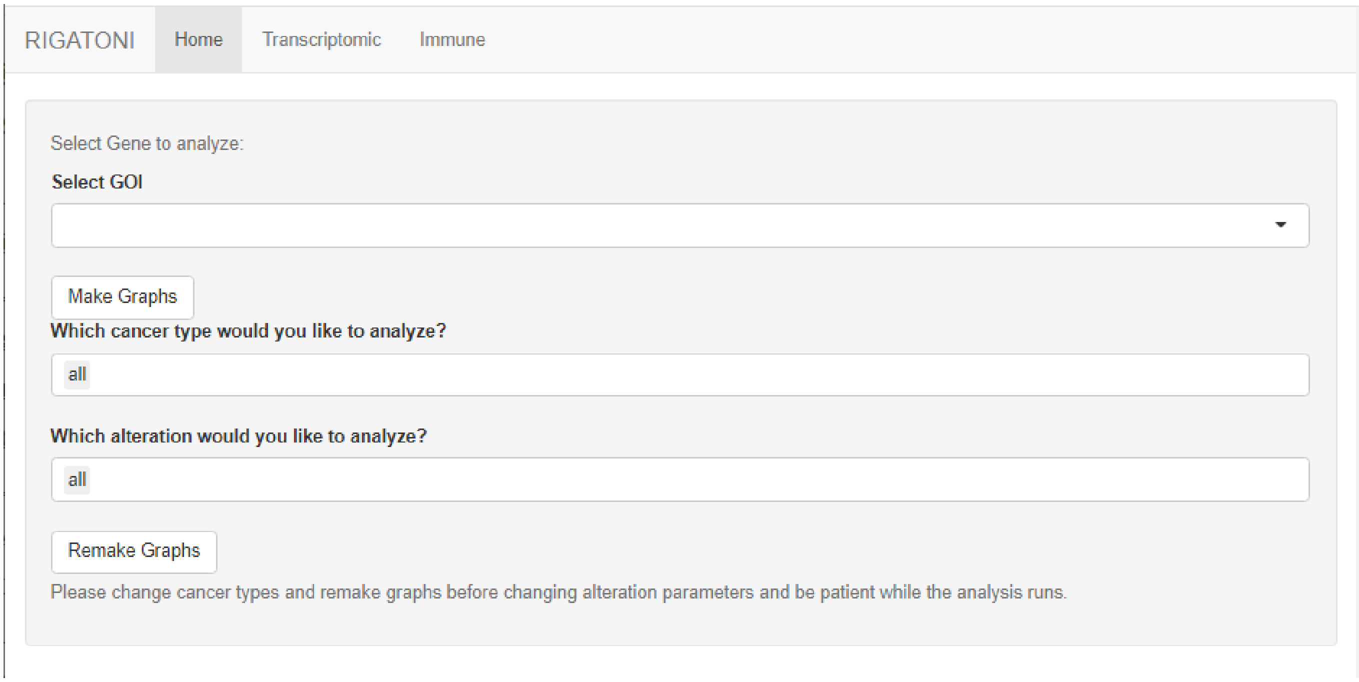


**Supplemental Figure 3: RIGATonI homepage screenshot.** Users select a gene of interest by typing into the drop down menu and selecting their gene. If the gene they are looking for does not appear, that indicates there were no function and immune affecting genomic variants uncovered in TCGA. By default, users will select all genomic alterations and all cancer types to analyze, but after making graphs can filter with their prefered options.

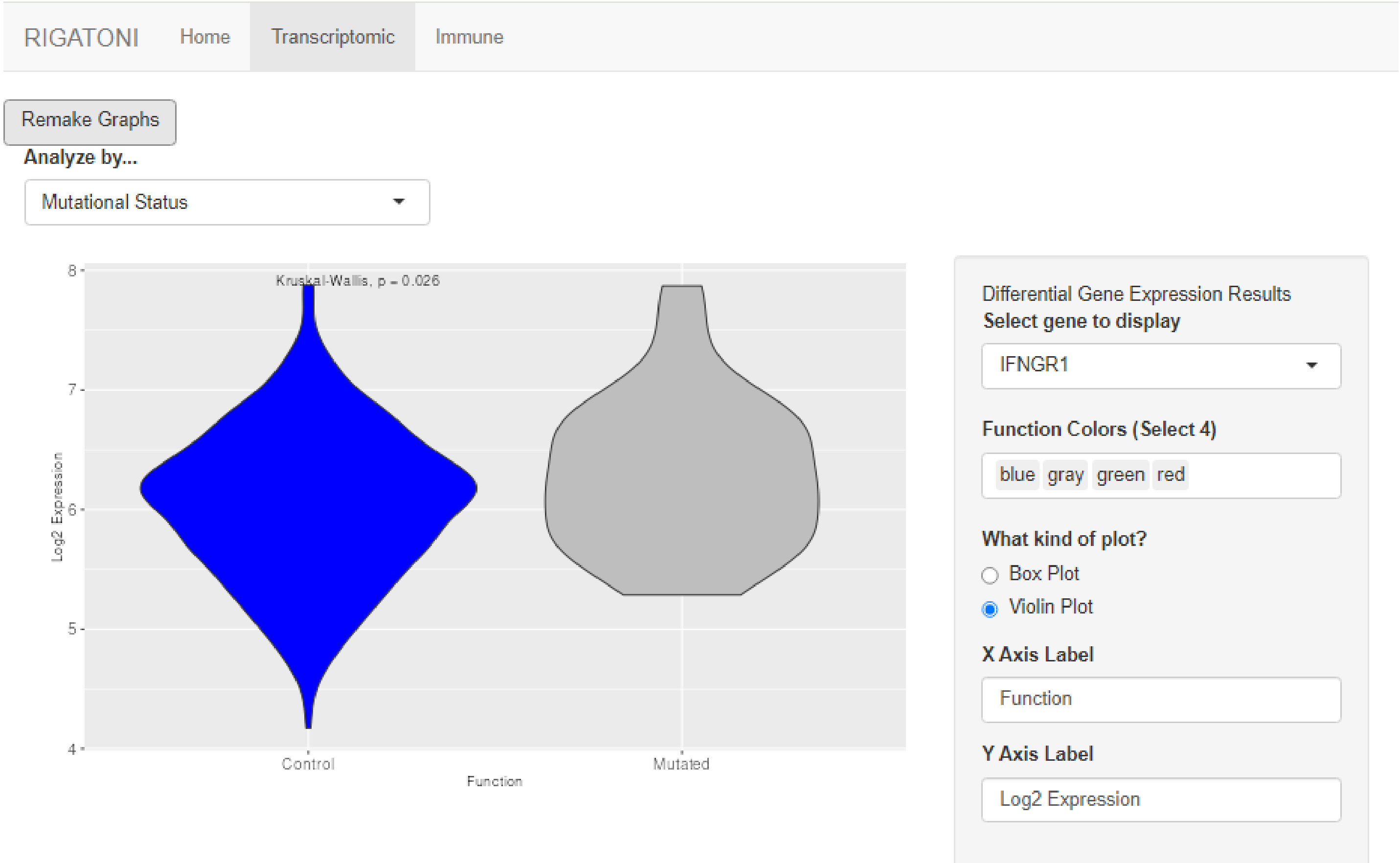


**Supplemental Figure 4: RIGATonI transcriptomic page screenshot.** Users select a gene of interest by typing into the drop down menu and selecting their gene. They can choose to display results by either mutation status, functional status, or immune state using the drop down in the top left. Users can switch between box plots and violin plots, change axis labels, and change colors using the panel to the right.

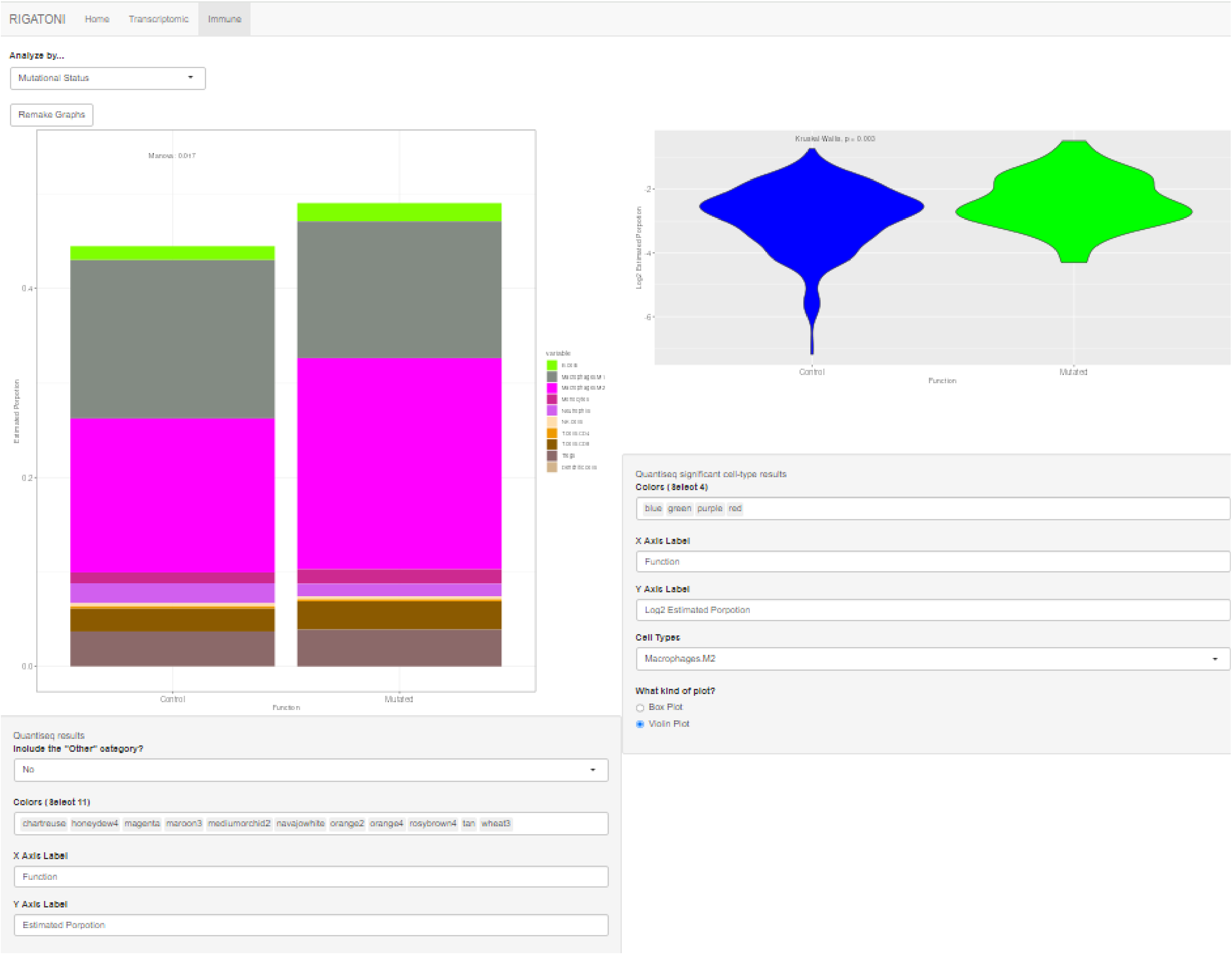


**Supplemental Figure 5: RIGATonI immune page screenshot.** Users can choose to display results by either mutation status, functional status, or immune state using the drop down in the top left. Users can switch between box plots and violin plots, change axis labels, and change colors using the panel beneath each image. On the left we see the complete immune microenvironment visualized in a bar plot. On the right box plots or violin plots of specific cell types can be displayed. Users can change the cell type of interest using the drop down menu beneath the figure.

**Supplemental Table 2: Confusion matrix of pathologist annotations and RIGATonI Immunity Module predictions.** 69 samples from the OSU-ORIEN training data were reserved for testing. Pathologist annotations are shown in column, and RIGATonI predictions are shown across rows.

| | Pathologist Annotations | | |
|---|---|---|---|
| RIGATonI Predictions | Low | Medium | High |
| Low | 30 | 6 | 0 |
| Medium | 5 | 8 | 4 |
| High | 1 | 4 | 11 |

**Supplemental Table 3: Detailed statistics for RIGATonI's Immunity module across all classes (low, medium, and high)**. 69 samples from the OSU-ORIEN training data were reserved for testing. The model was applied to these samples using pathologist annotations as a ground truth for calculation of statistics.

| Overall Statistics | All Classes |
|---|---|
| Accuracy | 0.7101 |
| 95% CI | (0.5884, 0.8131) |
| No Information Rate | 0.5217 |
| P-Value [Acc > NIR] | 0.001111 |
| Kappa | 0.5272 |
| Mcnemar's Test P-Value | 0.7792 |

**Supplemental Table 4: Detailed statistics for RIGATonI's Immunity module within classes (low, medium, and high)**. 69 samples from the OSU-ORIEN training data were reserved for testing. The model was applied to these samples using pathologist annotations as a ground truth for calculation of statistics.

| Statistics by Class: | Low | Medium | High |
|---|---|---|---|
| Sensitivity | 0.8333 | 0.4444 | 0.7333 |
| Specificity | 0.8182 | 0.8235 | 0.9074 |
| Pos Pred Value | 0.8333 | 0.4706 | 0.6875 |
| Neg Pred Value | 0.8182 | 0.8077 | 0.9245 |
| Prevalence | 0.5217 | 0.2609 | 0.2174 |
| Detection Rate | 0.4348 | 0.1159 | 0.1594 |
| Detection Prevalence | 0.5217 | 0.2464 | 0.2319 |
| Balanced Accuracy | 0.8258 | 0.634 | 0.8204 |

**Supplemental Table 6: Accuracy of possible RIGATonI models on testing data.** To build RIGATonI's Immunity Module, six different machine learning algorithms were considered using different machine learning methods, number of predictors, and pathologist annotations. Ultimately, model 6 was selected because of its high accuracy in the 69 samples from the OSU-ORIEN training data were reserved for testing.

| Training data | Method | Number of Predictors | AUC High | AUC Med | AUC Low |
|---|---|---|---|---|---|
| Path Combo | Ordinal | 121 | 54% | 70% | 74.24% |
| Path Combo | XgBoost | 121 | 62% | 66.06% | 74% |
| Pathologist 1 | Ordinal | 133 | 68.22% | 58% | 78.41% |
| Pathologist 1 | XgBoost | 133 | 50% | 50% | 50% |
| Pathologist 2 | Ordinal | 114 | 58% | 50% | 53.90% |
| Pathologist 2 | XgBoost | 114 | 82.04% | 63.40% | 82.58% |
